# Expanding Parkinson’s disease genetics: novel risk loci, genomic context, causal insights and heritable risk

**DOI:** 10.1101/388165

**Authors:** Mike A. Nalls, Cornelis Blauwendraat, Costanza L. Vallerga, Karl Heilbron, Sara Bandres-Ciga, Diana Chang, Manuela Tan, Demis A. Kia, Alastair J. Noyce, Angli Xue, Jose Bras, Emily Young, Rainer von Coelln, Javier Simón-Sánchez, Claudia Schulte, Manu Sharma, Lynne Krohn, Lasse Pihlstrom, Ari Siitonen, Hirotaka Iwaki, Hampton Leonard, Faraz Faghri, J. Raphael Gibbs, Dena G. Hernandez, Sonja W. Scholz, Juan A. Botia, Maria Martinez, Jean-Christophe Corvol, Suzanne Lesage, Joseph Jankovic, Lisa M. Shulman, The 23andMe Research Team, System Genomics of Parkinson’s Disease (SGPD) Consortium, Margaret Sutherland, Pentti Tienari, Kari Majamaa, Mathias Toft, Ole A. Andreassen, Tushar Bangale, Alexis Brice, Jian Yang, Ziv Gan-Or, Thomas Gasser, Peter Heutink, Joshua M Shulman, Nicolas Wood, David A. Hinds, John A. Hardy, Huw R Morris, Jacob Gratten, Peter M. Visscher, Robert R. Graham, Andrew B. Singleton, for the International Parkinson’s Disease Genomics Consortium

## Abstract

We performed the largest genome-wide association study of PD to date, involving the analysis of 7.8M SNPs in 37.7K cases, 18.6K UK Biobank proxy-cases, and 1.4M controls. We identified 90 independent genome-wide significant signals across 78 loci, including 38 independent risk signals in 37 novel loci. These variants explained 26-36% of the heritable risk of PD. Tests of causality within a Mendelian randomization framework identified putatively causal genes for 70 risk signals. Tissue expression enrichment analysis suggested that signatures of PD loci were heavily brain-enriched, consistent with specific neuronal cell types being implicated from single cell expression data. We found significant genetic correlations with brain volumes, smoking status, and educational attainment. In sum, these data provide the most comprehensive understanding of the genetic architecture of PD to date by revealing many additional PD risk loci, providing a biological context for these risk factors, and demonstrating that a considerable genetic component of this disease remains unidentified.

## INTRODUCTION

Parkinson’s disease (PD) is a neurodegenerative disorder, affecting up to 2% of the population older than 60 years, an estimated 1 million individuals in the United States alone. PD patients suffer from a combination of progressive motor and non-motor symptoms that increasingly impair daily function and quality of life. There are no treatments that delay or alter PD^1^. As the global population continues to age, the prevalence of PD is projected to double in some age groups by 2030, creating a substantial burden on healthcare systems.^1, 2, 3^

Early investigations into the role of genetic factors in PD focused on the identification of rare mutations underlying familial forms of the disease,^4–6^ but over the past decade there has been a growing appreciation for the important contribution of genetics in sporadic disease^7, 8^. Genetic studies of sporadic PD have altered the foundational view of disease etiology as much of sporadic disease was formerly thought to be environmental.

With this in mind, we executed a series of experiments to further explore the genetics of PD (summarized in Figure 1). We performed the largest-to-date GWAS for PD, including 7.8M SNPs, 37.7K cases, 18.6K UK Biobank (UKB) “proxy-cases” and 1.4M controls. We identified putatively causal genes for PD, providing valuable targets for therapeutic research. We assessed the function of these putatively causal genes on a larger scale than in previous studies of PD via Mendelian randomization (MR), expression enrichment, and protein-protein interaction network analysis ^9, 10, 11^. We estimated PD heritability, developed a polygenic risk score that predicted a substantial proportion of this heritability, and leveraged these results to inform future studies of PD genetics. Finally, we identified putative PD biomarkers and risk factors using genetic correlation and Mendelian randomization.

**Figure 1:**
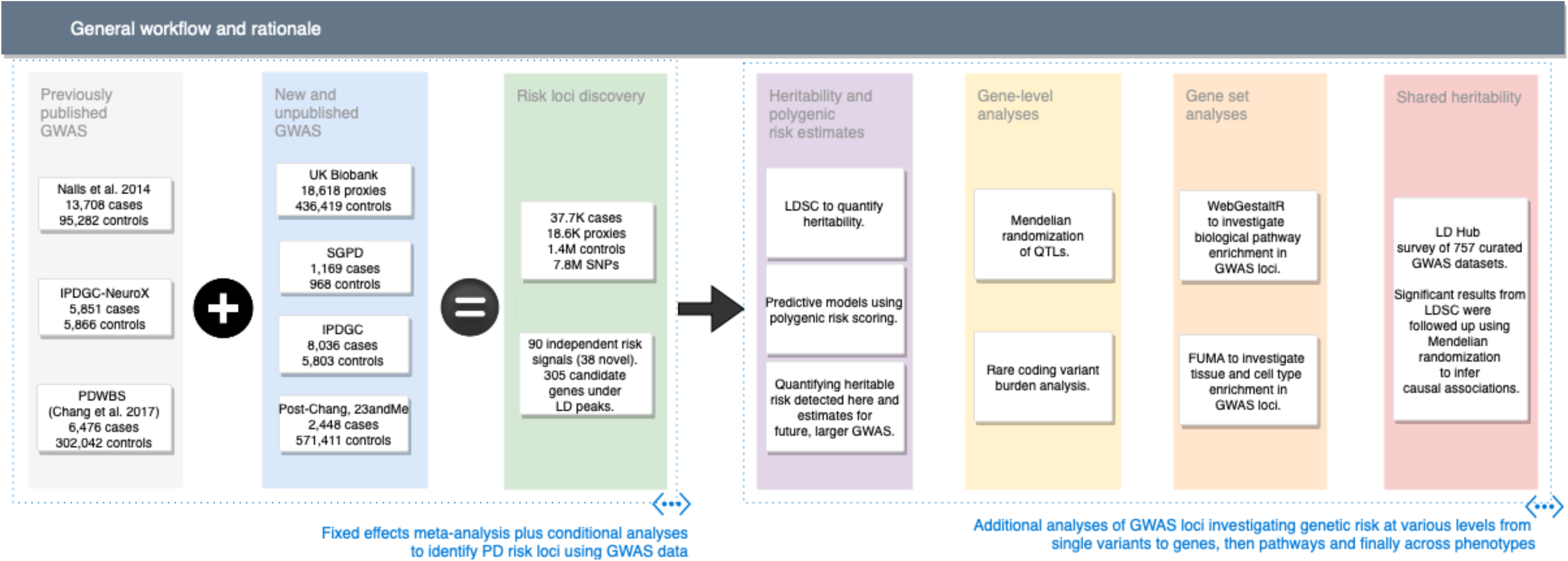
Workflow and rationale summary. This figure describes study design and rationale behind the analyses included in this report.

## METHODS

See Supplementary Methods

## RESULTS

### Novel loci and multiple signals in known loci identified

To maximize our power for locus discovery we used a single stage design, meta-analyzing all available GWAS summary statistics. In support of this design, we found strong genetic correlations between GWAS using PD cases ascertained by clinicians compared to 23andMe self-reported cases (rG = 0.85, SE = 0.06) and UKB proxy cases (rG = 0.84, SE = 0.134).

We identified a total of 90 independent genome-wide significant association signals through our meta-analysis and conditional analyses of 37,688 cases, 18,618 UKB proxy-cases and 1,417,791 controls at 7,784,415 SNPs (Figure 2, Table 1, Supplementary Appendices, Table S1, Table S2). Of these, 38 signals are new and more than 1MB from loci described in a previous report by Chang *et al*. 2017 (Table S3).

**Figure 2:**
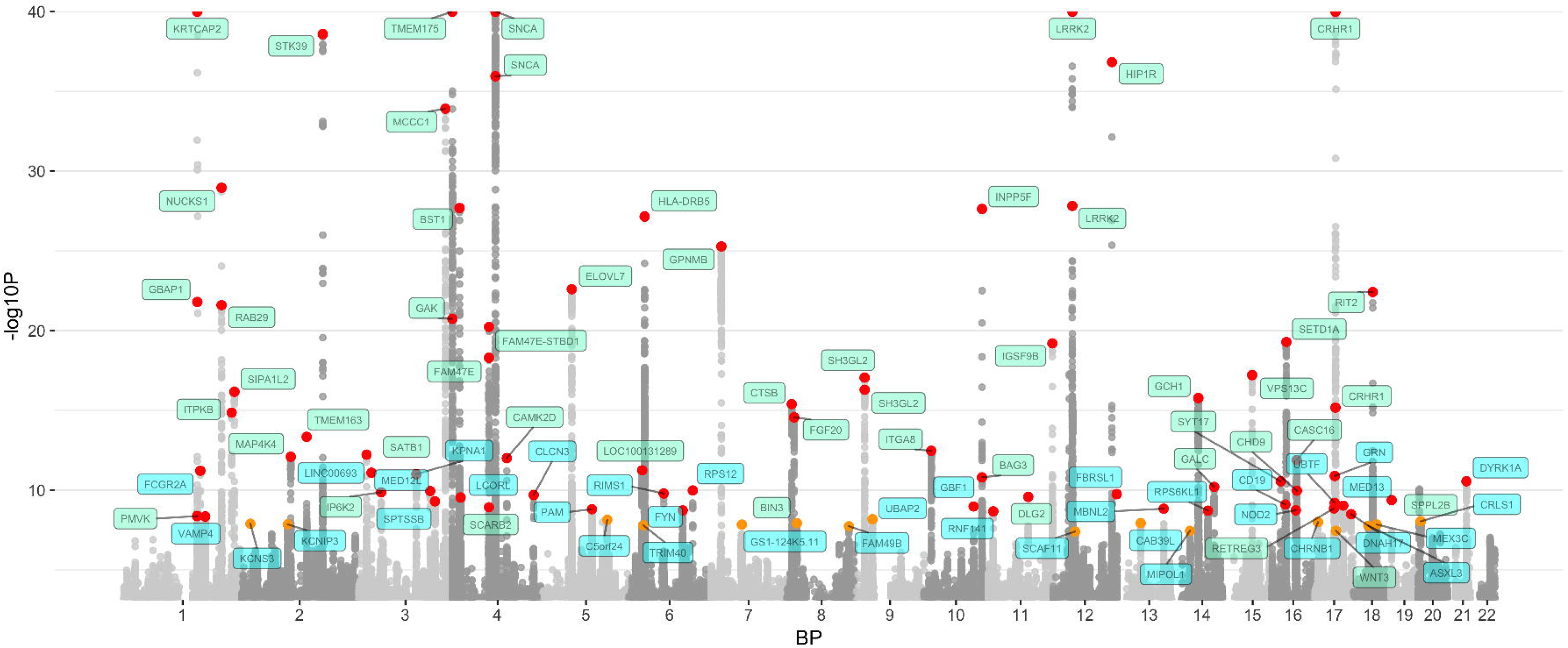
Manhattan plot. The nearest gene to each of the 90 significant variants are labeled in green for previously-identified loci and in blue for novel loci. –log10P values were capped at 40. Variant points are color coded red and orange, with orange representing significant variants at P 5E-08 and 5E-9 and red representing significant variants at P < 5E-9. The X axis represents the base pair position of variants from smallest to largest per chromosome (1-22).

**Table 1:**
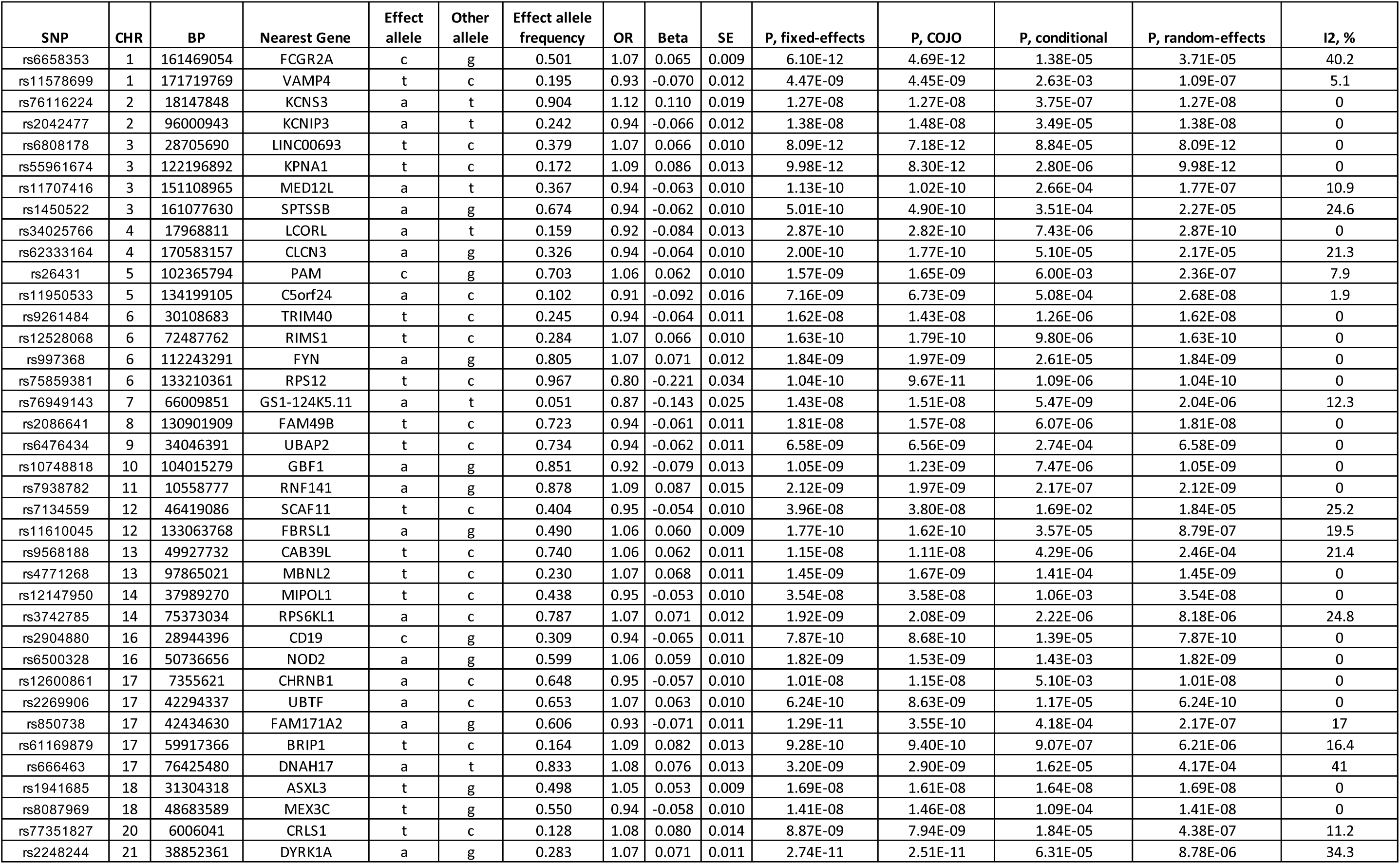
Novel loci associated with Parkinson’s disease. Summary statistics for all 90 genome-wide significant PD variants. Columns include single nucleotide polymorphism ID (SNP), chromosome (CHR), base pair position (BP), nearest gene annotation for the variant, effect allele designation and frequency, as well as metrics for the odds ratio (OR), regression coefficient (beta), and standard error of the beta for the SNP from fixed-effects meta-analysis as well as the index of heterogeneity (I2). We also include four p-values from: fixed-effects meta-analyses, random-effects meta-analyses, standard conditional analyses in 23andMe, and a conditional joint analysis approach (COJO).

In an attempt to detect multiple independent signals within loci we implemented conditional and joint analysis (GCTA-COJO, http://cnsgenomics.com/software/gcta/) with a large study-specific reference genotype series, as well as a participant-level conditional analysis using 23andMe data ^12^. We considered independent risk signals from conditional analyses to share the same locus if they were within 250kb of each other. We detected 10 loci containing more than one independent risk signal (22 risk SNPs in total across these loci), of which nine had been identified by previous GWAS, including multi-signal loci in the vicinity of *GBA, NUCKS1 / RAB29, GAK / TMEM175, SNCA* and *LRRK2*. The novel multi-signal locus comprised independent risk variants rs2269906 (*UBTF* / *GRN*) and rs850738 (*FAM171A2*). Detailed summary statistics on all nominated loci can be found in Table S2.

### Refining heritability estimates and determining extant genetic risk

To quantify how much of the genetic liability we have explained and what direction to take with future PD GWAS we calculated updated heritability estimates and polygenic risk scores (PRS). Using LD score regression (LDSC) on a meta-analysis of all 11 clinically-ascertained datasets from our GWAS and estimated the liability-scale narrow-sense heritability of PD as 0.22 (95% CI 0.18 - 0.26), only slightly lower than a previous estimate derived using GCTA (0.27, 95% CI 0.17 - 0.38)^10, 13, 14^. This may be because LDSC is known to be more conservative than GCTA, however, our LDSC heritability estimate does fall within the 95% confidence interval of the GCTA estimate.

Next, we sought to determine the proportion of SNP-based heritability explained by our PD GWAS results using polygenic risk scores (PRSs). We utilized a two-stage design for our PRS analyses, with variant selection and training in the NeuroX-dbGaP dataset (5,851 cases and 5,866 controls) and then validation in the Harvard Biomarker Study (HBS, 527 cases and 472 controls). We focused on the NeuroX-dbGaP and HBS cohort as both of these clinically characterized cohorts were genotyped on the same PD-focused array (NeuroX) and have been used in previous studies of PRSs ^8, 15–18^. In addition, both of these studies directly genotyped larger effect, rare variants within *LRRK2* (rs34637584, G2019S) and *GBA* (rs76763715, N370S) of great interest in previous PRS analyses.

In order to prevent bias, we estimated the effect size of each SNP contributing to the PRS using a meta-analysis of all PD GWAS datasets except NeuroX-dbGAP and HBS. Using permutation testing in the NeuroX-dbGAP training cohort, we found that the optimal *P* threshold for variant inclusion was 1.35E-03, which included 1809 variants. Two PRSs were tested in HBS, one limited to 88 of the 90 genome-wide significant variants (two variants failed to pass quality control in the HBS study), and the other incorporating 1805 variants from the training phase (four variants failed to pass quality control in HBS due to low imputation quality). The 88 variant PRS had an area under the curve (AUC) of 0.651 (95% CI 0.617 - 0.684), while the 1805 variant PRS had an AUC of 0.692 (95% CI 0.660 - 0.725). The AUCs from our 88 variant PRS in both the NeuroX-dbGAP cohort and the HBS cohort were significantly larger than the AUCs in those same cohorts using a published PRS (Chang *et al.* 2017, AUC = 0.624, P < 0.002 from DeLong’s test). Although the HBS cohort was used to discover the 90 PD GWAS risk variants, therefore potentially biasing our 88 variant PRS, all 90 variants remained genome-wide significant in a meta-analysis of all GWAS datasets excluding the HBS study. Extended results for all included studies at all P-value thresholds can be found in the Supplementary Appendix.

Using equations from Wray *et al*. 2010 and our current heritability estimates, the 88 variant PRS explained approximately 16% of the genetic liability of PD assuming a global prevalence of 0.5%^13, 19^. The 1805 variant PRS explained roughly 26% of PD heritability. In a high-risk population with a prevalence of 2%, the 1805 variant PRS explained 36% of PD heritable risk ^13, 19^ (Table S4).

We then attempted to quantify strata of risk in our more inclusive PRS. Compared to individuals with PRS values in the lowest quartile, the PD odds ratio for individuals with PRS values in the highest quartile was 3.74 (95% CI = 3.35 - 4.18) in the NeuroX-dbGaP cohort and 6.25 (95% CI = 4.26 - 9.28) in the HBS cohort (Table 2, Figure 3, Figure S1).

**Table 2:** Summary of genetic predictive model performance. These are estimates of performance for predictive models including single study estimates, estimates from meta-analyses across studies, as well as a two stage design. Here the best P threshold column denotes the filtering value for SNP inclusion to achieve the maximal pseudo (Nagelkerke’s) R2. The odds ration (OR) colum is the exponent of the regression coefficient (beta) from logistic regression of the polygenic risk score (PRS) on case status, with the standard error (SE) representing the precision of these estimates. These same metrics are derived across array types and datasets using random-effects meta-analyses. The area under the curve (AUC) is included as the most common metric for predictive model performance. In the table, * denotes R2 approximation adjusted for an estimated prevalence of 0.5%, equivalent to roughly half of the unadjusted R2 estimates for the PRS. All calculations and reported statistics include only the PRS and no other parameters after adjusting for principal components 1-5, age and sex at variant selection in the NeuroX-dbGaP dataset.

| Study | Max P threshold | pseudo R2<br>from PRS* | OR | Beta | SE | P | OR, highest<br>quartile PRS | 95% CI, highest<br>quartile PRS | N SNPs | N samples | AUC | 95% CI (DeLong) | Sensitivity | Specificity | Positive predictive<br>value (PPV) | Negative predictive<br>value (NPV) | Balanced<br>accuracy |
| --- | --- | --- | --- | --- | --- | --- | --- | --- | --- | --- | --- | --- | --- | --- | --- | --- | --- |
| Training dataset: IPDGC - Neurox | 1.35E-03 | 0.029 | 1.74 | 0.553 | 0.022 | 8.99E-135 | 3.74 | 3.35 - 4.18 | 1809 | 11,243 | 0.640 | 0.630 - 0.650 | 0.569 | 0.632 | 0.591 | 0.611 | 0.601 |
| Test dataset: HBS | 4.00E-02 | 0.054 | 2.03 | 0.709 | 0.072 | 8.28E-23 | 6.25 | 4.26 - 9.28 | 1805 | 999 | 0.692 | 0.660 - 0.725 | 0.628 | 0.686 | 0.691 | 0.623 | 0.657 |

**Figure 3:**
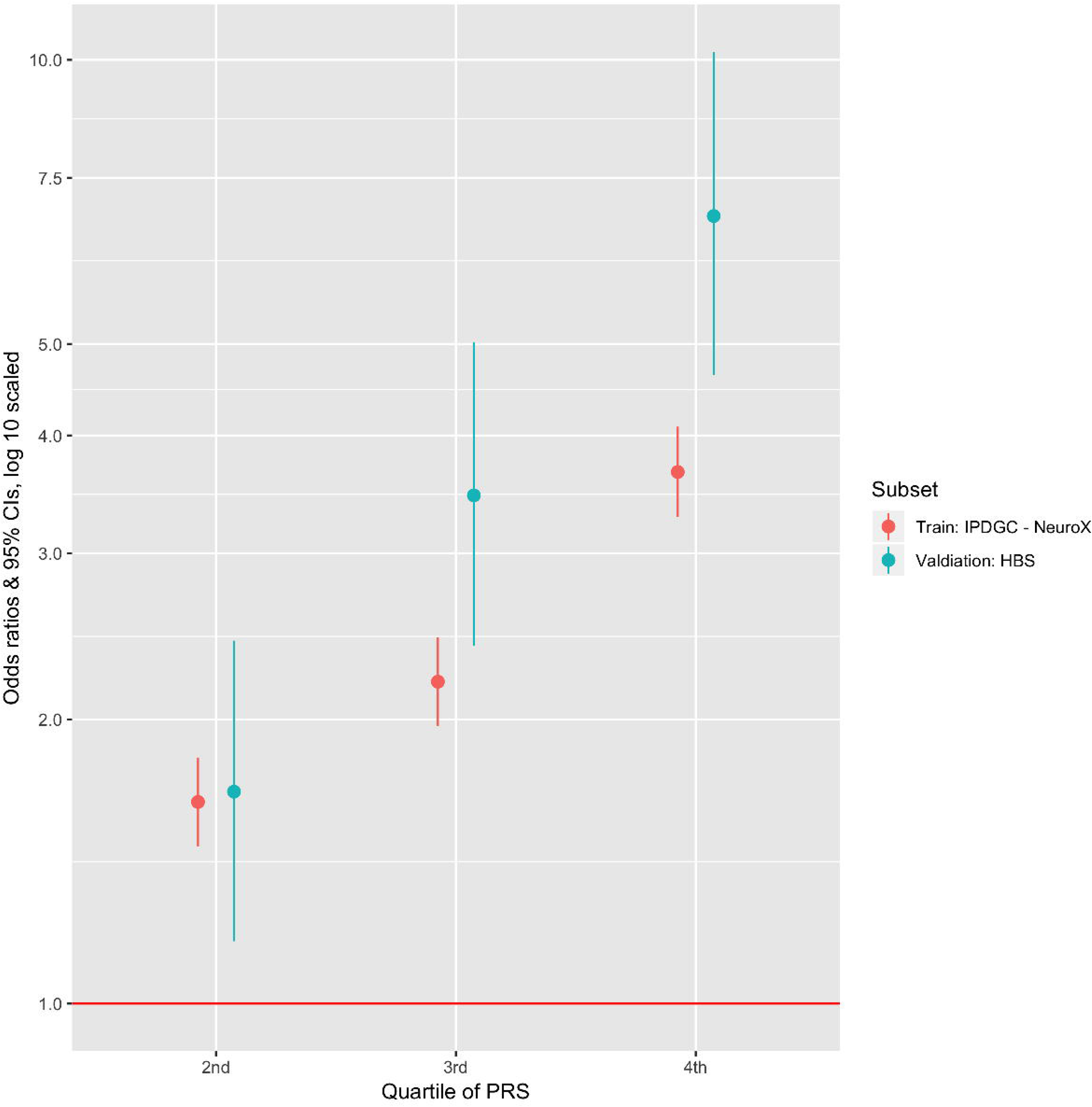
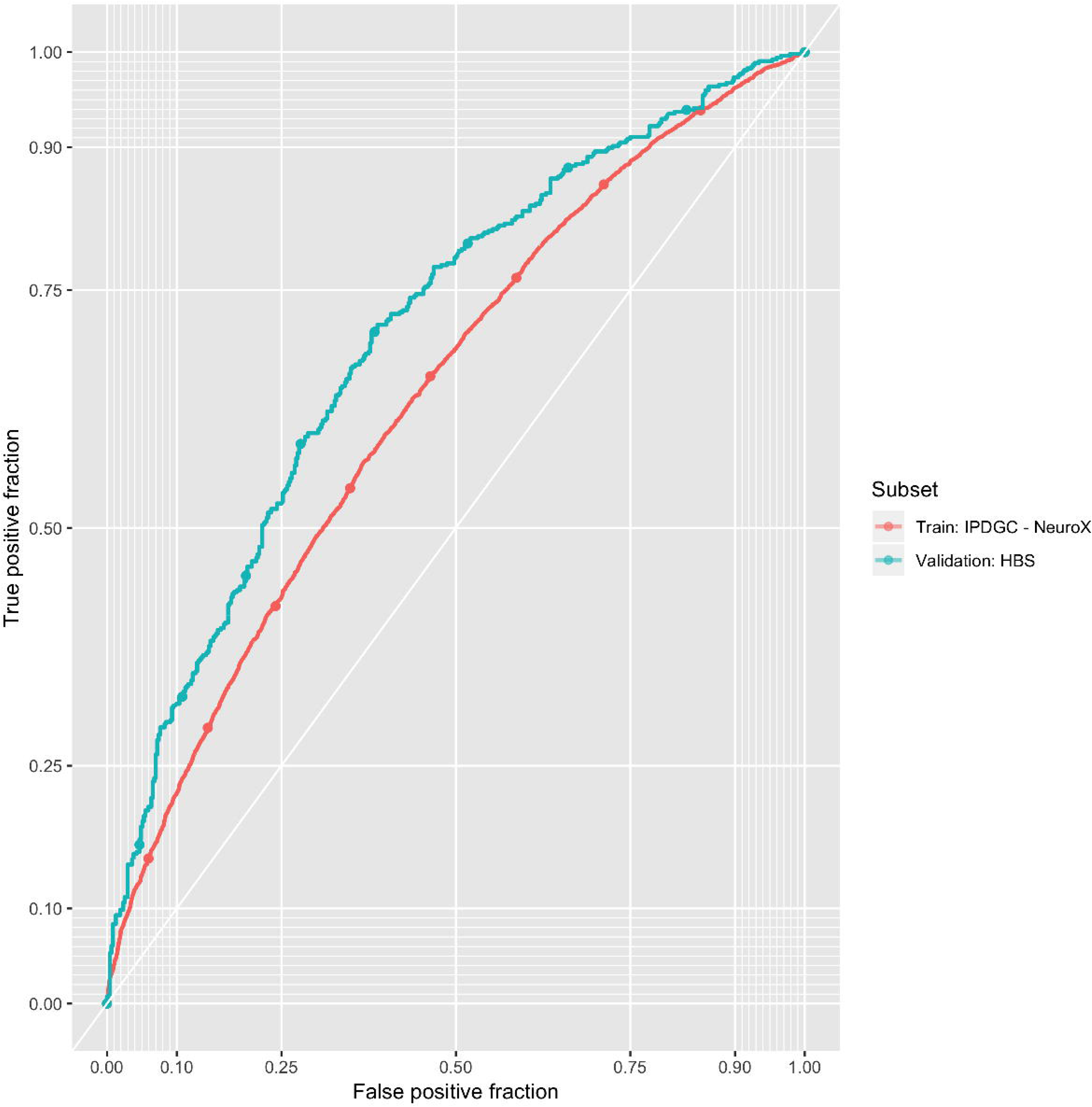
Predictive model details. The odds ratio of developing PD for each quartile of polygenic risk score (PRS) compared to the lowest quartile of genetic risk. B. PRS receiver-operator curves for each array type and sampling design.

Variants in the range of 5E-08 < P < 1.35E-03 (used in the 1805 variant PRS) were rarer and had smaller effect estimates than variants reaching genome-wide significance. These sub-significant variants had a median minor allele frequency of 21.3% and a median effect estimate (absolute value of the log odds ratio of the SNP parameter from regresion) of 0.047. Genome-wide significant risk variants were more common with a median minor allele frequency of 25.1%, and had a median effect estimate of 0.081. We performed power calculations to forecast the number of additional PD cases needed to achieve genome-wide significance at 80% power for a variant with a minor allele frequency of 21.3% and an effect estimate of 0.047 ^20^. Assuming that all incoming data is well harmonized with current data and that disease prevalence is 0.5%, we estimated that we would need a total of ∼99 K cases, ∼2.3 times as many as our current analysis. Variant discovery at this point will help us work towards the maximum achievable AUC for a genetic predictor in PD (estimated 85%). Past this point it is possible that effect estimates get too marginal, variants get too rare and they are no longer useful in predictions or in estimating heritability ^21^.

### Functional causal inferences via Quantitative Trait Loci (QTL)

There were 305 genes within the 78 GWAS loci. We sought to identify the causal gene in each locus in order to help direct future high-throughput functional studies. Specifically, we used large QTL datasets and summary-data-based Mendelian randomization (SMR) to test whether the expression or methylation of these genes led to a causal change in PD risk (Table 3, Table S5, Table S6)^11^. This method allows for functional inferences between two datasets to be made within a similar framework as a randomized controlled trial, treating the genotype as the randomizing factor.

**Table 3:**
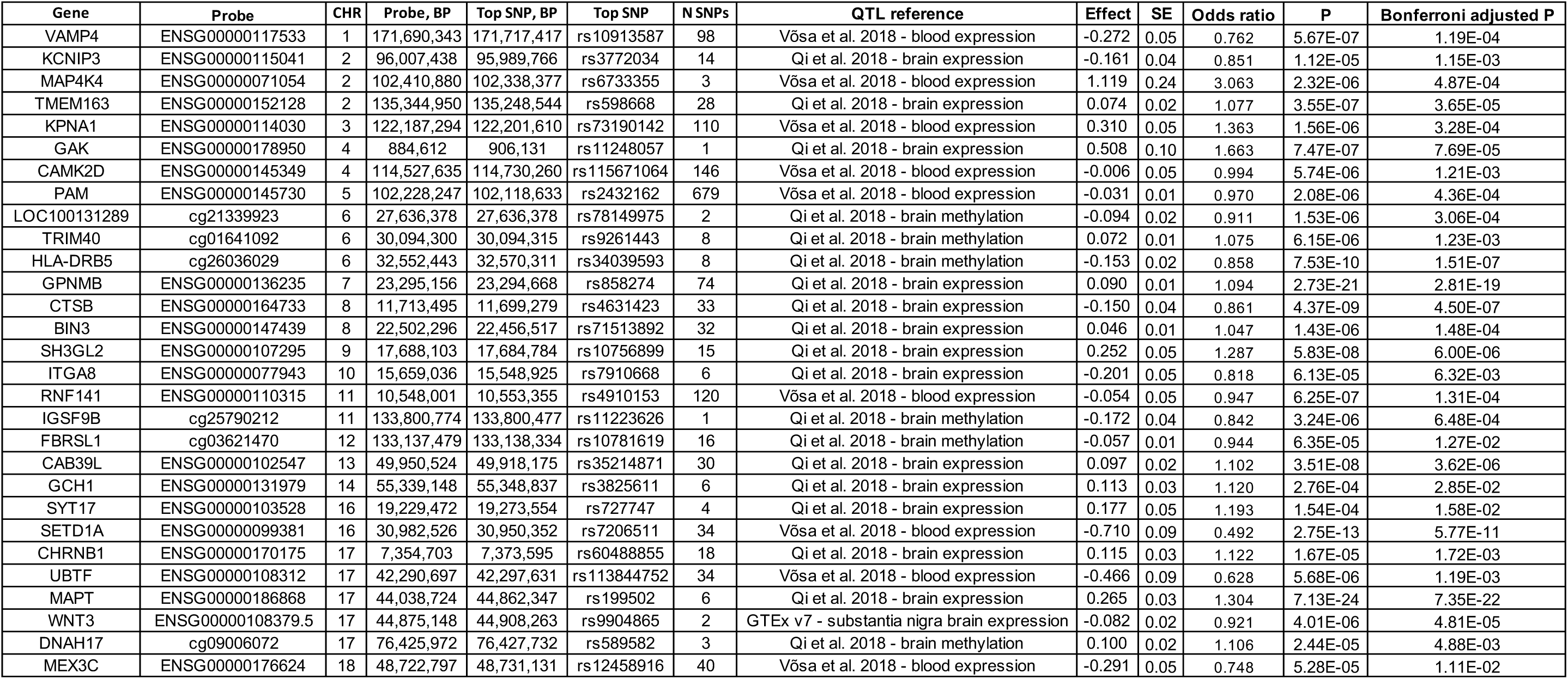
Summary of significant functional inferences from QTL associations via Mendelian randomization for nominated genes of interest. Multi-SNP eQTL Mendelian randomization results focusing only on the most significant association per nearest genes to PD risk loci after Bonferroni correction. If a locus was significantly associated with both brain and blood QTLs after multiple test correction, we opted to show the most signficant brain tissue derived association here after filtering for possible polygenicity (HEIDI P > 0.01). All tested QTL summary statistics can be found in Supplementary Table S6. Effect estimates represent the change in PD odds ratio per one standard deviation increase in gene expression or methylation.

We used four QTL datasets: a large meta-analysis of mRNA expression across brain tissues, mRNA expression in the substantia nigra, mRNA expression in blood, and methylation in blood ^22–26^. Of the 305 genes under linkage disequilibrium (LD) peaks around our risk variants of interest, 237 were possibly associated with at least one QTL and were therefore testable via SMR (Supplementary Methods, Table S6). The expression or methylation of 151 of these 237 genes (63.7%) was significantly linked to a causal change in PD risk.

Of the 90 PD GWAS risk variants, 70 were in loci containing at least one of these putatively causal genes after multiple test correction (Table 3 summarizes top QTL per gene). For 53 out of these 70 PD GWAS hits (75.7%), the gene nearest to the sentinel SNP was a putatively causal gene (Table S2). Most loci tested contained multiple putatively causal genes. Interestingly, the nearest putatively causal gene to the rs850738 / *FAM171A2* GWAS risk signal is *GRN*, a gene known to be associated with frontotemporal dementia (FTD)^27^. Mutations in this gene (*GRN*) have also been shown to be connected with another lysosomal storage disorder, neuronal ceroid lipofuscinosis^28^.

### Rare coding variant burden analysis

As an orthogonal approach for nominating putatively causal genes, we also carried out rare coding variant burden analyses. While the main GWAS analysis was limited to MAF ≥ 1% (except for known coding risk variants), we carried out rare variant burden analyses in a subset of studies (Supplementary Methods). We performed kernel-based burden tests on the 113 genes in our PD GWAS loci that contained two or more rare coding variants (MAF < 5% or MAF < 1%). After Bonferroni correction for 113 genes, we identified 7 significant putatively causal genes: *LRRK2*, *GBA*, *CATSPER3* (rs11950533/*C5orf24* locus), *LAMB2* (rs12497850/*IP6K2* locus), *LOC442028* (rs2042477/*KCNIP3* locus), *NFKB2* (rs10748818/*GBF1* locus), and *SCARB2* (rs6825004 locus). These results suggest that some of the risk associated with these loci may be due to rare coding variants. The *LRRK2* and *NFKB2* associations at MAF < 1% remained significant after correcting for all ∼20,000 genes in the human genome (*P* = 2.15E-10 and *P* = 4.02E-07, Table S7, Table S5).

### Tissue and cell specific expression enrichment plus protein-protein interactions

In order to better understand the function of the genes highlighted by this study, we tested whether these genes were enriched in 10,651 biological pathways. We tested for gene expression and pathway enrichment in PD loci using Functional Mapping and Annotation of Genome-Wide Association Studies (FUMA) and webgestaltR, respectively ^9, 29^. We found 10 significantly enriched pathways (false discovery rate [FDR]-adjusted P < 0.05, Table S8), including four related to vacuolar function and three related to known drug targets (calcium transporters: ikeda_mir1_targets_dn and ikeda_mir30_targets_up, kinase signaling: kim_pten_targets_dn). Known pathways of interest relating to lysosomal function, endocytosis, and dopamine metabolism were significantly enriched when using a more lenient *P* value (FDR-adjusted *P* < 0.1). At least three candidate genes within novel loci are involved in lysosomal storage disorder (*GUSB*, *GRN,* and *NEU1)*, a pathway of interest in recent PD research ^30^.

Next, we sought to determine the tissues and cell types most relevant to PD etiology using FUMA^9^. We tested whether the genes highlighted by our PD GWAS were enriched for expression in 53 tissues from across the body. We found 13 significant tissues, all of which were brain-derived (Figure S2A), in contrast to what has been seen in Alzheimer’s disease which shows a strong bias towards blood, spleen, lungs and microglial enrichments ^31^. To further disentangle the enrichment in brain tissues, we tested whether our PD GWAS genes were enriched for expression in 88 brain cell types using single cell RNA sequencing reference data from DropViz (http://dropviz.org)^32^. After false discovery rate correction we found seven significant brain cell types, all of which were neuronal (Figure S2B). The strongest enrichment was for neurons in the substantia nigra (SN) at P = 1.0E-06, with additional significant results at P < 5.0E-4 for the globus pallidus (GP), thalamus (TH), posterior cortex (PC), frontal cortex (FC), hippocampus (HC) and entopeduncular nucleus (ENT).

Finally, we analyzed protein-protein interaction networks using webgestaltR^29^ and found that the genes highlighted by our PD GWAS were enriched in six functional ontological networks (FDR-adjusted *P* < 0.1). The majority of these networks were related to chemical signaling pathways or response to some type of stressor. The most significant protein-protein interaction was related to response to interferon-gamma (Table S9, Figure S3A, Figure S3B).

### Genetic correlations and Mendelian randomization across phenotypes

Next, we used cross-trait genetic correlation and Mendelian randomization to identify putative PD biomarkers and risk factors. We estimated the cross-trait genetic correlation between our PD GWAS and 757 other GWAS datasets curated by LD hub ^33^. We found four significant genetic correlations (FDR-adjusted *P* < 0.05, Table 4, Table S10) including positive correlations with intracranial volume (rG = 0.351, SE = 0.077, P = 4.64E-06) and putamen volume (rG = 0.248, SE = 0.064, P = 9.55E-05, respectively)^34^, and negative correlations with current tobacco use (rG = −0.134, SE = 0.034, P = 7.92E-05) and “academic qualifications: National Vocational Qualifications (NVQ) or Higher National Diploma (HND) or Higher National Certificate (HNC) or equivalent” (rG = −0.169, SE 0.045, P = 2.00E-04)^35^. The negative association with one’s academic qualifications suggests that individuals without a college education may be at less risk of PD than individuals with higher levels of education. The correlation between PD and smoking status may not be independent from the correlation between PD and education as smoking status and years of education were significantly correlated (rG = −0.361, SE = 0.064, P = 1.64E-08) ^36^.

**Table 4:** Significant cross-trait genetic correlations. The genetic correlations between PD and four significantly-associated traits from LD Hub. An extended version of this table included in the Supplementary Materials (Supplementary Table S10) showing data for all tested correlations. In this table, h2 represents the heritability estimate of the trait.

| Trait of interest | PMID | correlation | SE, RG | Z, RG | P, RG | P, FDR adjusted | Observed H2 | SE, H2 | H2 intercept | SE, H2 intercept | Cross trait intercept | SE, Cross trait intercept |
| --- | --- | --- | --- | --- | --- | --- | --- | --- | --- | --- | --- | --- |
| Intracranial volume | 25607358 | 0.351 | 0.077 | 4.580 | 4.64E-06 | 3.51E-03 | 0.166 | 0.045 | 1.003 | 0.007 | -0.013 | 0.005 |
| Current tobacco smoking | Not available (UKB) | -0.134 | 0.034 | -3.947 | 7.92E-05 | 2.41E-02 | 0.055 | 0.003 | 1.014 | 0.010 | 0.004 | 0.007 |
| Mean Putamen | 25607358 | 0.248 | 0.064 | 3.902 | 9.55E-05 | 2.41E-02 | 0.282 | 0.047 | 0.952 | 0.007 | -0.007 | 0.006 |
| Qualifications: NVQ or HND or HNC or equivalent | Not available (UKB) | -0.169 | 0.045 | -3.726 | 2.00E-04 | 3.79E-02 | 0.015 | 0.002 | 1.011 | 0.007 | 0.005 | 0.005 |

We used Mendelian randomization to assess whether there was evidence of a causal relationship between PD and five phenotypes related to academic qualification, smoking, and brain volumes described above (Figure S4. Cognitive performance had a large, significant causal effect on PD risk (MR effect = 0.213, SE = 0.041, Bonferroni-adjusted P = 8.00E-07), while PD risk did not have a significant causal effect on cognitive performance (Bonferroni-adjusted P = 0.125). Educational attainment also had a significant causal effect on PD risk (MR effect = 0.162, SE = 0.040, Bonferroni-adjusted P = 2.06E-04), but PD risk also had a weak but significant causal effect on educational attainment (MR effect = 0.007, SE = 0.002, Bonferroni-adjusted P = 7.45E-3). There was no significant causal relationship between PD and current smoking status (forward analysis: MR effect = −0.069, SE = 0.031, Bonferroni-adjusted P = 0.125; reverse analysis: MR effect = 0.004, SE = 0.010, Bonferroni-adjusted P = 1). Smoking initiation (the act of ever starting smoking) did not have a causal effect on PD risk (MR effect = - 0.063, SE = 0.034, Bonferroni-adjusted P = 0.315), whereas PD had a small, but significantly positive causal effect on smoking initiation (MR effect = 0.027, SE = 0.006, Bonferroni-adjusted P = 1.62E-05). Intracranial volume could not be tested because its GWAS did not contain any genome-wide significant risk variants. There was no significant causal relationship between PD and putamen volume (P > 0.05 in both the forward and reverse directions).

## DISCUSSION

Our work marks a significant step forward in our understanding of the genetic architecture of PD and provides a genetic reference set for the broader research community. We identified 90 independent common genetic risk factors for PD, nearly doubling the number of known PD risk variants. We re-evaluated the cumulative contribution of genetic risk variants, both genome-wide significant and not-yet discovered, in order to refine our estimates of heritable Parkinson’s disease risk. We also nominated likely causal genes at each locus for further follow-up using QTL analyses and rare variant burden analyses. Our work has highlighted the pathways, tissues, and cell types involved in PD etiology. Finally, we identified intracranial and putaminal volume as potential PD biomarkers, and cognitive performance as a PD risk factor. Altogether, the data presented here has significantly expanded the resources available for future investigations into potential PD interventions.

Using a PRS constructed from our GWAS results, we were able to explain up to 36% of PD heritability. Power estimates suggest that expansions of case numbers to 99 K cases will continue to reveal additional insights into PD genetics. While these yet-to-be defined risk variants will have relatively small effects, cumulatively they will improve our ability to predict PD and will help to further expand our knowledge of the genes and pathways that drive PD risk. Population-wide screening for individuals who are likely to develop PD is currently not feasible using our 1805 variant PRS alone. There would be roughly 14 false positives per true positive assuming a prevalence of 0.5%. While large-scale genome sequencing and non-linear machine learning methods will likely improve these predictive models, we have previously shown that we will need to incorporate other data sources (*e.g.* smell tests, family history, age, sex) in order to generate algorithms that are useful for population-wide screening ^18^.

Evaluating these results in the larger context of pathway, tissue, and cellular functionality revealed that genes near PD risk variants showed enrichment for expression in the brain, contrasting with previous work in Alzheimer’s disease. Notably, we showed that the expression enrichment of genes at PD loci occured exclusively in neuronal cell types. We also found that PD genes were enriched in chemical signaling pathways and pathways involving the response to a stressor. These observations may be informative for disease modeling efforts, highlighting the importance of disease modeling in neurons and of incorporating a cellular stress component. This will help inform and focus stem cell derived therapeutic development efforts that are currently underway.

Using cross-trait LD score regression, we found four phenotypes that were genetically correlated with PD. Putamen and intracranial volumes may prove to be valuable PD biomarkers. Our bi-directional GSMR results suggest a complex etiological connection between smoking initiation and PD that will require further follow-up and should be viewed with some caution. One of the implications of this work is that PD trials of nicotine or other smoking-related compound(s) may be less likely to succeed. The strong causal effect of cognitive performance on PD is supported by observational studies ^37^.

While this study marks major progress in assessing genetic risk factors for PD, there remains a great deal to be done. No defined external validation dataset was used, which may be seen as a limitation. Simulations have suggested that without replication variants with *P* values between 5E-08 and 5E-9 should be interpreted with greater caution ^38, 39^. We found 16 risk variants in this *P* value range, including two known variants near *WNT3* (proximal to the *MAPT* locus) and *BIN3*. To a degree, the fact that we filtered our variants with a secondary random-effects meta-analysis may make our 90 PD GWAS hits somewhat more robust due to the conservative nature of random-effects. Secondly, this study focused on PD risk in individuals of European ancestry. Adding datasets from non-European populations would be helpful to further improve our granularity in association testing and ability to fine-map loci through integration of more variable LD signatures while also evaluating population specific associations. Additionally, large ancestry-specific PD LD reference panels, such as those for Ashkenazi Jewish patients, will help us further unravel the genetic architecture of loci such as GBA and LRRK2. This may be particularly crucial at these loci where LD patterns may be quite variable within European populations, accentuating the possible influence of LD reference series on conditional analyses in some cases ^40^. Finally, our work utilized state-of-the-art QTL datasets to nominate candidate genes, but many QTL associations are hampered by both small sample size and low *cis*-SNP density. Larger QTL studies and PD-specific network data from large scale cellular screens would allow us to build a more robust functional inference framework.

As the field moves forward there are some critical next steps that should be prioritized. First, allowing researchers to share participant-level data in a secure environment would facilitate inclusiveness and uniformity in analyses while maintaining the confidentiality of study participants. Our work suggests that GWASes including up to 99,000 cases will continue to provide useful biological insights into PD. In addition to studies of the genetics of PD risk, studies of disease onset, progression, and subtype will be important and will require large series of well-characterized patients. We also believe that work across diverse populations is important, not only to be able to best serve these populations but also to aid in fine mapping of loci. Notably, the use of genome sequencing technologies could further improve discovery by capturing rare variants and structural variants, but with the caveat that very large samples sizes will be required. While there is still much left to do, we believe that our current work represents a significant step forward and that the results and data will serve as a foundational resource for the community to pursue this next phase of PD research.

## SUPPLEMENTARY MATERIALS

**Supplementary Methods:** Detailed methods section.

**Supplemental Appendix:** This appendix is split into four sections detailing: first comparisons of effect estimates across GWAS cohorts (beta∼beta plots), second forest plots for each significant variant, thirdly locus plots showing regional GWAS results, and QTL and burden associations for each variant, finally the fourth section including extended PRS results. Beta∼beta plots compare the regression coefficients for up to 90 of the significant variants in one study to a meta-analysis of all others via linear regression. Forest plots communicate similar sensitivity analyses, for each of the 92 variants of interest. In the forest plots, box size indicates relative sample size for that study, and the width of the diamond representing the meta-analysis effect estimates indicate the 95% confidence interval. The locus plots are a zoomed-in version of Figure 2 for each of the 90 significant variants. These plots are truncated at a −log10 P value of 50 for display purposes and include the most significant burden test and QTL analysis results per gene denoted by label color-coding in each figure. In each locus plot, R2 is measured in our in-house LD reference dataset and shows the correlation between the most significant local SNP and all other proximal SNPs. Additional detailed PRS results for a subset of cohorts are available in the appendix summarizing PRS estimates at varied P thresholds. Each cohort specific PRS in the appendix is based on meta-analyses excluding that cohort when calculating SNP weights. A smaller table summarizing PRS associations at the P threshold with the highest r^2^ is also included. Column headers in the PRS section of the appendix mirror that of Table 2.

**Table S2:** Summary statistics for all nominated risk variants, known and novel. For binary variables, 0 = negative and 1 = positive. Some specific notes include: delineations of all studies, new studies and previous studies as discussed in the methods section. Betas and standard errors (StdErr) refer to effect estimates per SNP from logistic regression or fixed-effects meta-analyses.I2 is the index of heterogeneity. QTL Nominated Gene = genes which represent the nearest cis-QTL for that locus significant in MR.

**Table S4:** Estimates of genetic liability explained in different scenarios.

Here we compare how different AUC estimates and prevalence rates change the amount of genetic liability explained by GWAS.

**Table S6:** Complete summary statistics for QTL Mendelian randomization. Output from the SMR package for all QTLs of interest. Additional columns include QTL reference dataset, dataset-level Bonferroni corrected P values and a binary indicator if a candidate association passed multiple test correction. All columns prefixed by SMR indicate multi-SNP SMR results.

**Table S8:** FUMA expression pathway enrichment analysis results. Pathway enrichment from collapsed GWAS summary statistics.

**Table S10:** Bivariate LDscores. Default output from LD Hub. Abbreviations defined in main text and methods section.

**Text S1:** Authors and affiliations.

**Text S2:** Acknowledgements and Funding.

## AUTHOR CONTRIBUTIONS

## Study level analysis

MAN, CB, CLV, KH, SB-C, DC, MT, DK, LR, JS-S, LK, LP, ABS

## Additional analysis and data management

MAN, CB, SB-C, AJN, AX, JY, JG, PMV, ABS

## Design and funding

MAN, CB, CLV, KH, SB-C, LP, MS, KM, MT, AB, JY, ZG-O, TG, PH, JMS, NW, DAH, JH, HRM, JG, PMV, RRG, ABS

## Critical review and writing the manuscript

MAN, CB, CLV, KH, SB-C, DC, MT, DAK, AJN, AX, JB, EY, RvC, JS-S, CS, MS, LK, LP, AS,

HI, HL, FF, JRG, DGH, SWS, JAB, MM, J-CC, SL, JJ, LMS, MS, PT, KM, MT, AB, JY, ZG-O, TG, PH, JMS, NW, DAH, JH, HRM, JG, PMV, RRG, ABS

## DATA ACCESS

GWAS summary statistics for 23andMe datasets (post-Chang and data included in Chang et al. 2017 and Nalls et al. 2014) will be made available through 23andMe to qualified researchers under an agreement with 23andMe that protects the privacy of the 23andMe participants. Please visit <u>research.23andme.com/collaborate/#publication</u> for more information and to apply to access the data. An immediately accessible version of the summary statistics is available <u>here</u> https://drive.google.com/file/d/1FZ9UL99LAqyWnyNBxxlx6qOUlfAnublN/view?usp=sharing excluding Nalls et al. 2014, 23andMe post-Chang et al. 2017 and Web-Based Study of Parkinson’s Disease (PDWBS) but including all analyzed SNPs. After applying with 23andMe, the full summary statistics including all analyzed SNPs and samples in this GWAS meta-analysis will be accessible to the approved researcher(s). Underlying participant level IPDGC data is available to potential collaborators, please contact.

**Figure S1:** The odds ratio of developing PD for each decile of PRS, comparing each decile to all others for all samples in this analysis.

**Figure S2:** Results of FUMA analysis for tissue and cell type specific expression enrichment. A. Tissue enrichment. B. Cell type-specific enrichment. Red bars indicate levels of significance surpassing multiple test correction.

**Figure S3:** Panel A: Gene ontology term connectivity within protein-protein networks. This panel shows network of gene ontology (GO) terms from pathway analyses. Most significant GO terms are shown in green. Panel B: Gene level connectivity within protein-protein networks. This panel shows connectivity between genes across enriched pathways.

**Figure S4:** Comparison of regression coefficients in Mendelian randomization analyses across traits. Each cross represents a SNP, with the dashed lines representing the trend across all variants. Axes position are regression coefficients from GWAS for significant SNPs from either GWAS. Panel A includes results for cognitive performance, panel B includes results for educational attainment, panel C includes results for putamen volume, panel D includes results for smoking initiation and panel E includes results for current smoking status.

**Table S1:** Descriptive statistics and quality control summaries for meta-analyzed genome-wide association studies. ! denotes age at exam for both cases and controls. $ denotes age at death, onset not available. * based on 599 PD cases and 715 controls. ^ denotes samples checked for overlap across datasets as per Nalls et al. 2014 and Chang et al. 2017, ^^ denotes checked for overlap within IPDGC sample series, ^^^denotes a combination of both workflows for identifying sample overlap.

**Table S3:** Comparison with novel results from Chang et al., 2017. This table summarizes linkage disequilibrium estimates between Chang et al., 2017 novel loci and variants passing quality control in this report.

**Table S5:** SNPs of interest tagging genes for functional inferences and networks analysis. Nominated genes and SNPs for follow-up analyses based on minimum r^2^ > 0.5 within +/-1MB of one of our 90 risk loci.

**Table S7:** Rare coding variant burden analyses for genes under GWAS peaks. Detailed results of burden tests for genes proximal to risk loci. This includes variant counts, test statistics (rho, q, P, adjusted P) for each gene of interest.

**Table S9:** Protein network analysis for linked genes under association peaks. Gene ontology terms passing false discovery rate adjustment.

