## Additional Supplementary Files for "Expanding Parkinson’s disease genetics: novel risk loci, genomic context, causal insights and heritable risk": Text S1_ Authors and affiliations_.docx

**United Kingdom:** Alastair J Noyce (Preventive Neurology Unit, Wolfson Institute of Preventive Medicine, QMUL, London, UK and Department of Molecular Neuroscience, UCL, London, UK), Arianna Tucci (Department of Molecular Neuroscience, UCL Institute of Neurology, London, UK), Ben Middlehurst (Institute of Translational Medicine, University of Liverpool, Liverpool, UK), Demis A Kia (UCL Genetics Institute; and Department of Molecular Neuroscience, UCL Institute of Neurology, London, UK), Manuela Tan (Department of Clinical Neuroscience, University College London, London, UK), Henry Houlden (Department of Molecular Neuroscience, UCL Institute of Neurology, London, UK), Huw R Morris (Department of Clinical Neuroscience, University College London, London, UK), Helene Plun-Favreau (Department of Molecular Neuroscience, UCL Institute of Neurology, London, UK), Peter Holmans (Biostatistics & Bioinformatics Unit, Institute of Psychological Medicine and Clinical Neuroscience, MRC Centre for Neuropsychiatric Genetics & Genomics, Cardiff, UK), John Hardy (Department of Molecular Neuroscience, UCL Institute of Neurology, London, UK), Daniah Trabzuni (Department of Molecular Neuroscience, UCL Institute of Neurology, London, UK; Department of Genetics, King Faisal Specialist Hospital and Research Centre, Riyadh, 11211 Saudi Arabia), Jose Bras (UK Dementia Research Institute at UCL and Department of Molecular Neuroscience, UCL Institute of Neurology, London, UK), John Quinn (Institute of Translational Medicine, University of Liverpool, Liverpool, UK), Kin Y Mok (Department of Molecular Neuroscience, UCL Institute of Neurology, London, UK), Kerri J. Kinghorn (Institute of Healthy Ageing, University College London, London, UK), Kimberley Billingsley (Institute of Translational Medicine, University of Liverpool, Liverpool, UK), Nicholas W Wood (UCL Genetics Institute; and Department of Molecular Neuroscience, UCL Institute of Neurology, London, UK), Patrick Lewis (University of Reading, Reading, UK), Rita Guerreiro (UK Dementia Research Institute at UCL and Department of Molecular Neuroscience, UCL Institute of Neurology, London, UK), Ruth Lovering (University College London, London, UK), Lea R’Bibo (Department of Molecular Neuroscience, UCL Institute of Neurology, London, UK), Mie Rizig (Department of Molecular Neuroscience, UCL Institute of Neurology, London, UK), Mina Ryten (Department of Molecular Neuroscience, UCL Institute of Neurology, London, UK), Valentina Escott-Price (MRC Centre for Neuropsychiatric Genetics and Genomics, Cardiff University School of Medicine, Cardiff, UK), Viorica Chelban (Department of Molecular Neuroscience, UCL Institute of Neurology, London, UK), Thomas Foltynie (UCL Institute of Neurology, London, UK), Nigel Williams (MRC Centre for Neuropsychiatric Genetics and Genomics, Cardiff, UK),

**France:** Alexis Brice (Institut du Cerveau et de la Moelle épinière, ICM, Inserm U 1127, CNRS, UMR 7225, Sorbonne Universités, UPMC University Paris 06, UMR S 1127, AP-HP, Pitié-Salpêtrière Hospital, Paris, France), Fabrice Danjou (Institut du Cerveau et de la Moelle épinière, ICM, Inserm U 1127, CNRS, UMR 7225, Sorbonne Universités, UPMC University Paris 06, UMR S 1127, AP-HP, Pitié-Salpêtrière Hospital, Paris, France), Suzanne Lesage (Institut du Cerveau et de la Moelle épinière, ICM, Inserm U 1127, CNRS, UMR 7225, Sorbonne Universités, UPMC University Paris 06, UMR S 1127, AP-HP, Pitié-Salpêtrière Hospital, Paris, France), Jean-Christophe Corvol (Institut du Cerveau et de la Moelle épinière, ICM, Inserm U 1127, CNRS, UMR 7225, Sorbonne Universités, UPMC University Paris 06, UMR S 1127, Centre d’Investigation Clinique Pitié Neurosciences CIC-1422, AP-HP, Pitié-Salpêtrière Hospital, Paris, France), Maria Martinez (INSERM UMR 1220; and Paul Sabatier University, Toulouse, France),

**Germany:** Anamika Giri (Department for Neurodegenerative Diseases, Hertie Institute for Clinical Brain Research, University of Tübingen, and DZNE, German Center for Neurodegenerative Diseases, Tübingen, Germany), Claudia Schulte (Department for Neurodegenerative Diseases, Hertie Institute for Clinical Brain Research, University of Tübingen, and DZNE, German Center for Neurodegenerative Diseases, Tübingen, Germany), Kathrin Brockmann (Department for Neurodegenerative Diseases, Hertie Institute for Clinical Brain Research, University of Tübingen, and DZNE, German Center for Neurodegenerative Diseases, Tübingen, Germany), Javier Simón-Sánchez (Department for Neurodegenerative Diseases, Hertie Institute for Clinical Brain Research, University of Tübingen, and DZNE, German Center for Neurodegenerative Diseases, Tübingen, Germany), Peter Heutink (DZNE, German Center for Neurodegenerative Diseases and Department for Neurodegenerative Diseases, Hertie Institute for Clinical Brain Research, University of Tübingen, Tübingen, Germany), Patrizia Rizzu (DZNE, German Center for Neurodegenerative Diseases), Manu Sharma (Centre for Genetic Epidemiology, Institute for Clinical Epidemiology and Applied Biometry, University of Tubingen, Germany), Thomas Gasser (Department for Neurodegenerative Diseases, Hertie Institute for Clinical Brain Research, and DZNE, German Center for Neurodegenerative Diseases, Tübingen, Germany),

**United States of America:** Aude Nicolas (Laboratory of Neurogenetics, National Institute on Aging, Bethesda, MD, USA), Mark R Cookson (Laboratory of Neurogenetics, National Institute on Aging, Bethesda, USA), Sara Bandres-Ciga (Laboratory of Neurogenetics, National Institute on Aging, Bethesda, MD, USA), Cornelis Blauwendraat (National Institute on Aging and National Institute of Neurological Disorders and Stroke, USA), Faraz Faghri (Laboratory of Neurogenetics, National Institute on Aging, Bethesda, USA; Department of Computer Science, University of Illinois at Urbana-Champaign, Urbana, IL, USA), J Raphael Gibbs (Laboratory of Neurogenetics, National Institute on Aging, National Institutes of Health, Bethesda, MD, USA), Dena G Hernandez (Laboratory of Neurogenetics, National Institute on Aging, Bethesda, MD, USA), Joshua M. Shulman (Departments of Neurology, Neuroscience, and Molecular & Human Genetics, Baylor College of Medicine, Houston, Texas, USA; Jan and Dan Duncan Neurological Research Institute, Texas Children’s Hospital, Houston, Texas, USA), Mike A. Nalls (Laboratory of Neurogenetics, National Institute on Aging, Bethesda, USA; CEO/Consultant Data Tecnica International, Glen Echo, MD, USA), Laurie Robak (Baylor College of Medicine, Houston, Texas, USA), Steven Lubbe (Ken and Ruth Davee Department of Neurology, Northwestern University Feinberg School of Medicine, Chicago, IL, USA), Steven Finkbeiner (Departments of Neurology and Physiology, University of California, San Francisco; Gladstone Institute of Neurological Disease; Taube/Koret Center for Neurodegenerative Disease Research, San Francisco, CA, USA), Niccolo E. Mencacci (Northwestern University Feinberg School of Medicine, Chicago, IL, USA), Codrin Lungu (National Institutes of Health Division of Clinical Research, NINDS, National Institutes of Health, Bethesda, MD, USA), Andrew B Singleton (Laboratory of Neurogenetics, National Institute on Aging, Bethesda, MD, USA), Sonja W. Scholz (Neurodegenerative Diseases Research Unit, National Institute of Neurological Disorders and Stroke, Bethesda, MD, USA), Xylena Reed (Laboratory of Neurogenetics, National Institute on Aging, Bethesda, MD, USA), Hampton Leonard (Laboratory of Neurogenetics, National Institute on Aging, Bethesda, MD, USA).

**Canada:** Ziv Gan-Or (Montreal Neurological Institute and Hospital, Department of Neurology & Neurosurgery, Department of Human Genetics, McGill University, Montréal, QC, H3A 0G4, Canada), Guy A. Rouleau (Montreal Neurological Institute and Hospital, Department of Neurology & Neurosurgery, Department of Human Genetics, McGill University, Montréal, QC, H3A 0G4, Canada), Lynne Krohan (Department of Human Genetics, McGill University, Montréal, QC, H3A 0G4, Canada),

**The Netherlands:** Jacobus J van Hilten (Department of Neurology, Leiden University Medical Center, Leiden, Netherlands), Johan Marinus (Department of Neurology, Leiden University Medical Center, Leiden, Netherlands)

**Spain:** Astrid D. Adarmes-Gómez (Instituto de Biomedicina de Sevilla (IBiS), Hospital Universitario Virgen del Rocío/CSIC/Universidad de Sevilla, Seville), Miquel Aguilar (Fundació Docència i Recerca Mútua de Terrassa and Movement Disorders Unit, Department of Neurology, University Hospital Mutua de Terrassa, Terrassa, Barcelona.),Ignacio Alvarez (Fundació Docència i Recerca Mútua de Terrassa and Movement Disorders Unit, Department of Neurology, University Hospital Mutua de Terrassa, Terrassa, Barcelona.),Victoria Alvarez (Hospital Universitario Central de Asturias, Oviedo), Francisco Javier Barrero (Hospital Universitario Parque Tecnologico de la Salud, Granada), Jesús Alberto Bergareche Yarza (Instituto de Investigación Sanitaria Biodonostia, San Sebastián), Inmaculada Bernal-Bernal (Instituto de Biomedicina de Sevilla (IBiS), Hospital Universitario Virgen del Rocío/CSIC/Universidad de Sevilla, Seville), Marta Blazquez (Hospital Universitario Central de Asturias, Oviedo), Marta Bonilla-Toribio (Instituto de Biomedicina de Sevilla (IBiS), Hospital Universitario Virgen del Rocío/CSIC/Universidad de Sevilla, Seville), Juan A. Botía (Universidad de Murcia, Murcia), María Teresa Boungiorno (Fundació Docència i Recerca Mútua de Terrassa and Movement Disorders Unit, Department of Neurology, University Hospital Mutua de Terrassa, Terrassa, Barcelona.) Dolores Buiza-Rueda (Instituto de Biomedicina de Sevilla (IBiS), Hospital Universitario Virgen del Rocío/CSIC/Universidad de Sevilla, Seville), Ana Cámara (Hospital Clinic de Barcelona), Maria Carcel (Fundació Docència i Recerca Mútua de Terrassa, Barcelona and Department of Neurology, Movement Disorders Unit, Hospital Universitari Mutua de Terrassa, Barcelona), Fátima Carrillo (Instituto de Biomedicina de Sevilla (IBiS), Hospital Universitario Virgen del Rocío/CSIC/Universidad de Sevilla, Seville), Mario Carrión-Claro (Instituto de Biomedicina de Sevilla (IBiS), Hospital Universitario Virgen del Rocío/CSIC/Universidad de Sevilla, Seville), Debora Cerdan (Hospital General de Segovia, Segovia), Jordi Clarimón (Memory Unit, Department of Neurology, IIB Sant Pau, Hospital de la Santa Creu i Sant Pau, Universitat Autònoma de Barcelona and Centro de Investigación Biomédica en Red en Enfermedades Neurodegenerativas (CIBERNED), Madrid),Yaroslau Compta (Hospital Clinic de Barcelona), Monica Diez-Fairen (Fundació Docència i Recerca Mútua de Terrassa and Movement Disorders Unit, Department of Neurology, University Hospital Mutua de Terrassa, Terrassa, Barcelona.), Oriol Dols-Icardo (Memory Unit, Department of Neurology, IIB Sant Pau, Hospital de la Santa Creu i Sant Pau, Universitat Autònoma de Barcelona, Barcelona, and Centro de Investigación Biomédica en Red en Enfermedades Neurodegenerativas (CIBERNED), Madrid), Jacinto Duarte (Hospital General de Segovia, Segovia), Raquel Duran (Centro de Investigacion Biomedica, Universidad de Granada, Granada), Francisco Escamilla-Sevilla (Hospital Universitario Virgen de las Nieves, Instituto de Investigación Biosanitaria de Granada, Granada), Mario Ezquerra (Hospital Clinic de Barcelona),Manel Fernández (Hospital Clinic de Barcelona), Rubén Fernández-Santiago (Hospital Clinic de Barcelona), Ciara Garcia (Hospital Universitario Central de Asturias, Oviedo), Pedro García-Ruiz (Instituto de Investigación Sanitaria Fundación Jiménez Díaz, Madrid), Pilar Gómez-Garre (Instituto de Biomedicina de Sevilla (IBiS), Hospital Universitario Virgen del Rocío/CSIC/Universidad de Sevilla, Seville), Maria Jose Gomez Heredia (Hospital Universitario Virgen de la Victoria, Malaga), Isabel Gonzalez-Aramburu (Hospital Universitario Marqués de Valdecilla-IDIVAL, Santander) ,Ana Gorostidi Pagola (Instituto de Investigación Sanitaria Biodonostia, San Sebastián), Janet Hoenicka (Institut de Recerca Sant Joan de Déu, Barcelona), Jon Infante (Hospital Universitario Marqués de Valdecilla-IDIVAL and University of Cantabria, Santander, and Centro de Investigación Biomédica en Red en Enfermedades Neurodegenerativas (CIBERNED), Silvia Jesús (Instituto de Biomedicina de Sevilla (IBiS), Hospital Universitario Virgen del Rocío/CSIC/Universidad de Sevilla, Seville), Adriano Jimenez-Escrig (Hospital Universitario Ramón y Cajal, Madrid), Jaime Kulisevsky (Movement Disorders Unit, Department of Neurology, IIB Sant Pau, Hospital de la Santa Creu i Sant Pau, Universitat Autònoma de Barcelona, Barcelona, and Centro de Investigación Biomédica en Red en Enfermedades Neurodegenerativas (CIBERNED)), Miguel A. Labrador-Espinosa (Instituto de Biomedicina de Sevilla (IBiS), Hospital Universitario Virgen del Rocío/CSIC/Universidad de Sevilla, Seville), Jose Luis Lopez-Sendon (Hospital Universitario Ramón y Cajal, Madrid), Adolfo López de Munain Arregui (Instituto de Investigación Sanitaria Biodonostia, San Sebastián), Daniel Macias (Instituto de Biomedicina de Sevilla (IBiS), Hospital Universitario Virgen del Rocío/CSIC/Universidad de Sevilla, Seville), Irene Martínez Torres (Department of Neurology, Instituto de Investigación Sanitaria La Fe, Hospital Universitario y Politécnico La Fe, Valencia), Juan Marín (Movement Disorders Unit, Department of Neurology, IIB Sant Pau, Hospital de la Santa Creu i Sant Pau, Universitat Autònoma de Barcelona, Barcelona, and Centro de Investigación Biomédica en Red en Enfermedades Neurodegenerativas (CIBERNED)), Maria Jose Marti (Hospital Clinic Barcelona), Juan Carlos Martínez-Castrillo (Instituto Ramón y Cajal de Investigación Sanitaria, Hospital Universitario Ramón y Cajal, Madrid), Carlota Méndez-del-Barrio (Instituto de Biomedicina de Sevilla (IBiS), Hospital Universitario Virgen del Rocío/CSIC/Universidad de Sevilla, Seville), Manuel Menéndez González (Hospital Universitario Central de Asturias, Oviedo), Adolfo Mínguez (Hospital Universitario Virgen de las Nieves, Granada, Instituto de Investigación Biosanitaria de Granada), Pablo Mir (Instituto de Biomedicina de Sevilla (IBiS), Hospital Universitario Virgen del Rocío/CSIC/Universidad de Sevilla, Seville), Elisabet Mondragon Rezola (Instituto de Investigación Sanitaria Biodonostia, San Sebastián), Esteban Muñoz (Hospital Clinic Barcelona), Javier Pagonabarraga (Movement Disorders Unit, Department of Neurology, IIB Sant Pau, Hospital de la Santa Creu i Sant Pau, Universitat Autònoma de Barcelona, Barcelona, and Centro de Investigación Biomédica en Red en Enfermedades Neurodegenerativas (CIBERNED)), Pau Pastor (Fundació Docència i Recerca Mútua de Terrassa and Movement Disorders Unit, Department of Neurology, University Hospital Mutua de Terrassa, Terrassa, Barcelona.), Francisco Perez Errazquin (Hospital Universitario Virgen de la Victoria, Malaga), Teresa Periñán-Tocino (Instituto de Biomedicina de Sevilla (IBiS), Hospital Universitario Virgen del Rocío/CSIC/Universidad de Sevilla, Seville), Javier Ruiz-Martínez (Hospital Universitario Donostia, Instituto de Investigación Sanitaria Biodonostia, San Sebastián), Clara Ruz (Centro de Investigacion Biomedica, Universidad de Granada, Granada), Antonio Sanchez Rodriguez (Hospital Universitario Marqués de Valdecilla-IDIVAL, Santander), María Sierra (Hospital Universitario Marqués de Valdecilla-IDIVAL, Santander), Esther Suarez-Sanmartin (Hospital Universitario Central de Asturias, Oviedo), Cesar Tabernero (Hospital General de Segovia, Segovia), Juan Pablo Tartari (Fundació Docència i Recerca Mútua de Terrassa and Movement Disorders Unit, Department of Neurology, University Hospital Mutua de Terrassa, Terrassa, Barcelona), Cristina Tejera-Parrado (Instituto de Biomedicina de Sevilla (IBiS), Hospital Universitario Virgen del Rocío/CSIC/Universidad de Sevilla, Seville), Eduard Tolosa (Hospital Clinic Barcelona), Francesc Valldeoriola (Hospital Clinic Barcelona), Laura Vargas-González (Instituto de Biomedicina de Sevilla (IBiS), Hospital Universitario Virgen del Rocío/CSIC/Universidad de Sevilla, Seville), Lydia Vela (Department of Neurology, Hospital Universitario Fundación Alcorcón, Madrid), Francisco Vives (Centro de Investigacion Biomedica, Universidad de Granada, Granada).

**Austria:** Alexander Zimprich (Department of Neurology, Medical University of Vienna, Austria)

**Norway:** Lasse Pihlstrom (Department of Neurology, Oslo University Hospital, Oslo, Norway)

**Estonia**: Sulev Koks (Department of Pathophysiology, University of Tartu, Tartu, Estonia; Department of Reproductive Biology, Estonian University of Life Sciences, Tartu, Estonia; Perron Institute for Neurological and Translational Science, Perth, Western Australia, Australia, Pille Taba (Department of Neurology and Neurosurgery, University of Tartu, Tartu, Estonia)

**Finland:** Kari Majamaa (Institute of Clinical Medicine, Department of Neurology, University of Oulu, Oulu, Finland; Department of Neurology and Medical Research Center, Oulu University Hospital, Oulu, Finland), Ari Siitonen (Institute of Clinical Medicine, Department of Neurology, University of Oulu, Oulu, Finland; Department of Neurology and Medical Research Center, Oulu University Hospital, Oulu, Finland)

**Nigeria**: Njideka U. Okubadejo (University of Lagos, Lagos State, Nigeria), Oluwadamilola O. Ojo (University of Lagos, Lagos State, Nigeria),

**23AndMe RESEARCH TEAM MEMBERS:**

Michelle Agee, Babak Alipanahi, Adam Auton, Robert K. Bell, Katarzyna Bryc, Sarah L. Elson, Pierre Fontanillas, Nicholas A. Furlotte, Karen E. Huber, Barry Hicks, Ethan M. Jewett, Yunxuan Jiang, Aaron Kleinman, Keng-Han Lin, Nadia K. Litterman, Jennifer C. McCreight, Matthew H. McIntyre, Kimberly F. McManus, Joanna L. Mountain, Elizabeth S. Noblin, Carrie A.M. Northover, Steven J. Pitts, G. David Poznik, J. Fah Sathirapongsasuti, Janie F. Shelton, Suyash Shringarpure, Chao Tian, Joyce Y. Tung, Vladimir Vacic, Xin Wang, and Catherine H. Wilson.

**SYSTEM GENOMICS OF PARKINSON’S DISEASE COLLABORATORS:**

Tim Anderson(New Zealand Brain Research Institute, Christchurch, New Zealand; Department of Medicine, University of Otago, Christchurch, New Zealand),Steven Bentley(Griffith Institute for Drug Discovery, Griffith University, Brisbane, Australia), John Dalrymple-Alford(Dept. Psychology, University of Canterbury, New Zealand Brain Research Institute), Javed Fowdar(Griffith Institute for Drug Discovery, Griffith University, Brisbane, Australia),Jacob Gratten (Institute for Molecular Bioscience, University of Queensland, Brisbane, Australia; Queensland Brain Institute, University of Queensland, Brisbane, Australia), Glenda Halliday(Brain and Mind Centre, Sydney Medical School, University of Sydney, Sydney, Australia),Anjali K. Henders(Institute for Molecular Bioscience, University of Queensland, Brisbane, Australia),Ian Hickie(Brain and Mind Centre, Sydney Medical School, University of Sydney, Sydney, Australia),Irfahan Kassam(Institute for Molecular Bioscience, University of Queensland, Brisbane, Australia),Martin Kennedy(Department of Pathology, University of Otago, Christchurch, New Zealand),John Kwok(Brain and Mind Centre, Sydney Medical School, University of Sydney, Sydney, Australia),Simon Lewis(Brain and Mind Centre, Sydney Medical School, University of Sydney, Sydney, Australia),George Mellick(Griffith Institute for Drug Discovery, Griffith University, Brisbane, Australia),Grant Montgomery(Institute for Molecular Bioscience, University of Queensland, Brisbane, Australia),John Pearson(Department of Pathology, University of Otago, Christchurch, New Zealand),Toni Pitcher(New Zealand Brain Research Institute, Christchurch, New Zealand; Department of Medicine, University of Otago, Christchurch, New Zealand),Julia Sidorenko(Institute for Molecular Bioscience, University of Queensland, Brisbane, Australia),Peter A. Silburn(Queensland Brain Institute, University of Queensland, Brisbane, Australia),Costanza L. Vallerga(Institute for Molecular Bioscience, University of Queensland, Brisbane, Australia), Peter M. Visscher(Institute for Molecular Bioscience, University of Queensland, Brisbane, Australia; Queensland Brain Institute, University of Queensland, Brisbane, Australia), Leanne Wallace, (Institute for Molecular Bioscience, University of Queensland, Brisbane, Australia), Naomi R. Wray(Institute for Molecular Bioscience, University of Queensland, Brisbane, Australia; Queensland Brain Institute, University of Queensland, Brisbane, Australia),Angli Xue(Institute for Molecular Bioscience, University of Queensland, Brisbane, Australia), Jian Yang (Institute for Molecular Bioscience, University of Queensland, Brisbane, Australia; Queensland Brain Institute, University of Queensland, Brisbane, Australia), Futao Zhang(Institute for Molecular Bioscience, University of Queensland, Brisbane, Australia)

**OTHER COLLABORATORS:**

**Department of Neurology, University of Maryland School of Medicine:** Lisa M. Shulman, Rainer von Coelln, Stephen Reich, Joseph Savitt

**Institute of Clinical Medicine, Department of Neurology, University of Turku, Turku, Finland; Division of Clinical Neurosciences, Turku University Hospital, Turku, Finland:** Pauli Ylikotila
