## Additional Supplementary Files for "Expanding Parkinson’s disease genetics: novel risk loci, genomic context, causal insights and heritable risk": Text S2_ Acknowledgements and Funding_.docx

**ACKNOWLEDGMENTS:**

We would like to thank all of the subjects who donated their time and biological samples to be a part of this study. This work was supported in part by the Intramural Research Programs of the National Institute of Neurological Disorders and Stroke (NINDS), the National Institute on Aging (NIA), and the National Institute of Environmental Health Sciences both part of the National Institutes of Health, Department of Health and Human Services; project numbers 1ZIA-NS003154, Z01-AG000949-02 and Z01-ES101986. In addition this work was supported by the Department of Defense (award W81XWH-09-2-0128), and The Michael J Fox Foundation for Parkinson’s Research. John Hardy’s contribution was in part supported by MR/N026004/1. This work was supported by National Institutes of Health grants R01NS037167, R01CA141668, P50NS071674, American Parkinson Disease Association (APDA); Barnes Jewish Hospital Foundation; Greater St Louis Chapter of the APDA. The KORA (Cooperative Research in the Region of Augsburg) research platform was started and financed by the Forschungszentrum für Umwelt und Gesundheit, which is funded by the German Federal Ministry of Education, Science, Research, and Technology and by the State of Bavaria. This study was also funded by the German Federal Ministry of Education and Research (BMBF) under the funding code 031A430A, the EU Joint Programme - Neurodegenerative Diseases Research (JPND) project under the aegis of JPND -[www.jpnd.eu](http://www.jpnd.eu)- through Germany, BMBF, funding code 01ED1406 and iMed - the Helmholtz Initiative on Personalized Medicine. This study is funded by the German National Foundation grant (DFG SH599/6-1) (grant to M.S), Michael J Fox Foundation, and MSA Coalition, USA (to M.S). The French GWAS work was supported by the French National Agency of Research (ANR-08-MNP-012). This study was also funded by France-Parkinson Association, Fondation de France, the French program “Investissements d’avenir” funding (ANR-10-IAIHU-06) and a grant from Assistance Publique-Hôpitaux de Paris (PHRC, AOR-08010) for the French clinical data. This study was also sponsored by the Landspitali University Hospital Research Fund (grant to SSv); Icelandic Research Council (grant to SSv); and European Community Framework Programme 7, People Programme, and IAPP on novel genetic and phenotypic markers of Parkinson’s disease and Essential Tremor (MarkMD), contract number PIAP-GA-2008-230596 MarkMD (to HP and JHu). Institutional research funding IUT20-46 was received of the Estonian Ministry of Education and Research (SK). Finnish exome sequencing study was partly funded by Sigrid Juselius Foundation. This study utilized the high-performance computational capabilities of the Biowulf Linux cluster at the National Institutes of Health, Bethesda, Md. (http://biowulf.nih.gov), and DNA panels, samples, and clinical data from the National Institute of Neurological Disorders and Stroke Human Genetics Resource Center DNA and Cell Line Repository. People who contributed samples are acknowledged in descriptions of every panel on the repository website. We thank the French Parkinson’s Disease Genetics Study Group and the Drug Interaction with genes (DIGPD) study group: Y Agid, M Anheim, F Artaud, A-M Bonnet, C Bonnet, F Bourdain, J-P Brandel, C Brefel-Courbon, M Borg, A Brice, E Broussolle, F Cormier-Dequaire, J-C Corvol, P Damier, B Debilly, B Degos, P Derkinderen, A Destée, A Dürr, F Durif, A Elbaz, D Grabli, A Hartmann, S Klebe, P. Krack, J Kraemmer, S Leder, S Lesage, R Levy, E Lohmann, L Lacomblez, G Mangone, L-L Mariani, A-R Marques, M Martinez, V Mesnage, J Muellner, F Ory-Magne, F Pico, V Planté-Bordeneuve, P Pollak, O Rascol, K Tahiri, F Tison, C Tranchant, E Roze, M Tir, M Vérin, F Viallet, M Vidailhet, A You. We also thank the members of the French 3C Consortium: A Alpérovitch, C Berr, C Tzourio, and P Amouyel for allowing us to use part of the 3C cohort, and D Zelenika for support in generating the genome-wide molecular data. We thank P Tienari (Molecular Neurology Programme, Biomedicum, University of Helsinki), T Peuralinna (Department of Neurology, Helsinki University Central Hospital), L Myllykangas (Folkhalsan Institute of Genetics and Department of Pathology, University of Helsinki), and R Sulkava (Department of Public Health and General Practice Division of Geriatrics, University of Eastern Finland) for the Finnish controls (Vantaa85+ GWAS data). We used genome-wide association data generated by the Wellcome Trust Case-Control Consortium 2 (WTCCC2) from UK patients with Parkinson’s disease and UK control individuals from the 1958 Birth Cohort and National Blood Service. Genotyping of UK replication cases on ImmunoChip was part of the WTCCC2 project, which was funded by the Wellcome Trust (083948/Z/07/Z). UK population control data was made available through WTCCC1. This study was supported by the Medical Research Council and Wellcome Trust disease centre (grant WT089698/Z/09/Z to NW, JHa, and ASc). As with previous IPDGC efforts, this study makes use of data generated by the Wellcome Trust Case-Control Consortium. A full list of the investigators who contributed to the generation of the data is available from www.wtccc.org.uk. Funding for the project was provided by the Wellcome Trust under award 076113, 085475 and 090355. This study was also supported by Parkinson’s UK (grants 8047 and J-0804) and the Medical Research Council (G0700943 and G1100643). Sequencing and genotyping done in McGill University was supported by grants from the Michael J Fox Foundation, the Canadian Consortium on Neurodegeneration in Aging (CCNA), the Canada First Research Excellence Fund, awarded to McGill University for the Healthy Brains for Healthy Lives (HBHL) program. The access to part of the participants in the McGill cohort has been made possible thanks to the Quebec Parkinson’s Network (http://rpq-qpn.ca/en/). Ziv Gan-Or is supported by the FRQS Chercheurs-boursiers Award, granted by the Fonds de recherche du Québec – Santé (FRQS) and Parkinson’s Quebec. We thank Jeffrey Barrett and Jason Downing for assistance with the design of the ImmunoChip and NeuroX arrays. DNA extraction work that was done in the UK was undertaken at University College London Hospitals, University College London, who received a proportion of funding from the Department of Health’s National Institute for Health Research Biomedical Research Centres funding. This study was supported in part by the Wellcome Trust/Medical Research Council Joint Call in Neurodegeneration award (WT089698) to the Parkinson’s Disease Consortium (UKPDC), whose members are from the UCL Institute of Neurology, University of Sheffield, and the Medical Research Council Protein Phosphorylation Unit at the University of Dundee. We thank the Quebec Parkinson’s Network (http://rpq-qpn.org) and its members. This work was supported by the Medical Research Council grant MR/N026004/1. The Braineac project was supported by the MRC through the MRC Sudden Death Brain Bank Grant (MR/G0901254) to J.H. P.A.L. was supported by the MRC (grants MR/N026004/1 and MR/L010933/1) and Michael J. Fox Foundation for Parkinson’s Research. D.T. was supported by the King Faisal Specialist Hospital and Research Centre, Saudi Arabia, and the Michael J. Fox Foundation for Parkinson’s Research and MRC grant (MR/N026004/1). Lasse Pihlstrøm is supported by the Norwegian Health Association and Michael J. Fox Foundation. Mathias Toft is supported by the Research Council of Norway and the South-Eastern Norway Regional Health Authority. We thank Ole Andreassen and the DemGene consortium, Norway for genotyping of the Oslo cohort. Mike A. Nalls’ participation is supported by a consulting contract between Data Tecnica International and the National Institute on Aging, NIH, Bethesda, MD, USA, as a possible conflict of interest Dr. Nalls also consults for SK Therapeutics Inc, Lysosomal Therapeutics Inc, the Michael J. Fox Foundation and Vivid Genomics among others. Joshua M. Shulman was supported by Huffington Foundation, the Jan and Dan Duncan Neurological Research Institute at Texas Children’s Hospital, and a Career Award for Medical Scientists from the Burroughs Wellcome Fund. The Baylor College of Medicine and University of Maryland replication cohorts were made possible in part due to assistance from Amanda Stillwell, Christina Griffin, Katrina Schrader, and Weidong Le with sample collection, preparation, and handling. Data used in the preparation of this article were obtained from the Parkinson’s Progression Markers Initiative (PPMI) database ([www.ppmi-info.org/data](http://www.ppmi-info.org/data)). For up-to-date information on the study, visit [www.ppmi-info.org](http://www.ppmi-info.org/). PPMI – a public-private partnership – is funded by The Michael J. Fox Foundation for Parkinson’s Research and funding partners, including Abbvie, Allergan, Avid Radiopharmaceuticals, Biogen, BioLegend, Bristol-Myers Squibb, Denali, GE Healthcare, Genentech, GlaxoSmithKline, Lilly, Lundbeck, Merck, Meso Scale Discovery, Pfizer, Piramal, Roche, Sanofi, Servier, Takeda, Teva, and UCB. . The SGPD’s contribution was supported by the Australian Research Council (ARC) (DP160102400) and the Australian National Health and Medical Research Council (NHMRC) (1078037,1078901, 1103418, 1107258, 1127440, 1113400). Support also came from ForeFront, a large collaborative research group dedicated to the study of neurodegenerative diseases and funded by the NHMRC (Program Grant 1132524, Dementia Research Team Grant 1095127, NeuroSleep Centre of Research Excellence 1060992) and ARC (Centre of Excellence in Cognition and its Disorders Memory Program CE10001021). Simon Lewis was supported by an NHMRC-ARC Dementia Fellowship (1110414) and Glenda Halliday was supported by an NHMRC Fellowship (1079679). The Queensland Parkinson’s Project (QPP) was supported by a grant from the Australian National Health and Medical Research Council (1084560) to George Mellick. The New Zealand Brain Research Institute (NZBRI) cohort was funded by a University of Otago Research Grant, together with financial support from the Jim and Mary Carney Charitable Trust (Whangarei, New Zealand). We thank Allison Miller for processing and handling of NZBRI samples.

**Additional acknowledgement for use of LD Hub**

We gratefully acknowledge all the studies and databases that made GWAS summary data available: **ADIPOGen** (Adiponectin genetics consortium),**C4D**(Coronary Artery Disease Genetics Consortium),**CARDIoGRAM** (Coronary ARtery DIsease Genome wide Replication and Meta-analysis), **CKDGen** (Chronic Kidney Disease Genetics consortium), **dbGAP** (database of Genotypes and Phenotypes), **DIAGRAM** (DIAbetes Genetics Replication And Meta-analysis), **ENIGMA**(Enhancing Neuro Imaging Genetics through Meta Analysis), **EAGLE**(EArly Genetics & Lifecourse Epidemiology Eczema Consortium, excluding 23andMe),**EGG** (Early Growth Genetics Consortium), **GABRIEL**(A Multidisciplinary Study to Identify the Genetic and Environmental Causes of Asthma in the European Community), **GCAN** (Genetic Consortium for Anorexia Nervosa),**GEFOS** (GEnetic Factors for OSteoporosis Consortium),**GIANT** (Genetic Investigation of ANthropometric Traits), **GIS**(Genetics of Iron Status consortium), **GLGC** (Global Lipids Genetics Consortium), **GPC** (Genetics of Personality Consortium), **GUGC**(Global Urate and Gout consortium), **HaemGen**(haemotological and platelet traits genetics consortium), **HRgene**(Heart Rate consortium), **IIBDGC** (International Inflammatory Bowel Disease Genetics Consortium),**ILCCO** (International Lung Cancer Consortium),**IMSGC** (International Multiple Sclerosis Genetic Consortium), **MAGIC** (Meta-Analyses of Glucose and Insulin-related traits Consortium),**MESA** (Multi-Ethnic Study of Atherosclerosis), **PGC** (Psychiatric Genomics Consortium), **Project MinE**consortium, **ReproGen** (Reproductive Genetics Consortium), **SSGAC (**Social Science Genetics Association Consortium) and**TAG** (Tobacco and Genetics Consortium), **TRICL**(Transdisciplinary Research in Cancer of the Lung consortium), **UK Biobank**. We gratefully acknowledge the contributions of Alkes Price (the systemic lupus erythematosus GWAS and primary biliary cirrhosis GWAS) and Johannes Kettunen (lipids metabolites GWAS).

**Additional acknowledgement for the use of the UKBB data**

This research has been conducted using the UK Biobank Resource under Application Number 33601.

**Additional 23andMe acknowledgement**

We thank the research participants and employees of 23andMe.
