## Supplementary figures and images for "Expanding Parkinson’s disease genetics: novel risk loci, genomic context, causal insights and heritable risk"

### Baylor_UMD_3phase.jpg

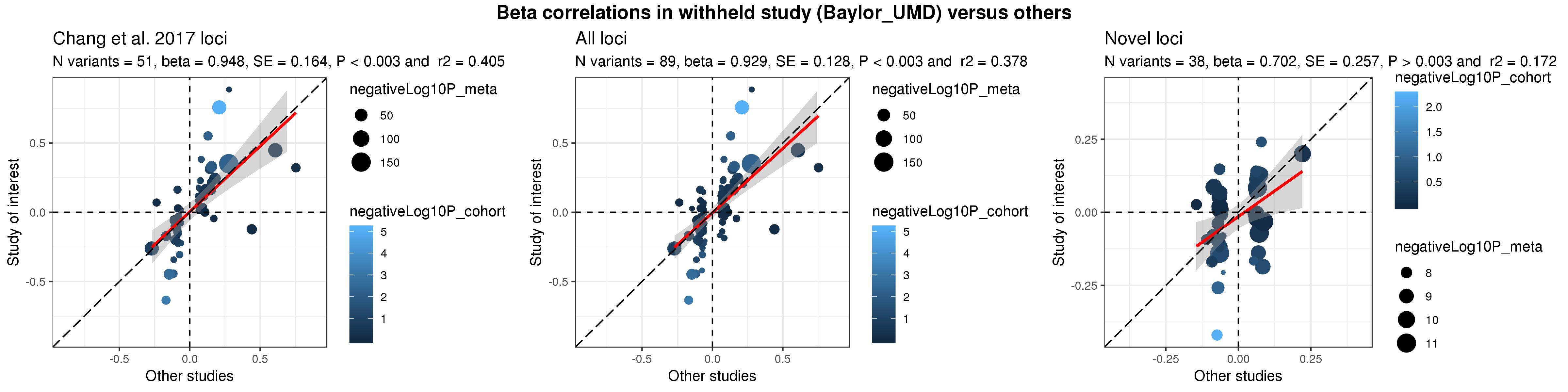

### chr1_154898185_rs114138760.pdf

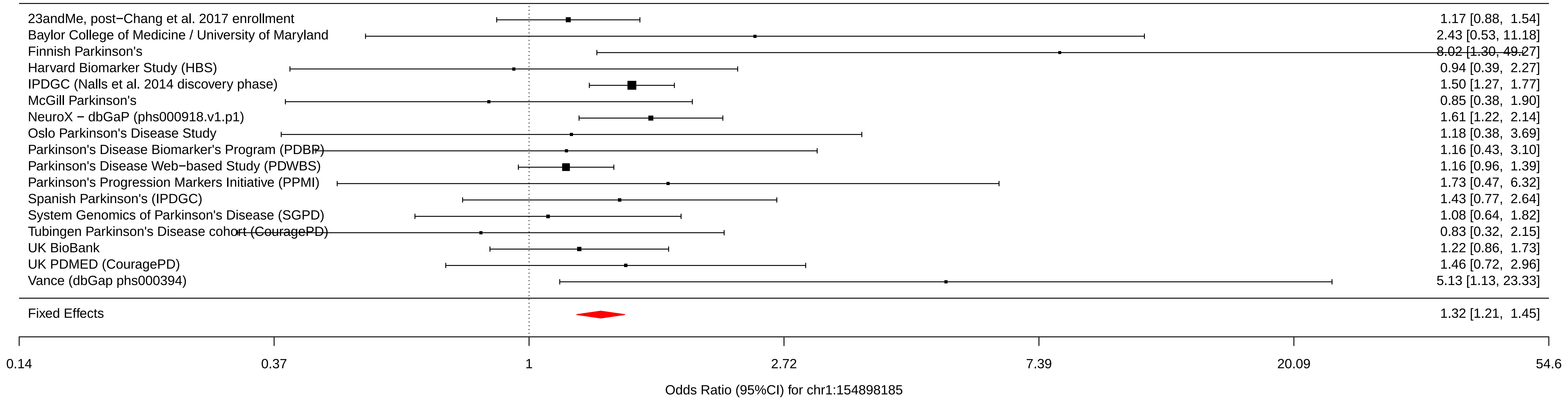

### chr1_155205634_rs76763715.pdf

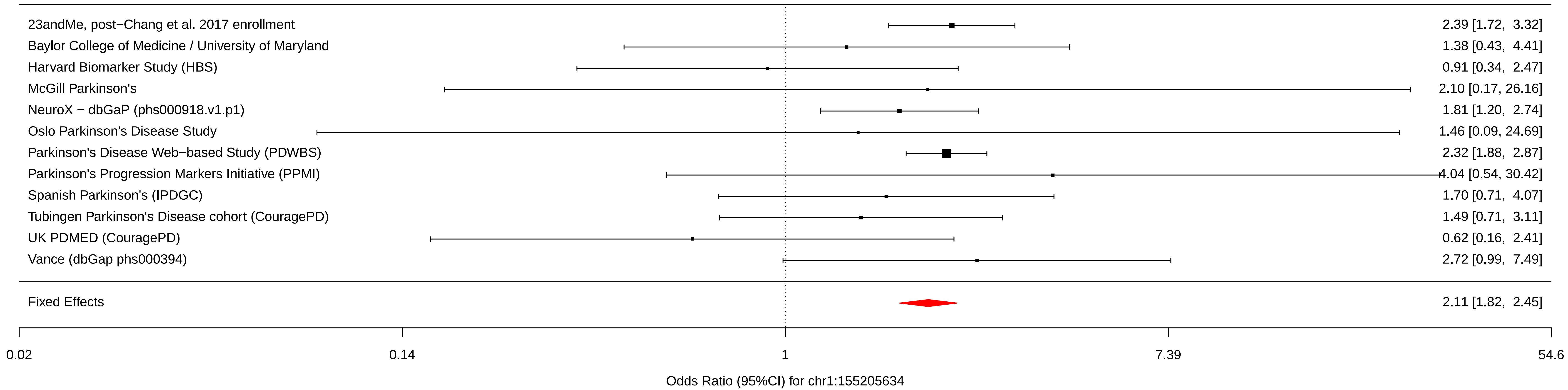

### chr1_161469054_rs6658353.pdf

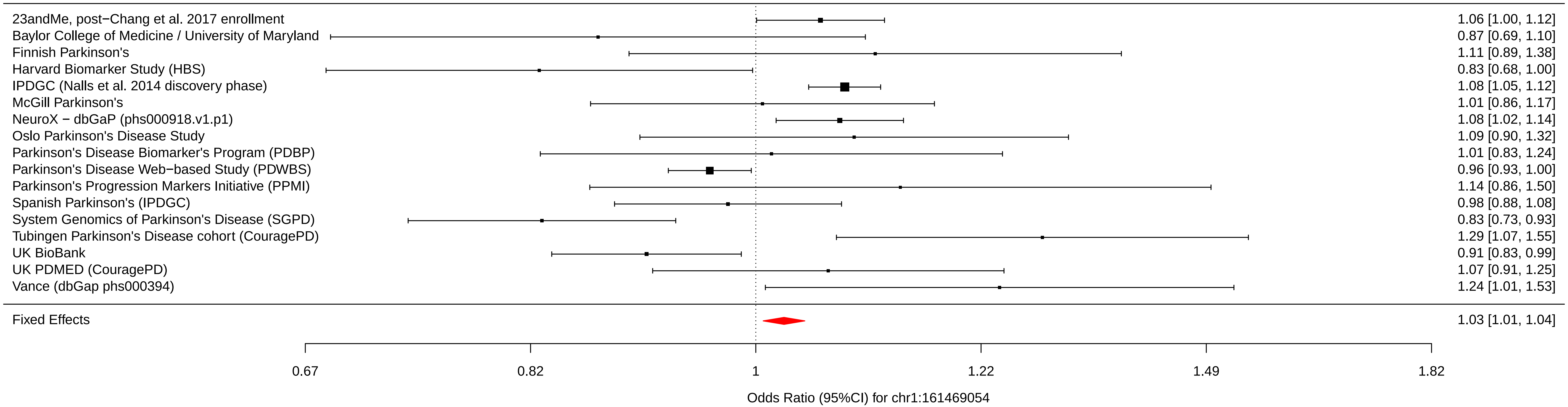

### chr1_171719769_rs11578699.pdf

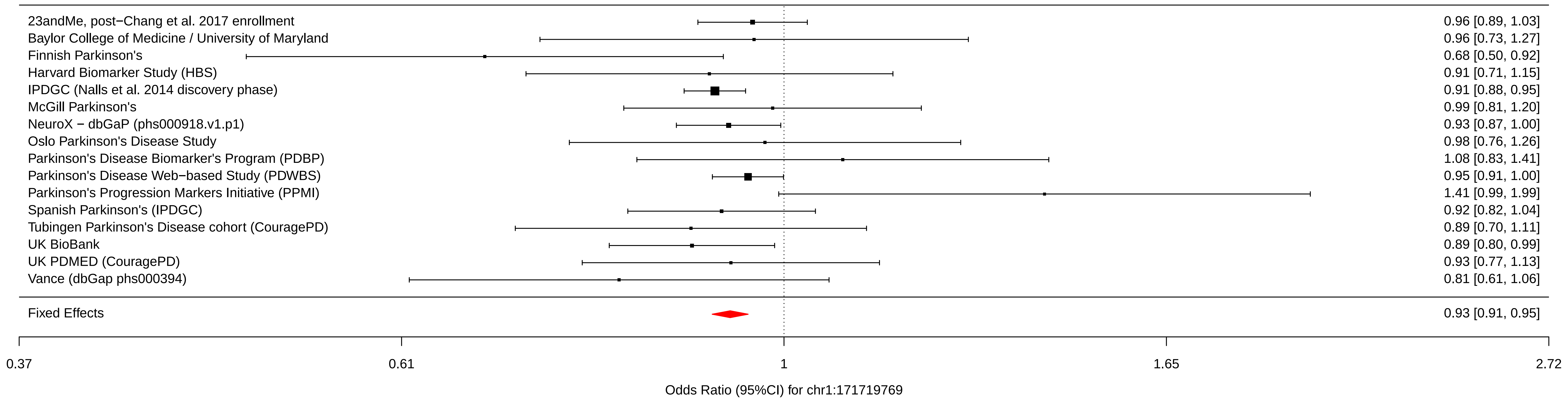

### chr1_205723572_rs823118.pdf

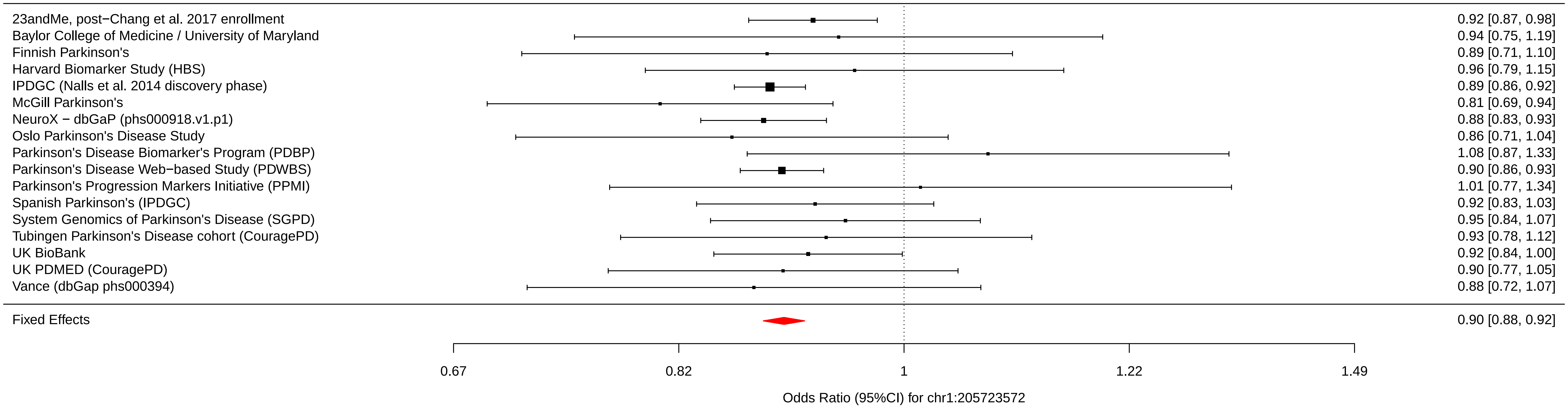

### chr1_205737739_rs11557080.pdf

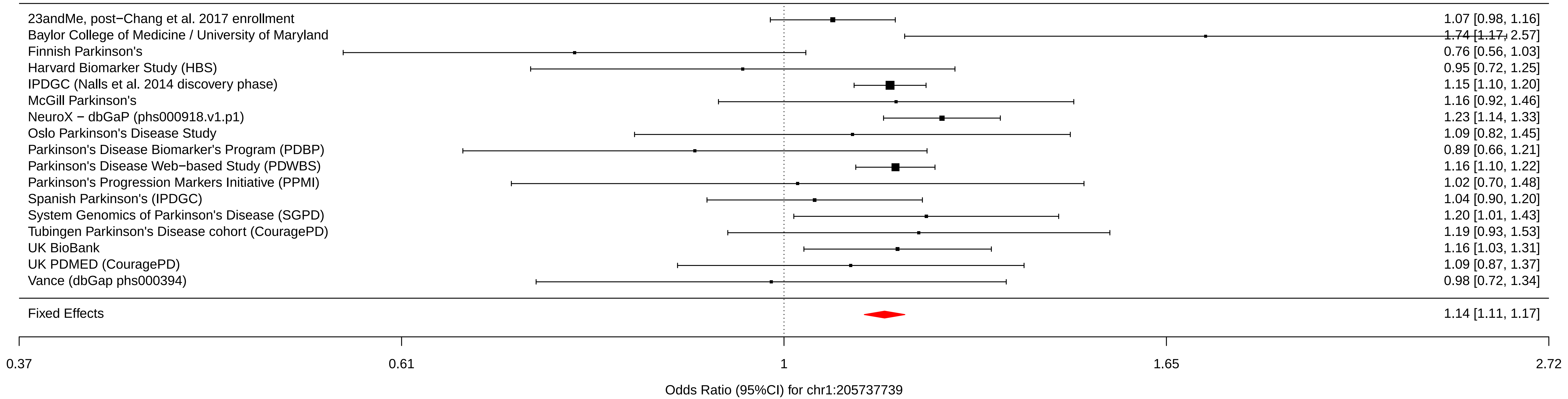

### chr1_226916078_rs4653767.pdf

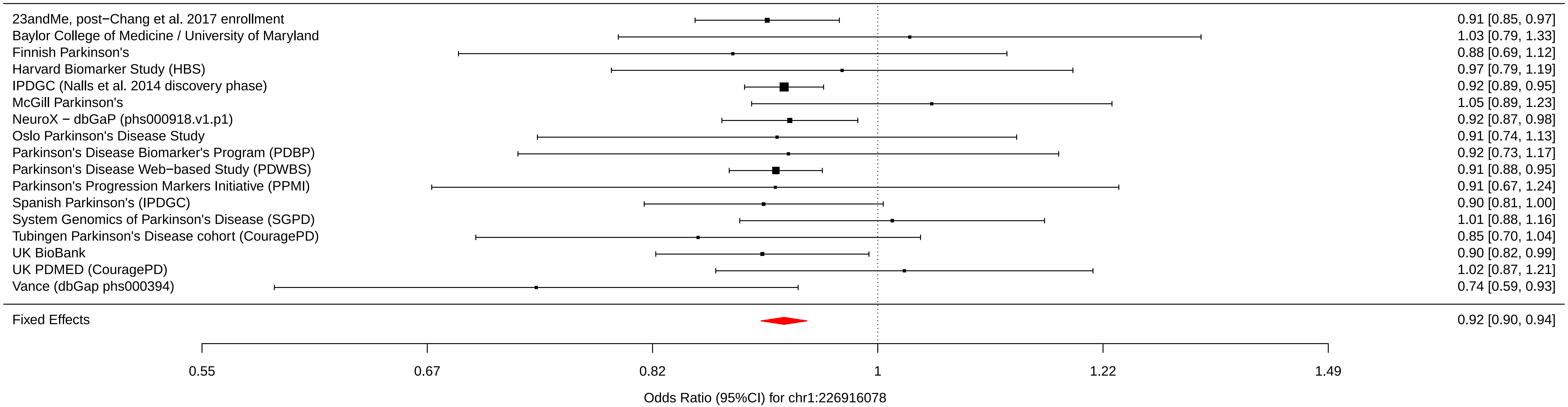

### chr1_232664611_rs10797576.pdf

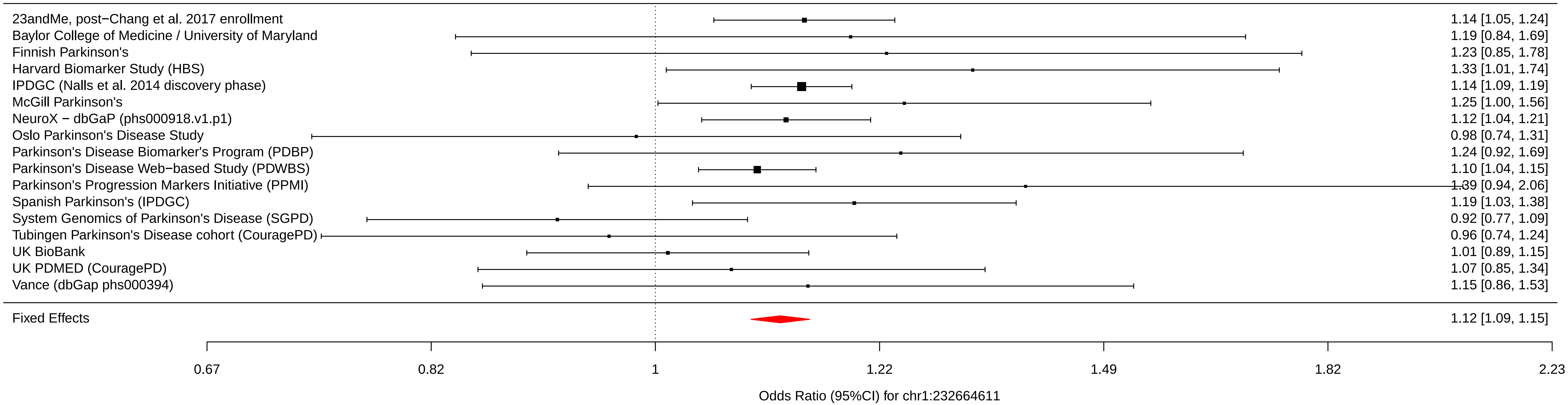

### chr2_18147848_rs76116224.pdf

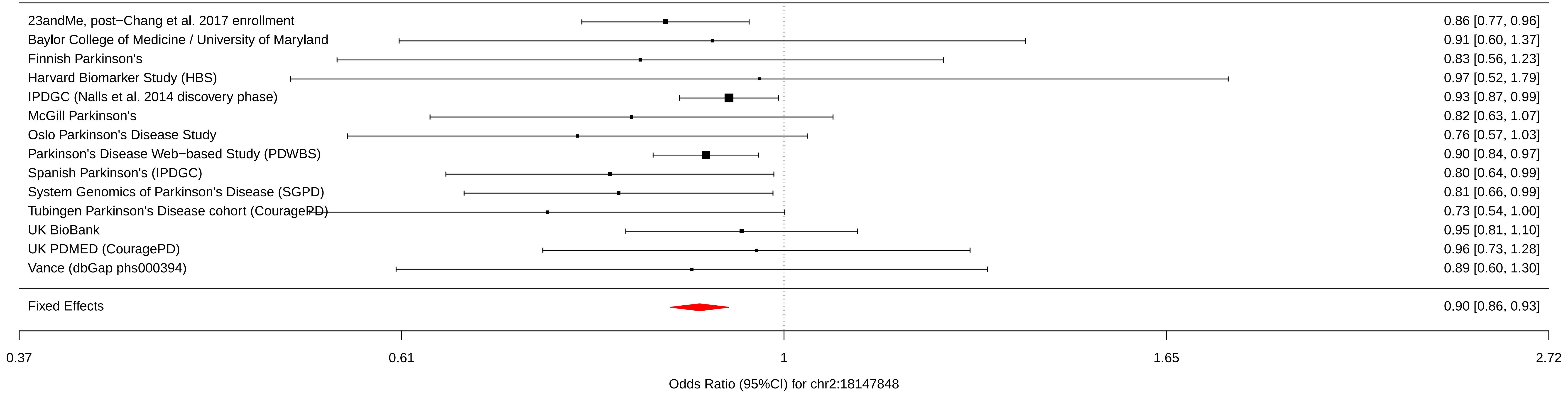

### chr2_96000943_rs2042477.pdf

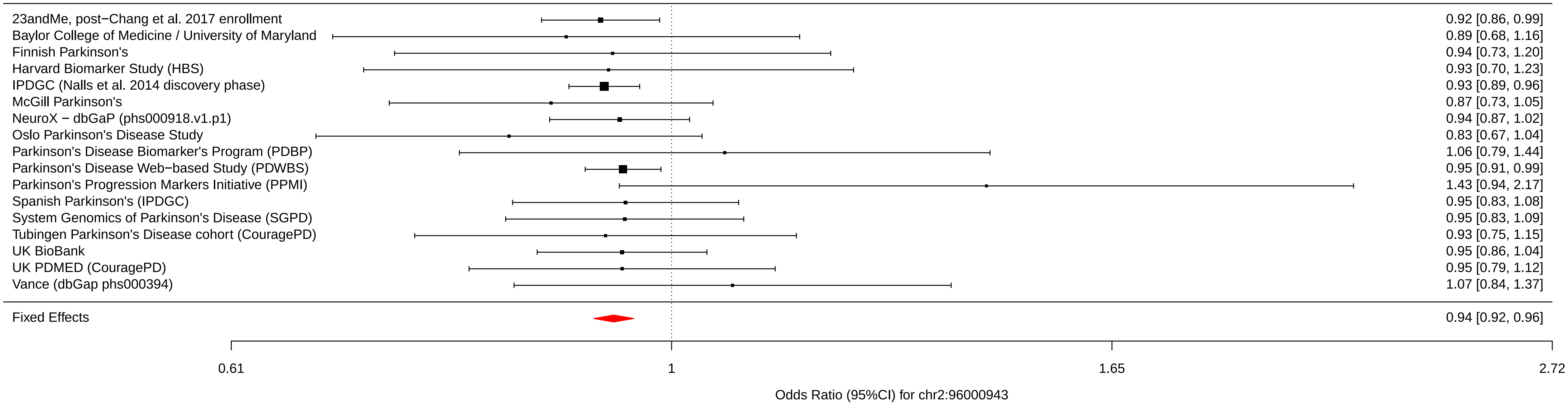

### chr2_102396963_rs11683001.pdf

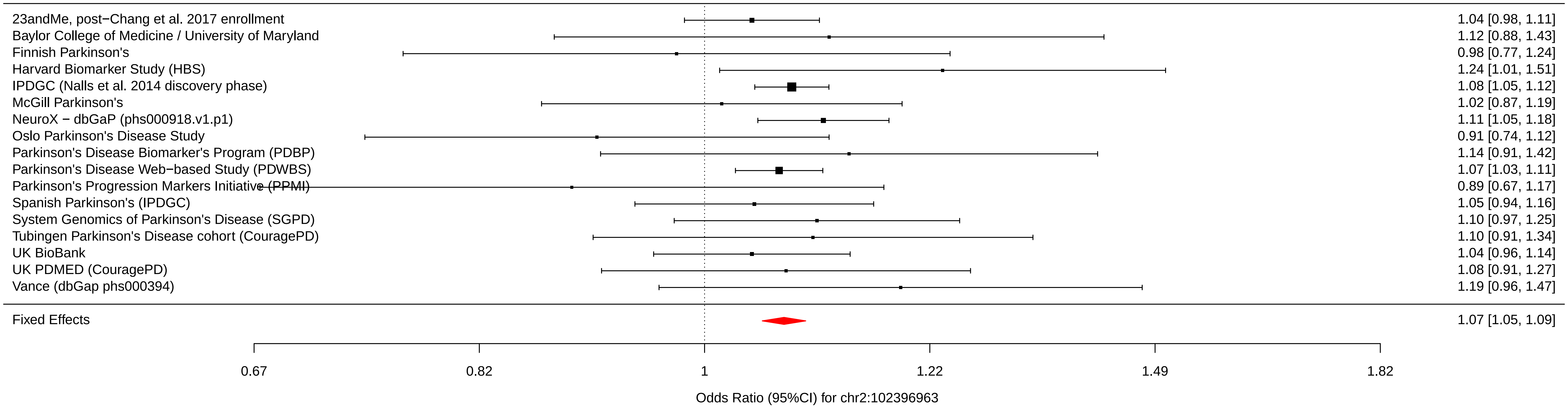

### chr2_135464616_rs57891859.pdf

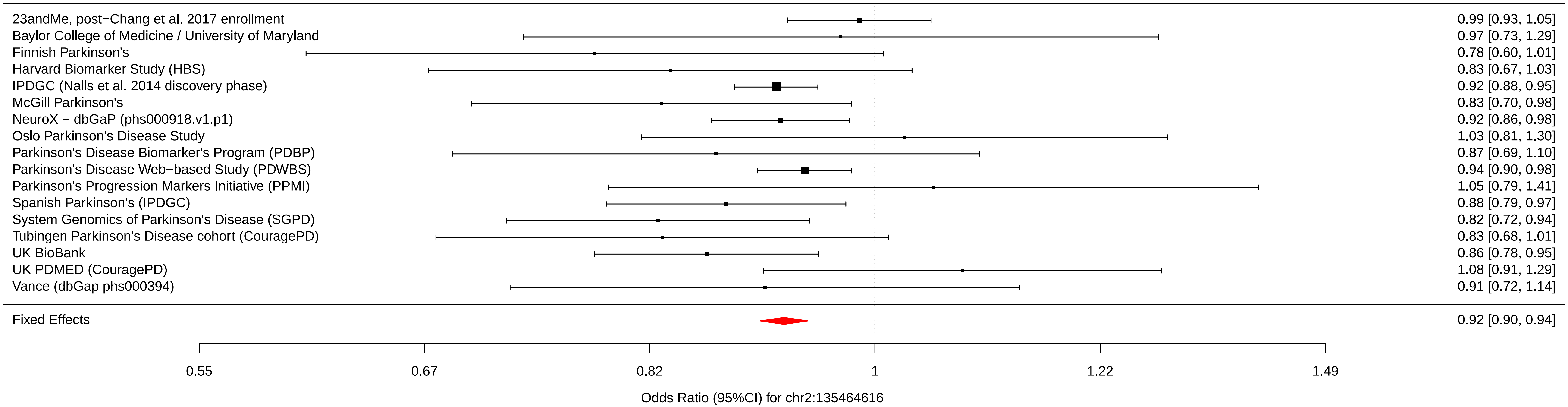

### chr3_18361759_rs73038319.pdf

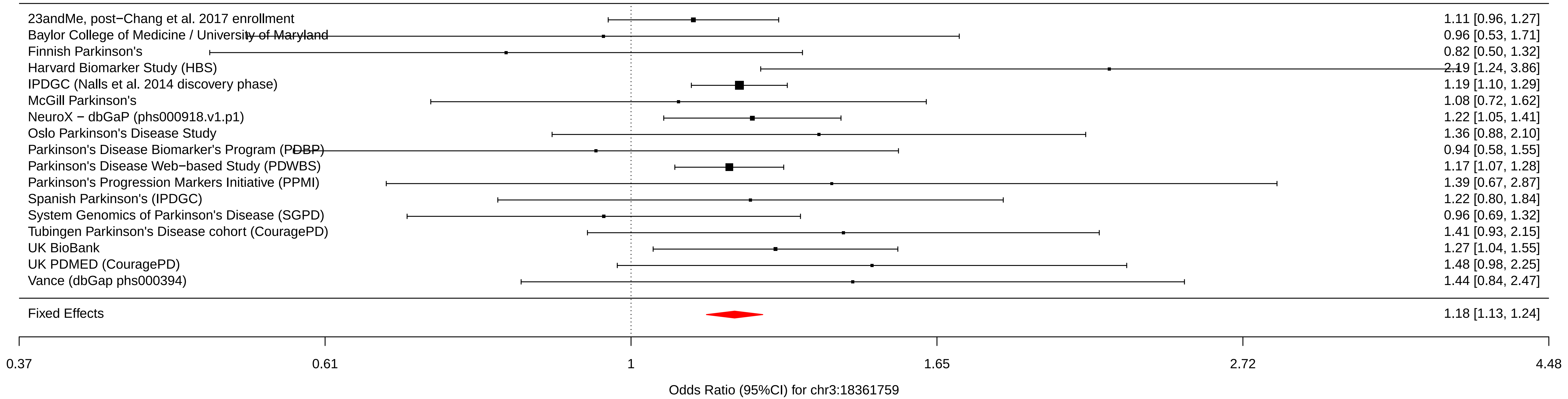

### chr3_48748989_rs12497850.pdf

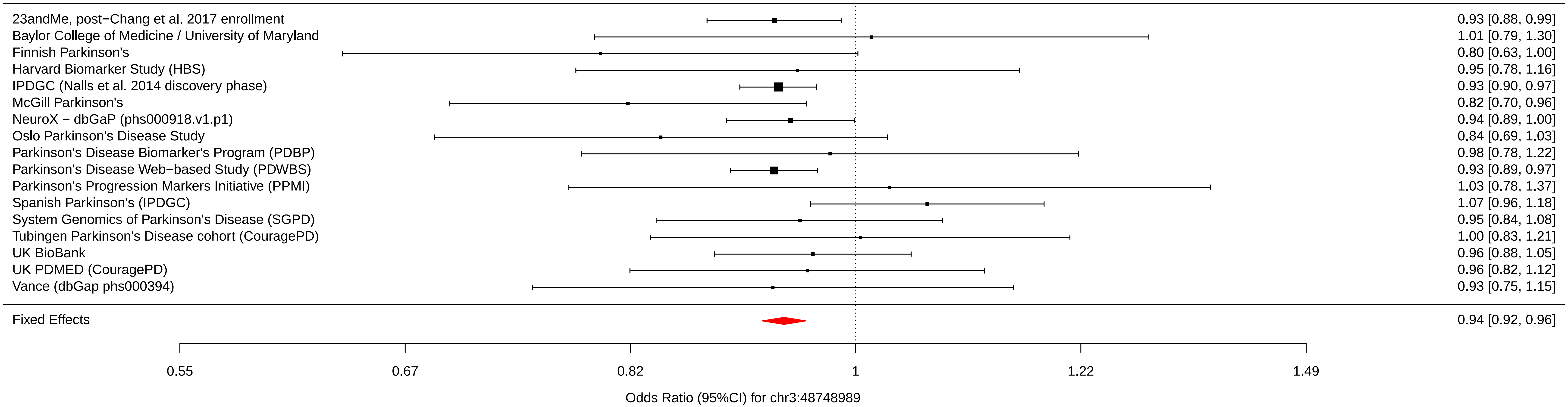

### chr3_122196892_rs55961674.pdf

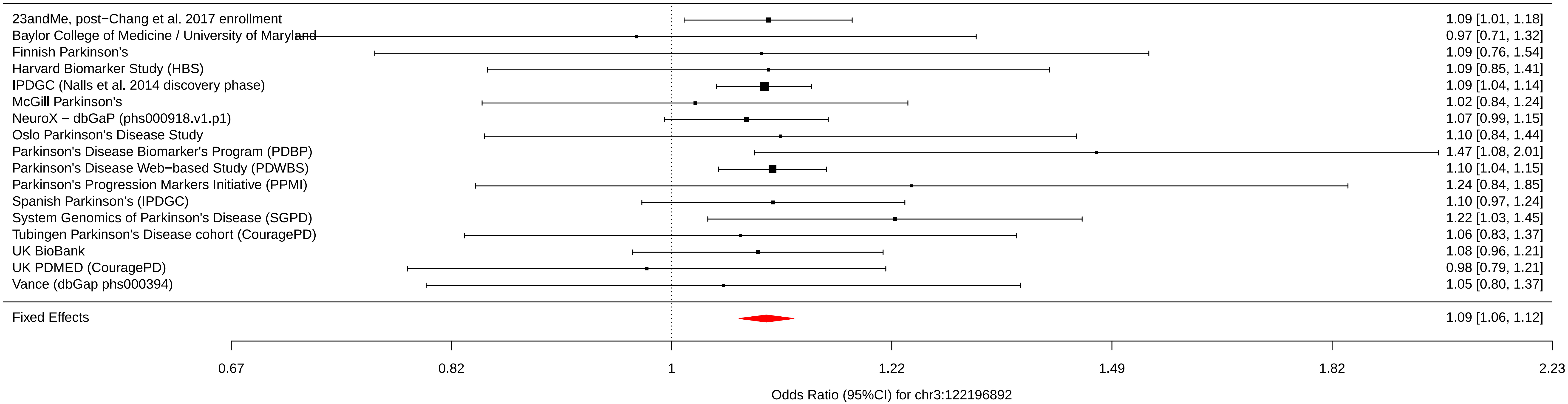

### chr3_151108965_rs11707416.pdf

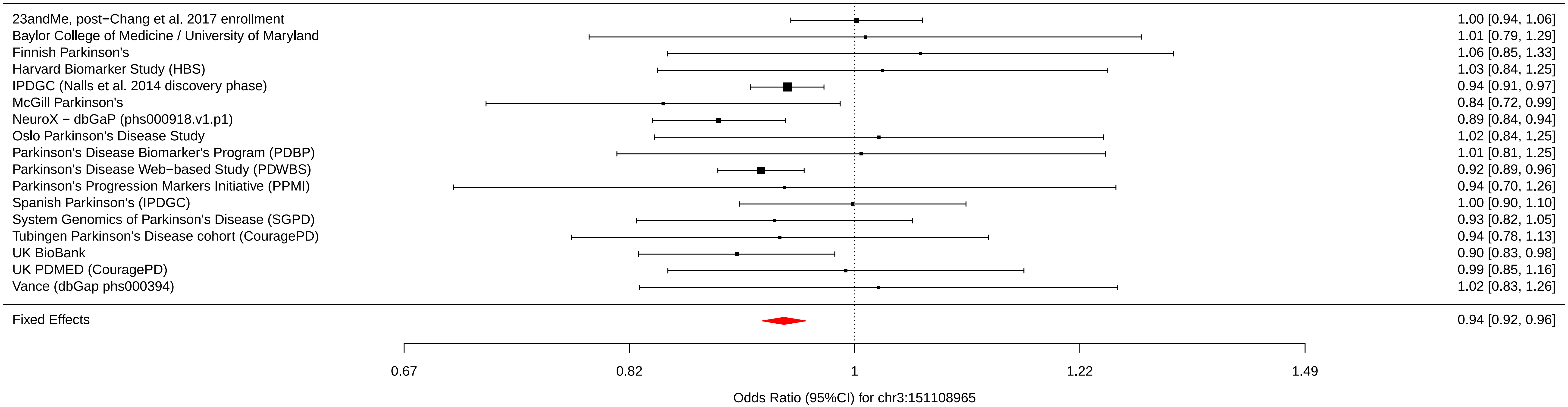

### chr3_161077630_rs1450522.pdf

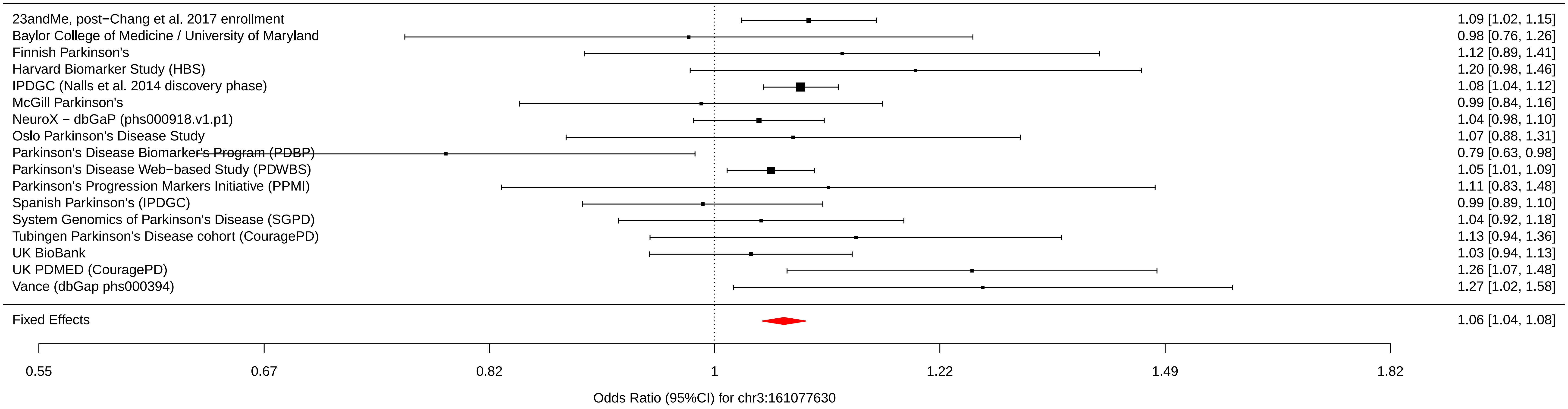

### chr3_182760073_rs10513789.pdf

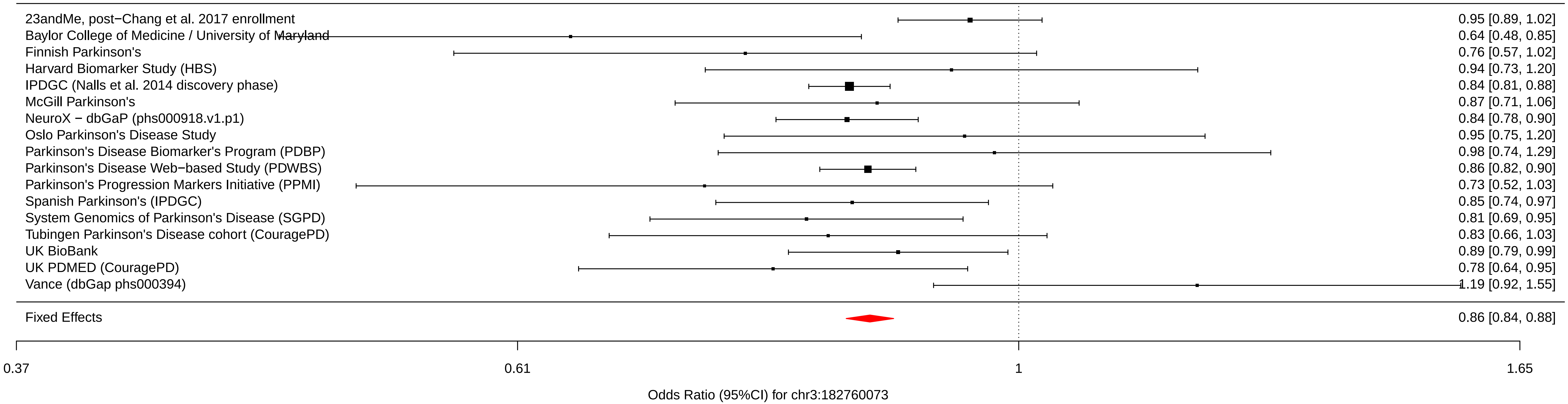

### chr4_925376_rs873786.pdf

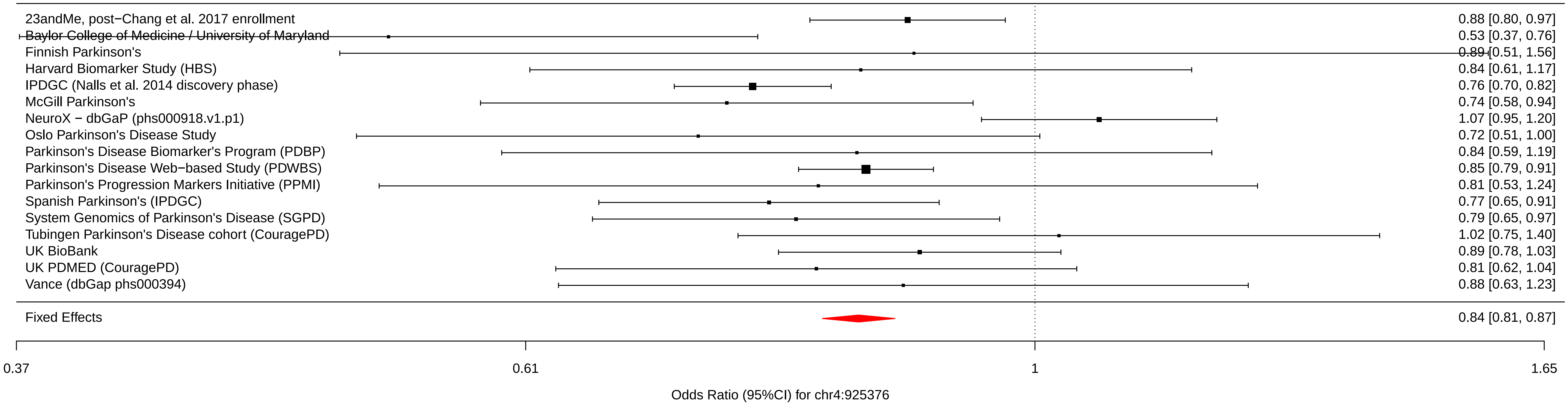

### chr4_951947_rs34311866.pdf

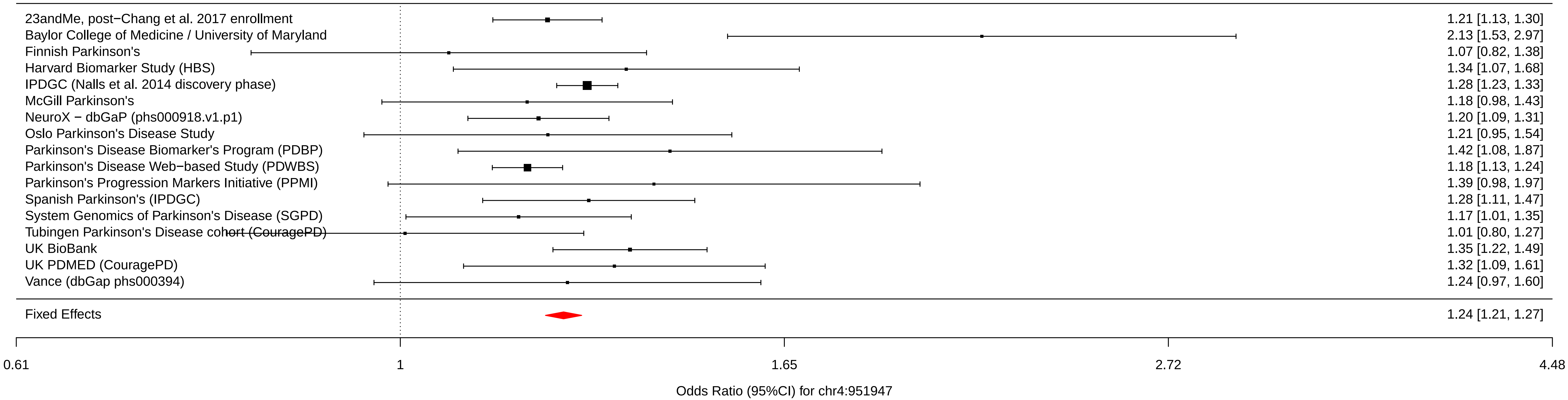

### chr4_15737348_rs4698412.pdf

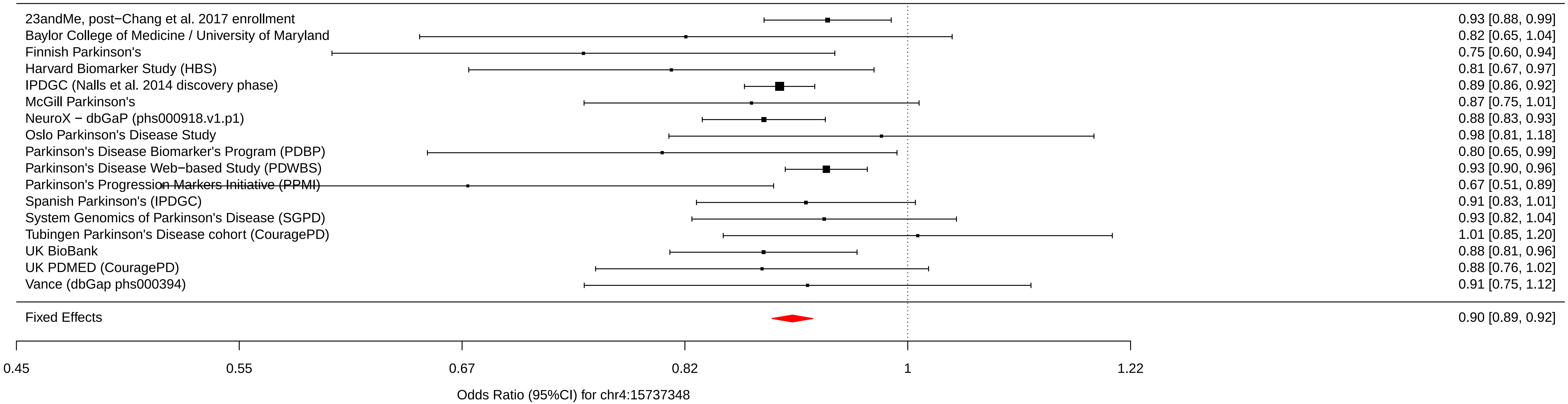

### chr4_17968811_rs34025766.pdf

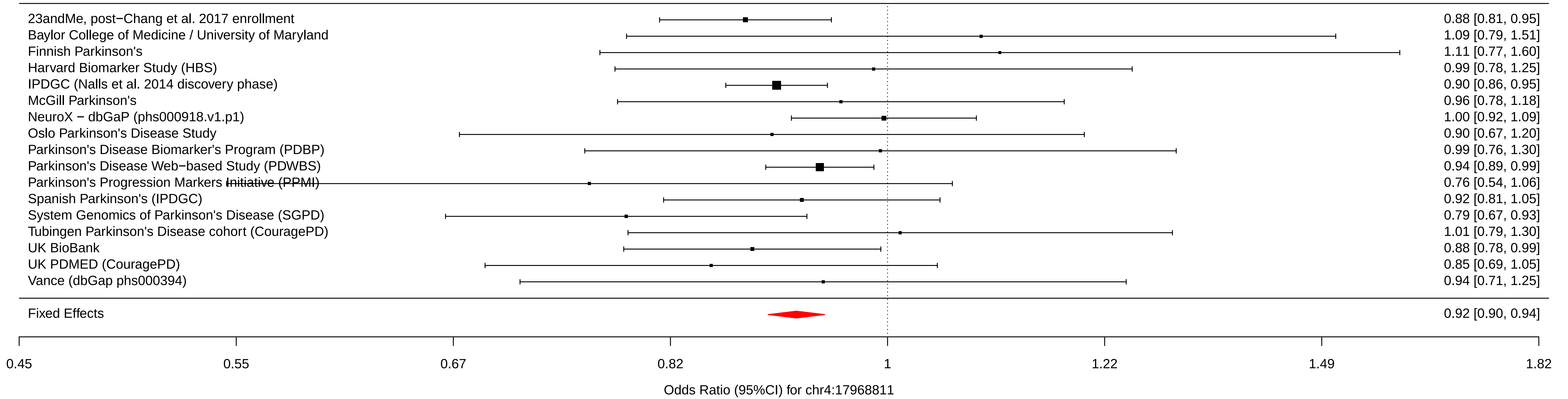

### chr4_77110365_rs6825004.pdf

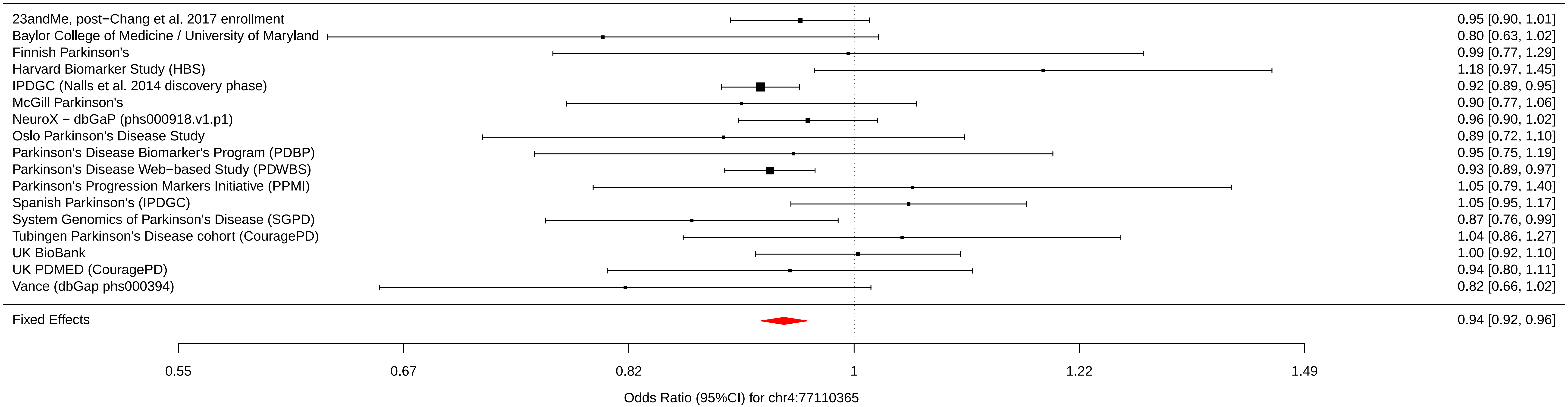

### chr4_77147969_rs4101061.pdf

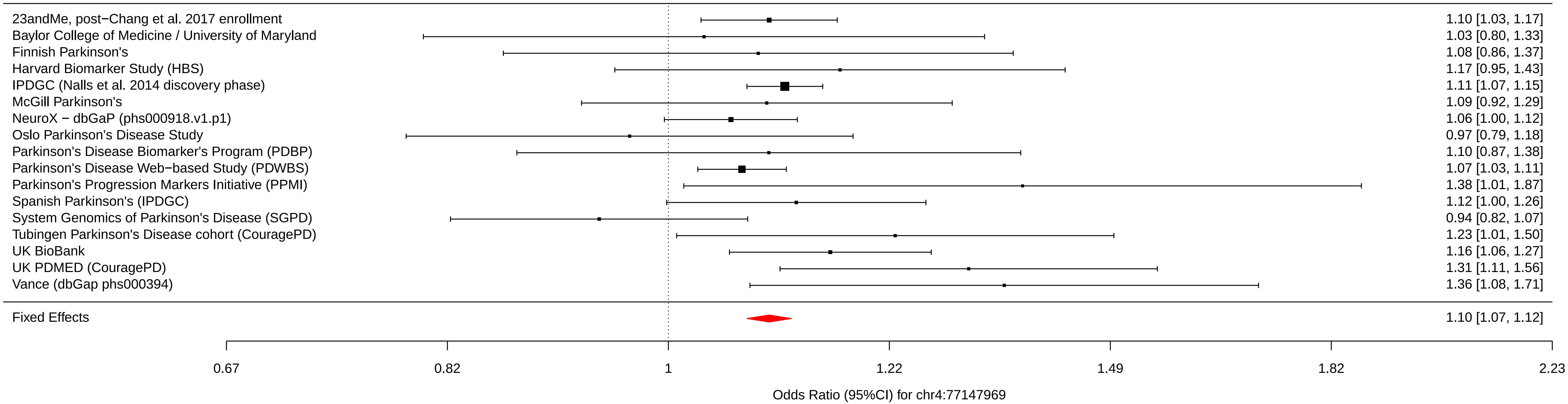

### chr4_77198054_rs6854006.pdf

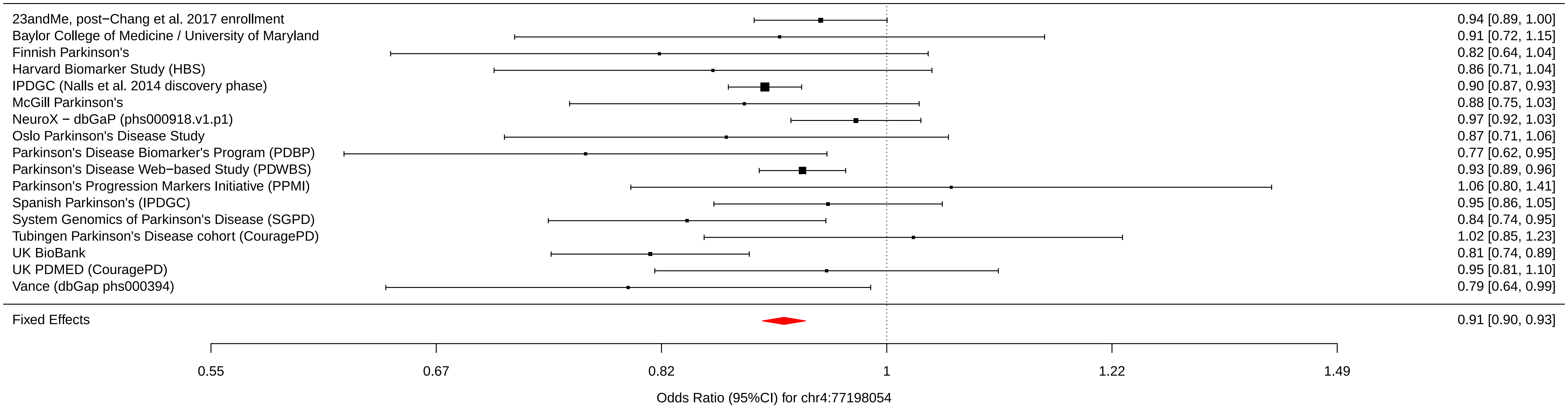

### chr4_90626111_rs356182.pdf

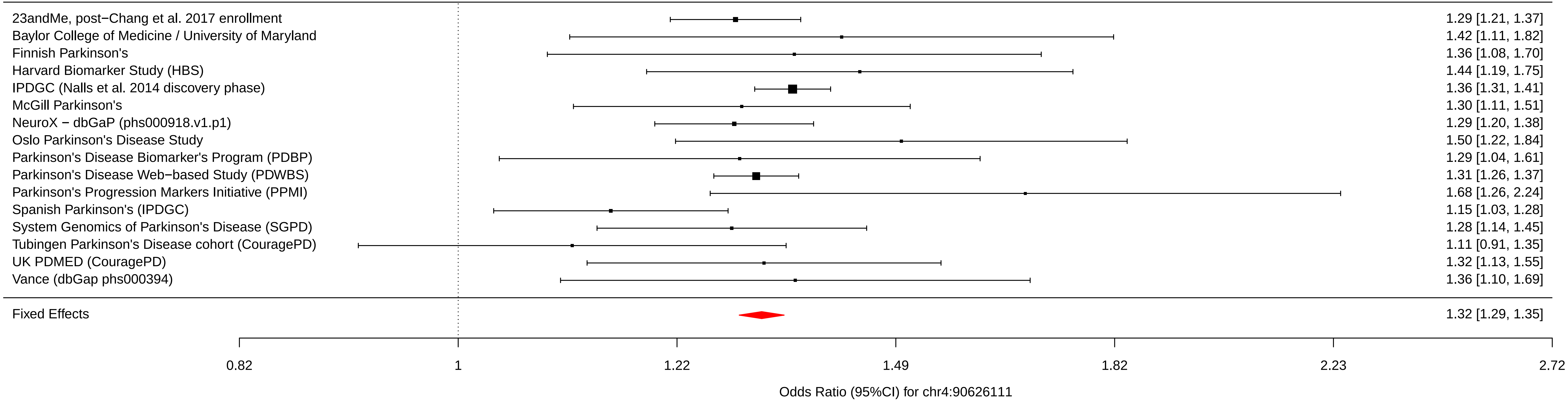

### chr4_90636630_rs5019538.pdf

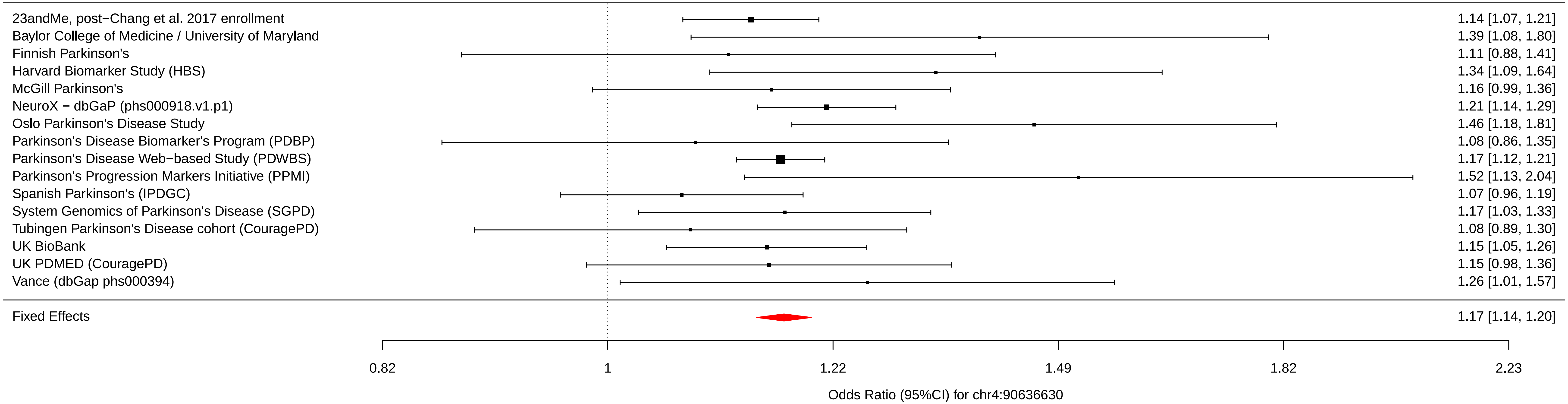

### chr5_60137959_rs1867598.pdf

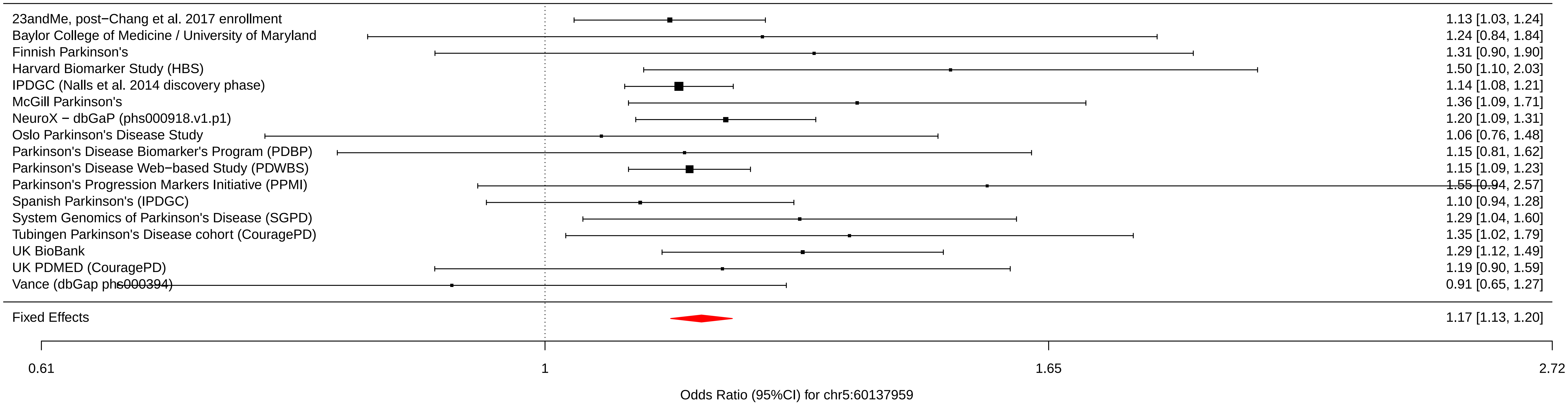

### chr5_102365794_rs26431.pdf

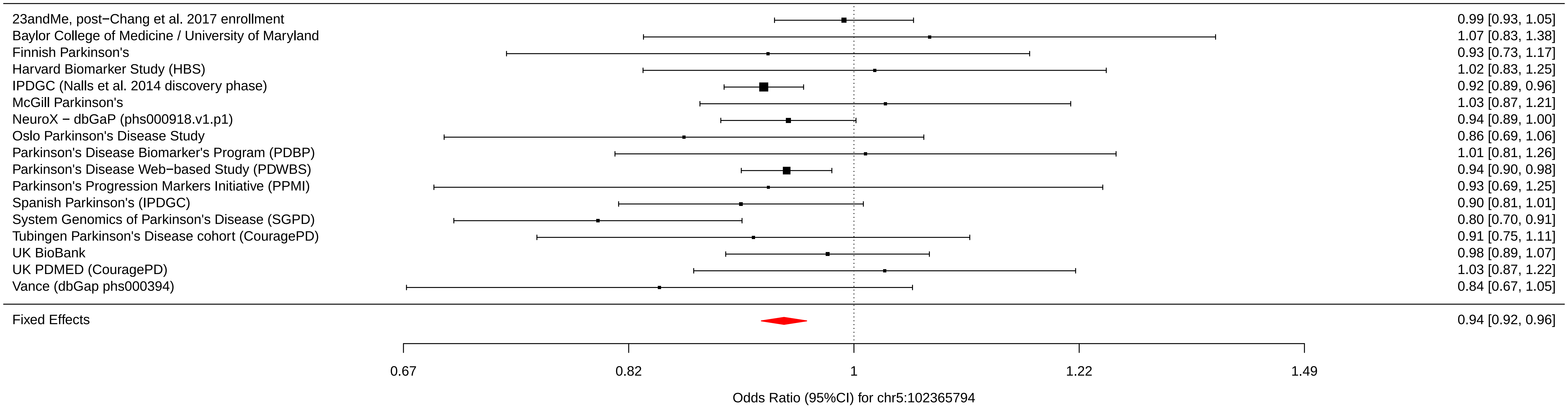

### chr5_134199105_rs11950533.pdf

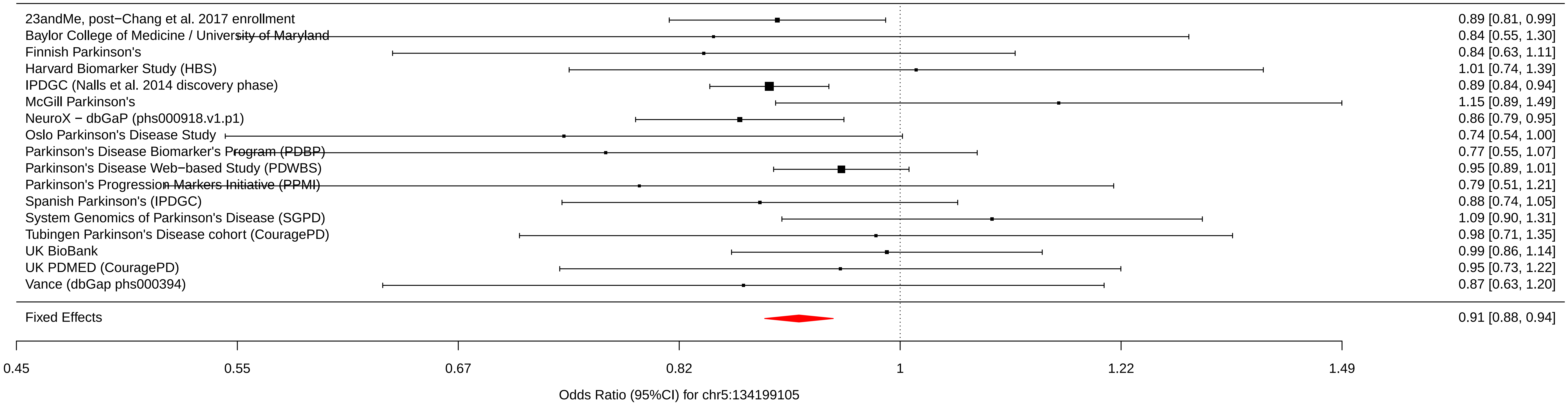

### chr6_27738801_rs4140646.pdf

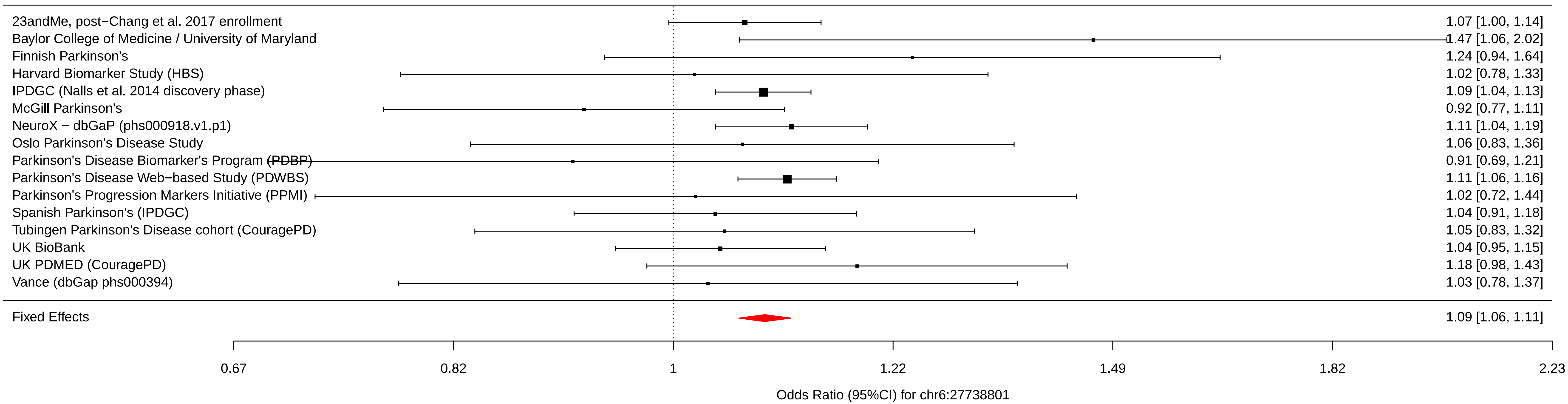

### chr6_72487762_rs12528068.pdf

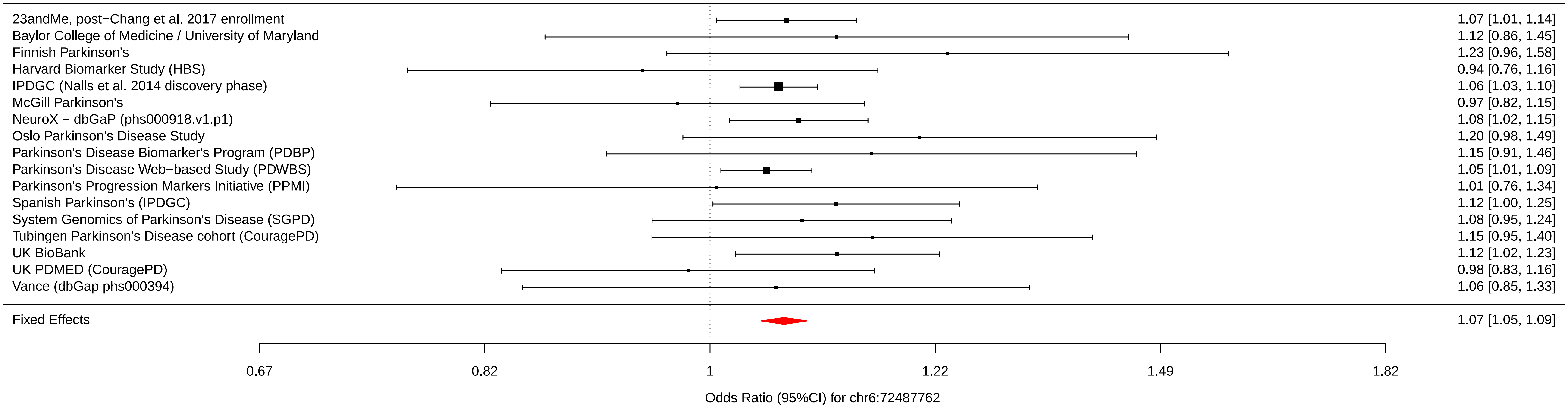

### chr6_112243291_rs997368.pdf

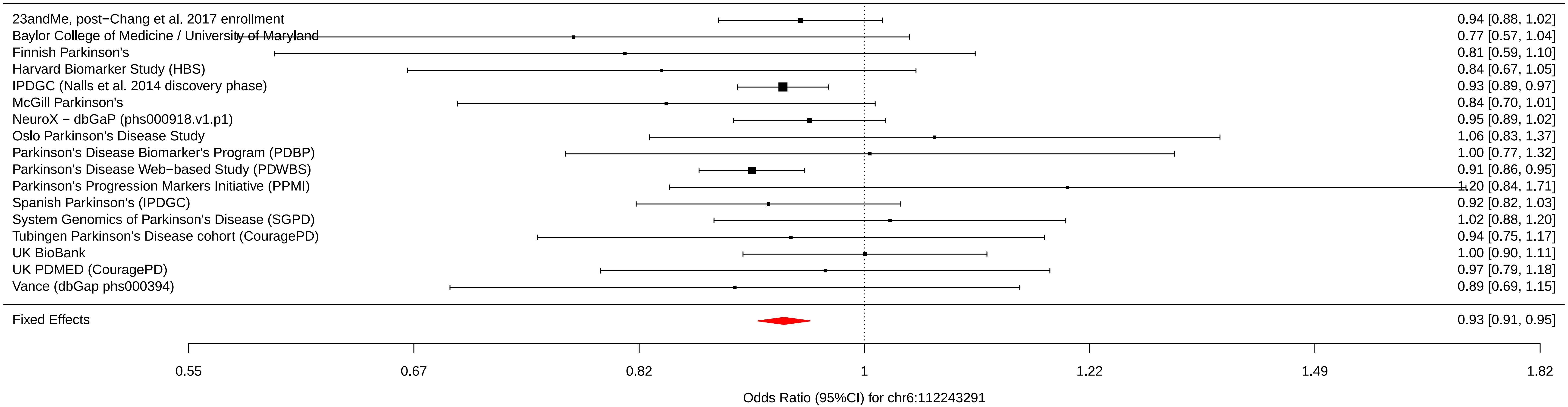

### chr6_133210361_rs75859381.pdf

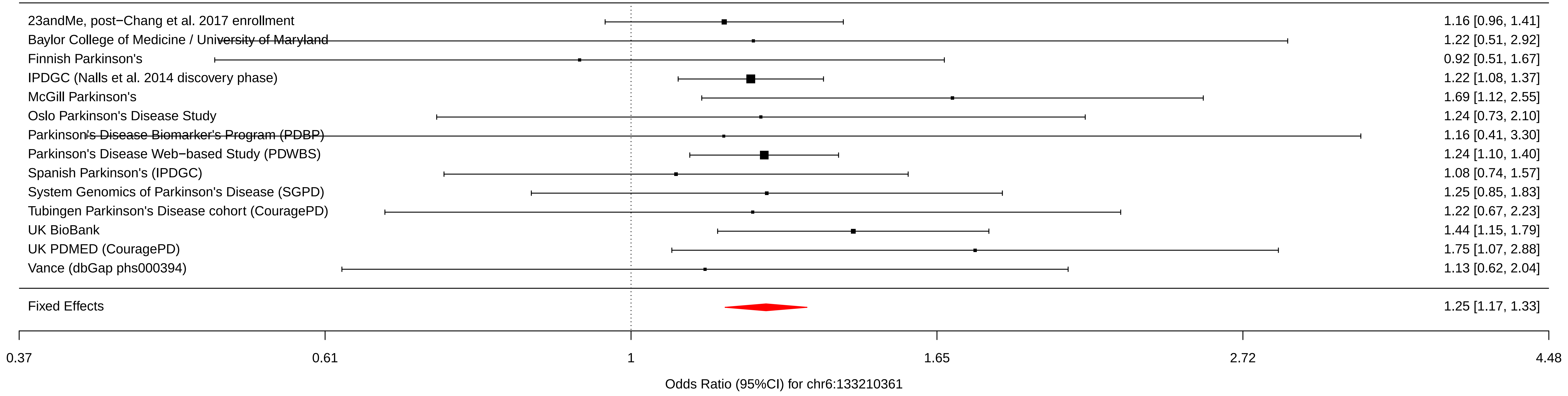

### chr7_23300049_rs199351.pdf

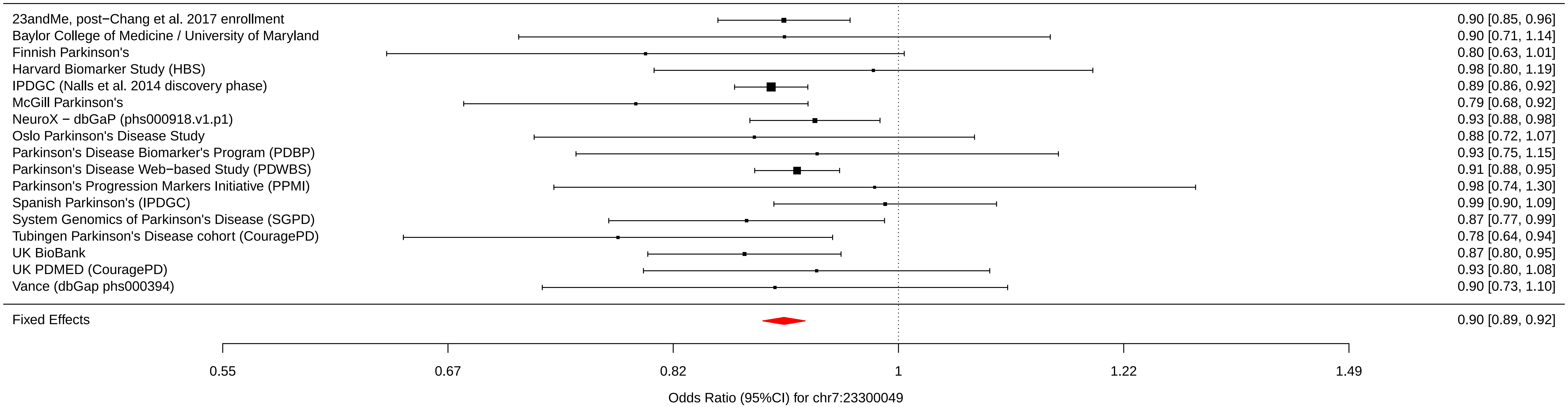

### chr7_66009851_rs76949143.pdf

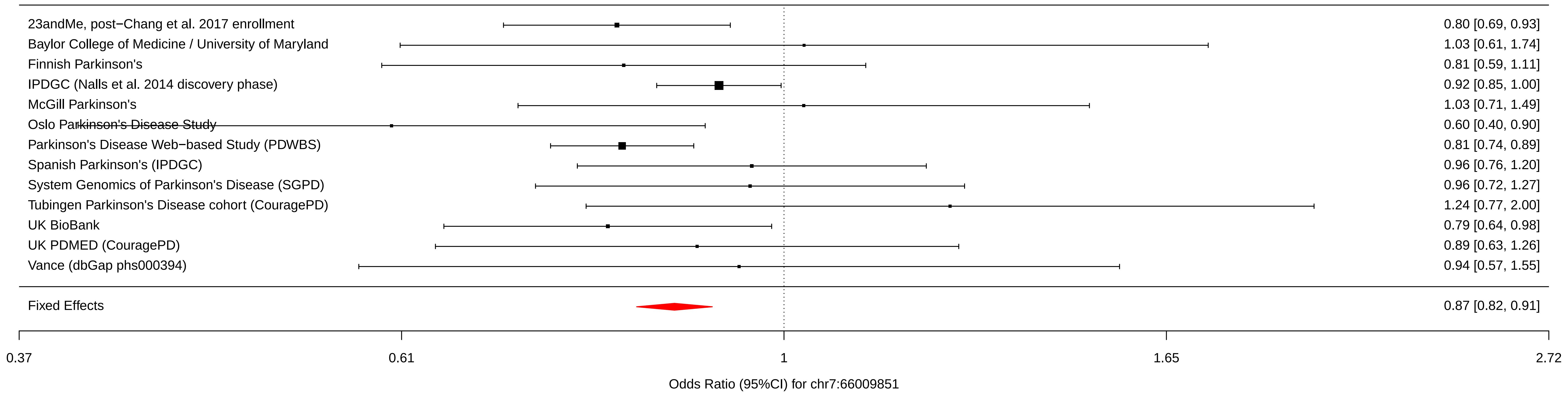

### chr8_11712443_rs1293298.pdf

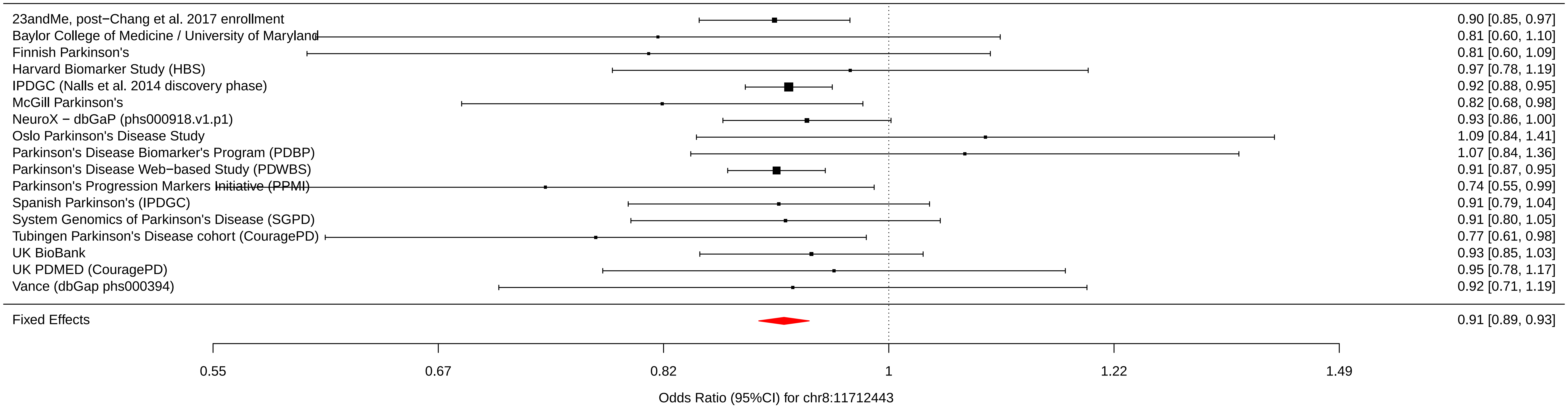

### chr8_16697593_rs620513.pdf

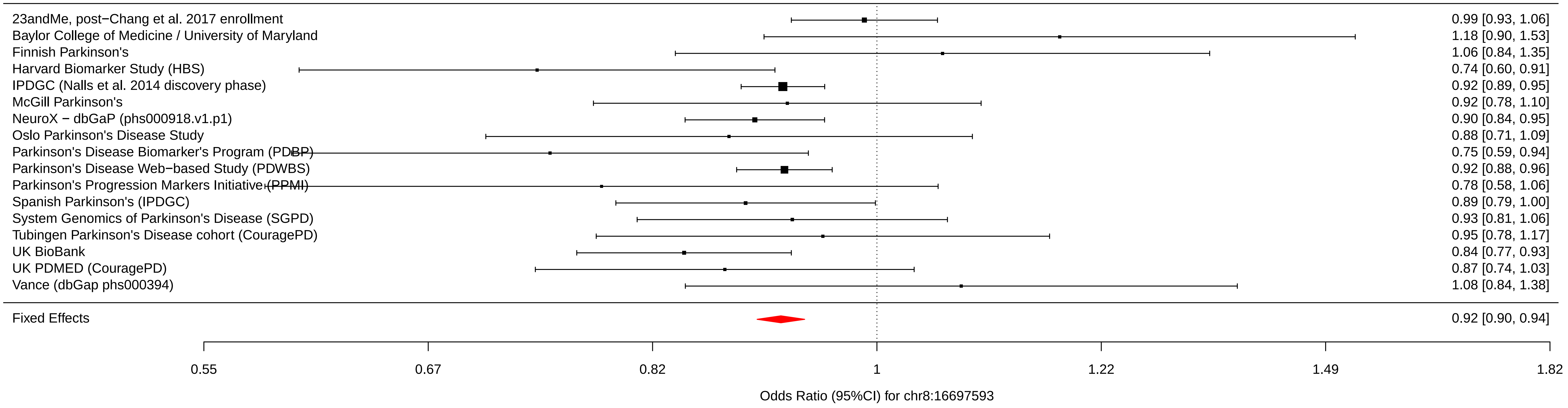

### chr8_22525980_rs2280104.pdf

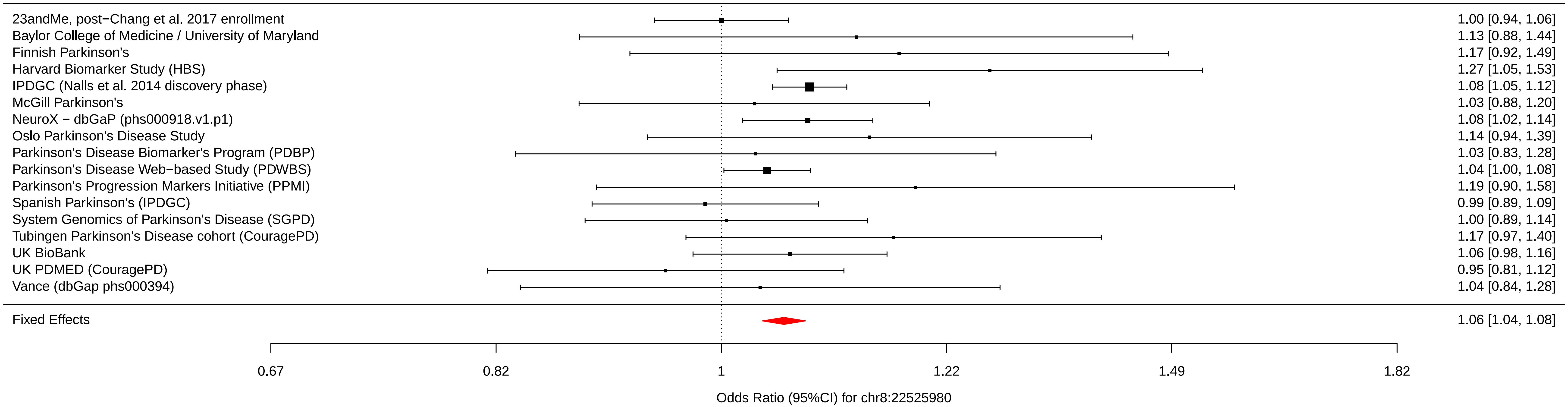

### chr8_130901909_rs2086641.pdf

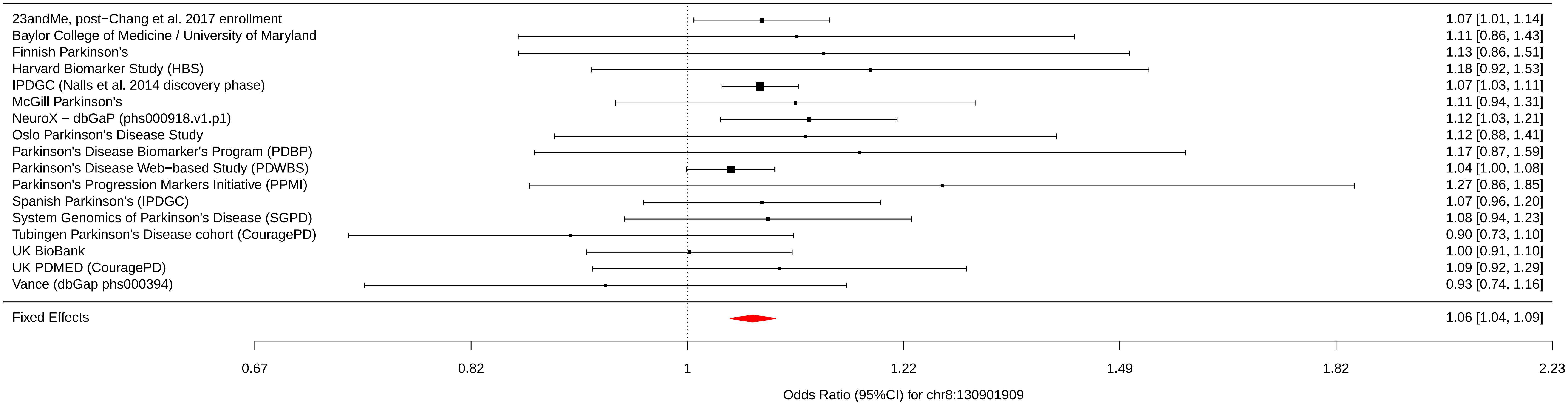

### chr9_17579690_rs13294100.pdf

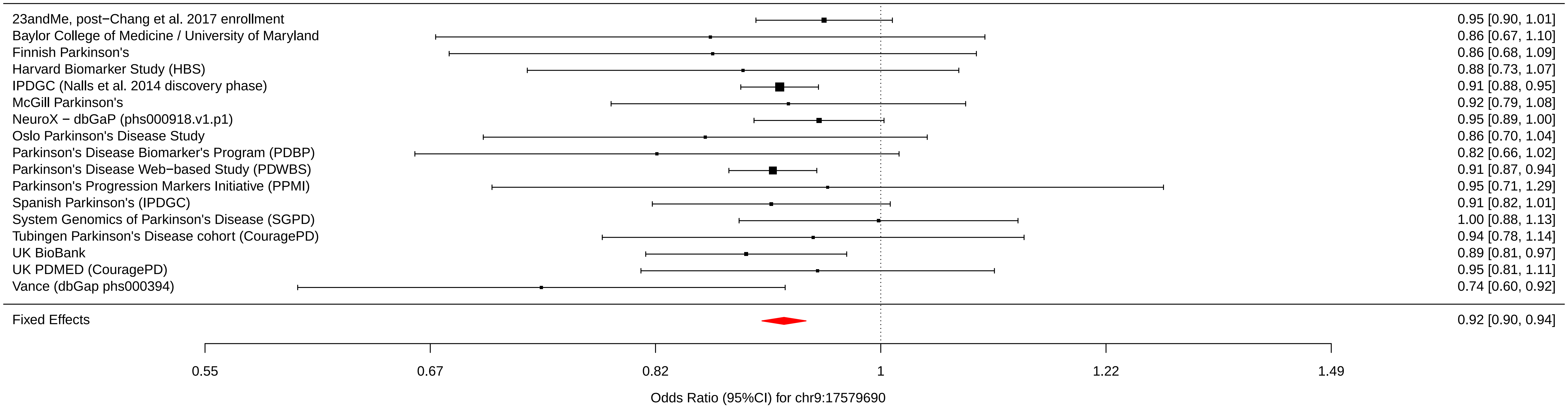

### chr9_17727065_rs10756907.pdf

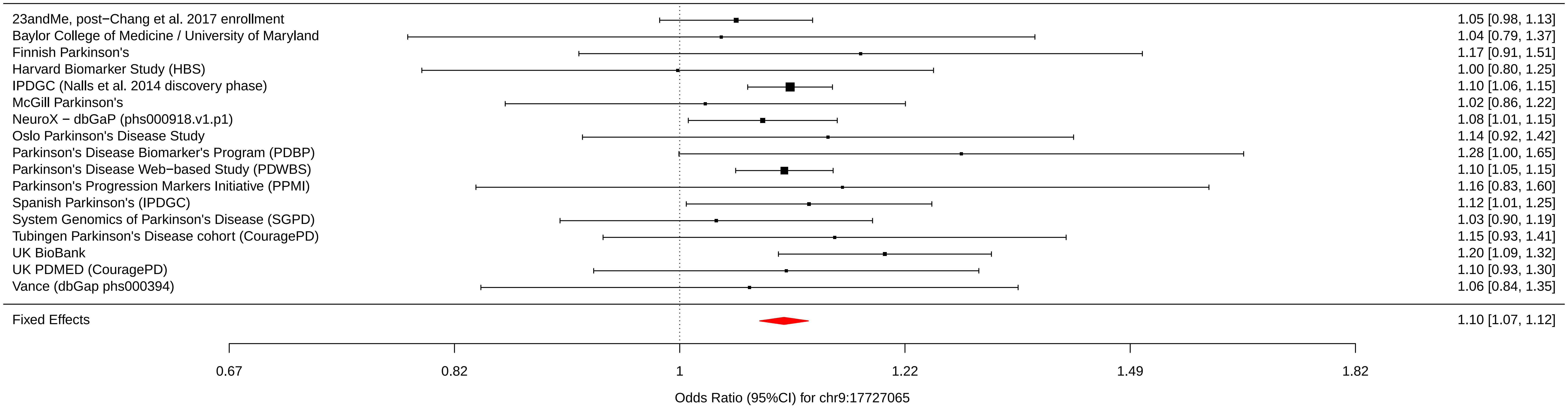

### chr9_34046391_rs6476434.pdf

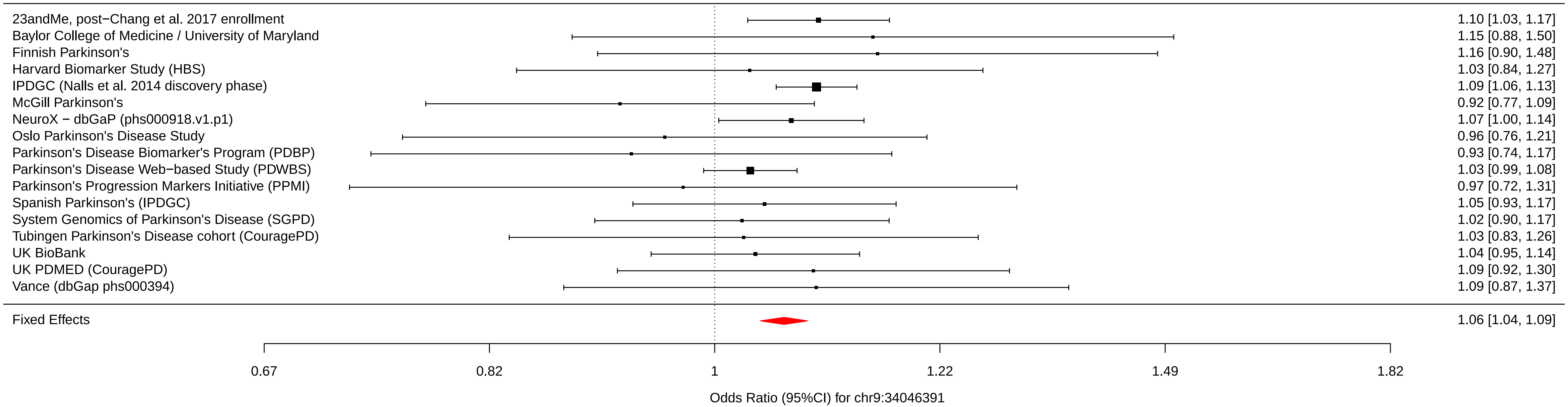

### chr10_15557406_rs896435.pdf

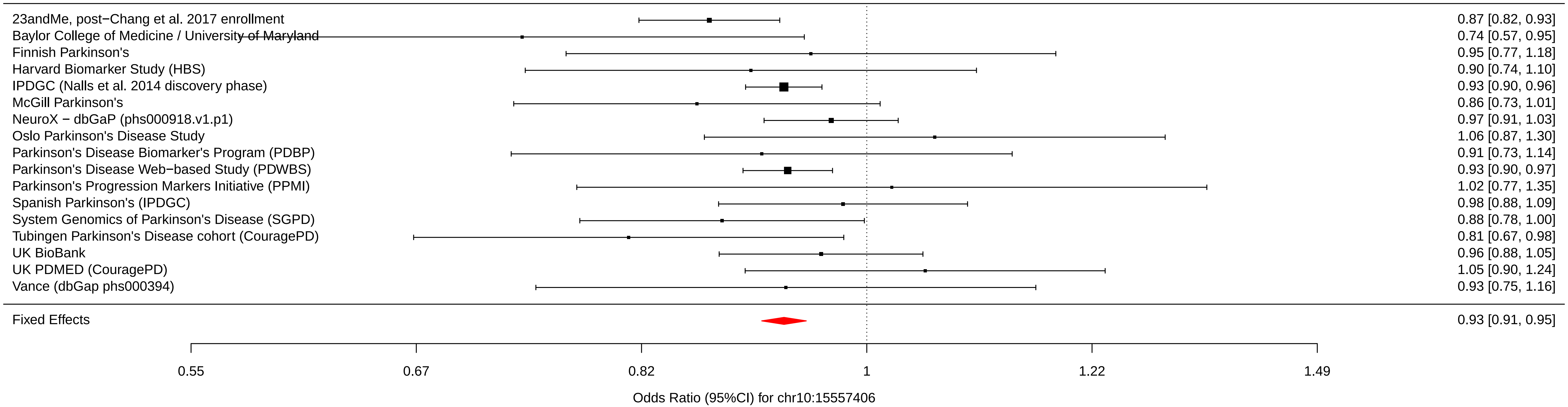

### chr10_104015279_rs10748818.pdf

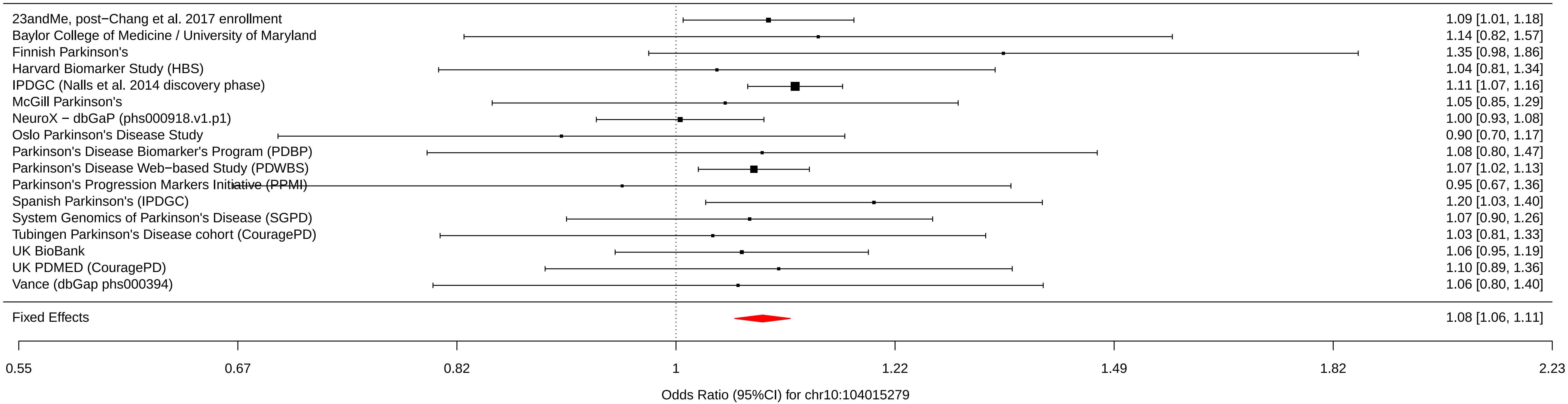

### chr10_121415685_rs72840788.pdf

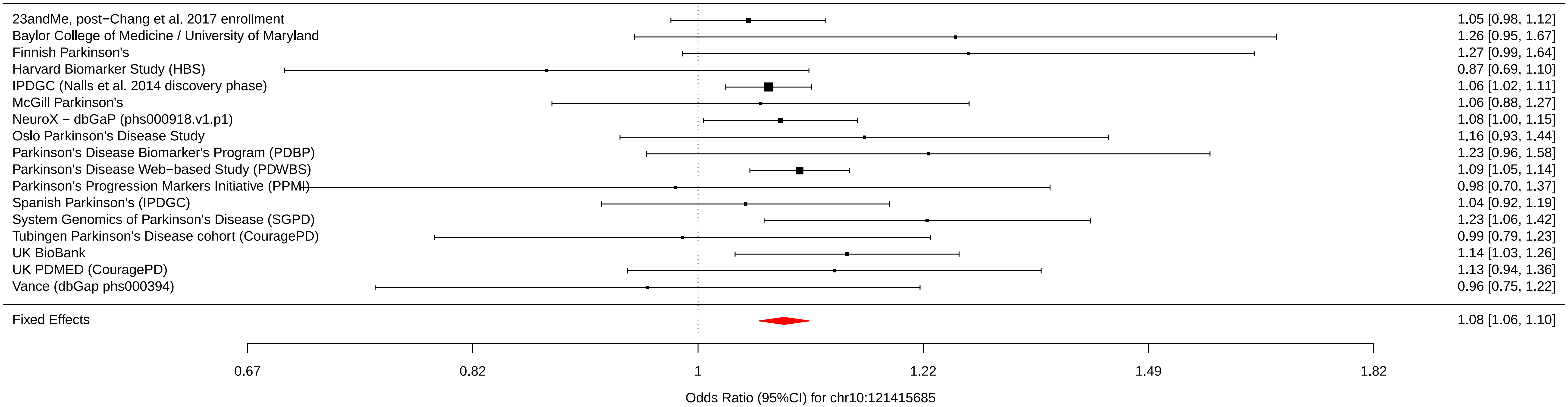

### chr10_121536327_rs117896735.pdf

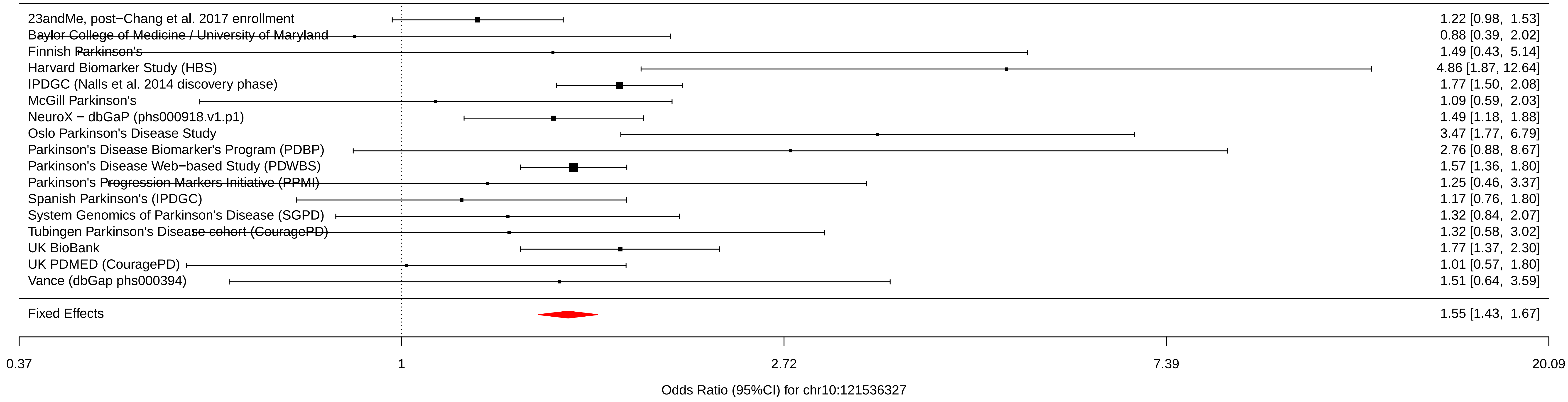

### chr11_10558777_rs7938782.pdf

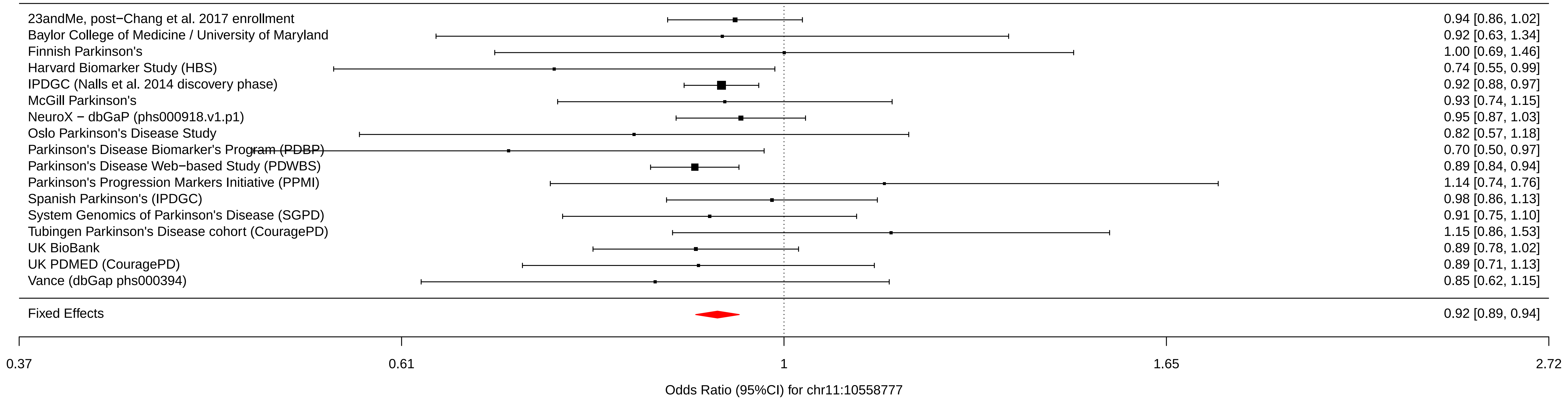

### chr11_83487277_rs12283611.pdf

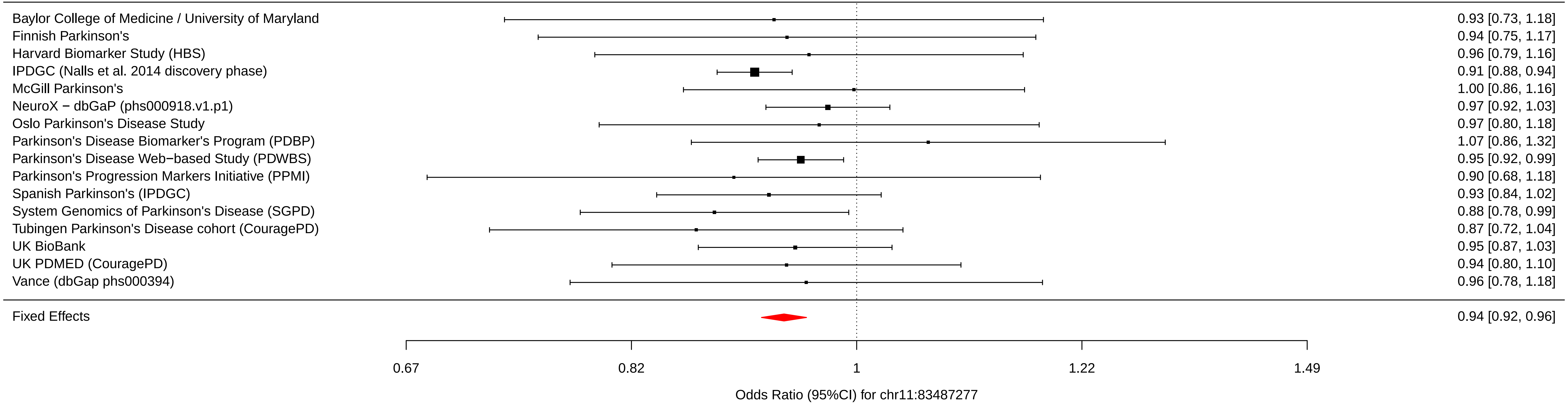

### chr11_133787001_rs3802920.pdf

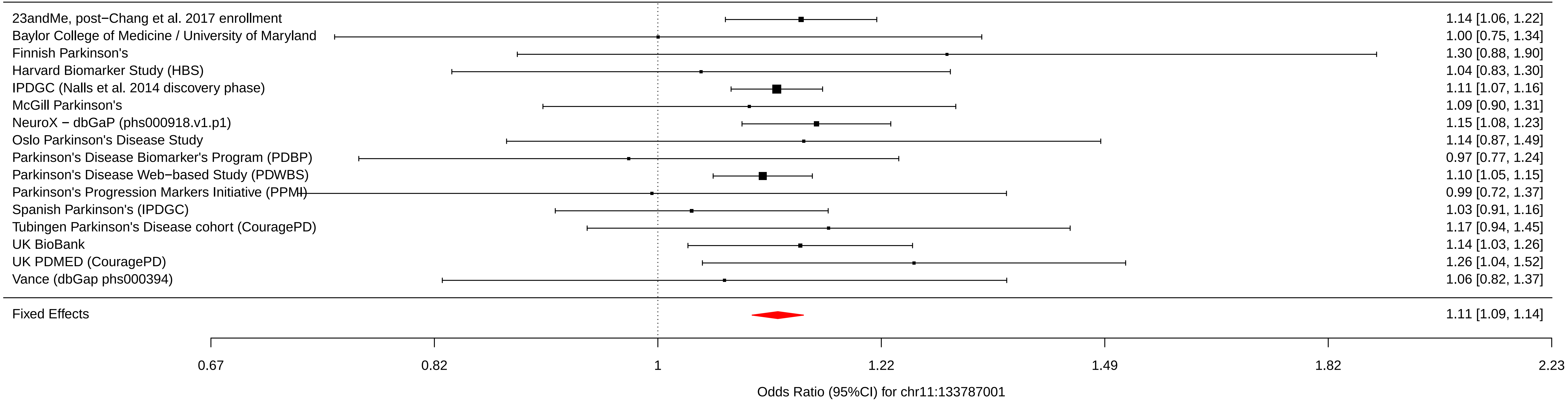

### chr12_40614434_rs76904798.pdf

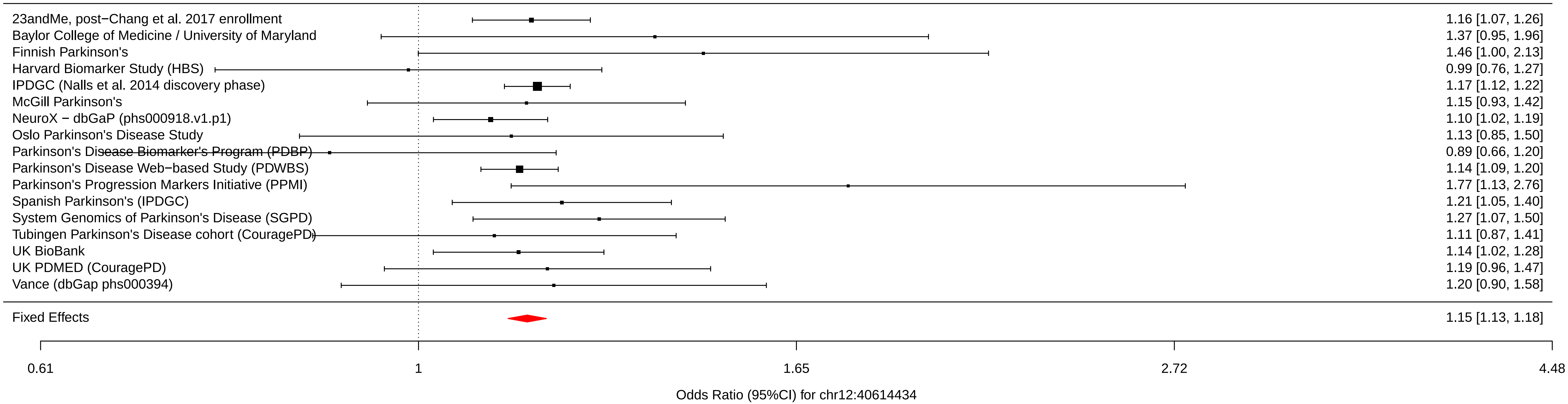

### chr12_40734202_rs34637584.pdf

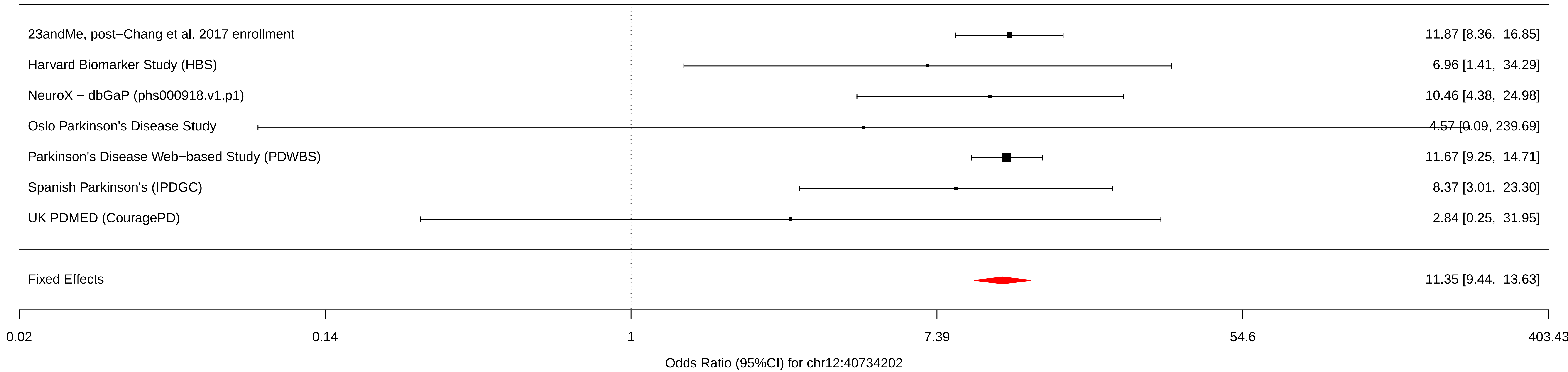

### chr12_46419086_rs7134559.pdf

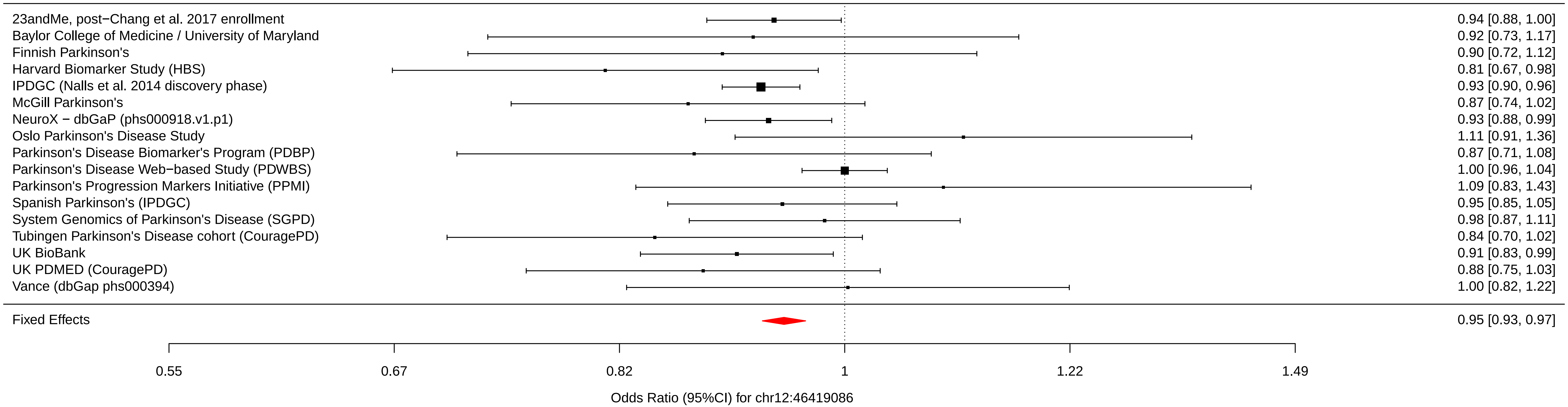

### chr12_123326598_rs10847864.pdf

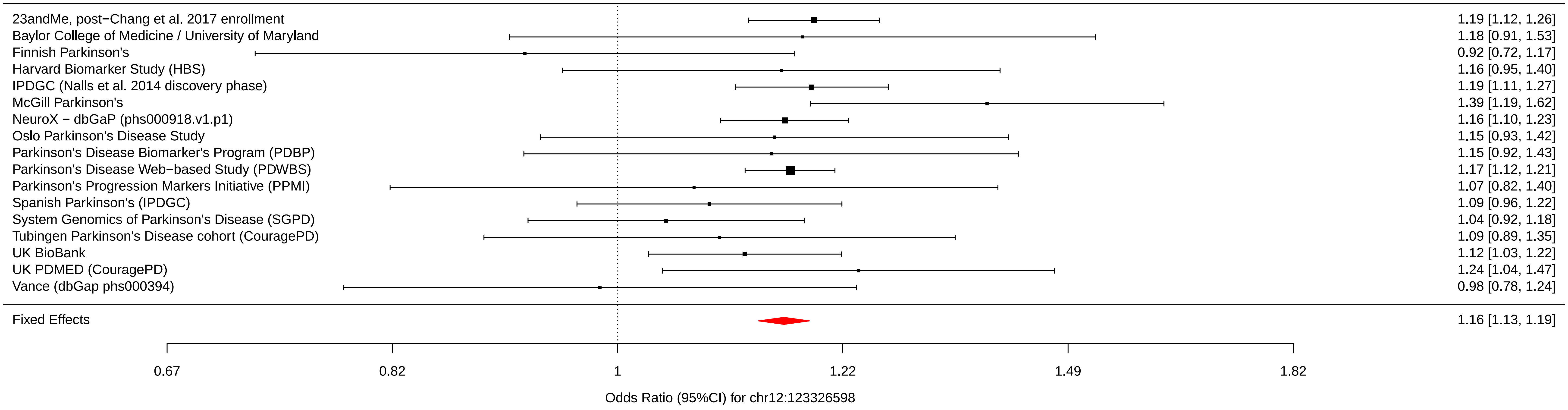

### chr12_133063768_rs11610045.pdf

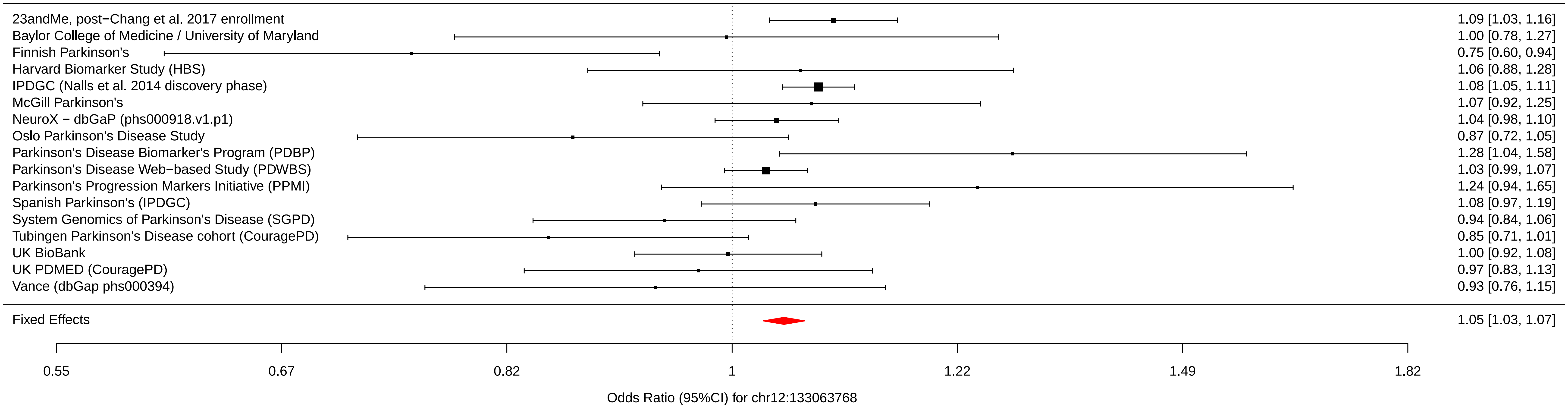

### chr13_49927732_rs9568188.pdf

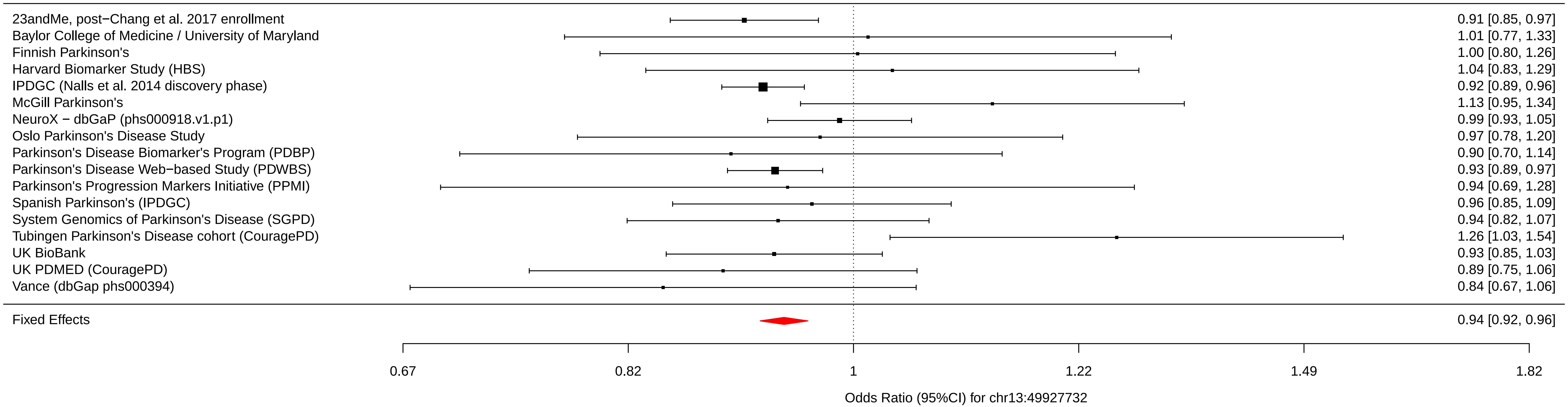

### chr13_97865021_rs4771268.pdf

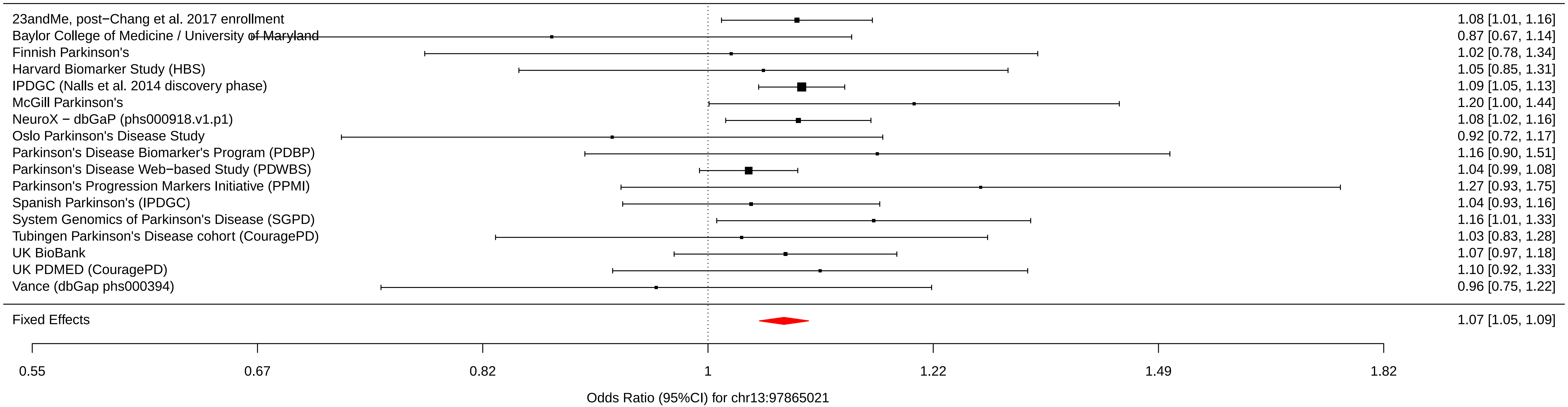

### chr14_37989270_rs12147950.pdf

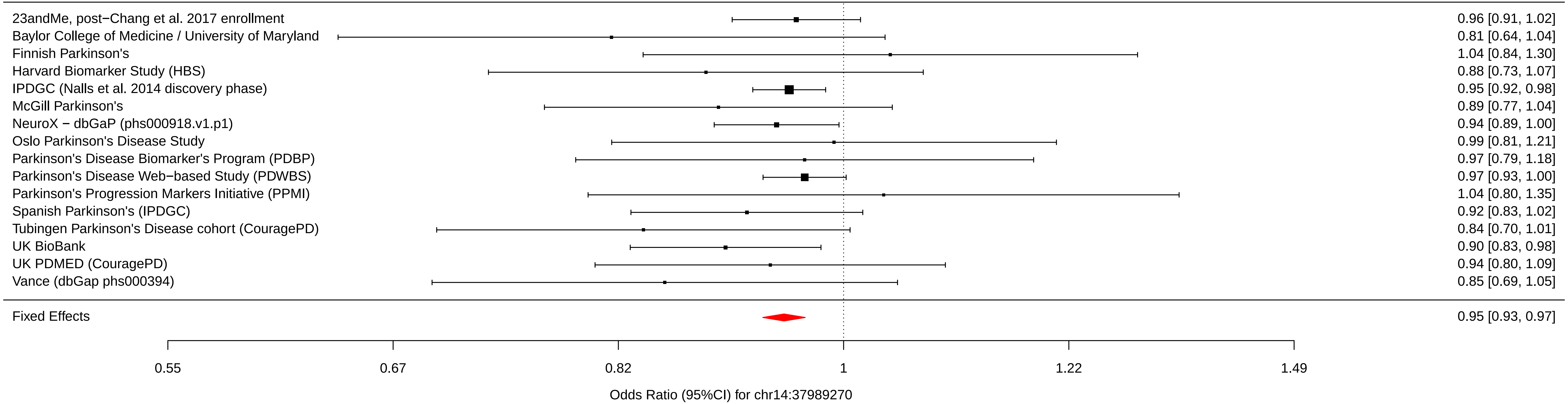

### chr14_55348869_rs11158026.pdf

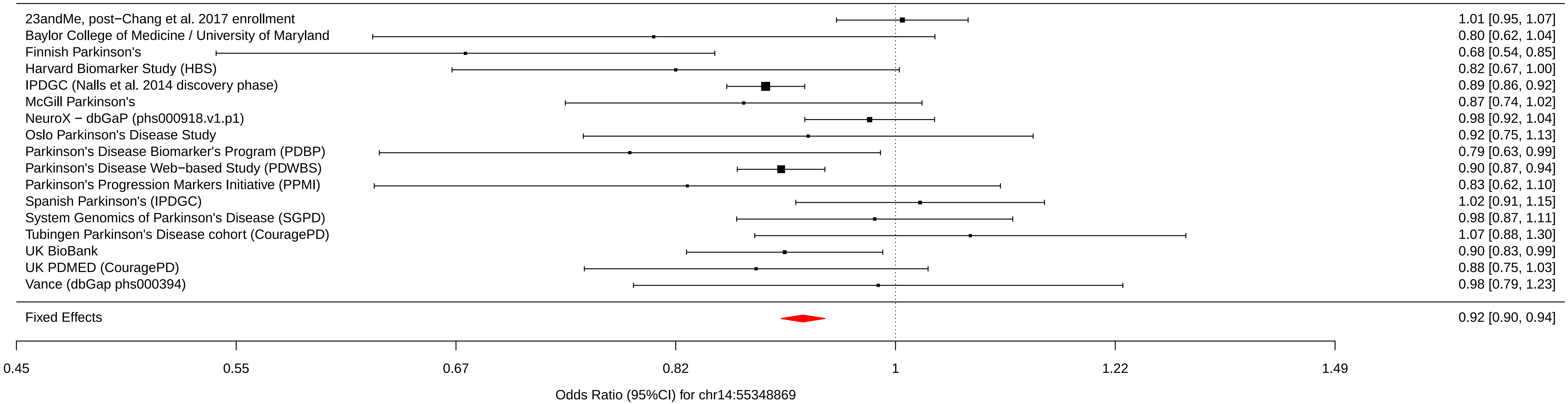

### chr14_75373034_rs3742785.pdf

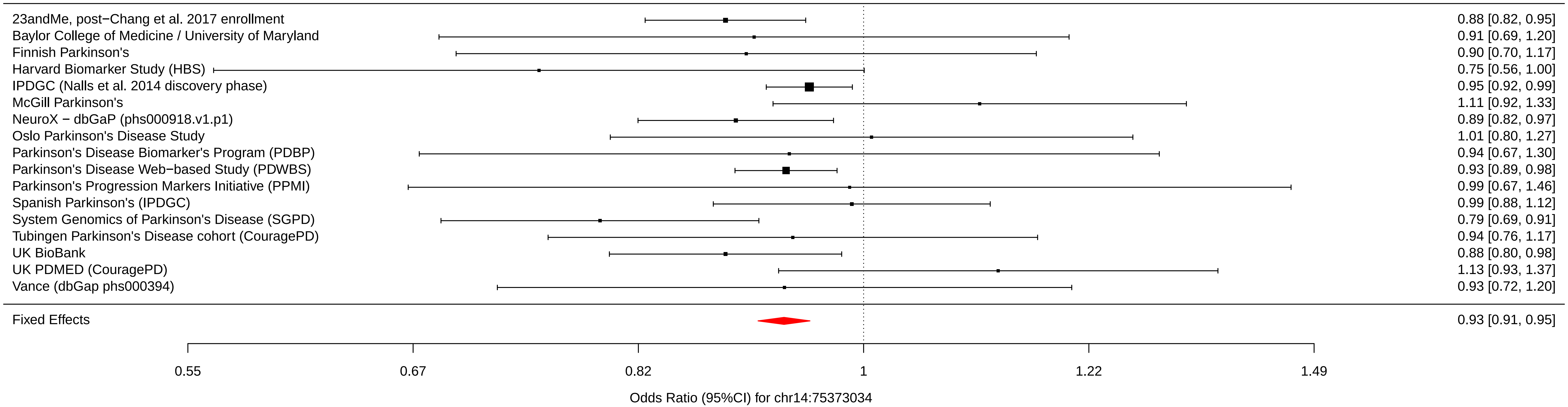

### chr14_88464264_rs979812.pdf

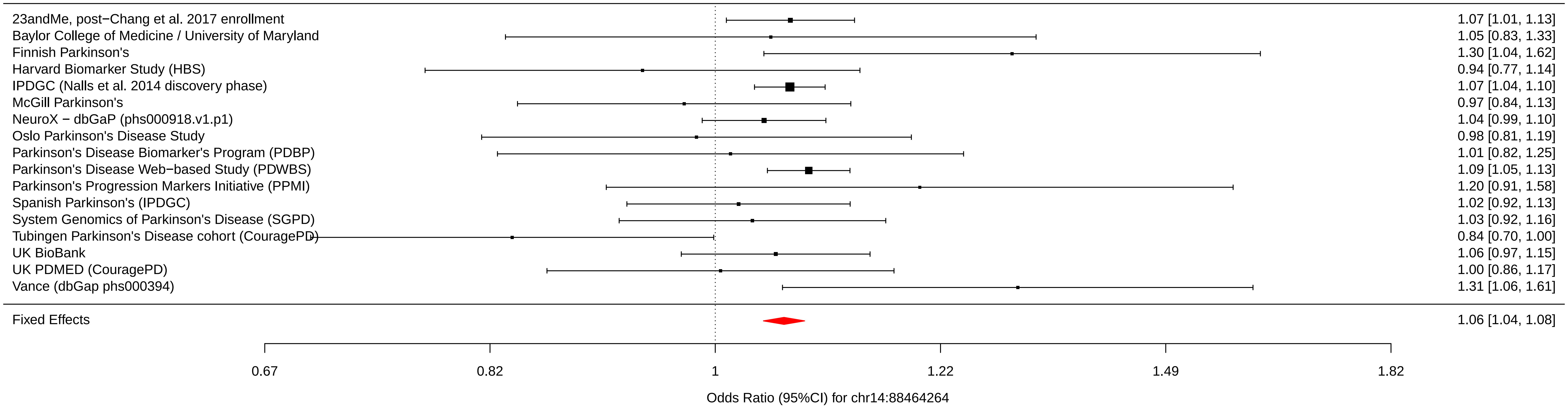

### chr15_61997385_rs2251086.pdf

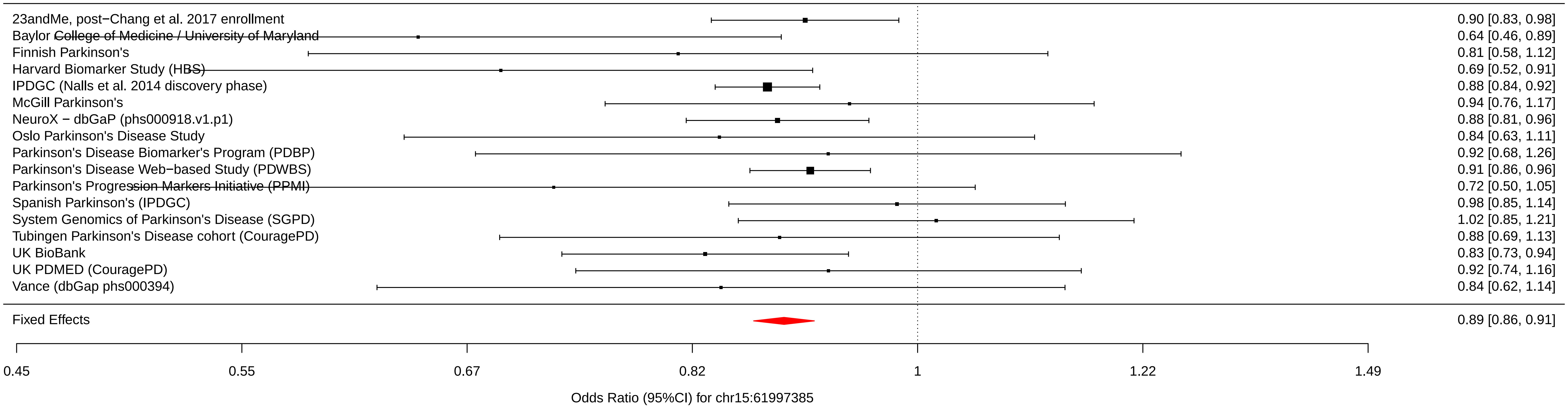

### chr16_19277493_rs6497339.pdf

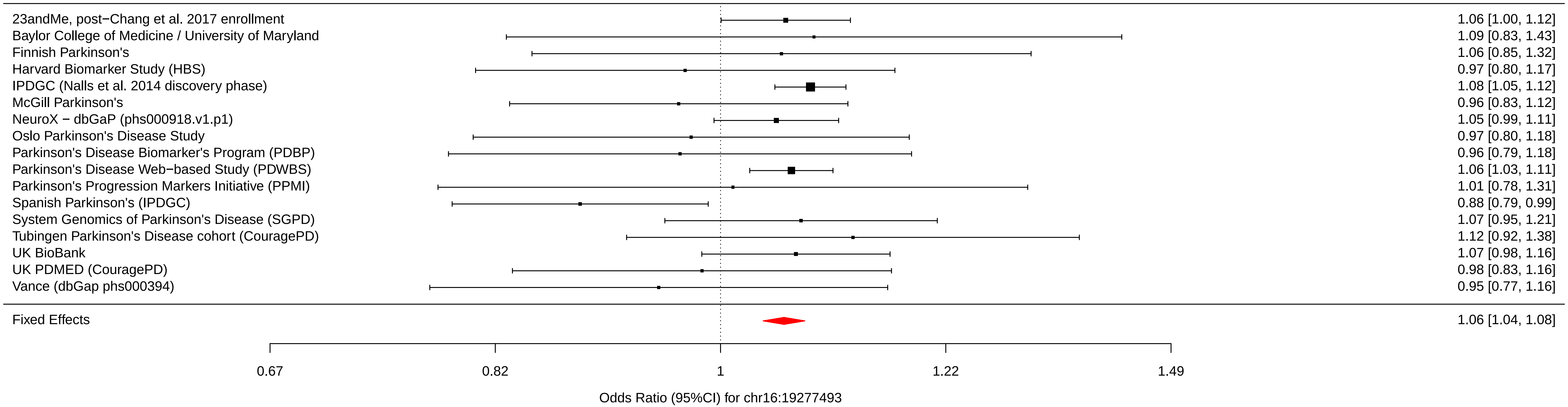

### chr16_28944396_rs2904880.pdf

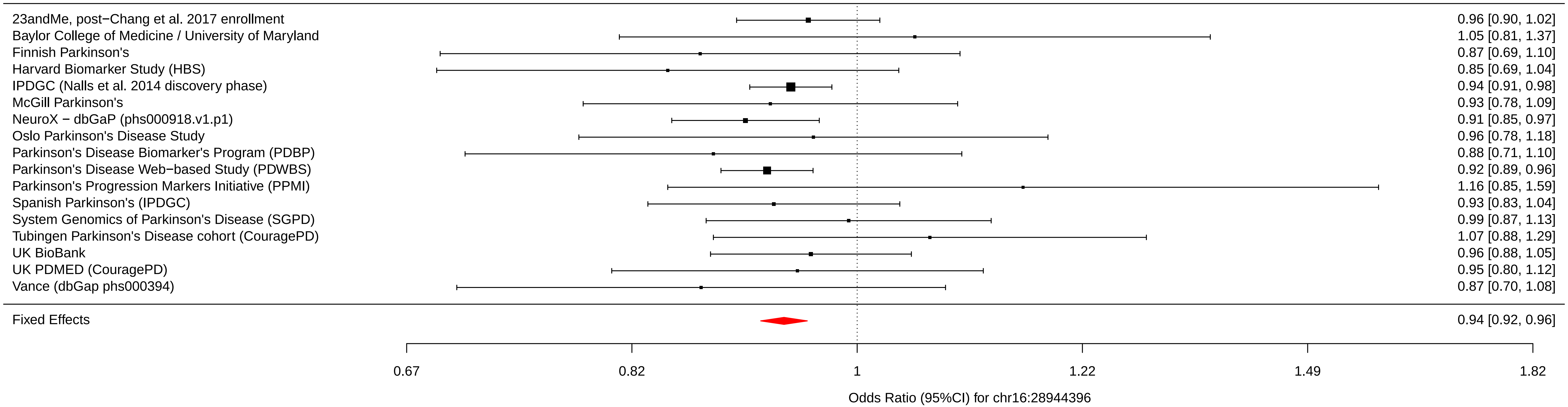

### chr16_30977799_rs11150601.pdf

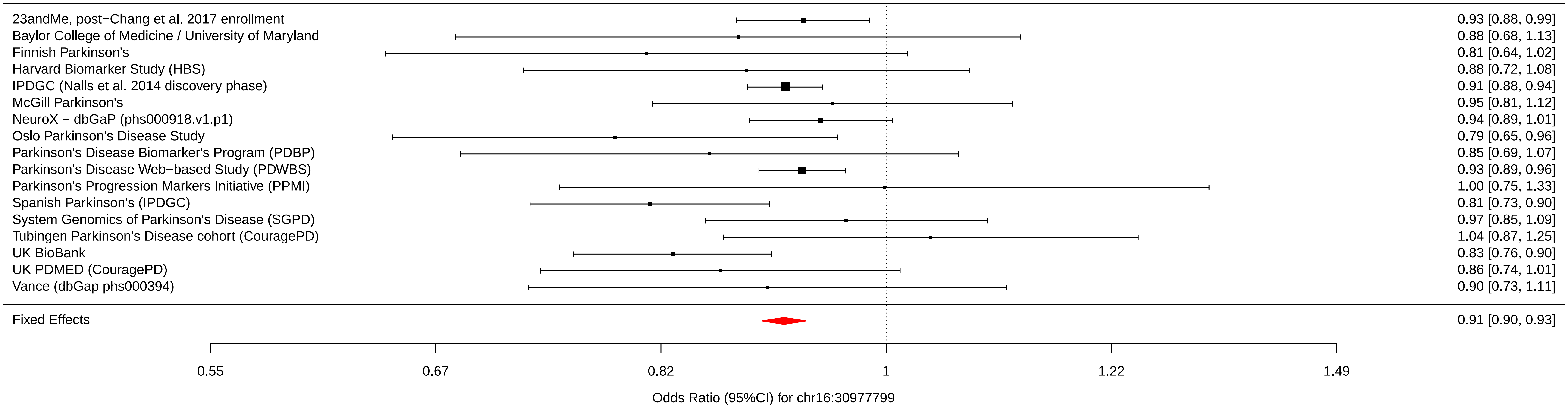

### chr16_50736656_rs6500328.pdf

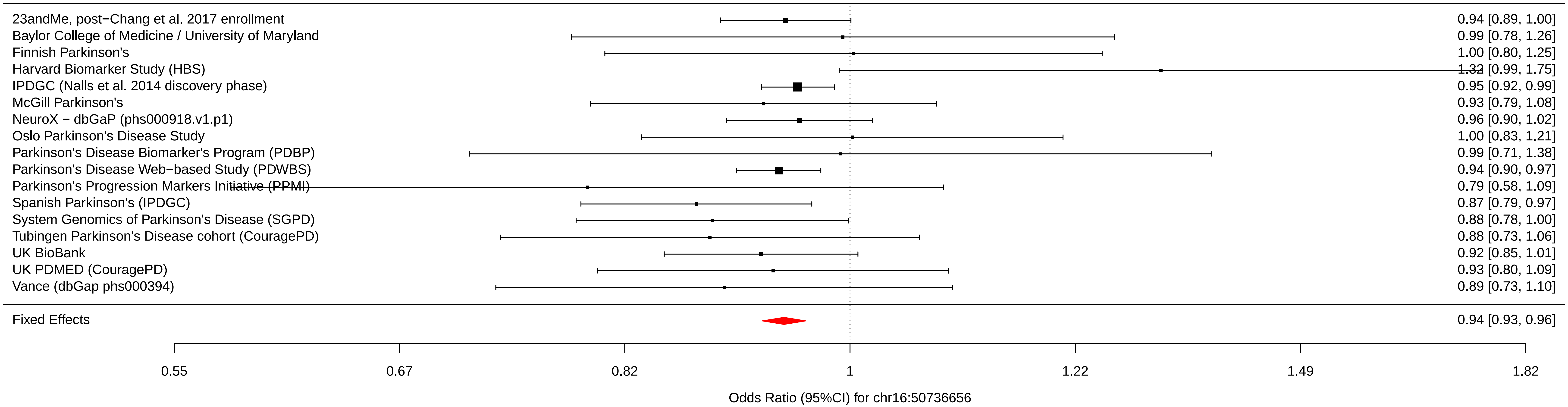

### chr16_52636242_rs3104783.pdf

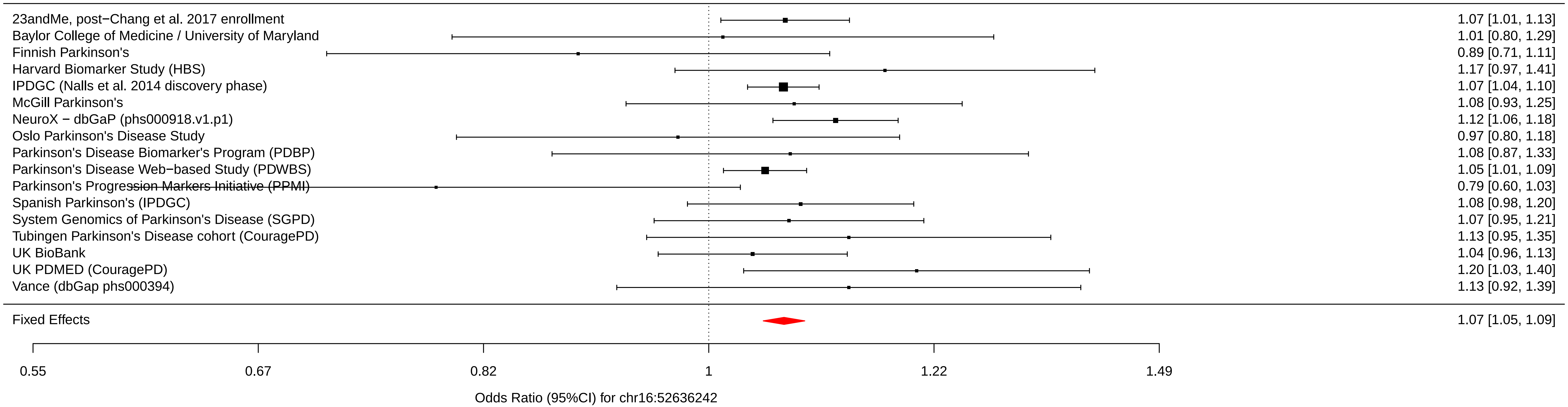

### chr16_52969426_rs10221156.pdf

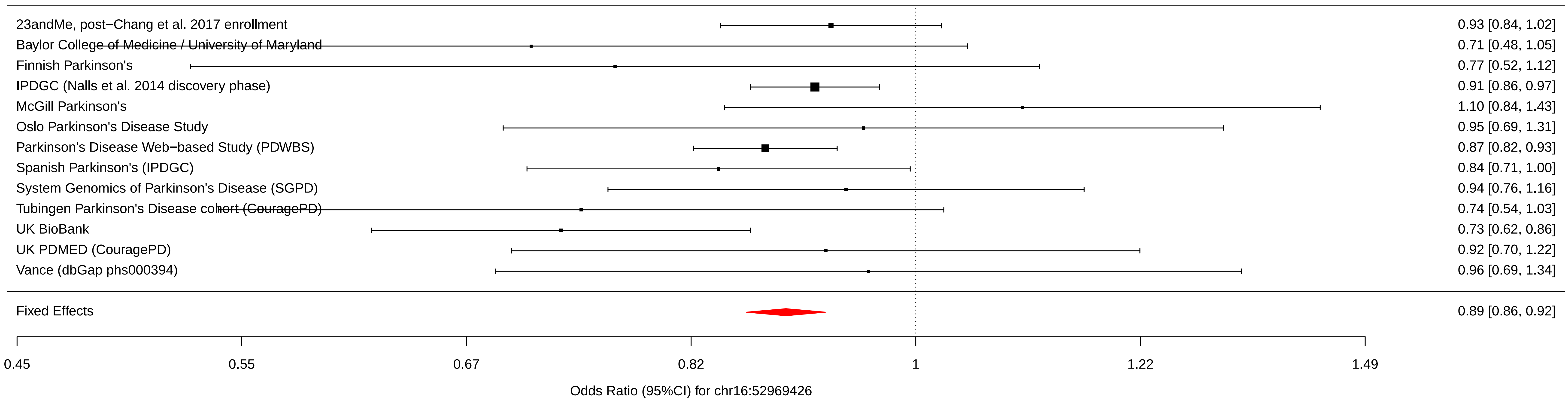

### chr17_7355621_rs12600861.pdf

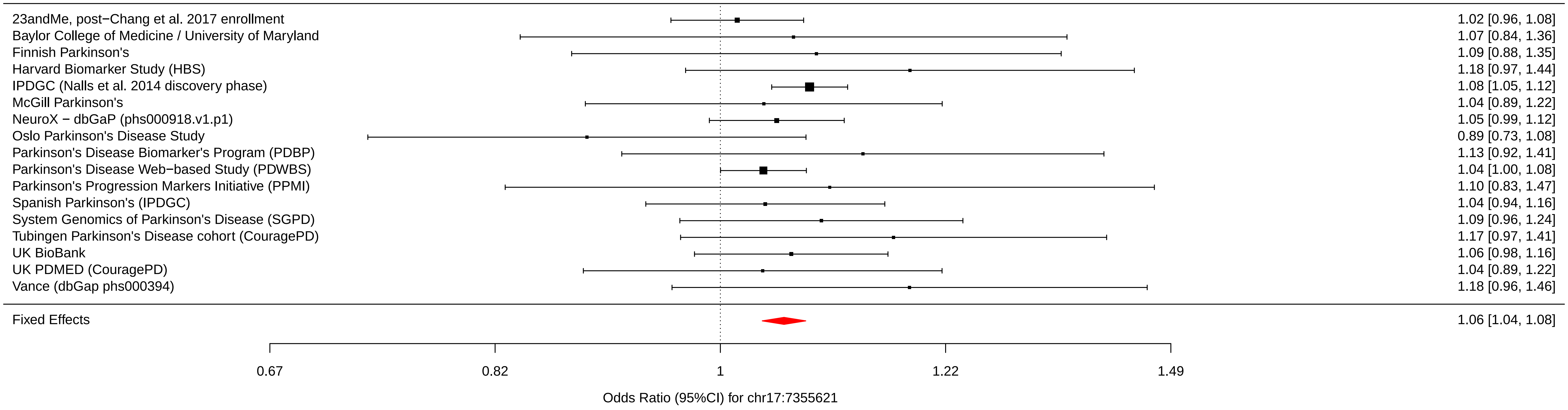

### chr17_40741013_rs12951632.pdf

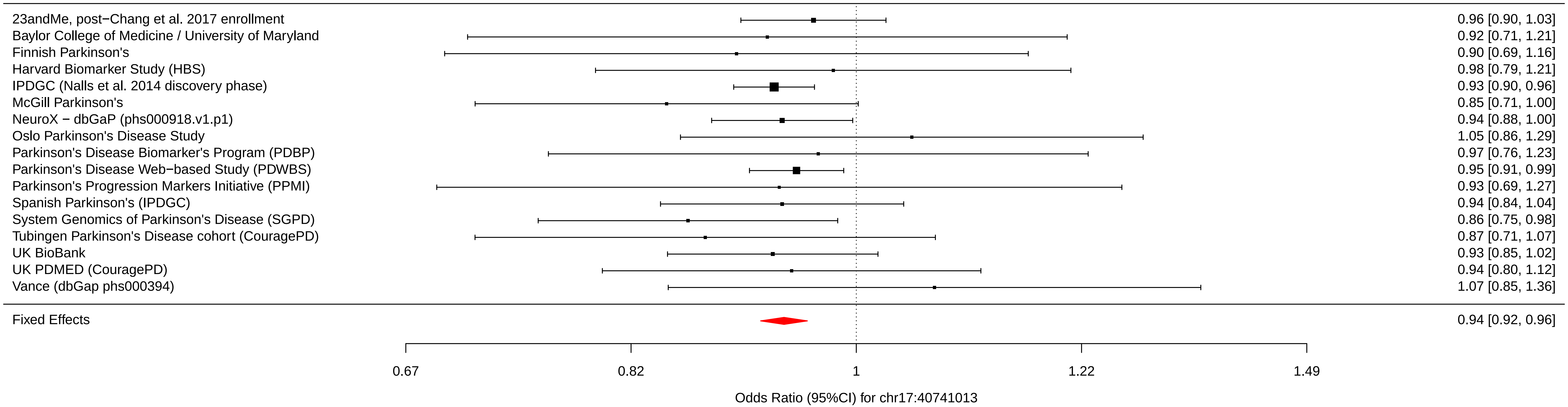

### chr17_42294337_rs2269906.pdf

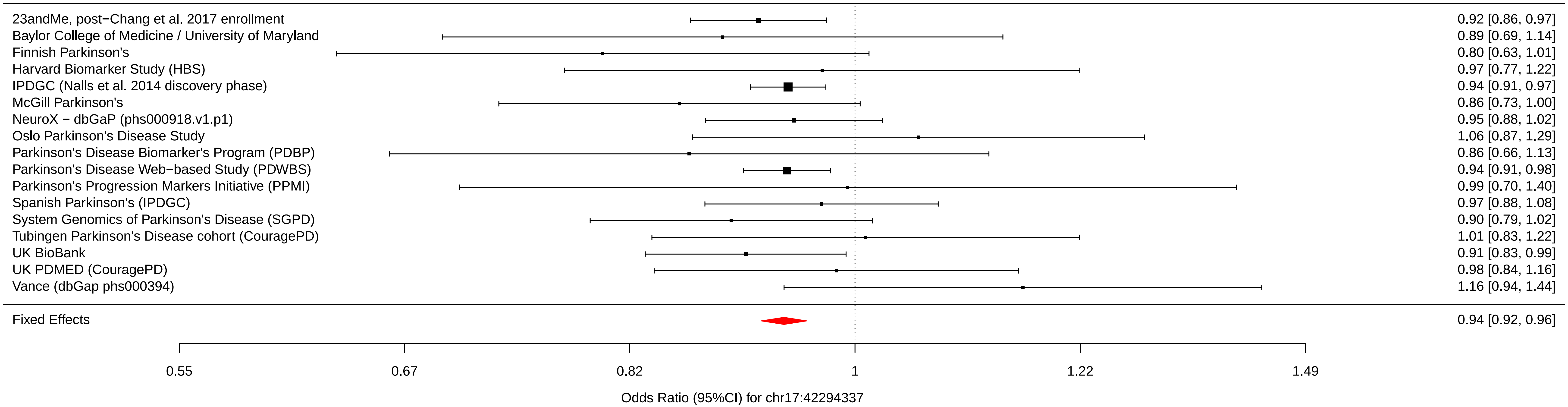

### chr17_42434630_rs850738.pdf

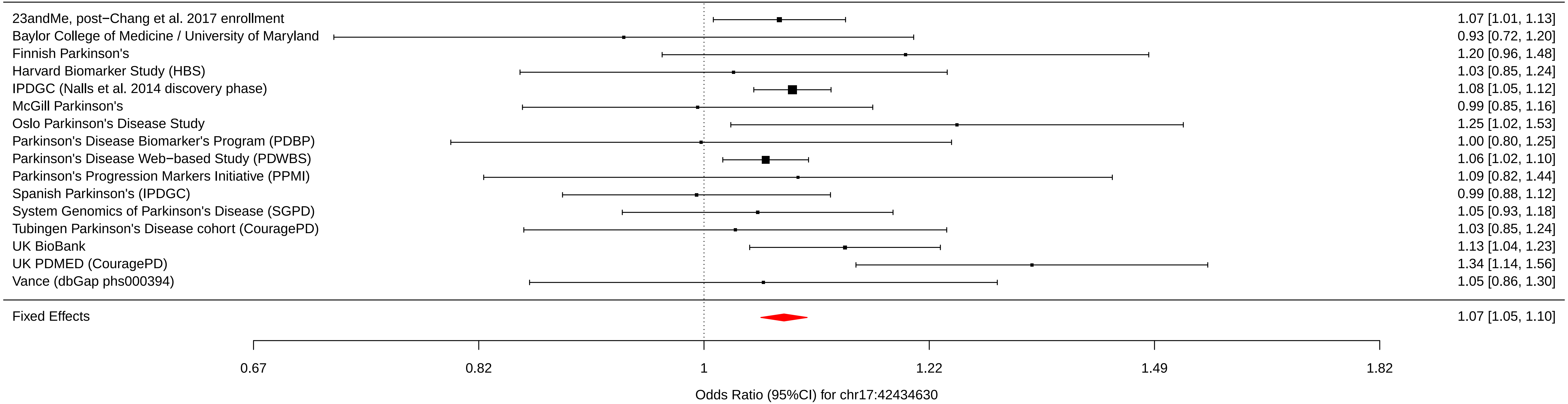

### chr17_43744203_rs62053943.pdf

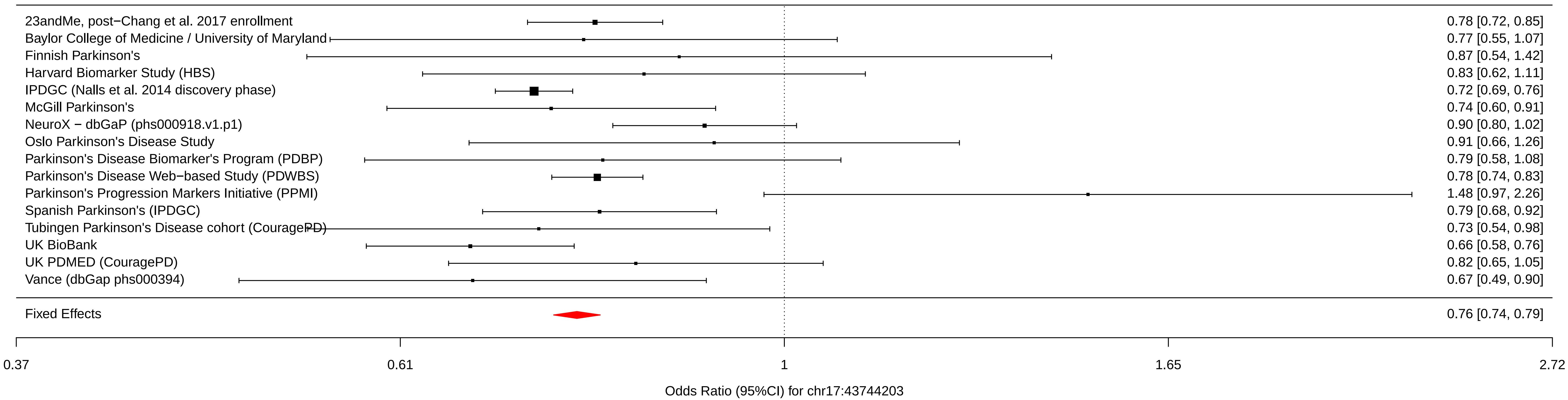

### chr17_43798308_rs117615688.pdf

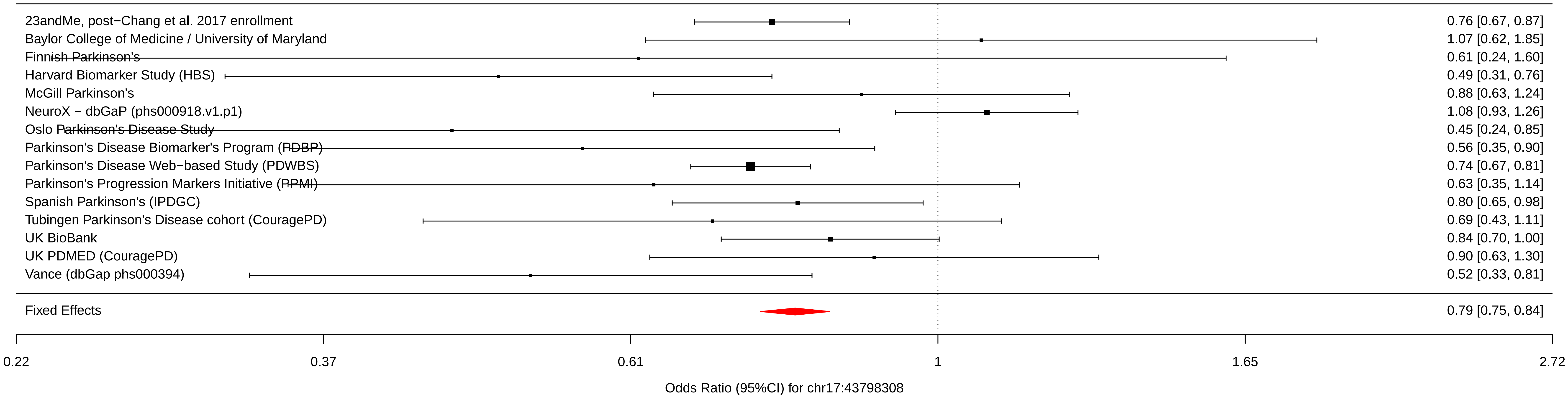

### chr17_44866805_rs11658976.pdf

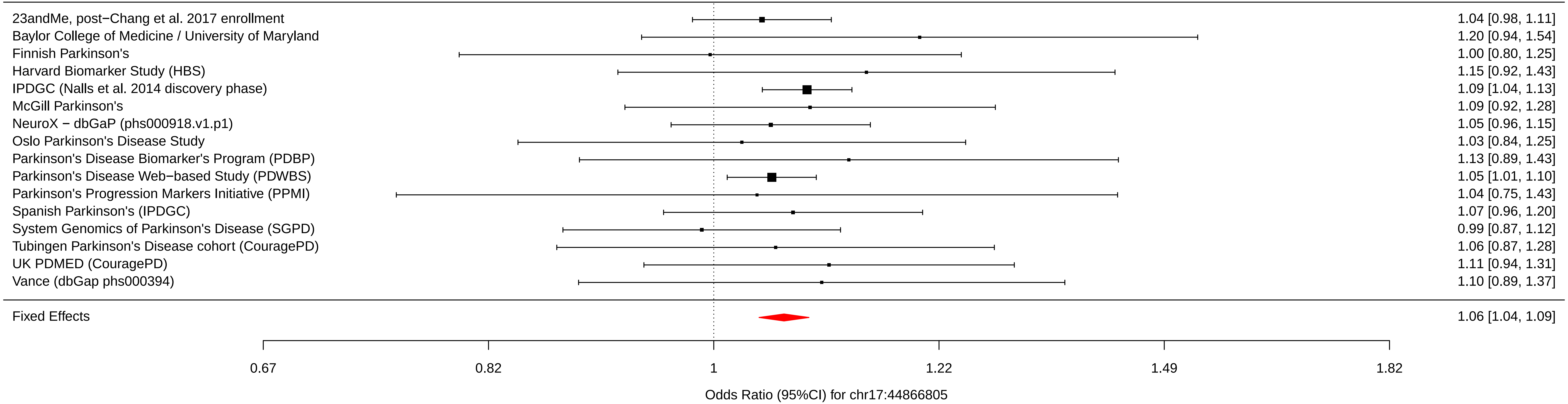

### chr17_59917366_rs61169879.pdf

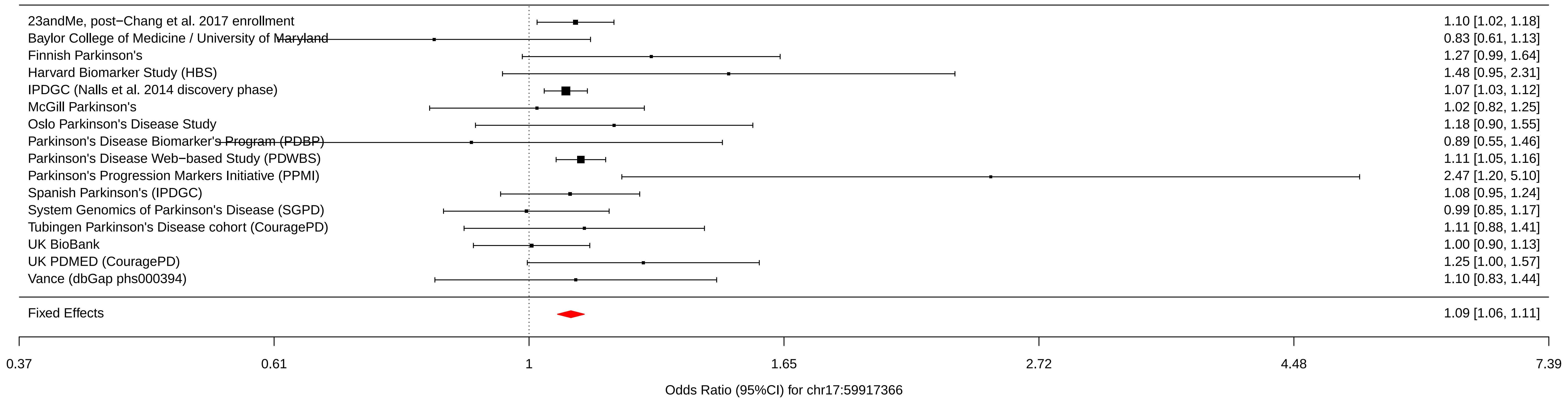

### chr17_76425480_rs666463.pdf

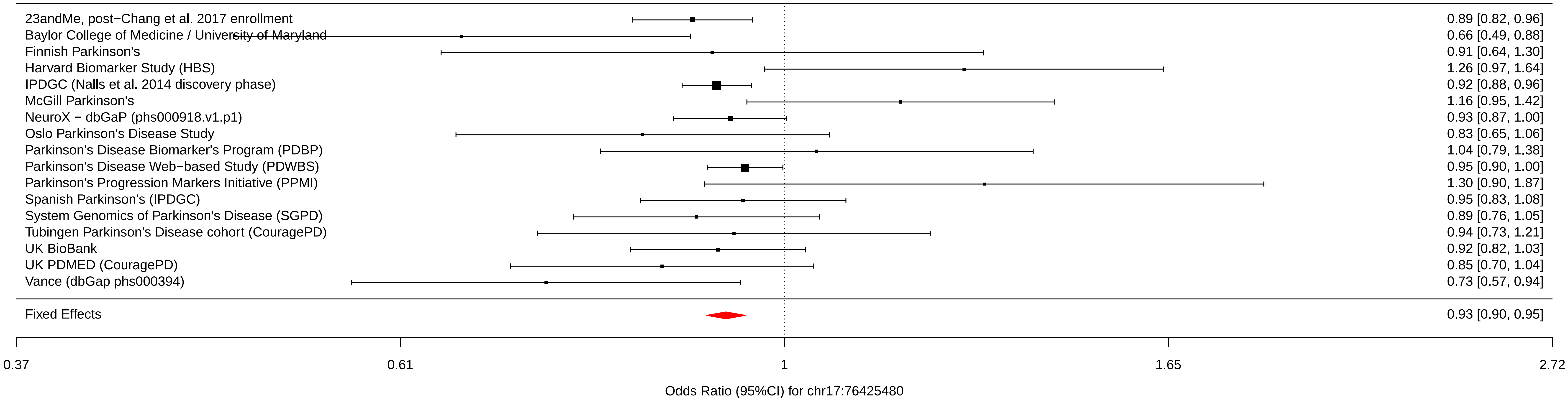

### chr18_31304318_rs1941685.pdf

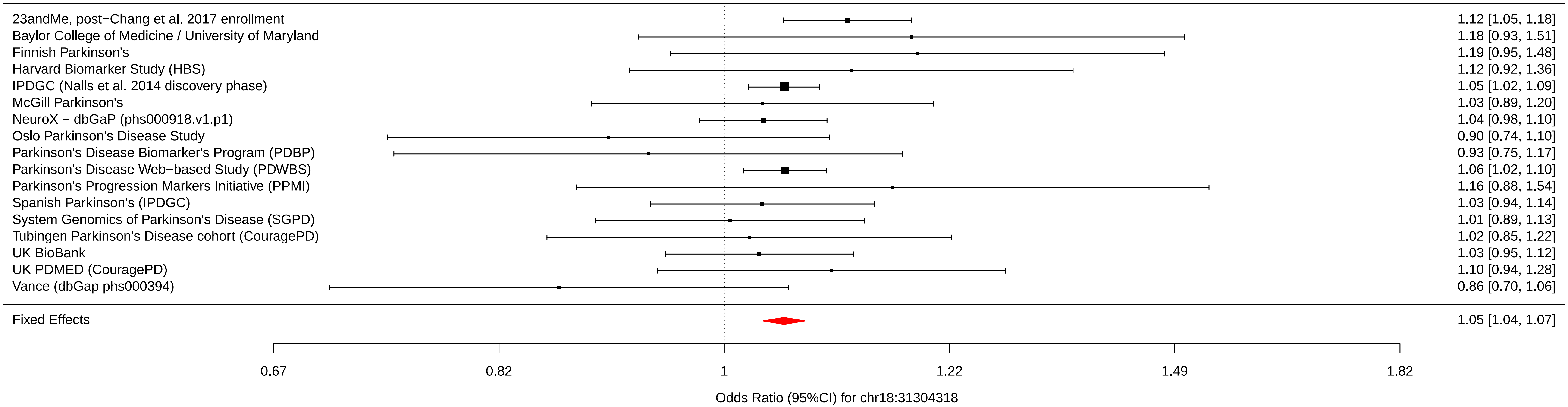

### chr18_48683589_rs8087969.pdf

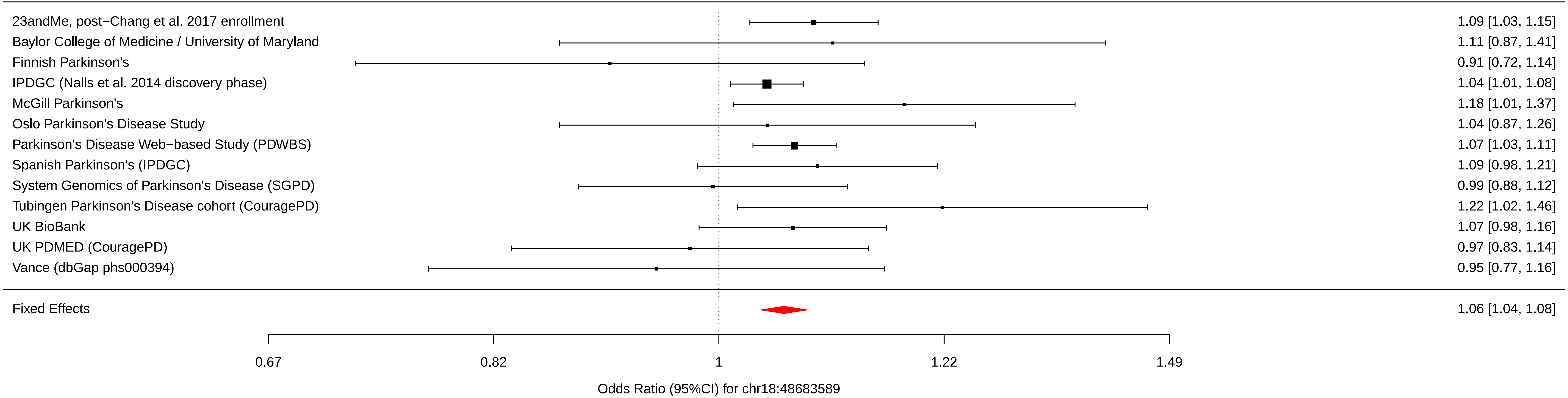

### chr19_2341047_rs55818311.pdf

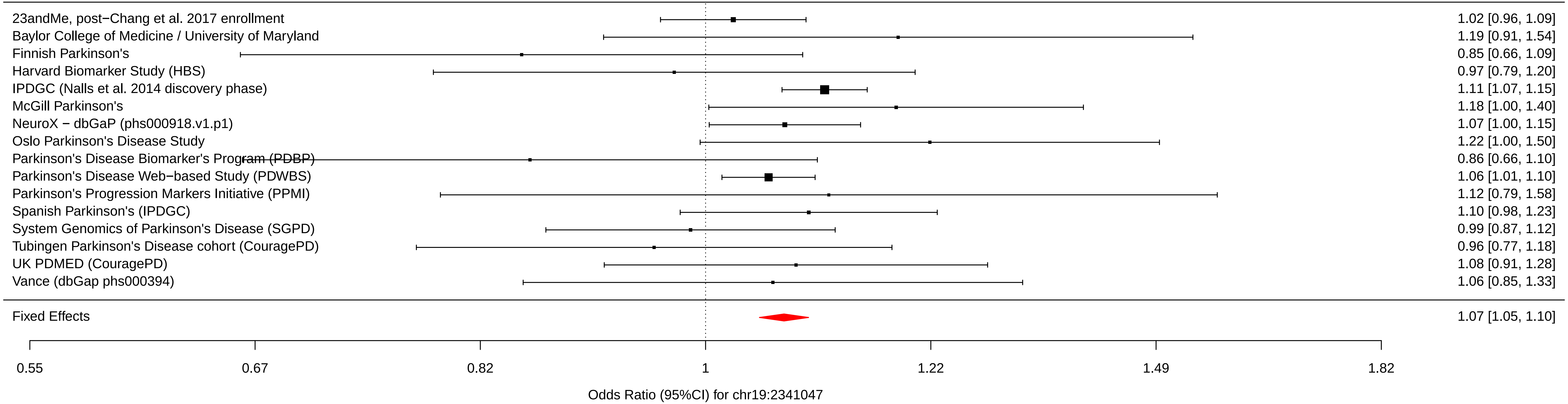

### chr20_6006041_rs77351827.pdf

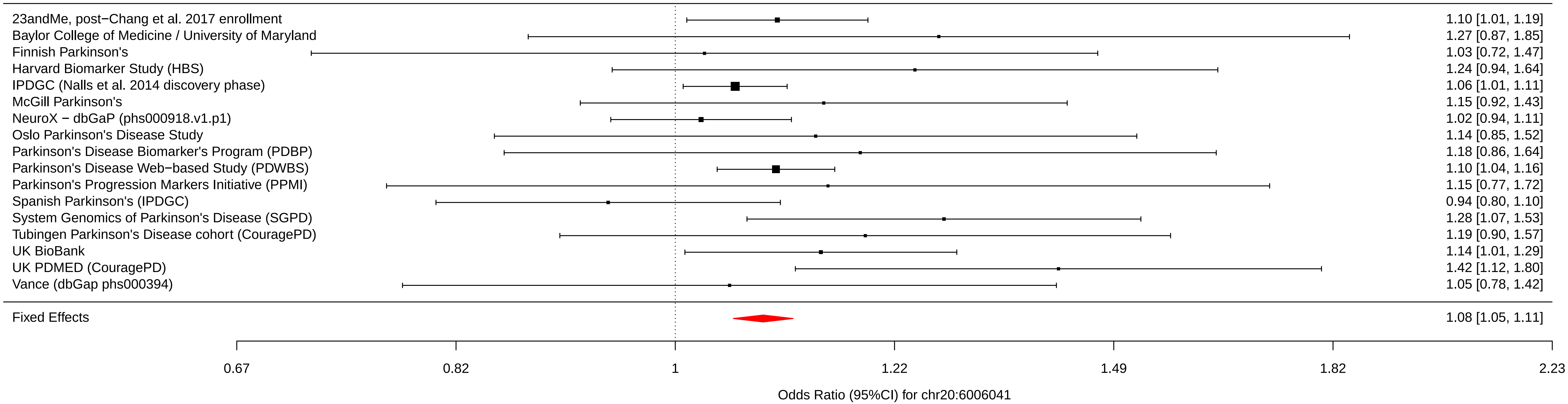

### chr21_38852361_rs2248244.pdf

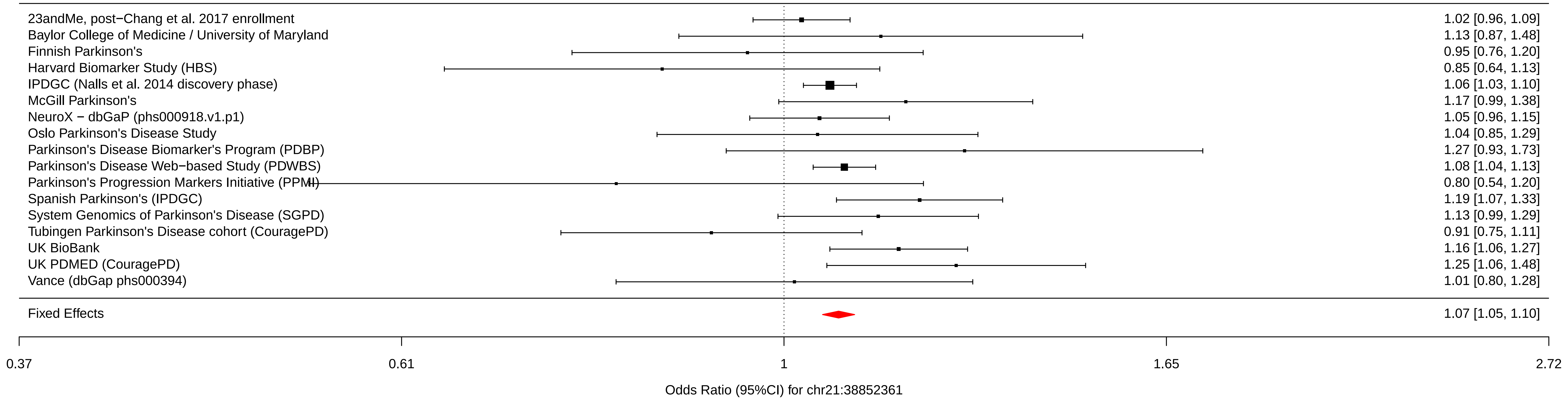

### Finnish_PD_3phase.jpg

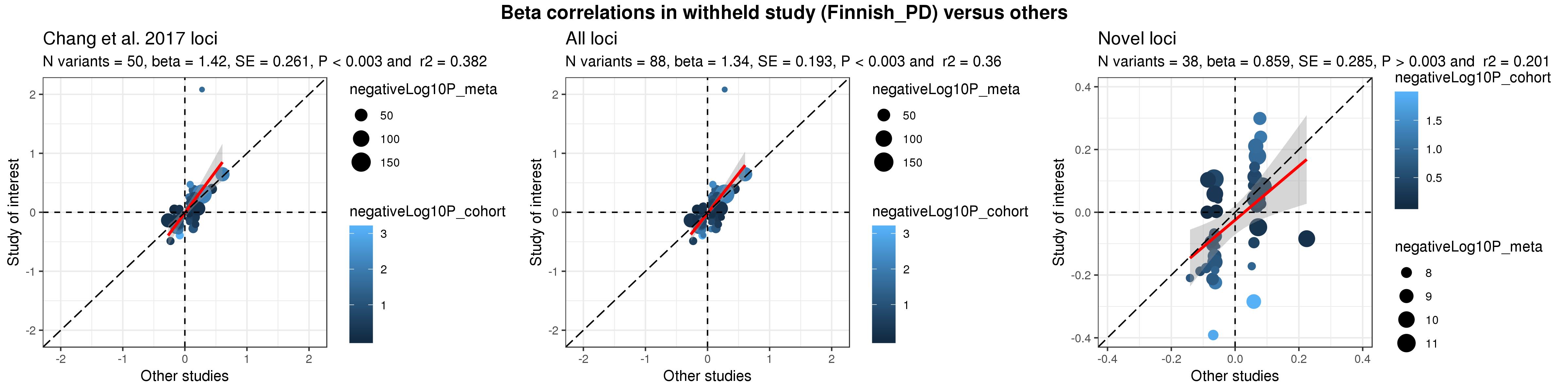

### Oslo_PD_3phase.jpg

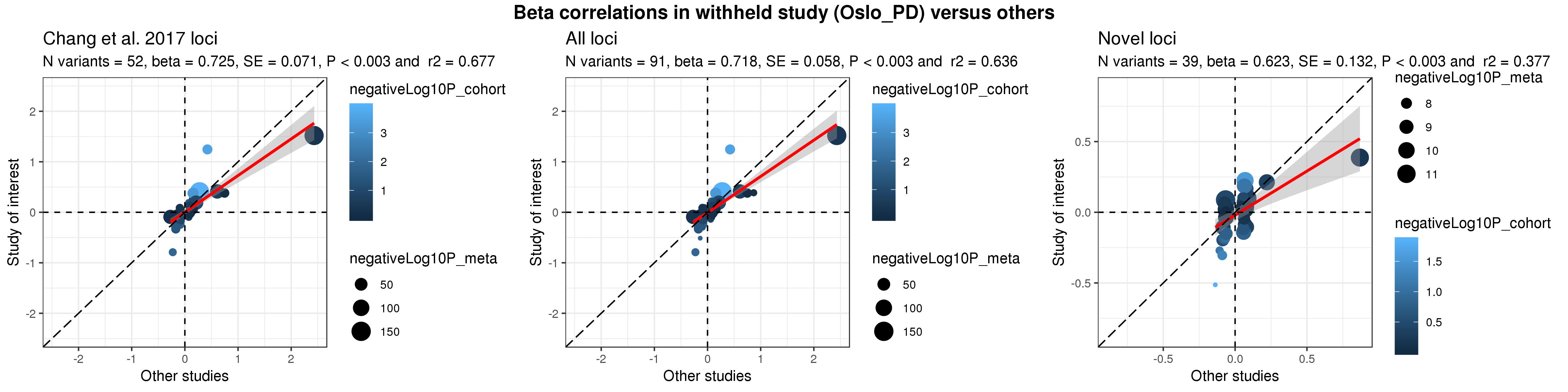

### PDBP_3phase.jpg

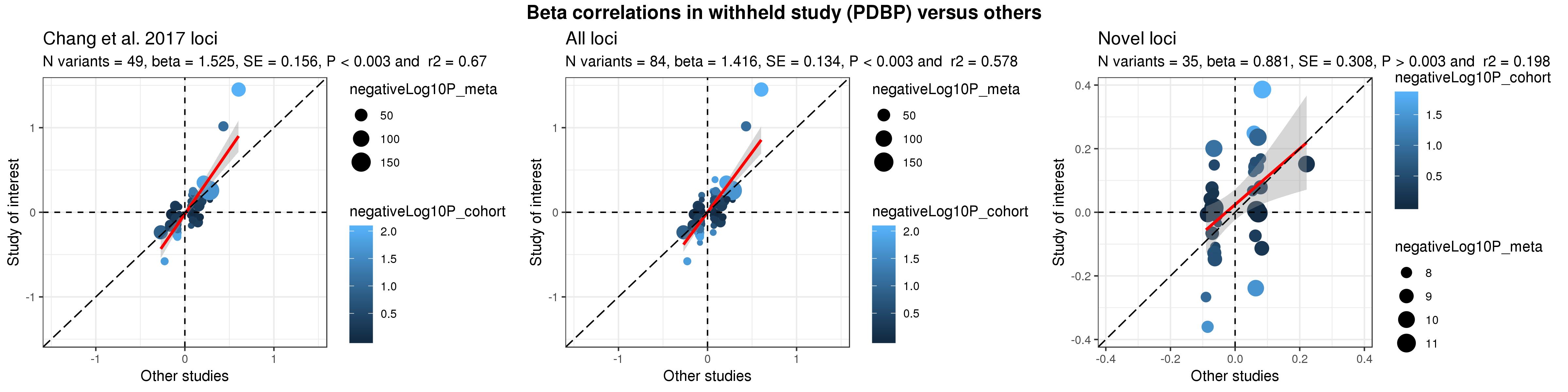

### PDMED_3phase.jpg

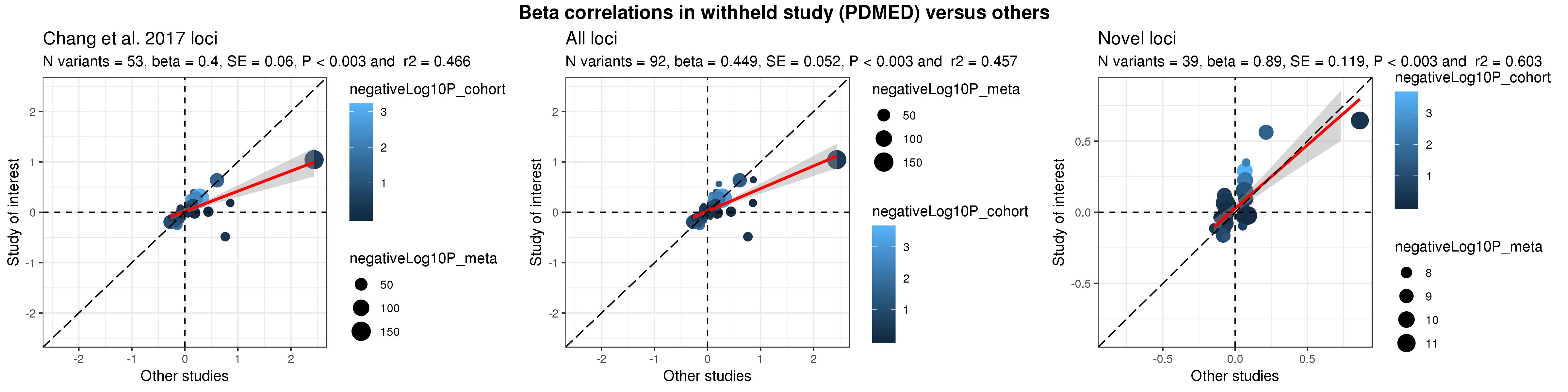

### PPMI_3phase.jpg

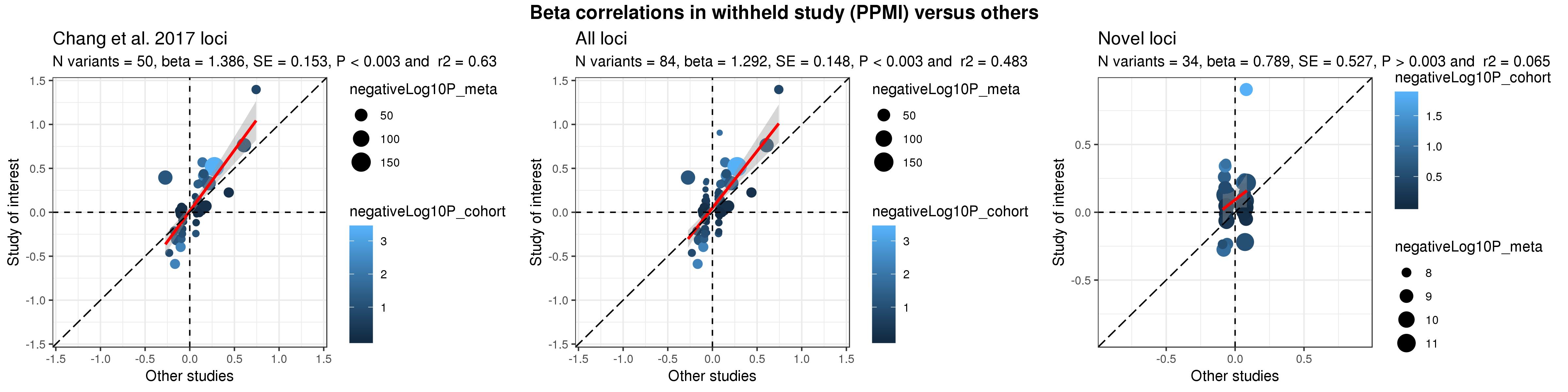

### SGPD_3phase.jpg

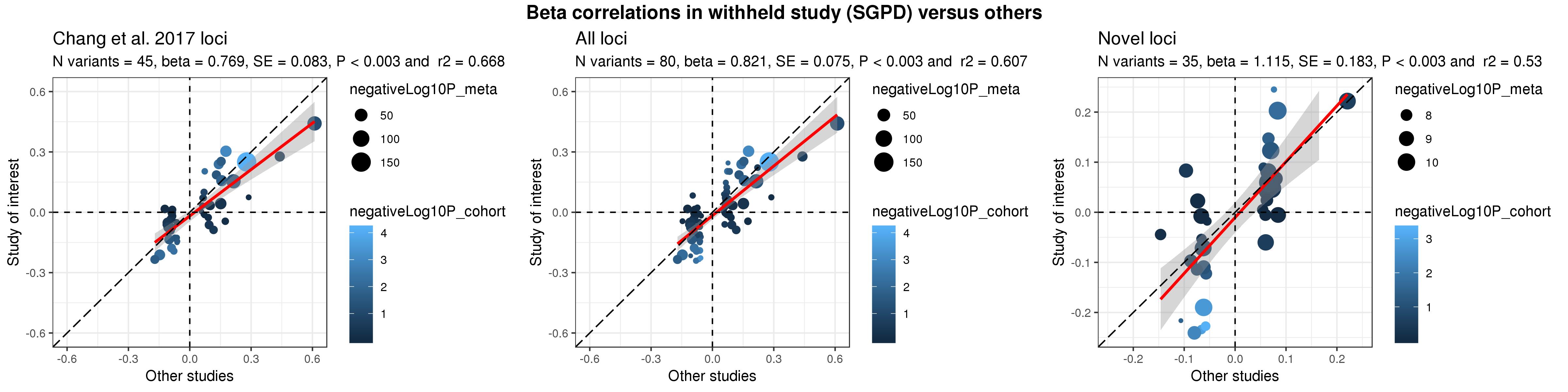

### UKB_3phase.jpg

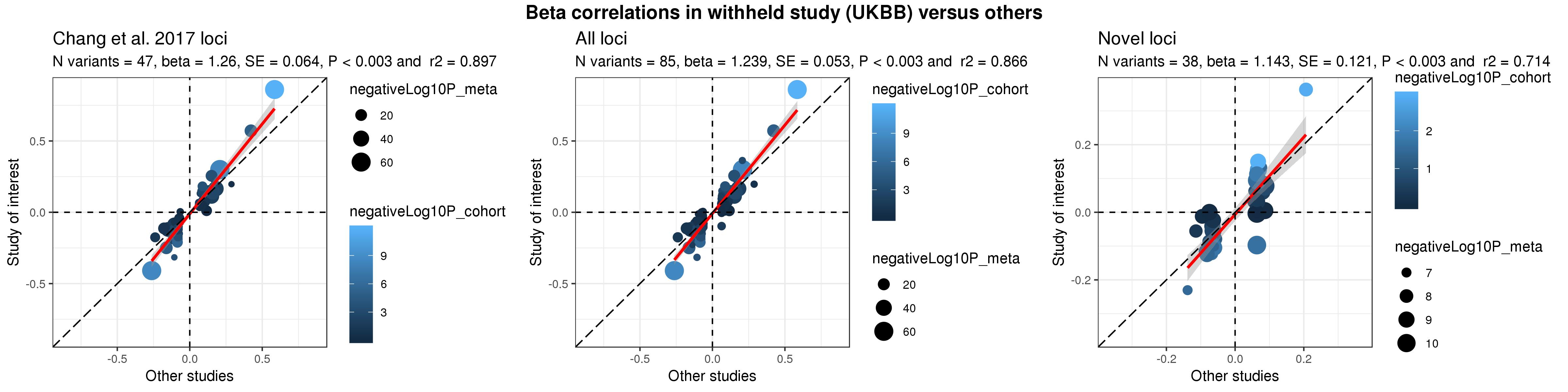
