## Supplementary figures and images for "Expanding Parkinson’s disease genetics: novel risk loci, genomic context, causal insights and heritable risk"

### chr1_155135036_rs35749011.pdf

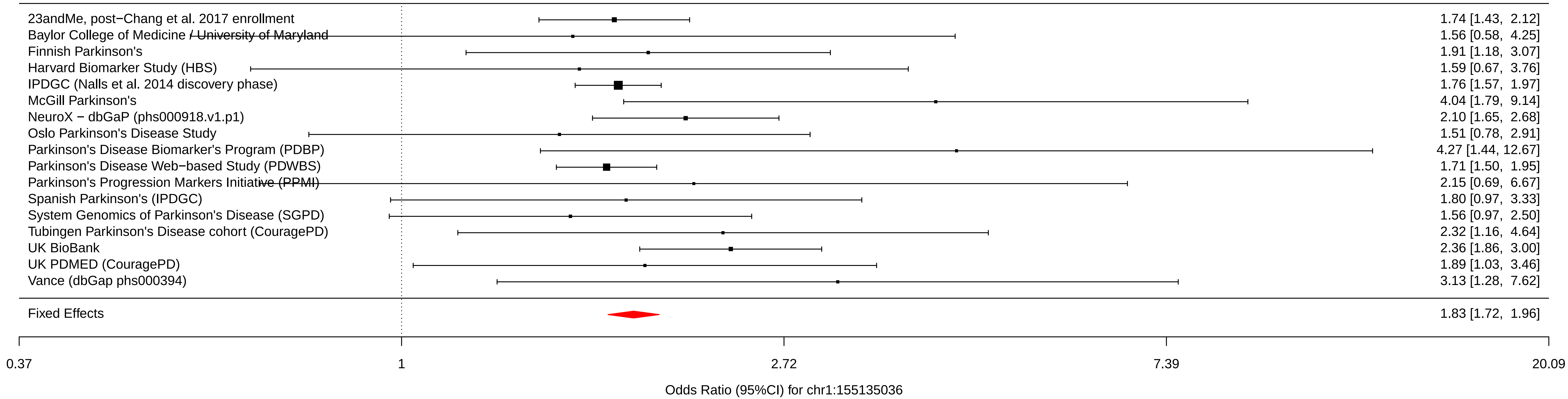

### chr2_169110394_rs1474055.pdf

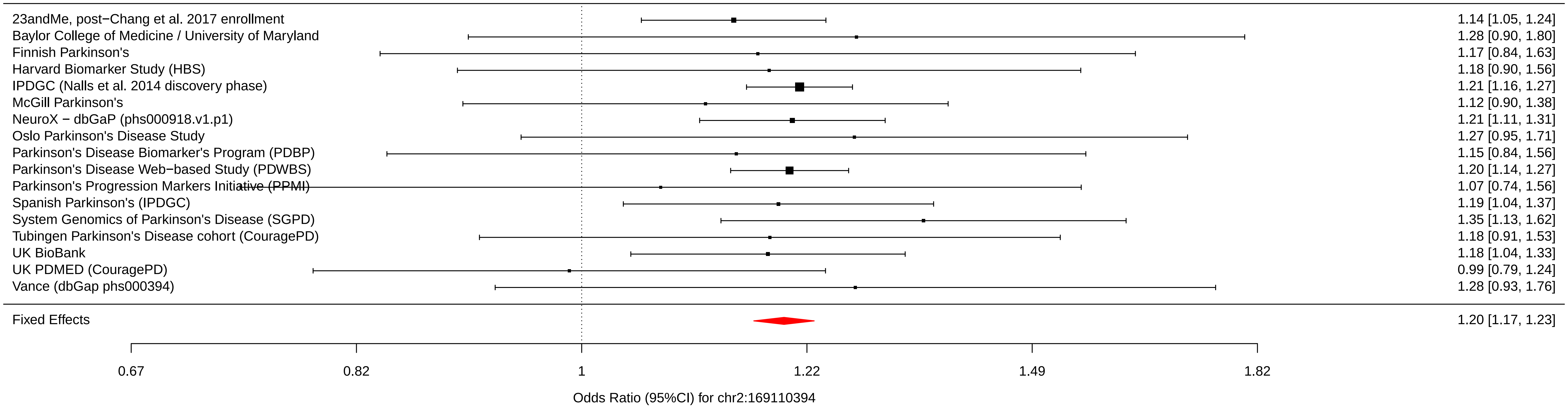

### chr3_28705690_rs6808178.pdf

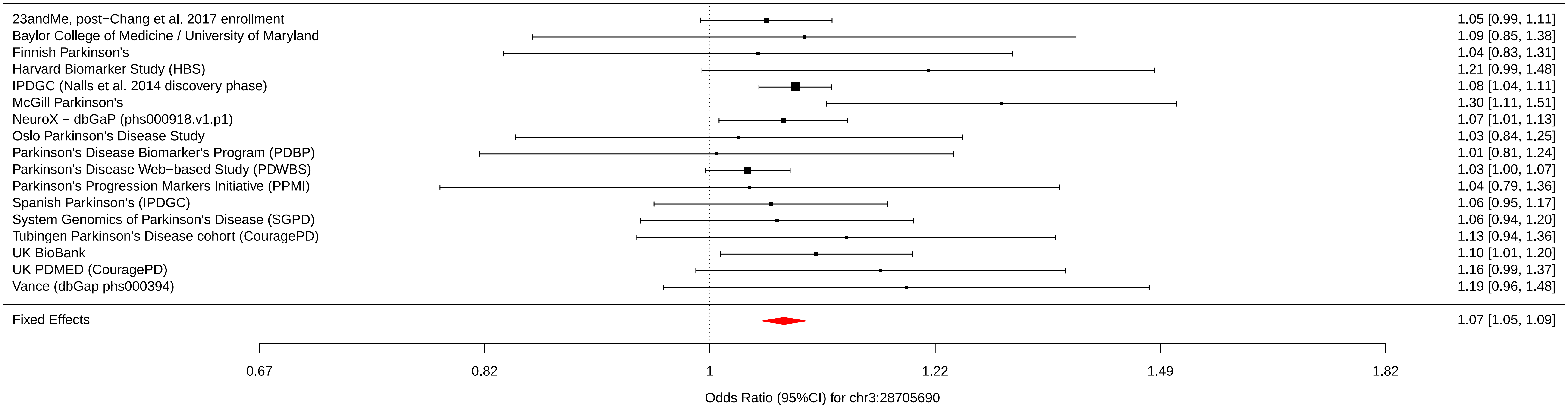

### chr4_114369065_rs13117519.pdf

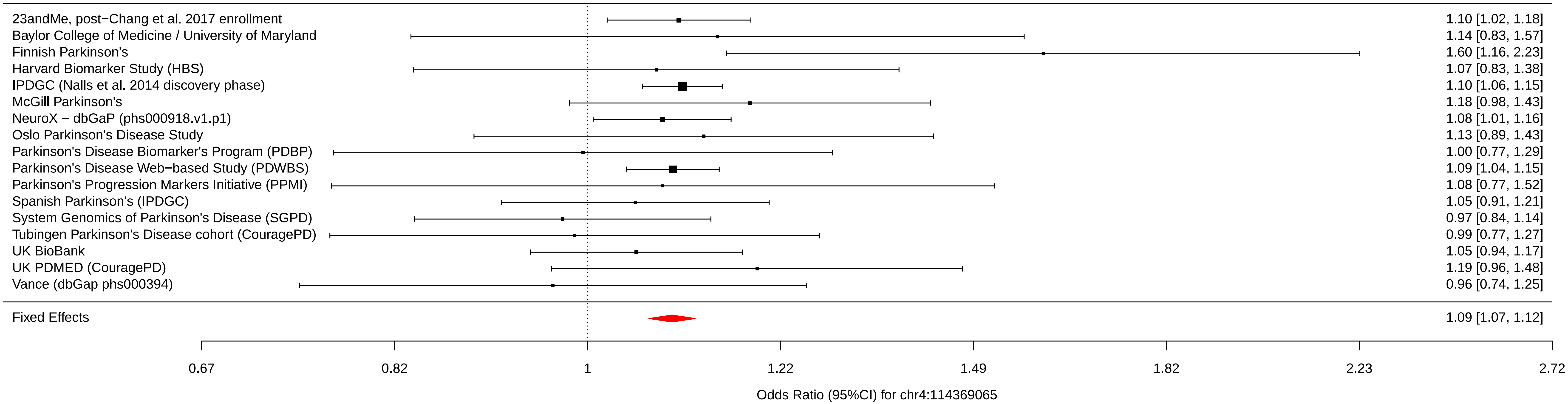

### chr4_170583157_rs62333164.pdf

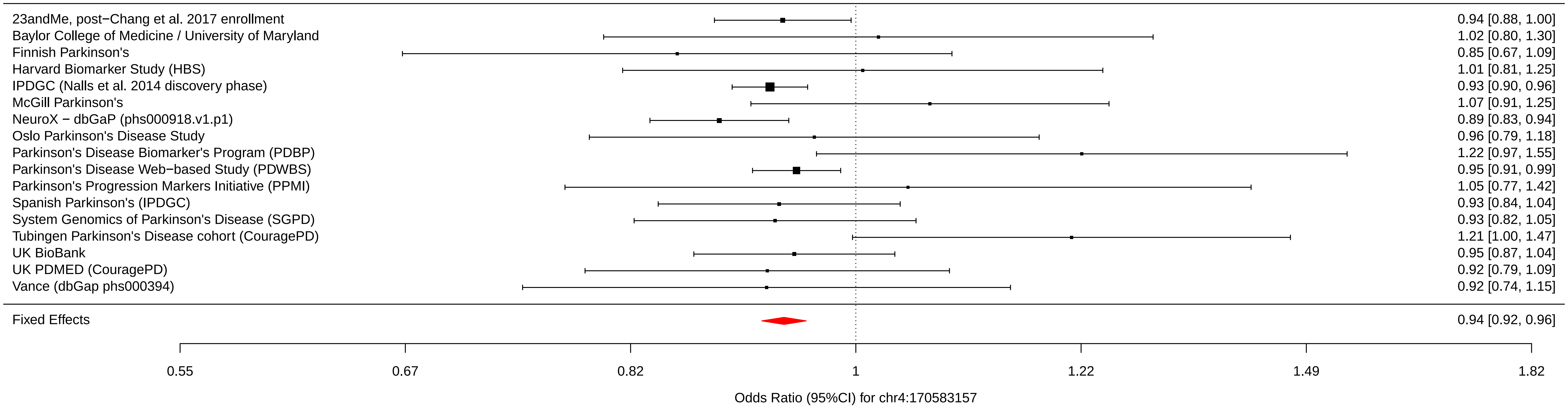

### chr6_30108683_rs9261484.pdf

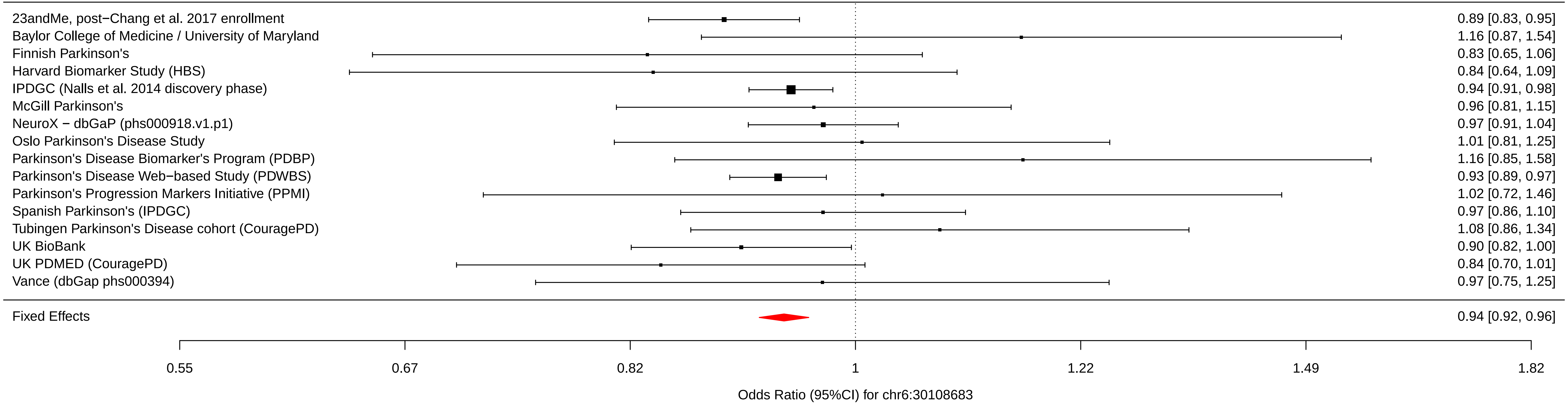

### chr6_32578772_rs112485576.pdf

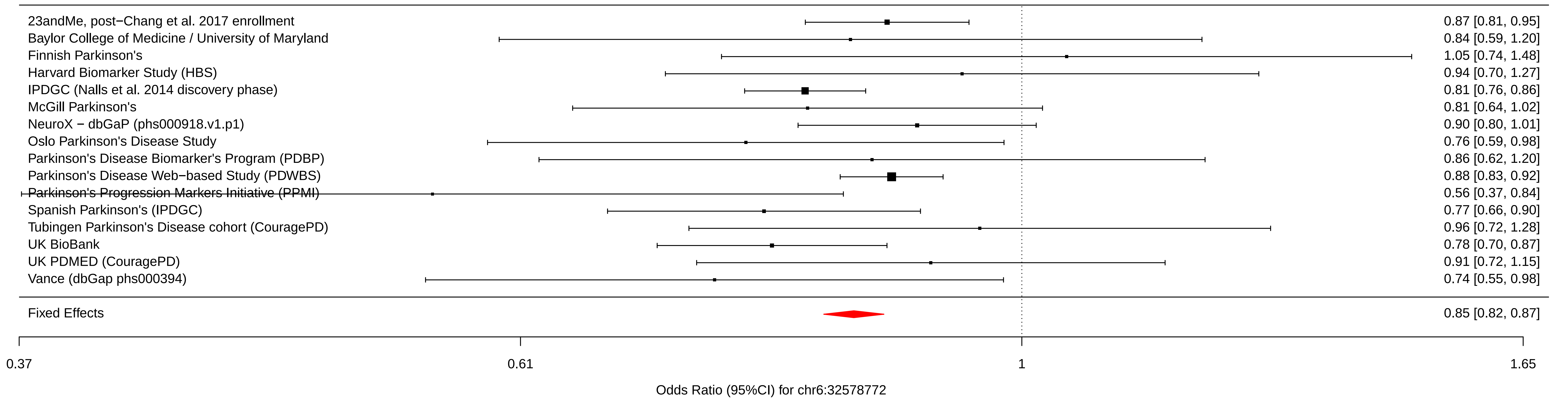

### chr18_40673380_rs12456492.pdf

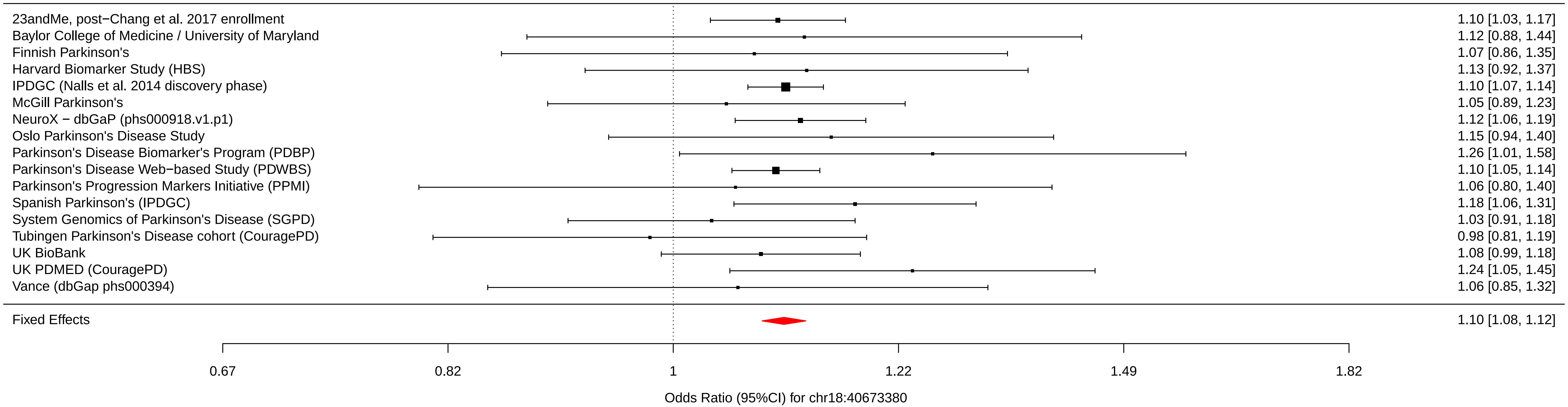

### Figure S2A. Expression enrichment by tissue from FUMA.png

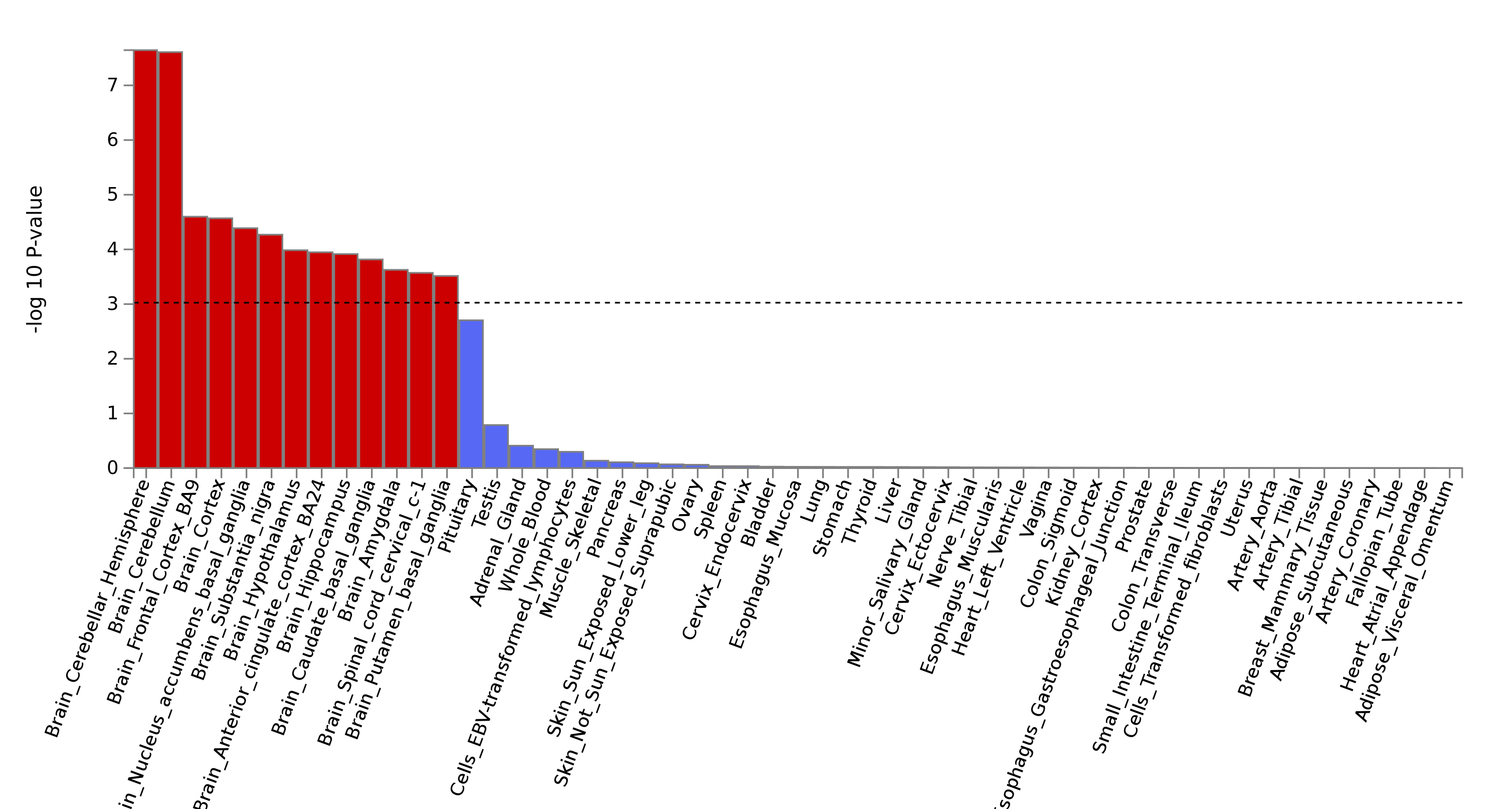

### Figure S3B. Gene level connectivity within significant networks.png

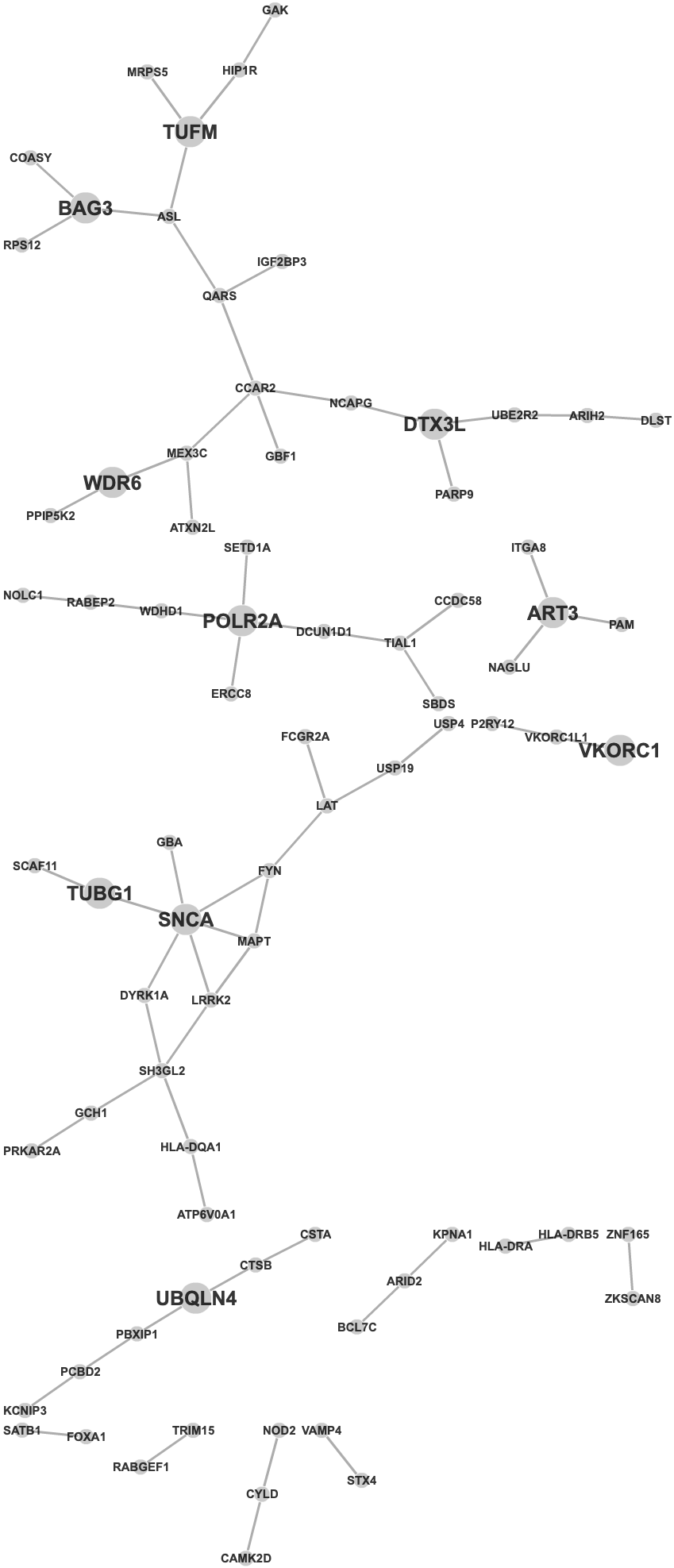

### rs26431.png

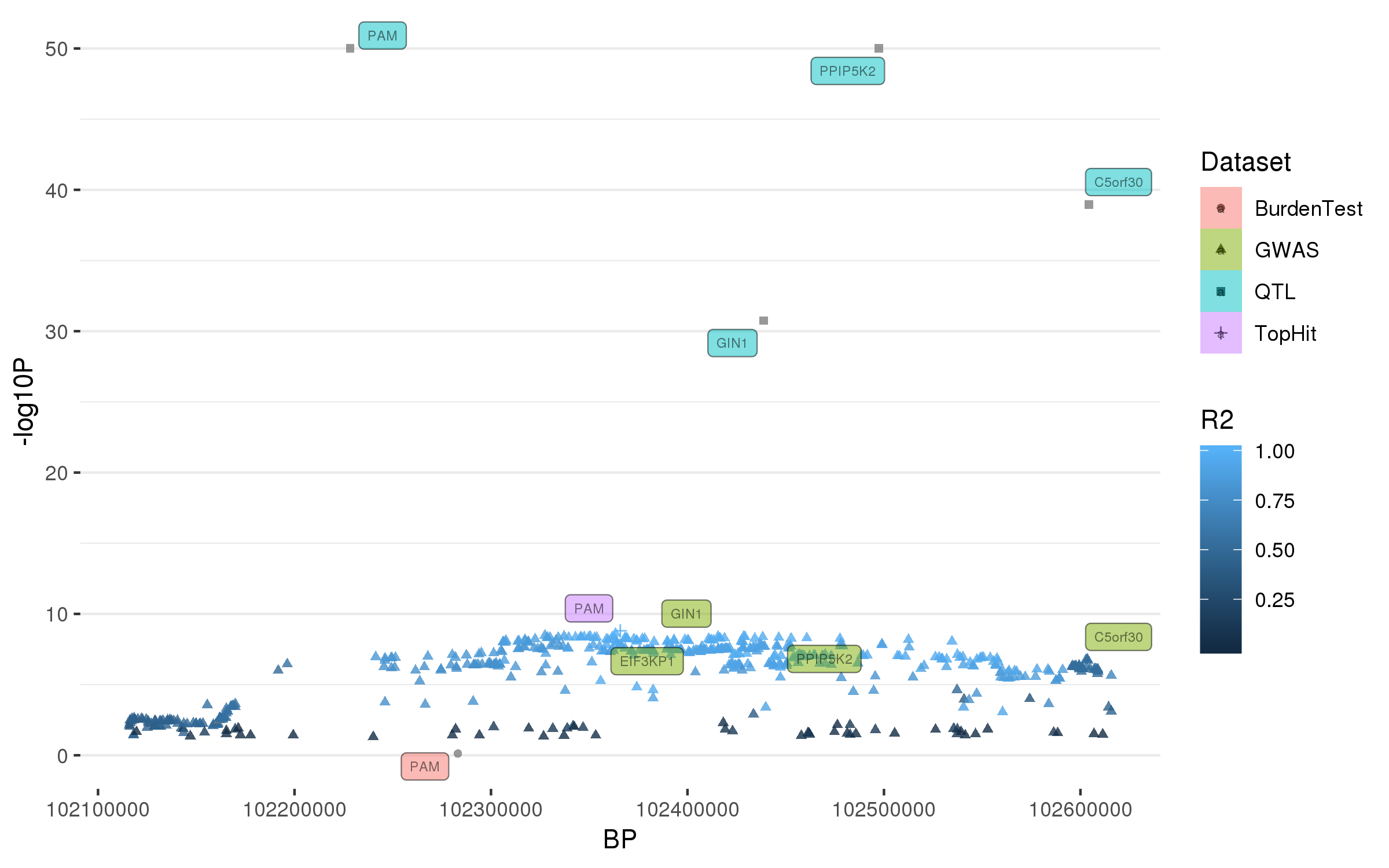

### rs199351.png

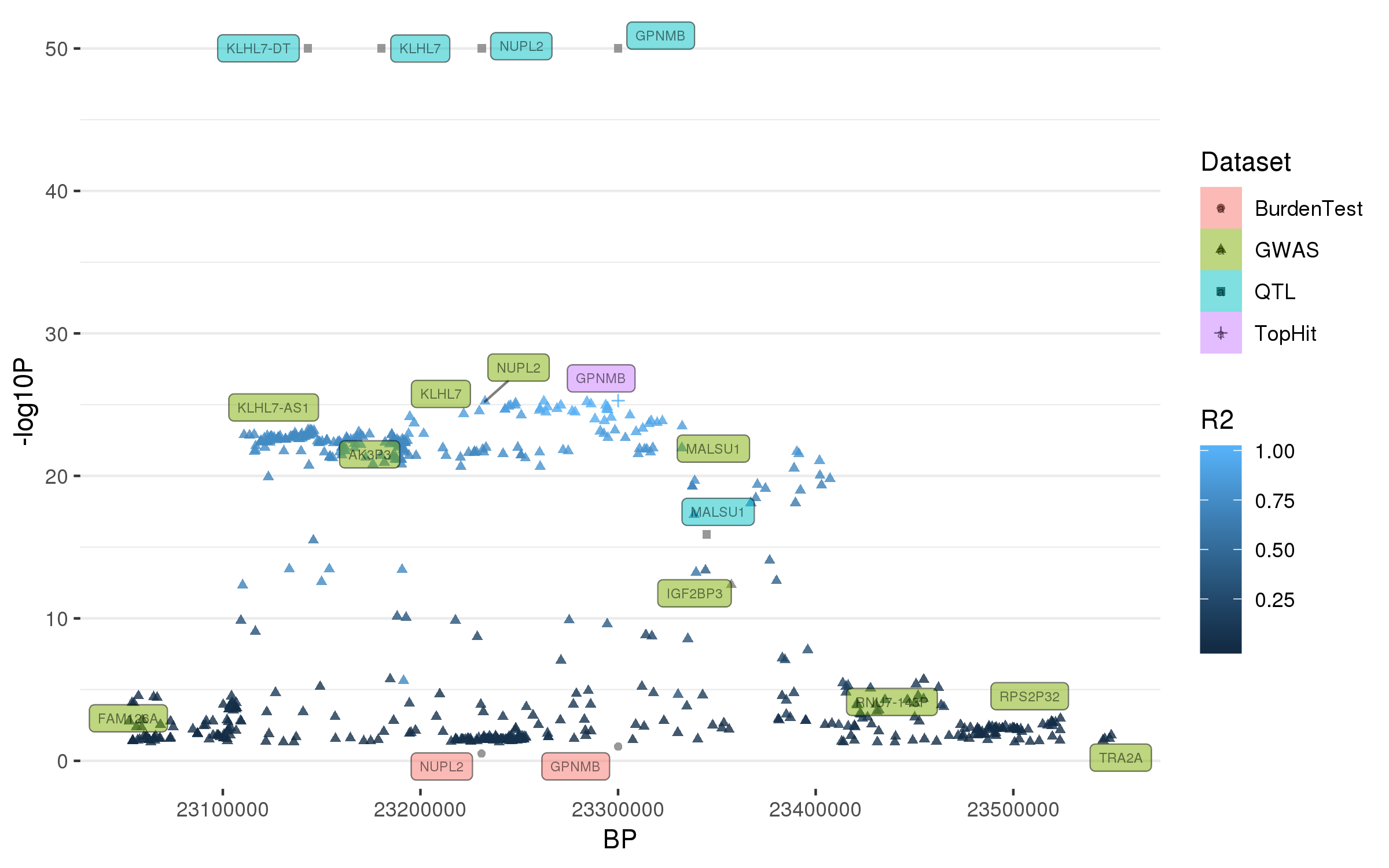

### rs356182.png

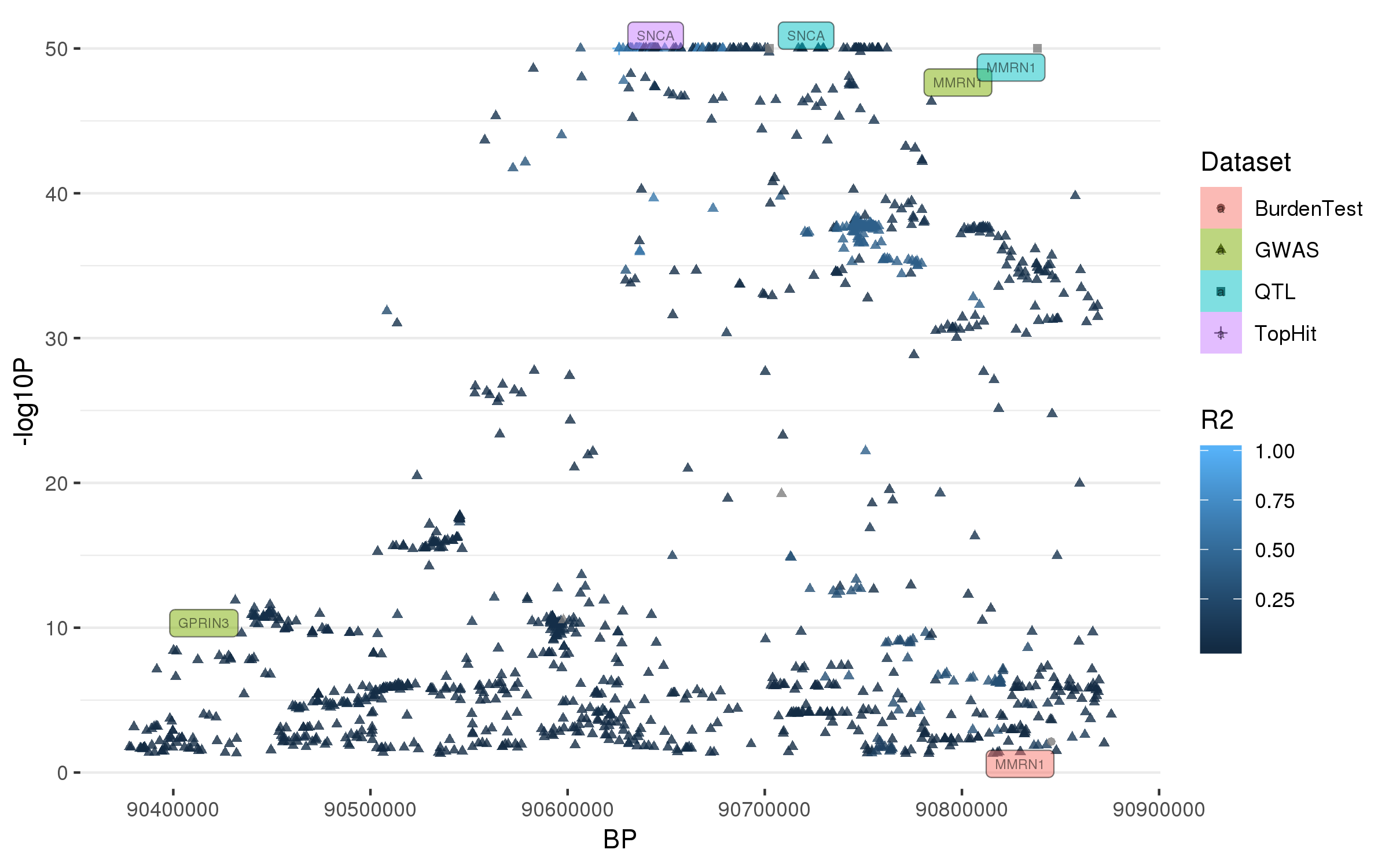

### rs620513.png

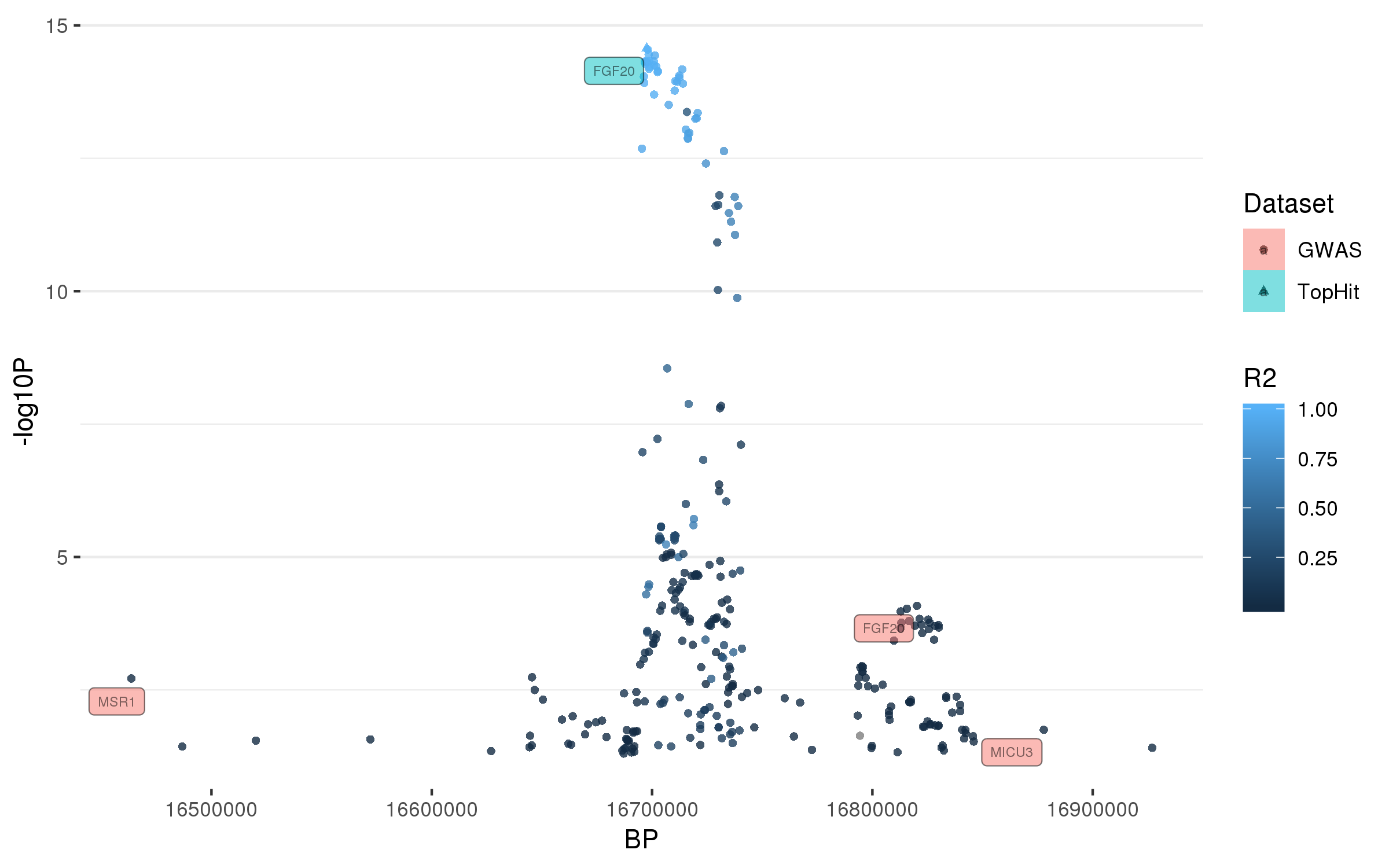

### rs666463.png

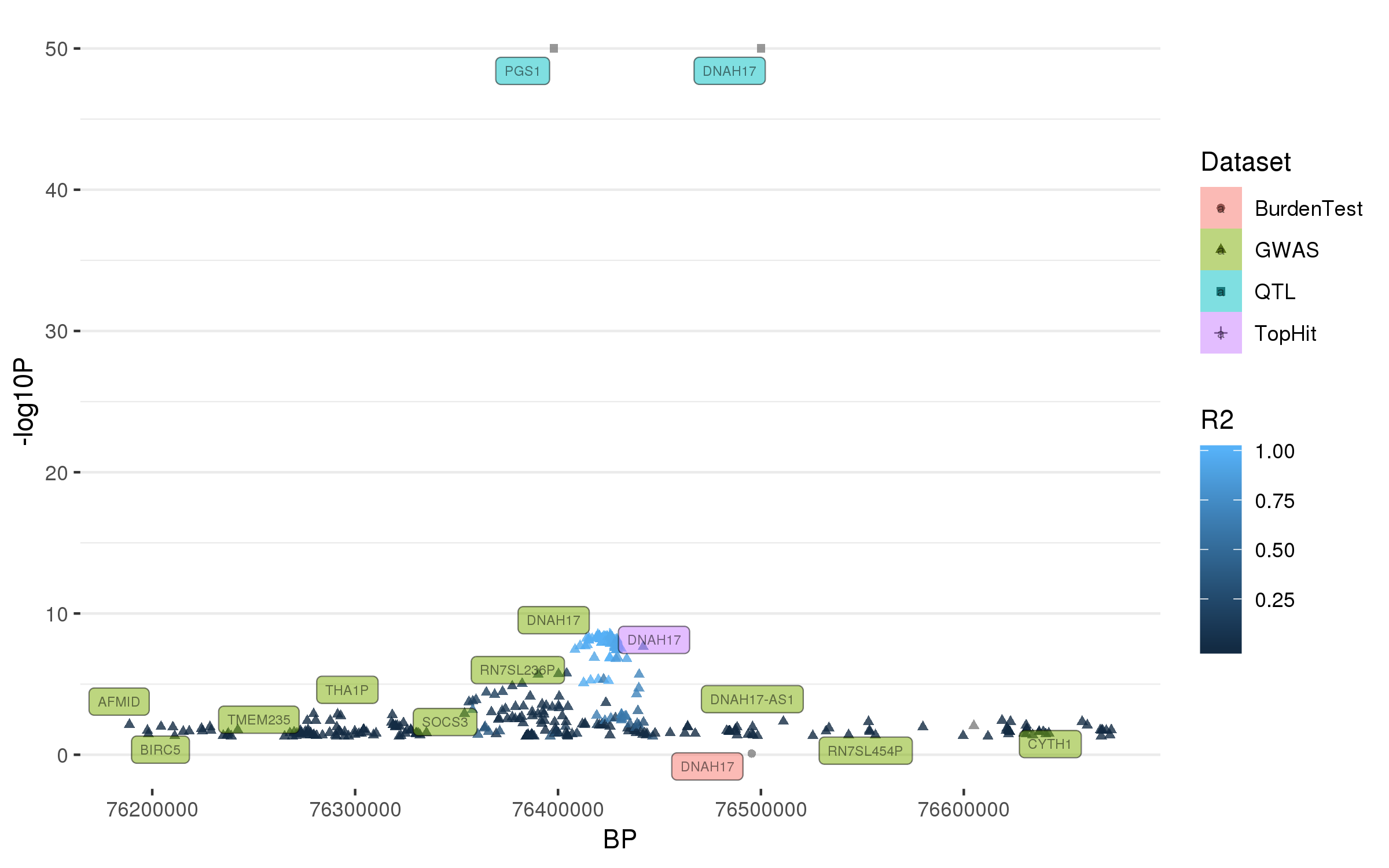

### rs823118.png

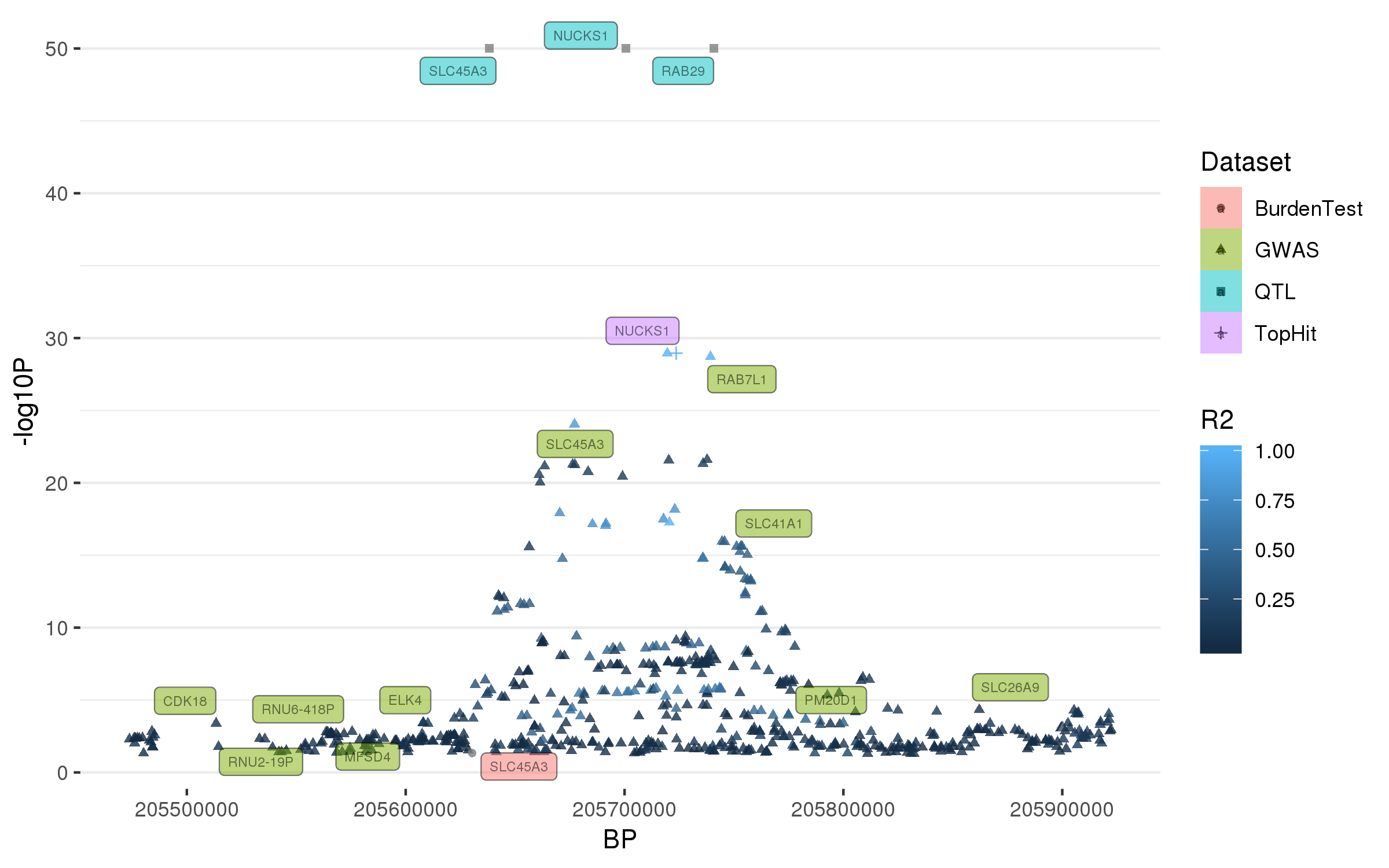

### rs850738.png

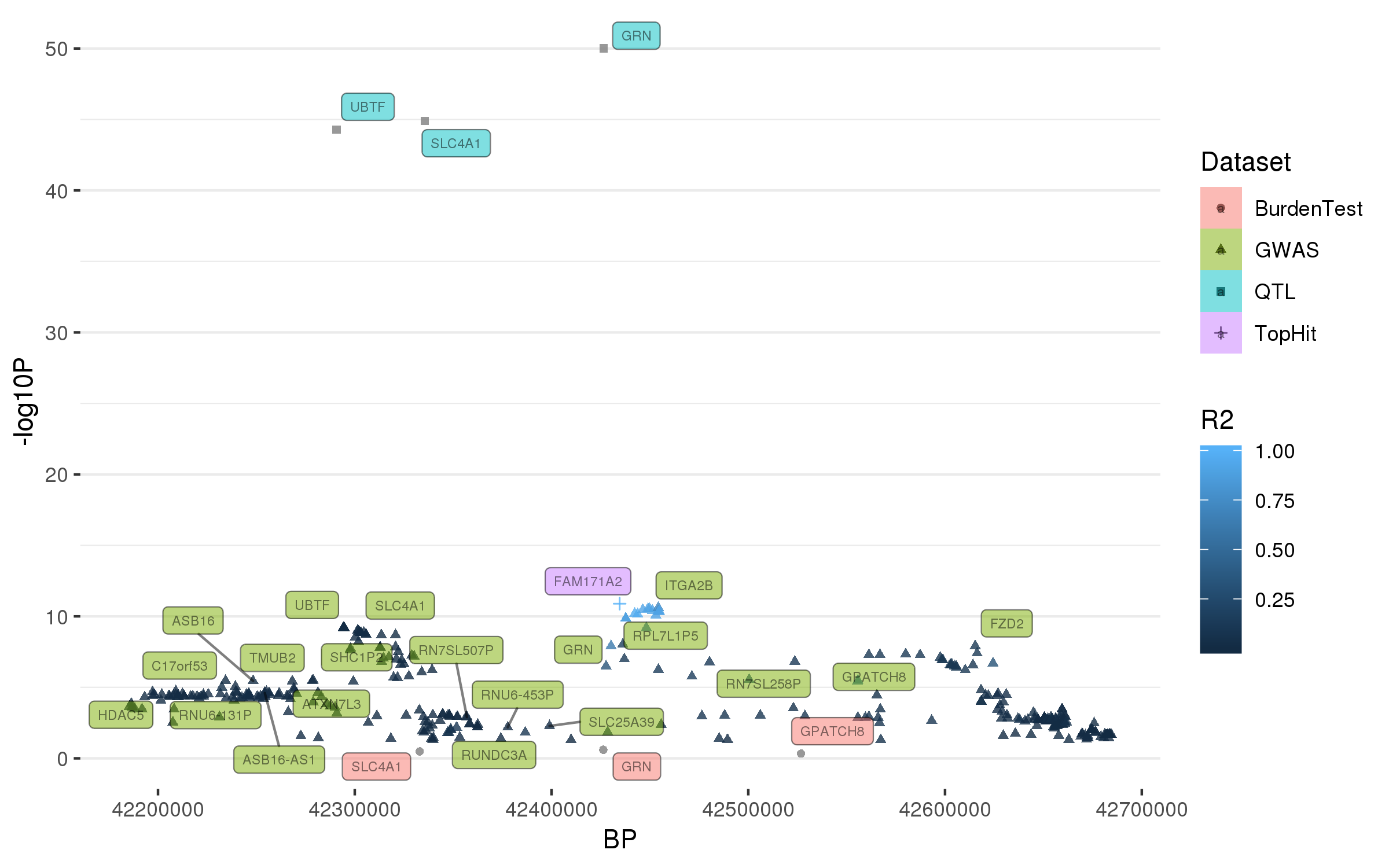

### rs873786.png

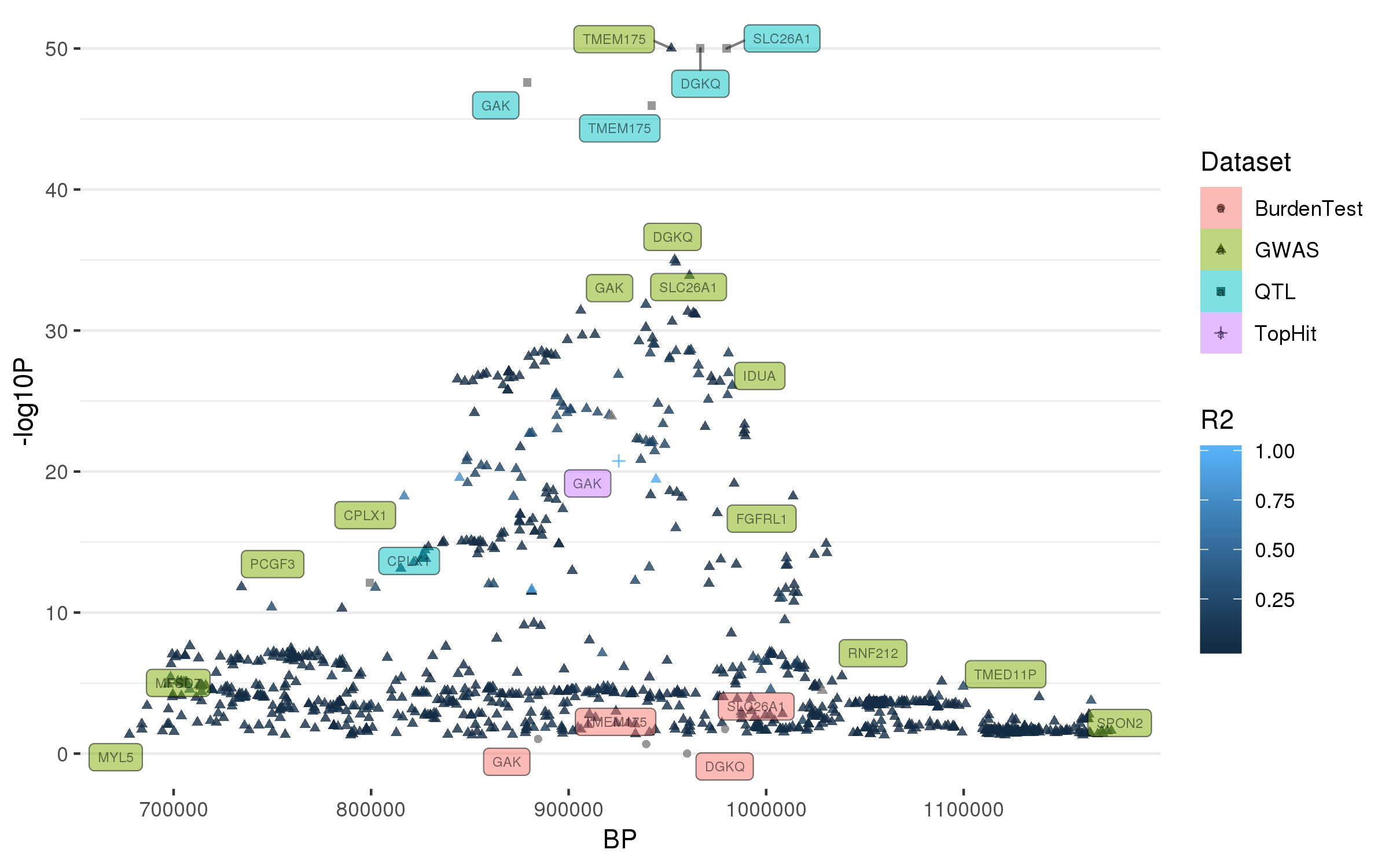

### rs896435.png

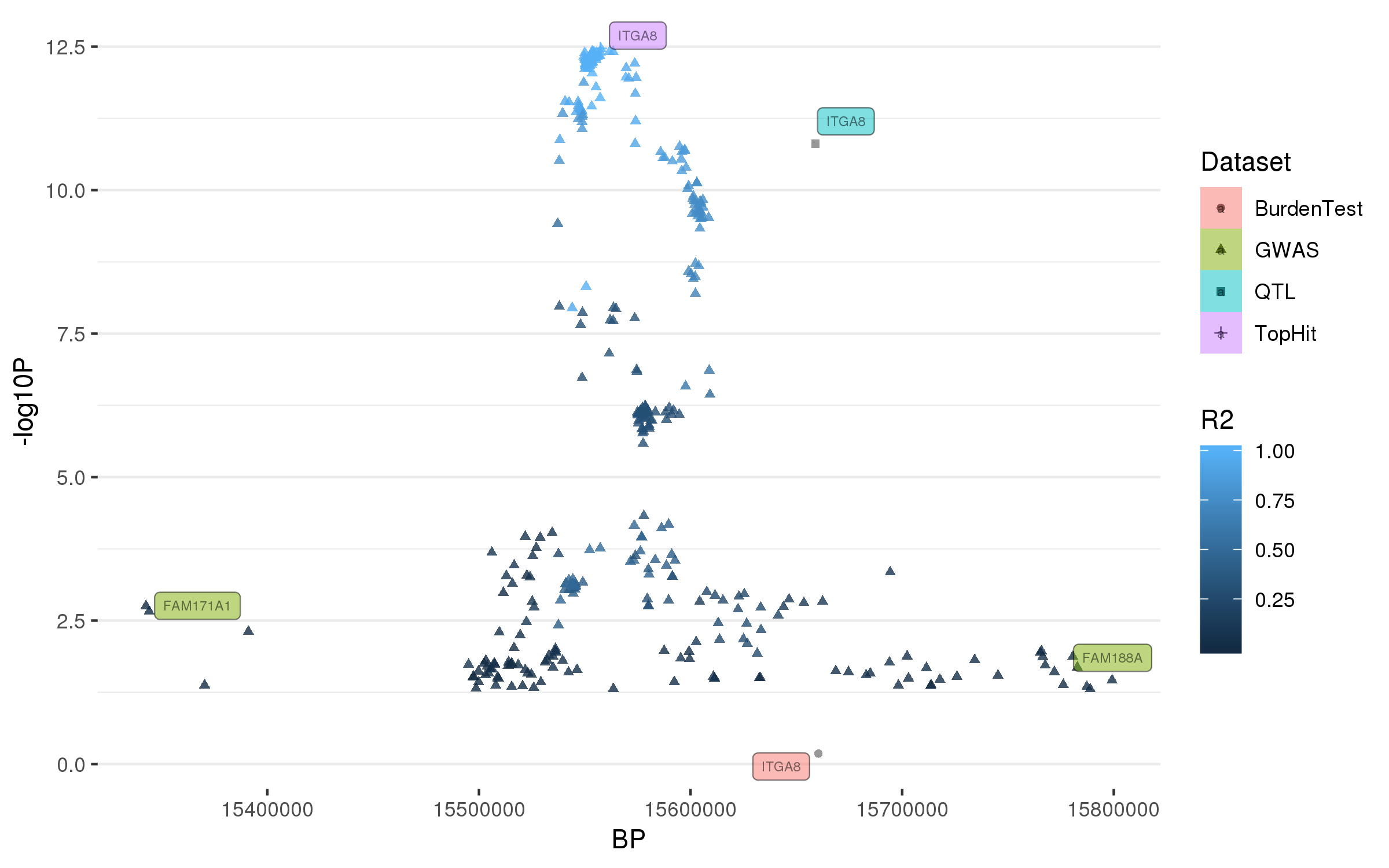

### rs979812.png

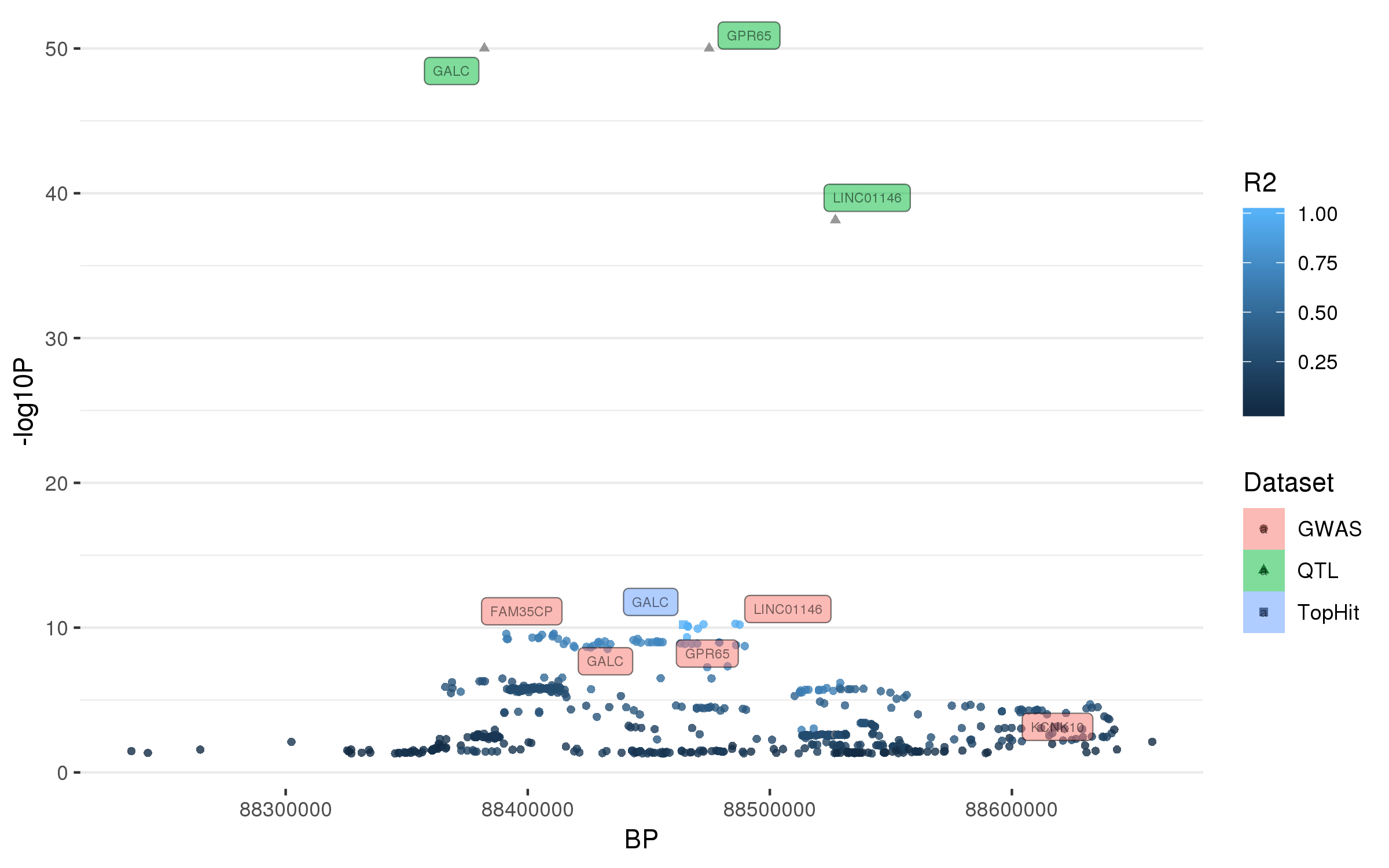

### rs997368.png

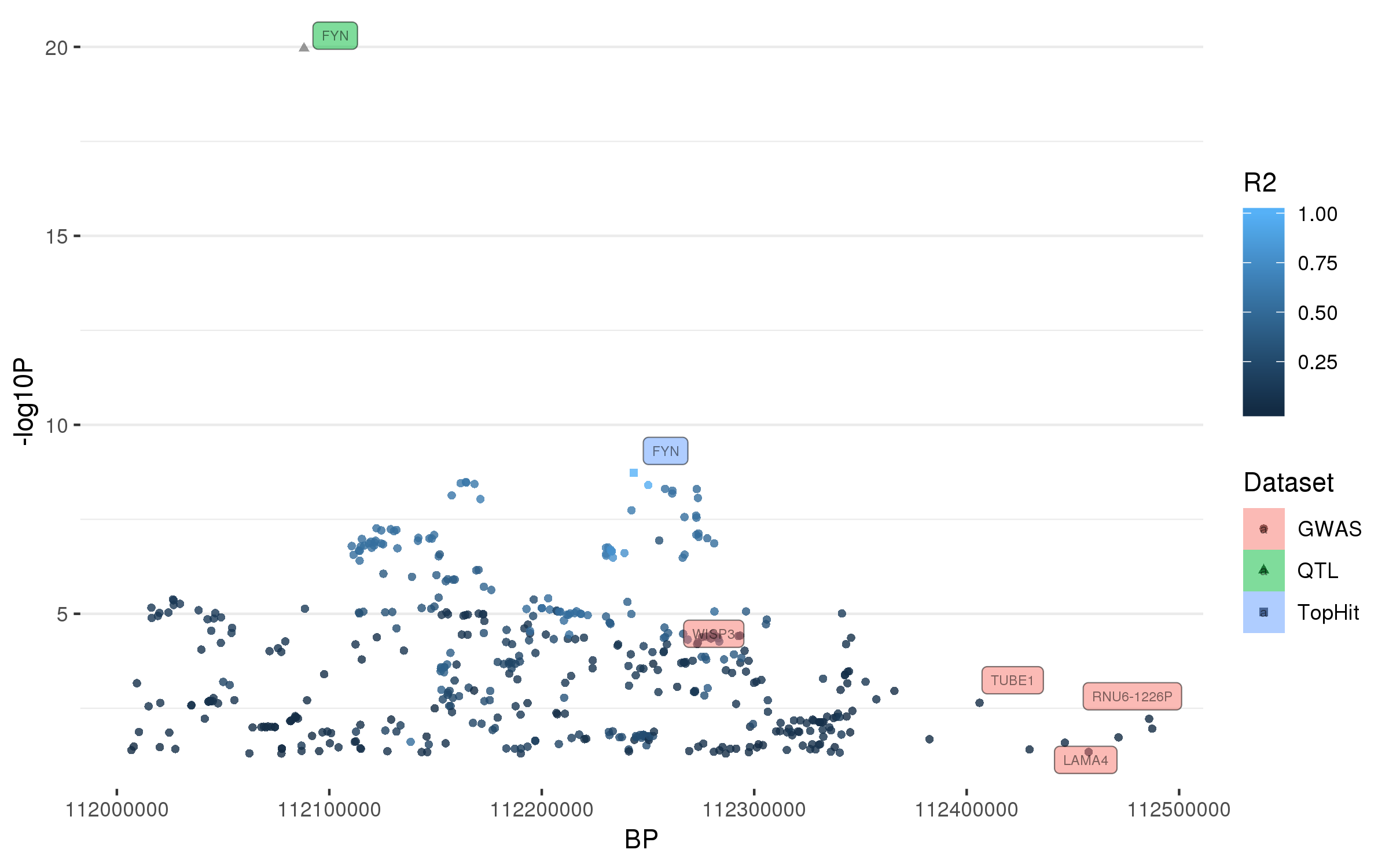

### rs1293298.png

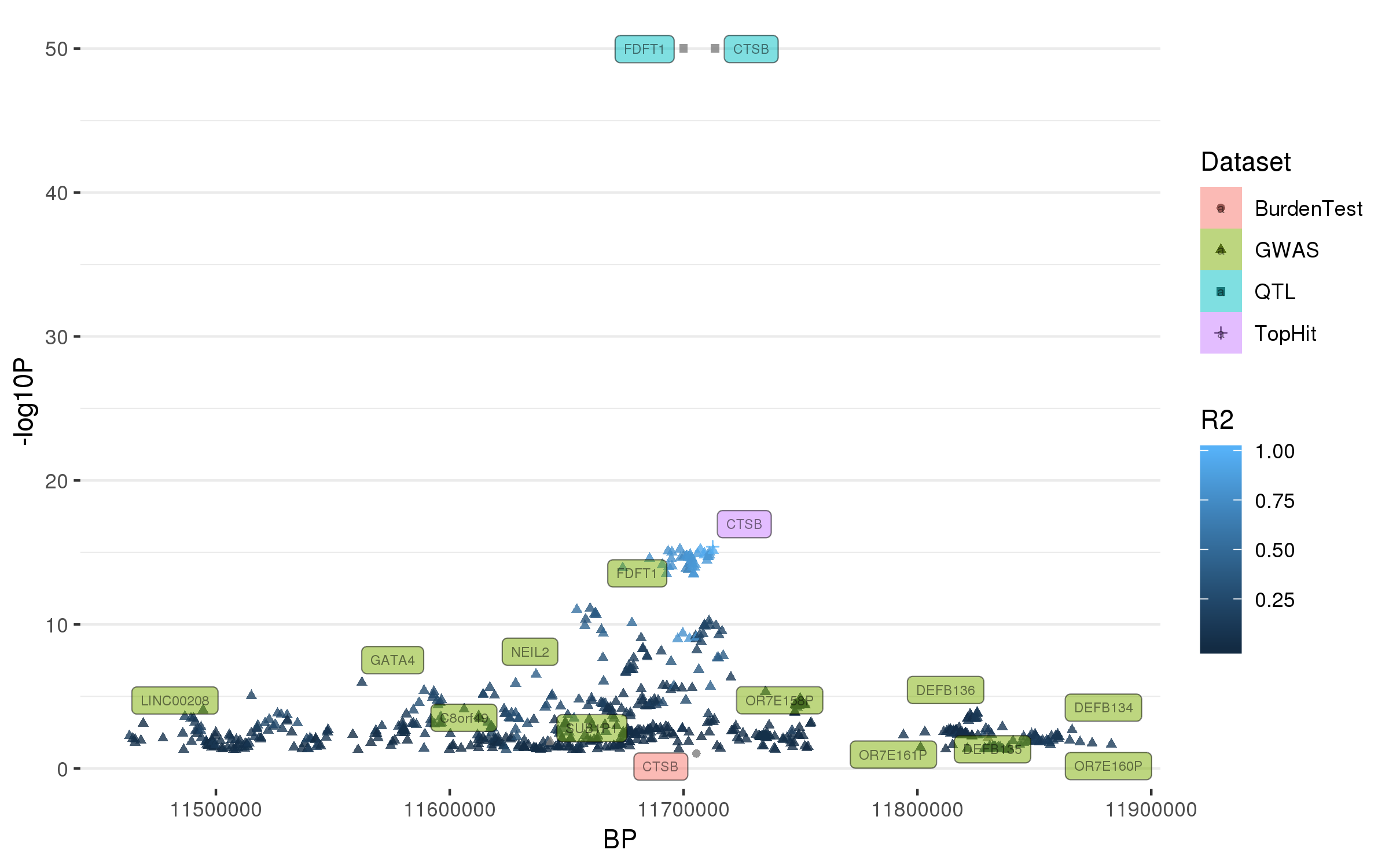

### rs1450522.png

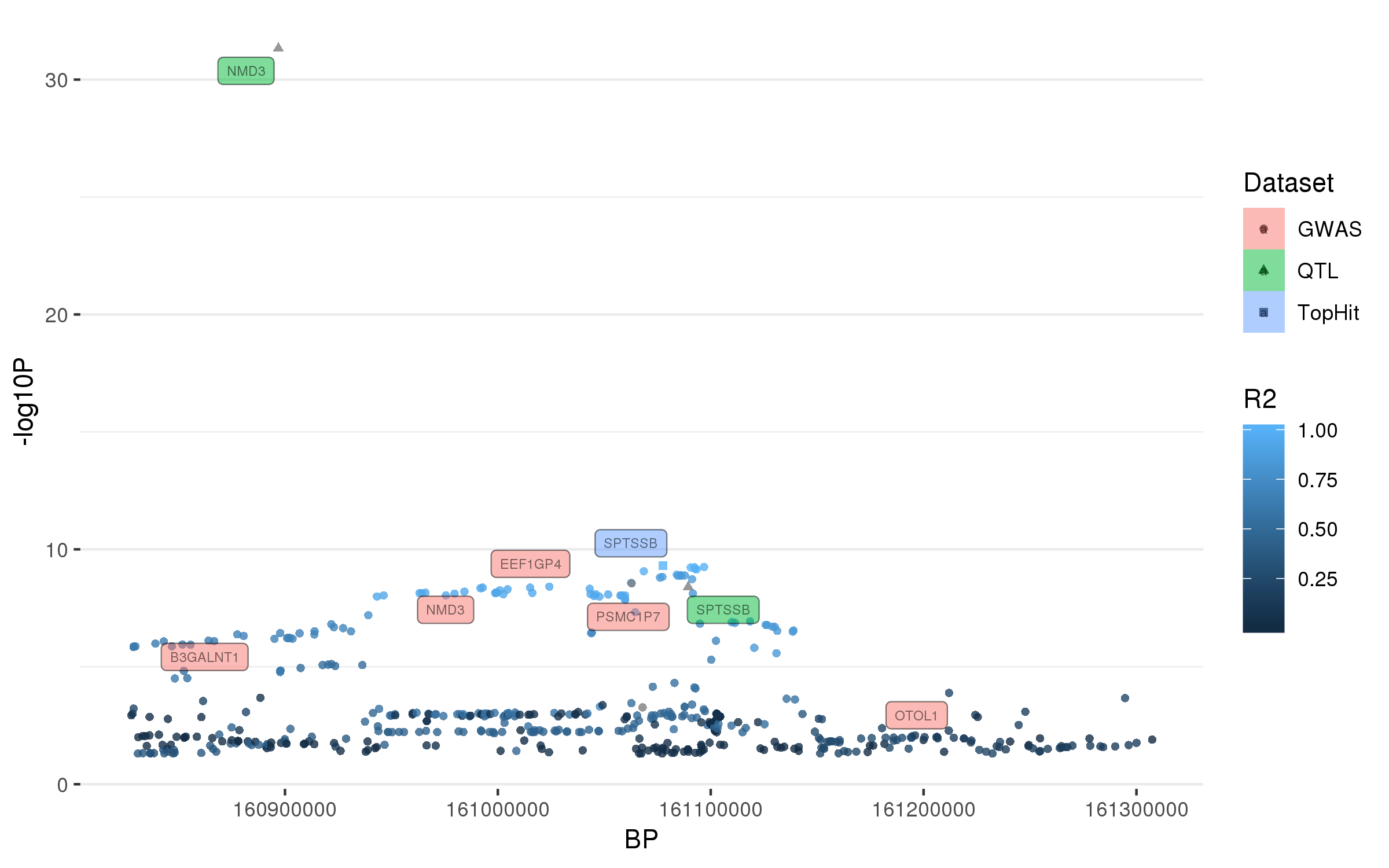

### rs1474055.png

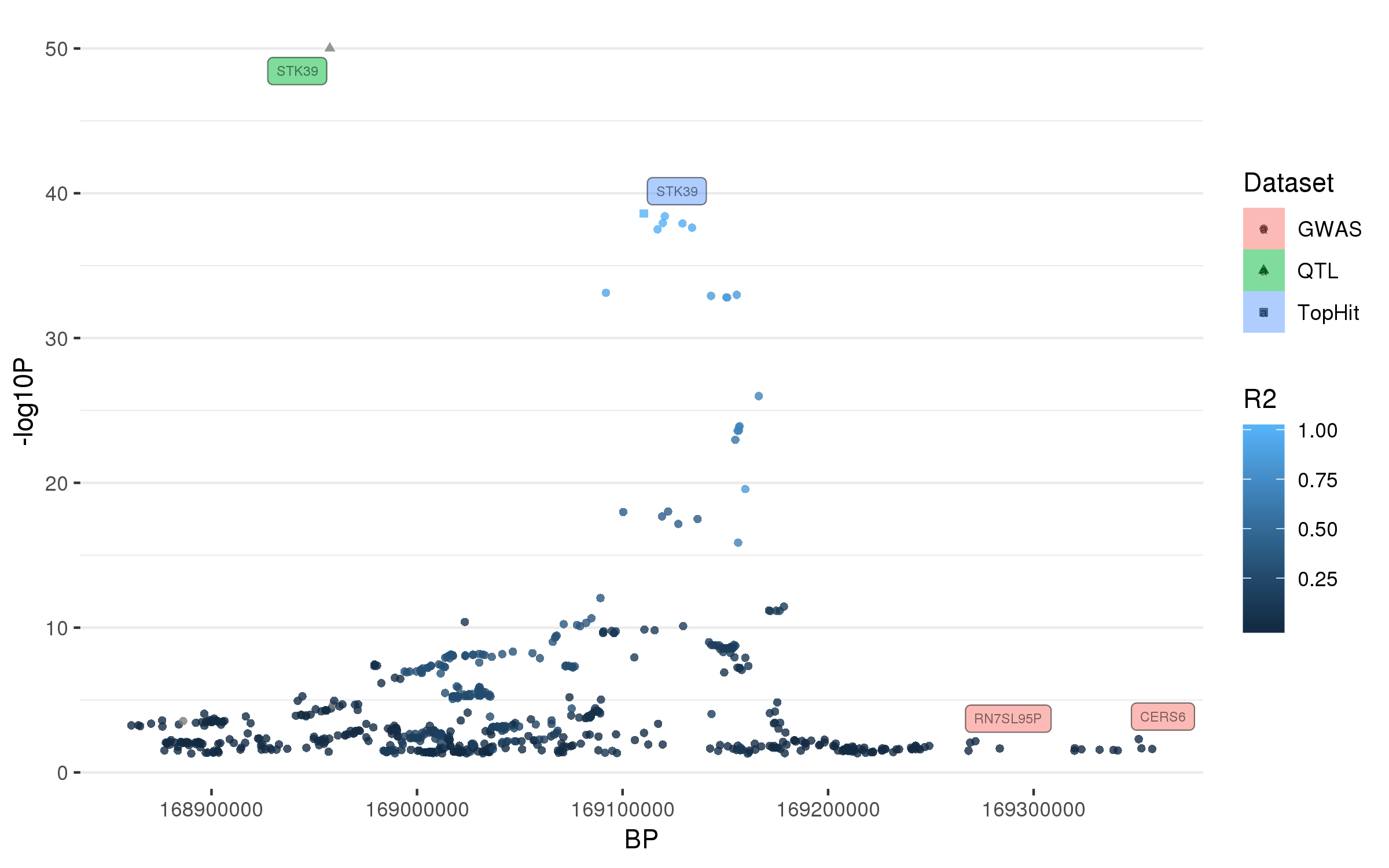

### rs1867598.png

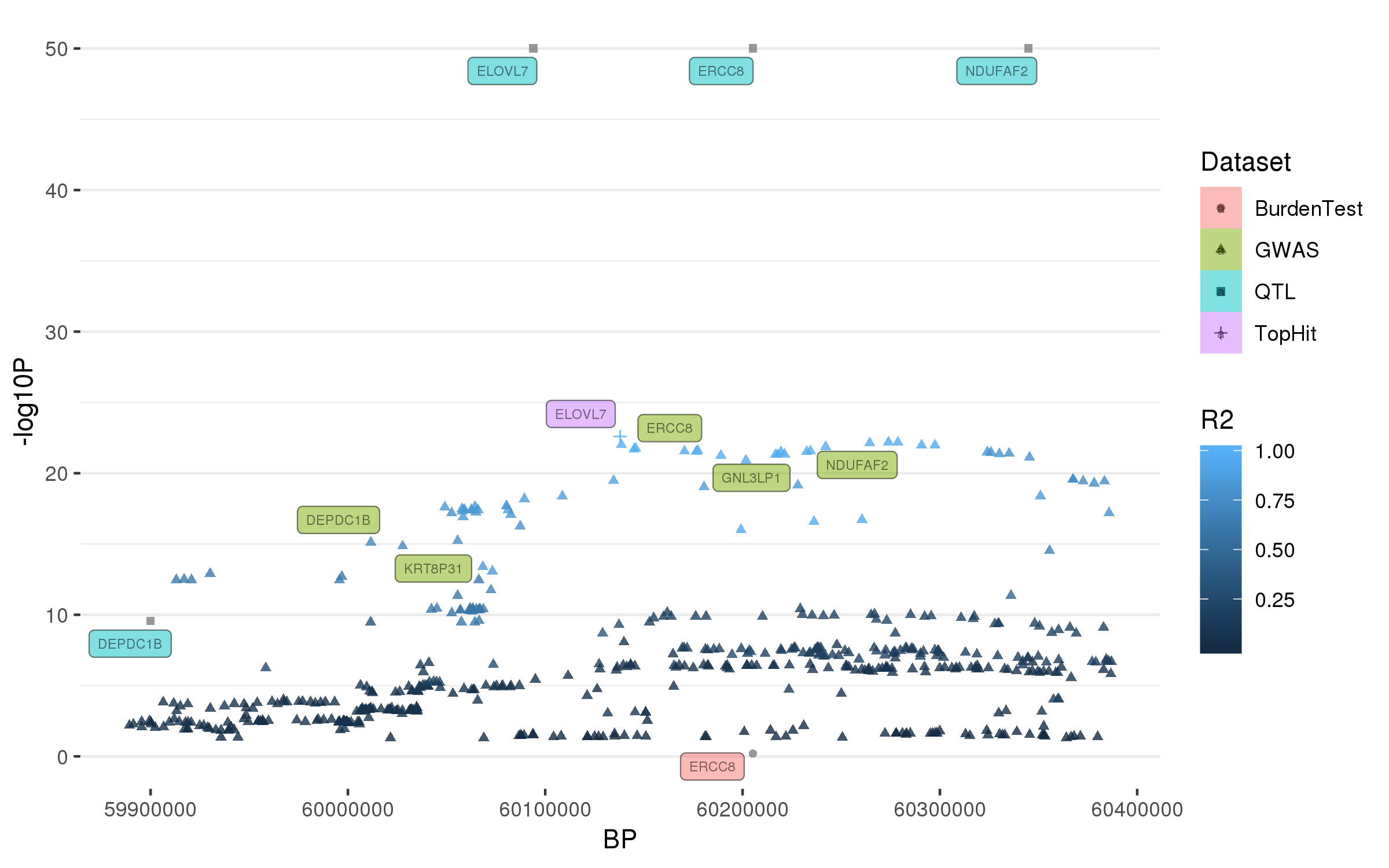

### rs1941685.png

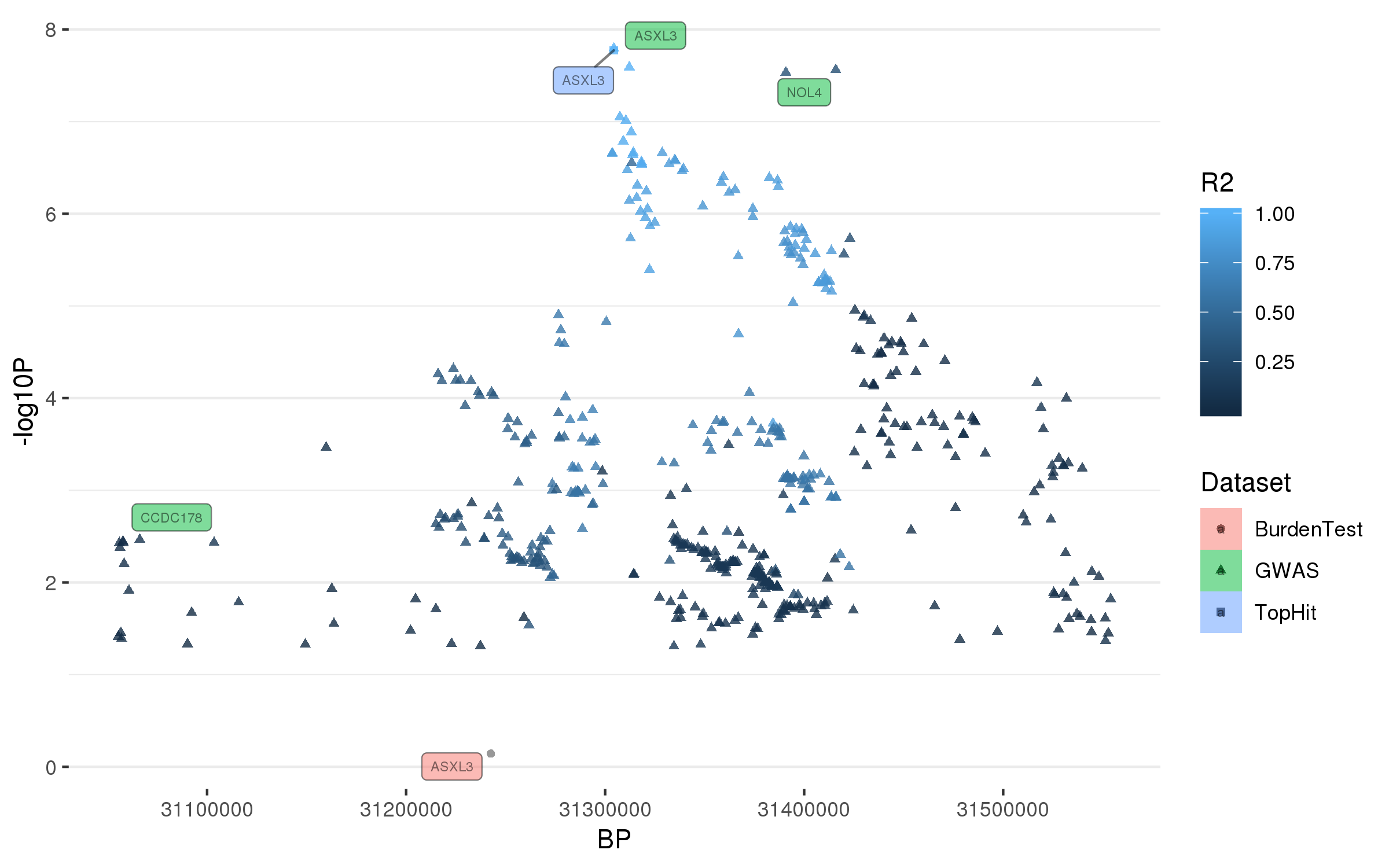

### rs2042477.png

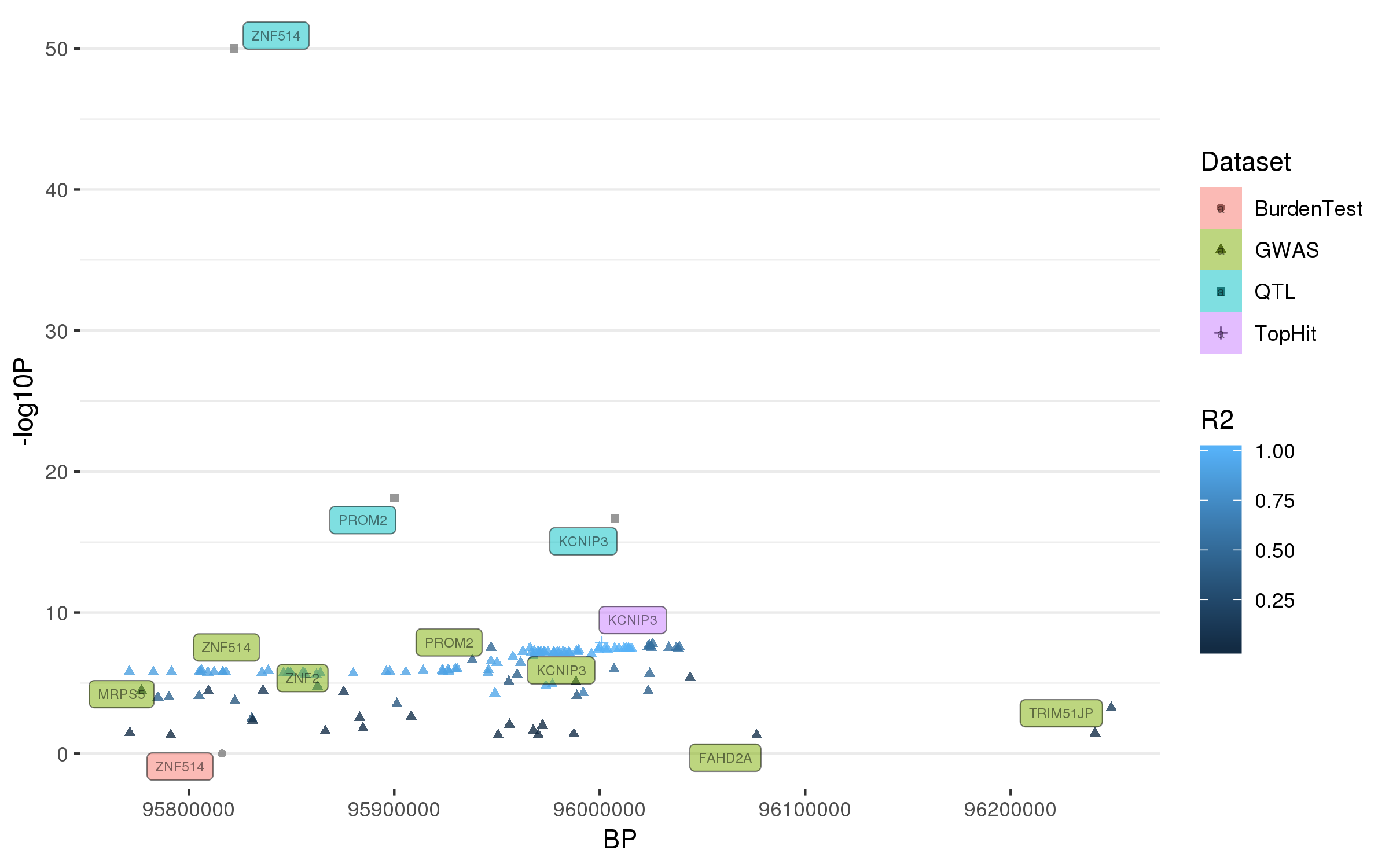

### rs2086641.png

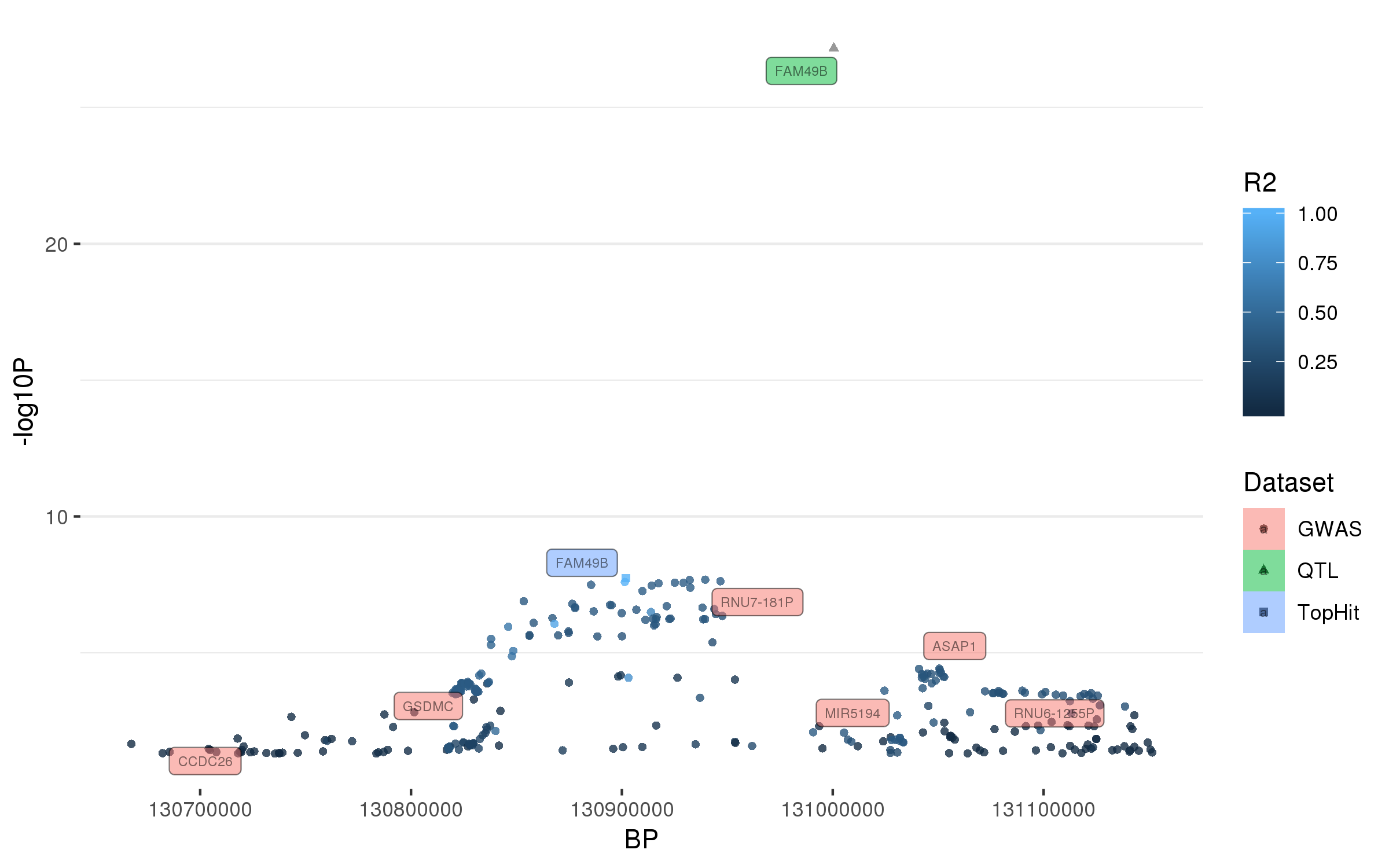

### rs2248244.png

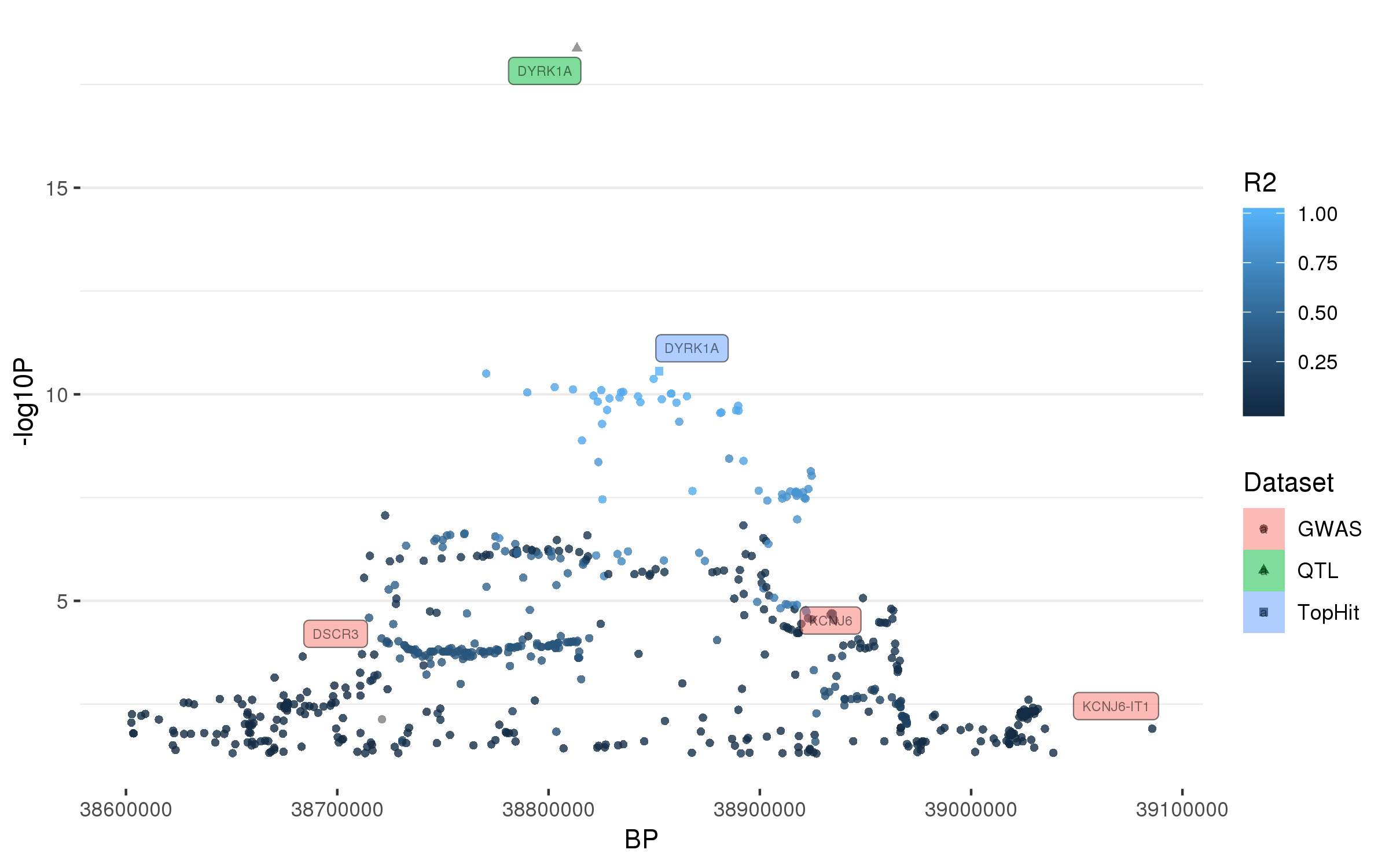

### rs2251086.png

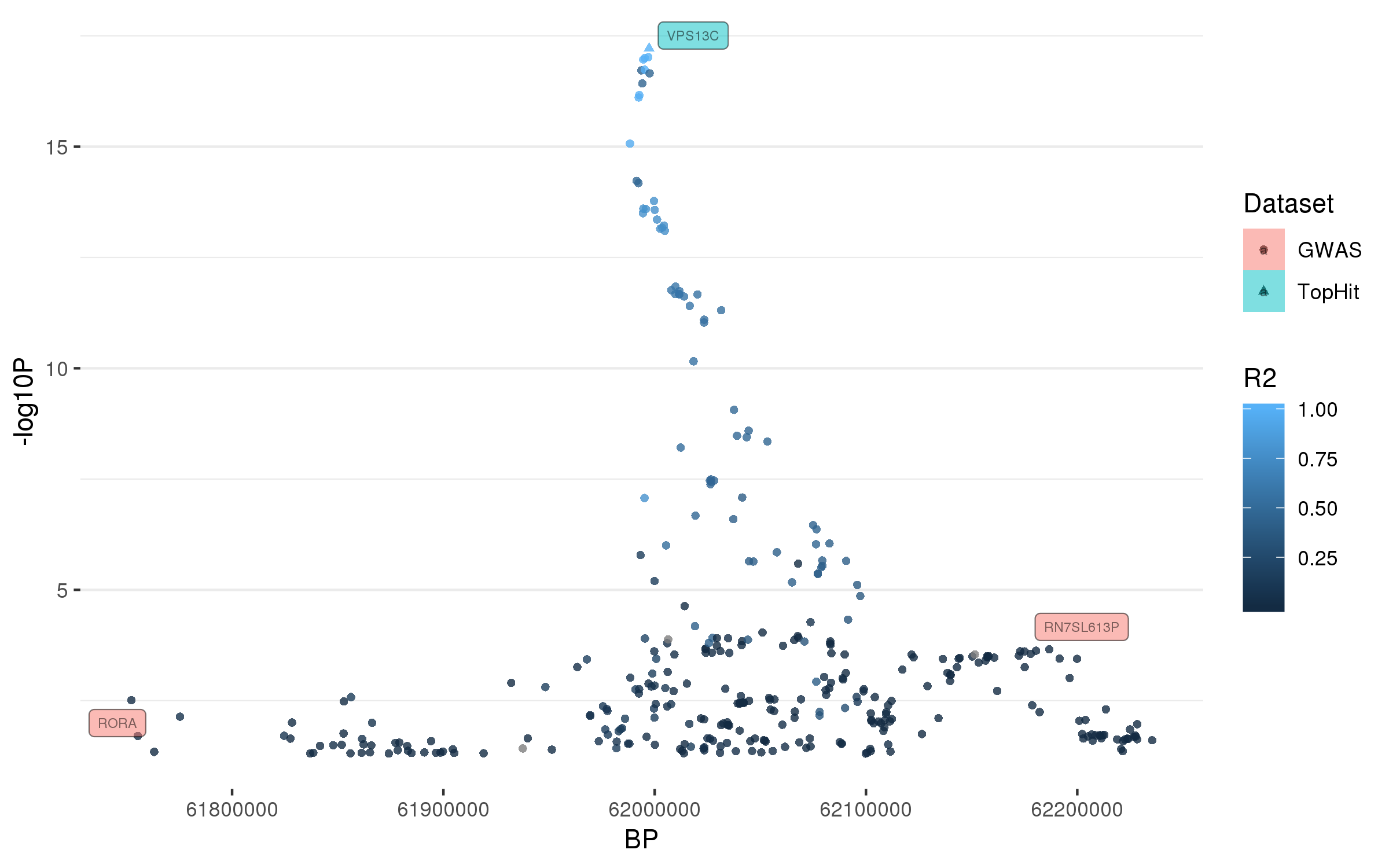

### rs2269906.png

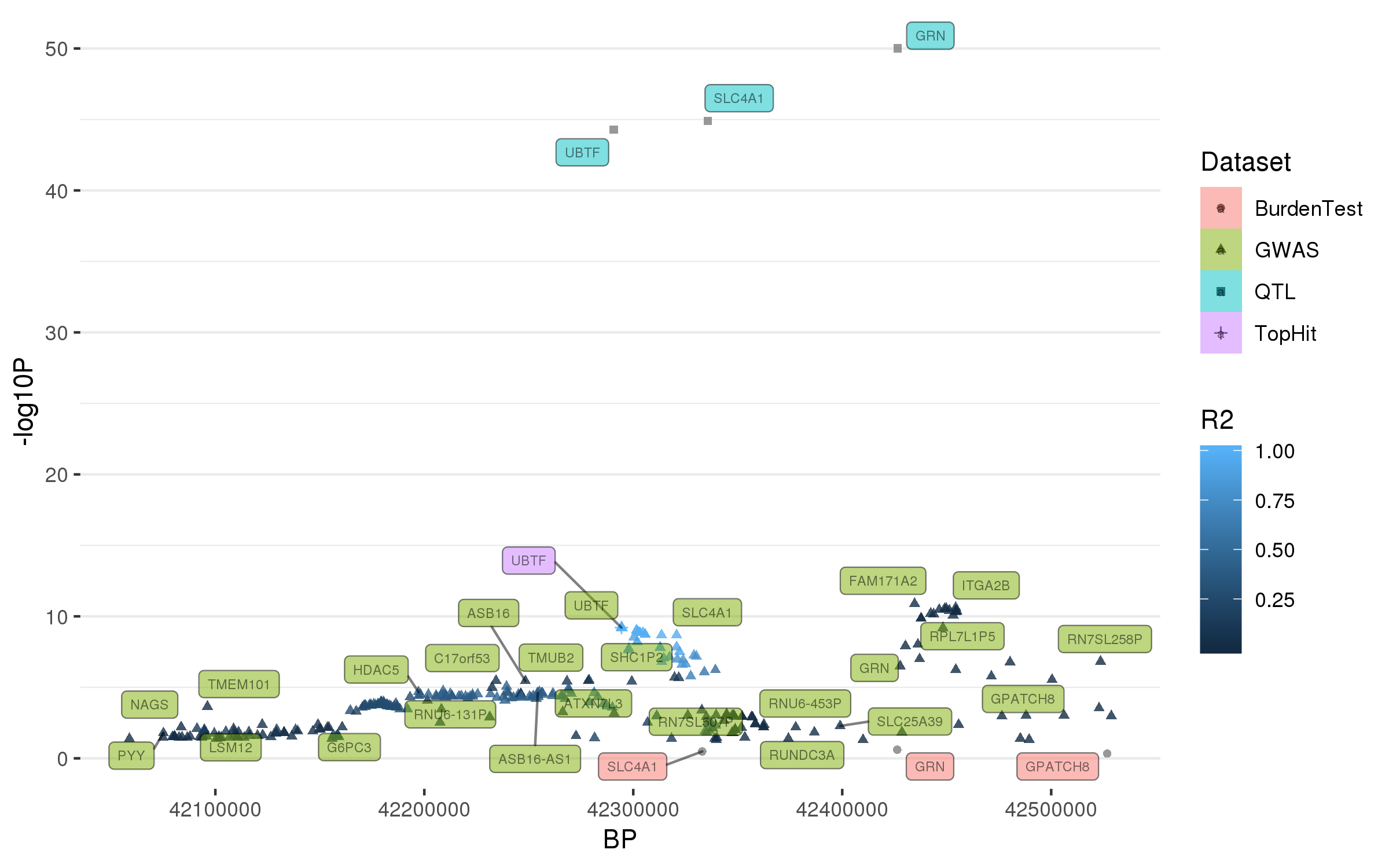

### rs2280104.png

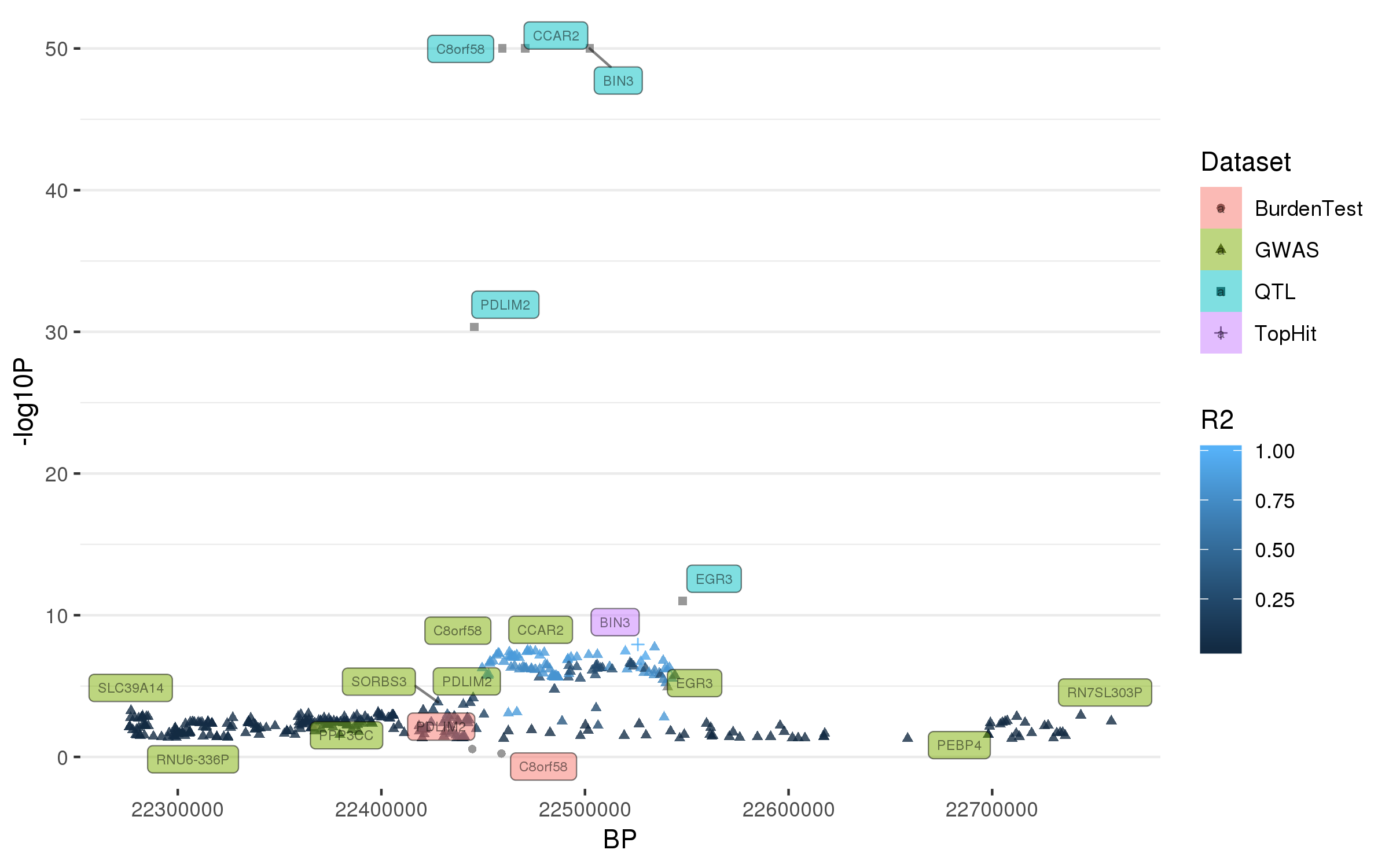

### rs2904880.png

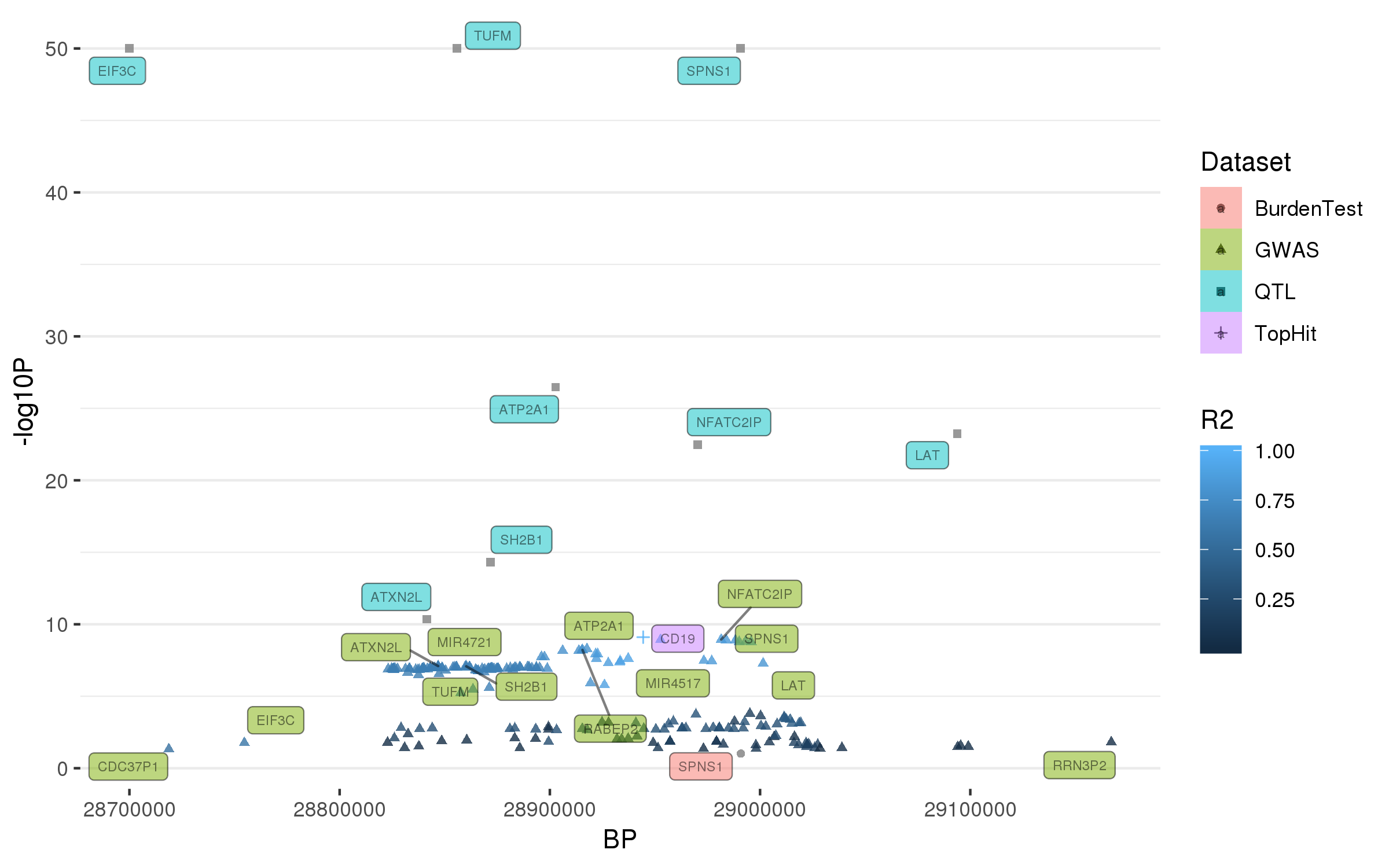

### rs3104783.png

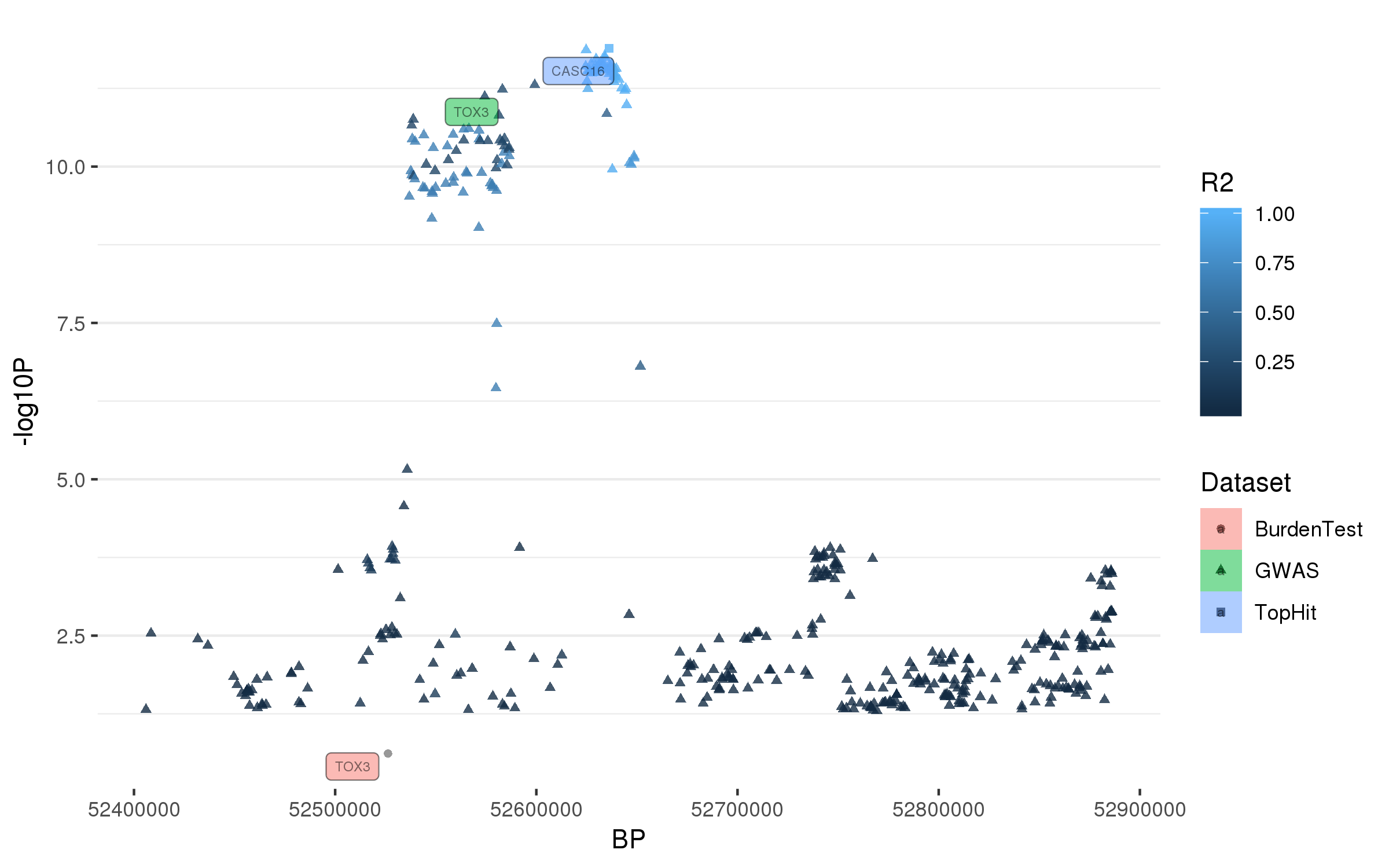

### rs3742785.png

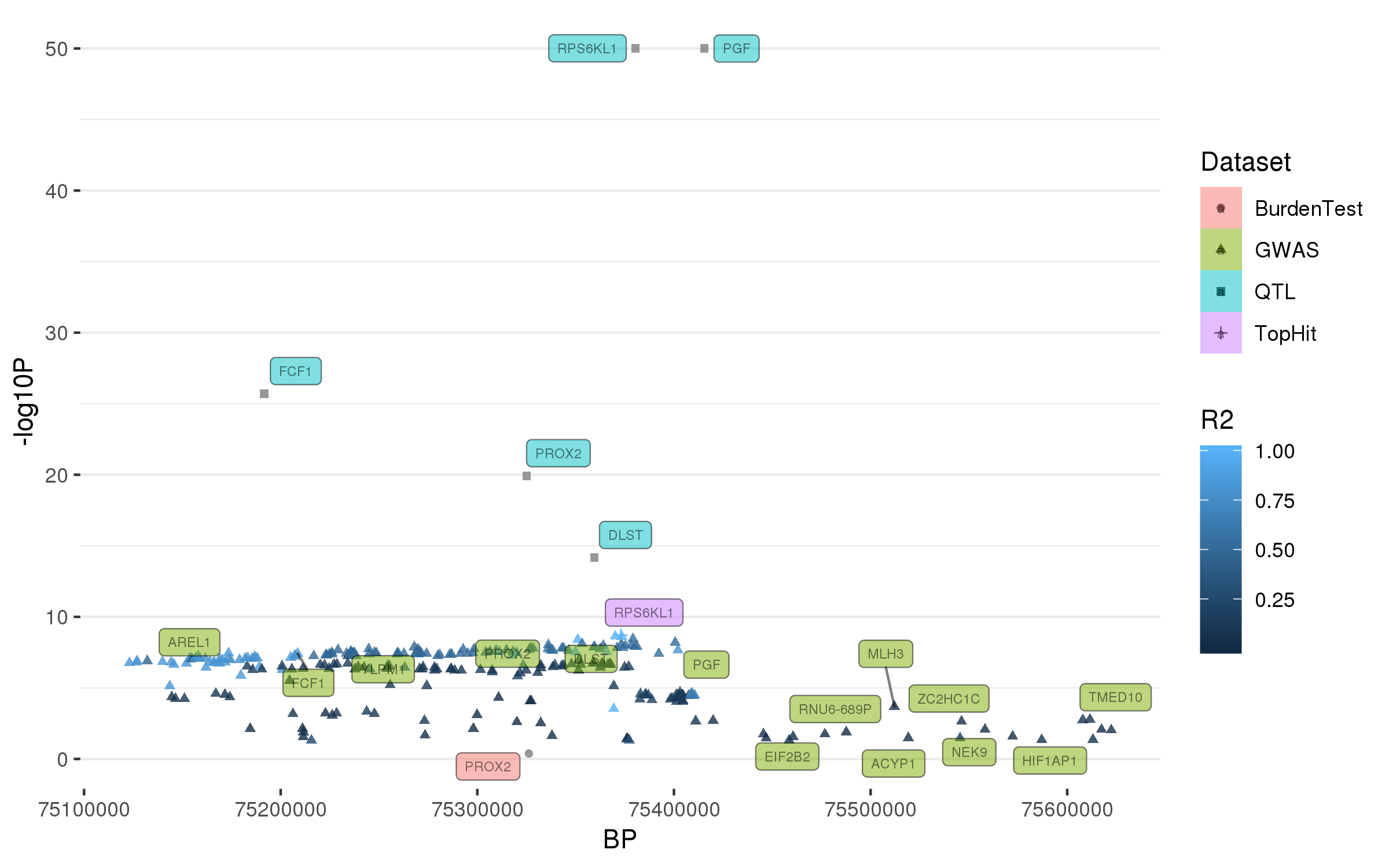

### rs3802920.png

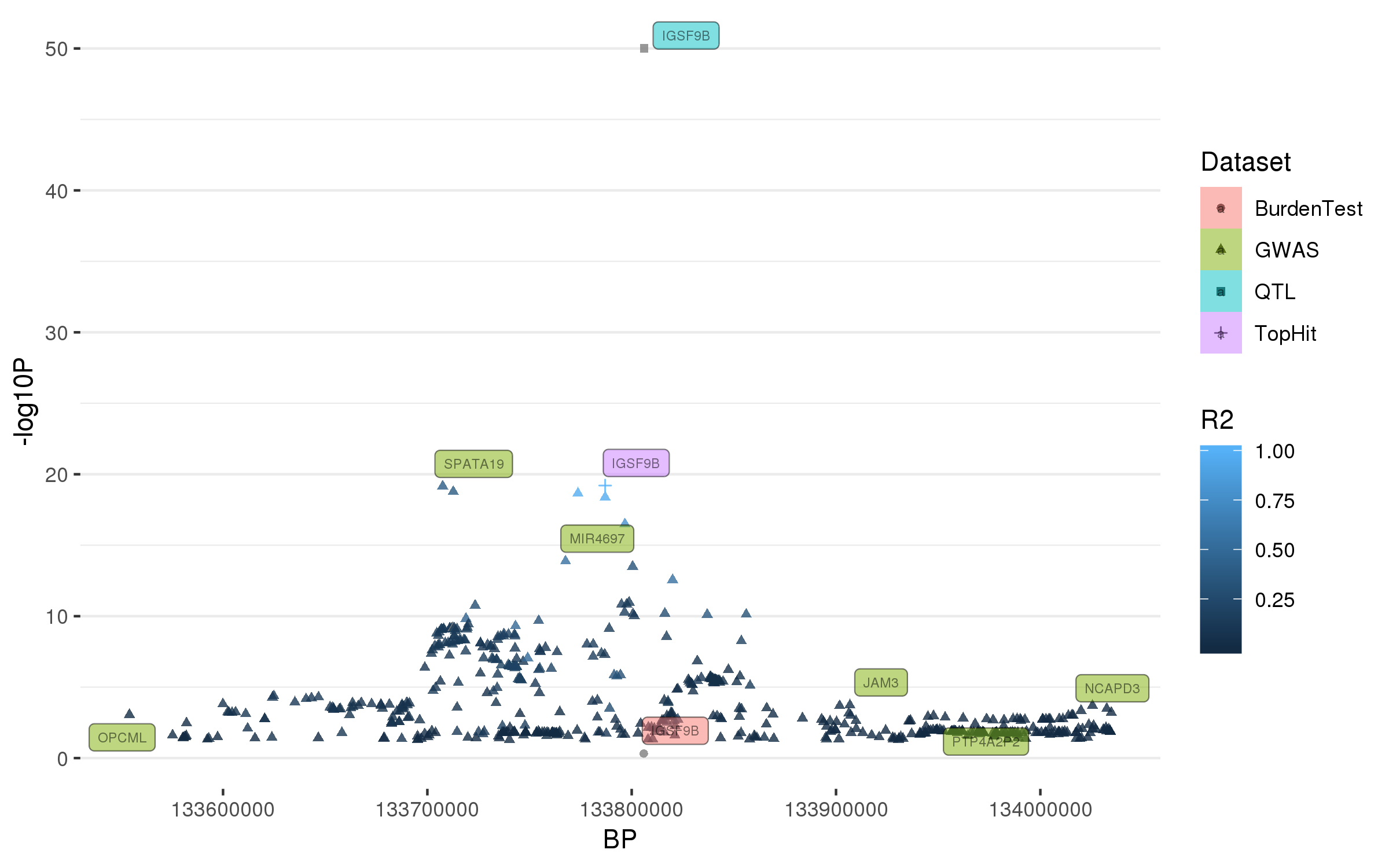

### rs4101061.png

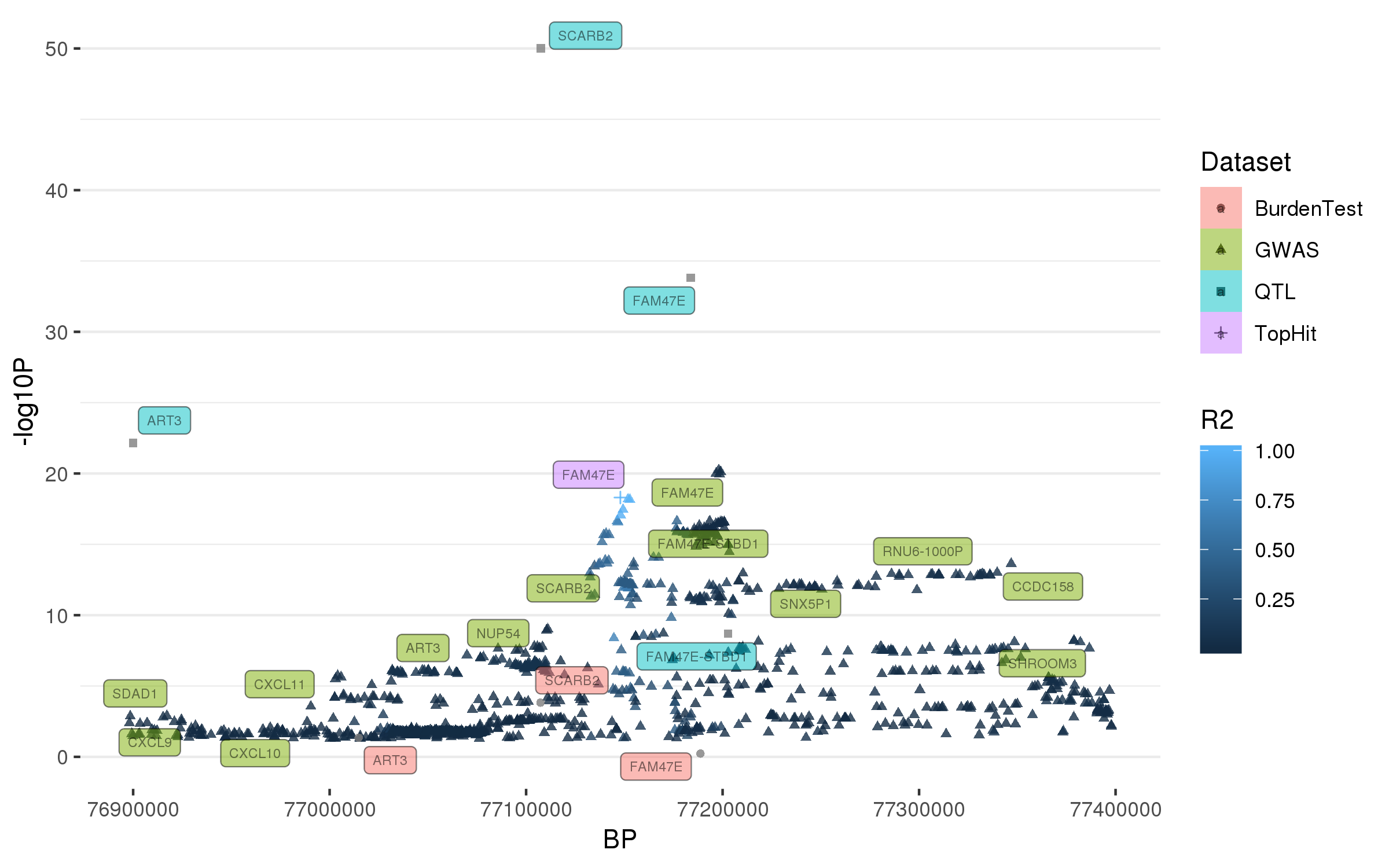

### rs4140646.png

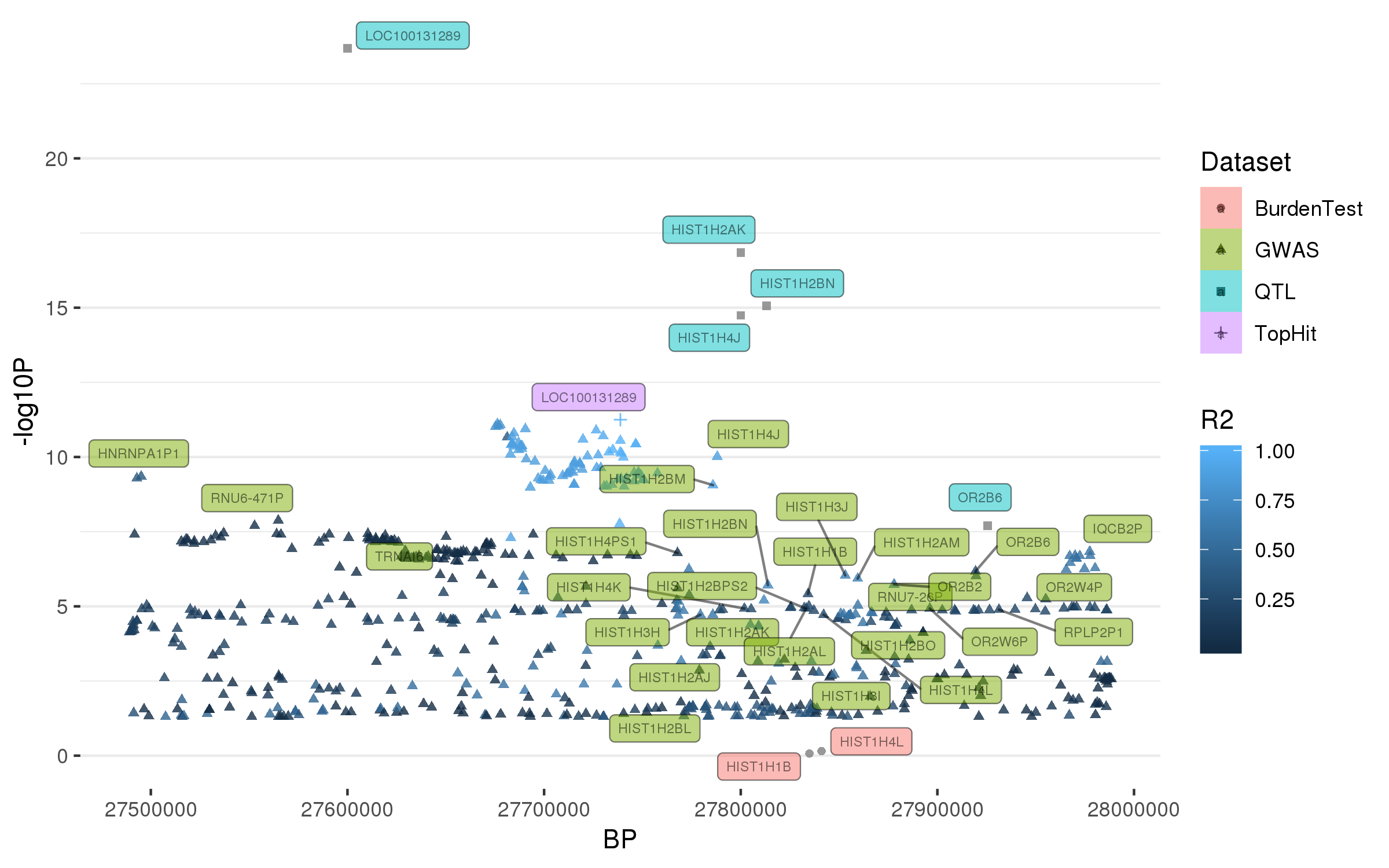

### rs4653767.png

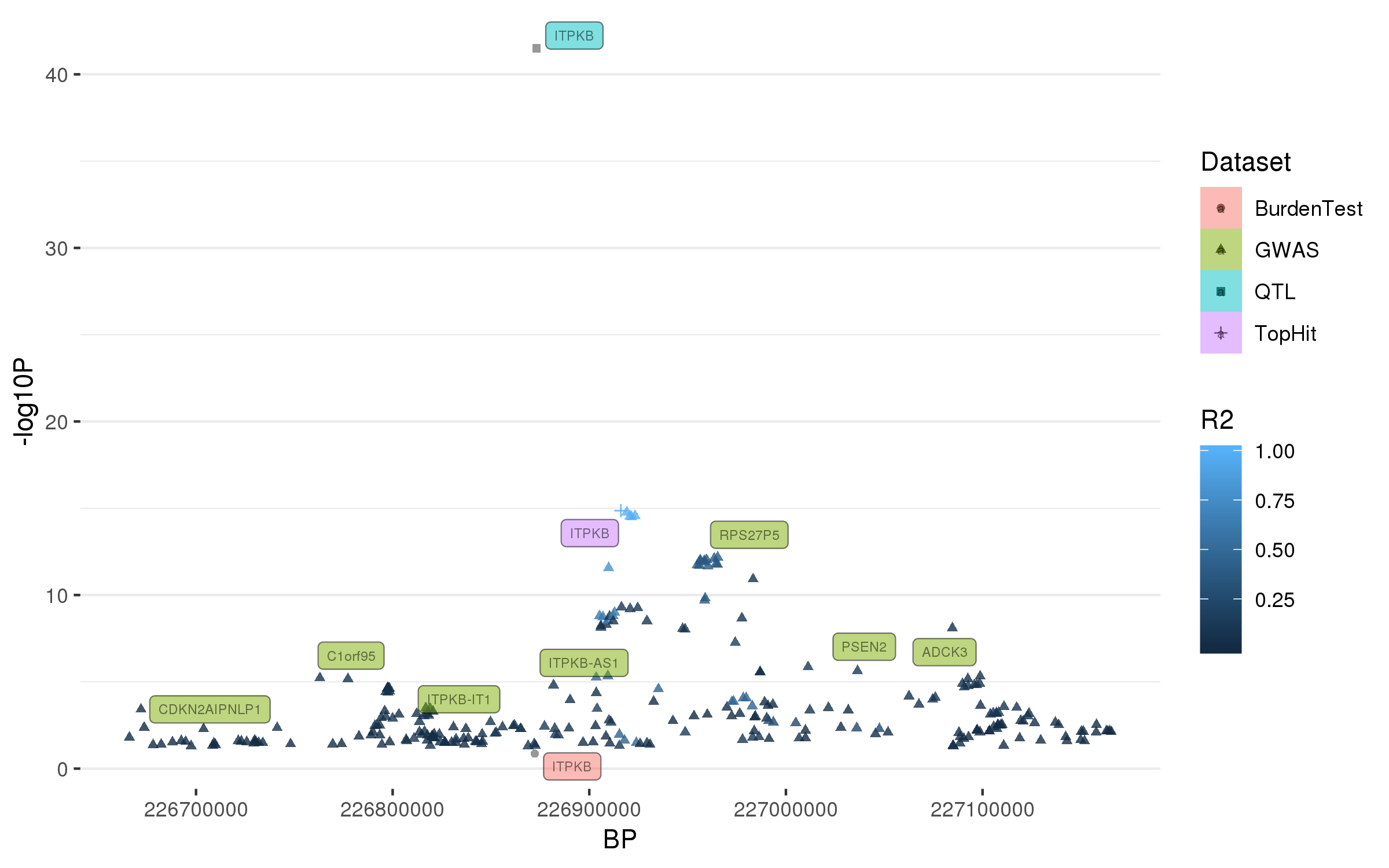

### rs4698412.png

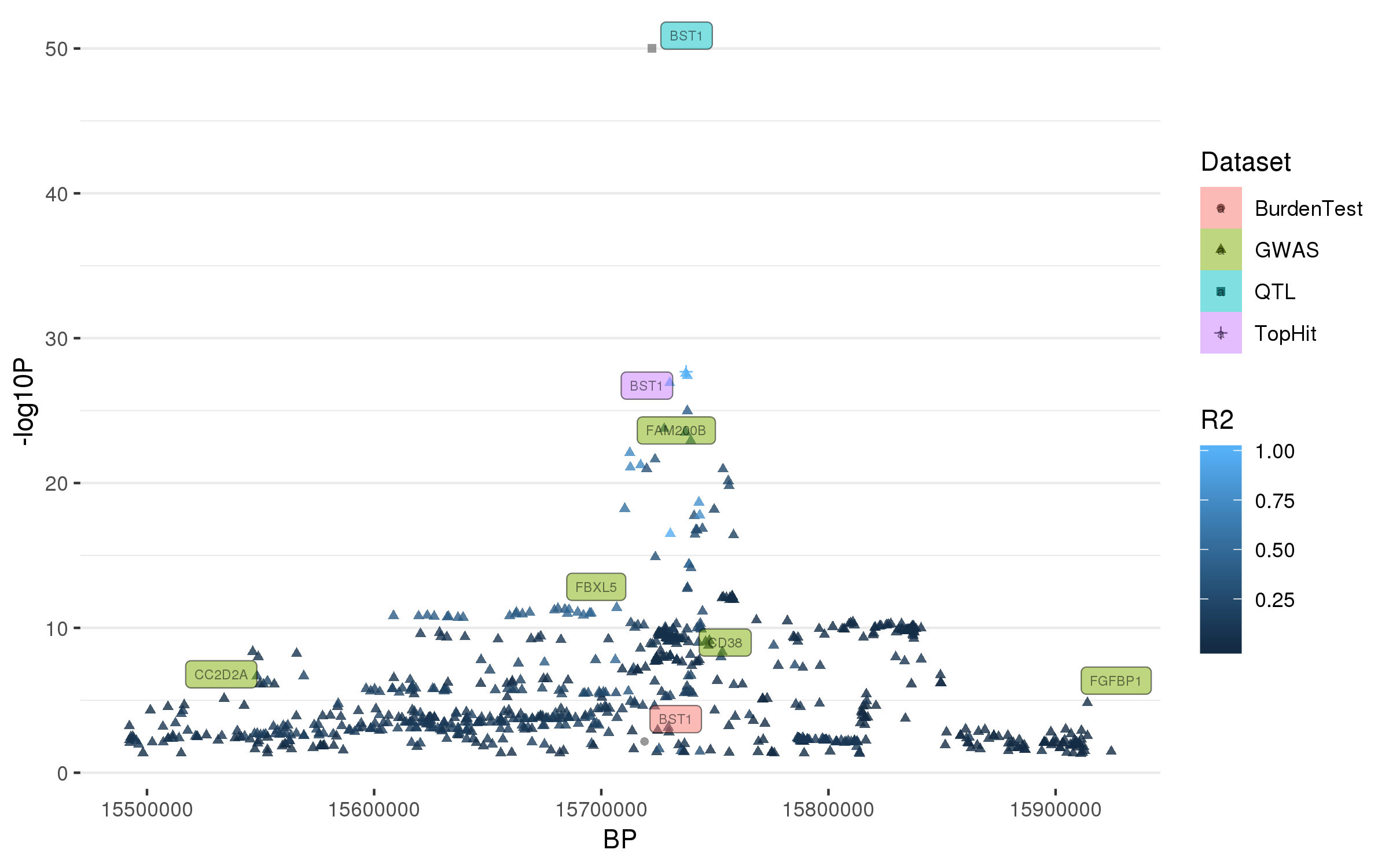

### rs4771268.png

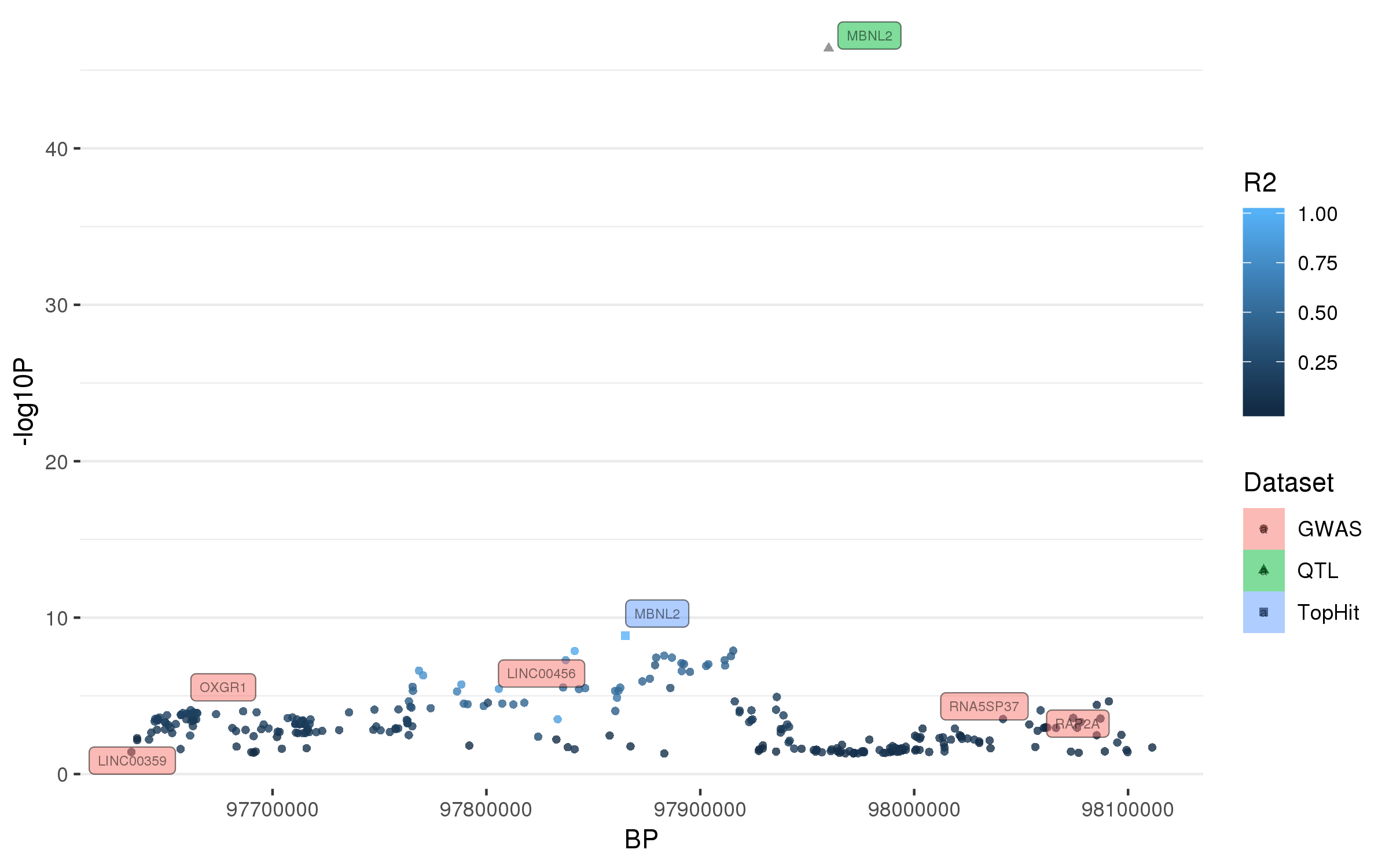

### rs5019538.png

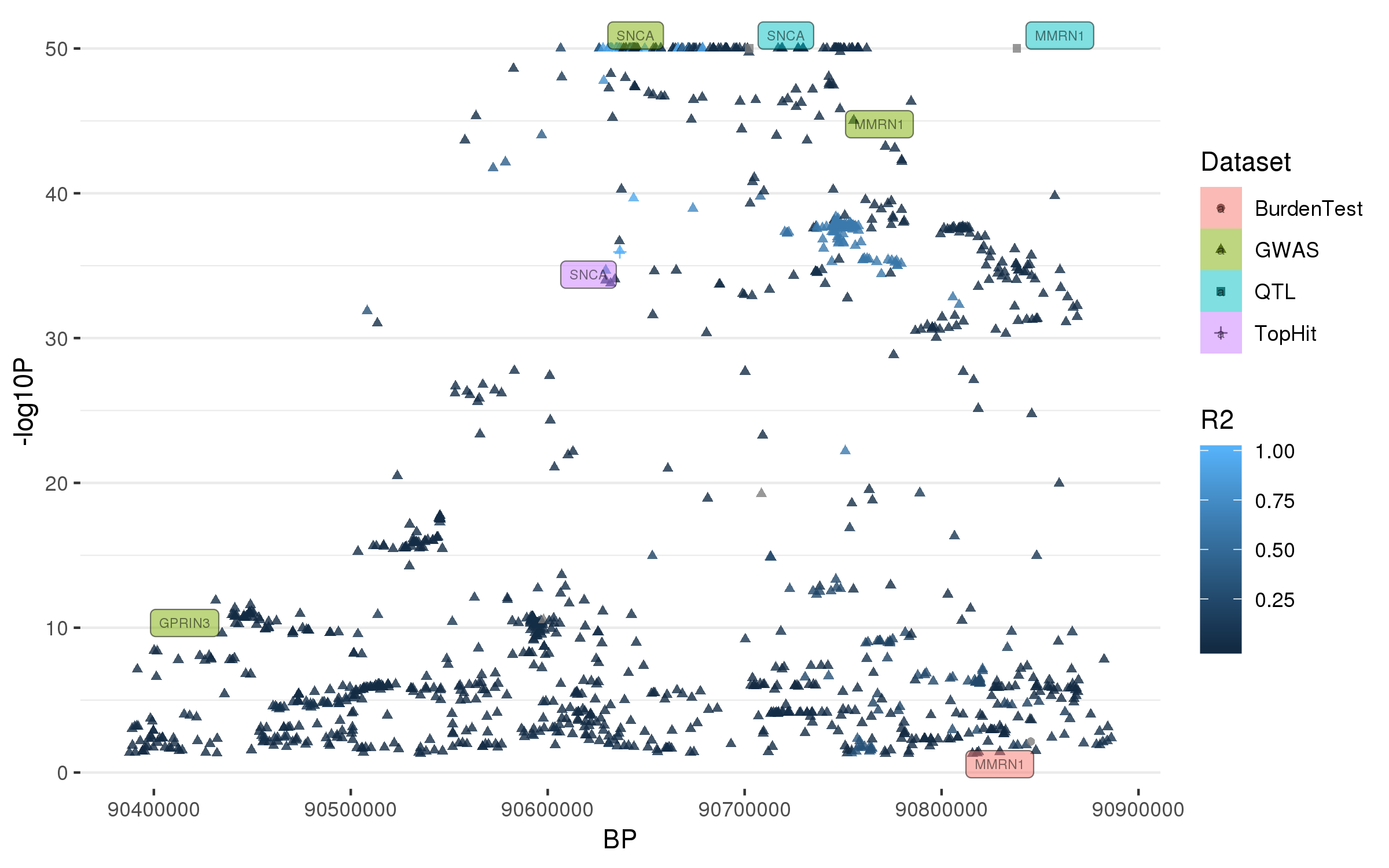

### rs6476434.png

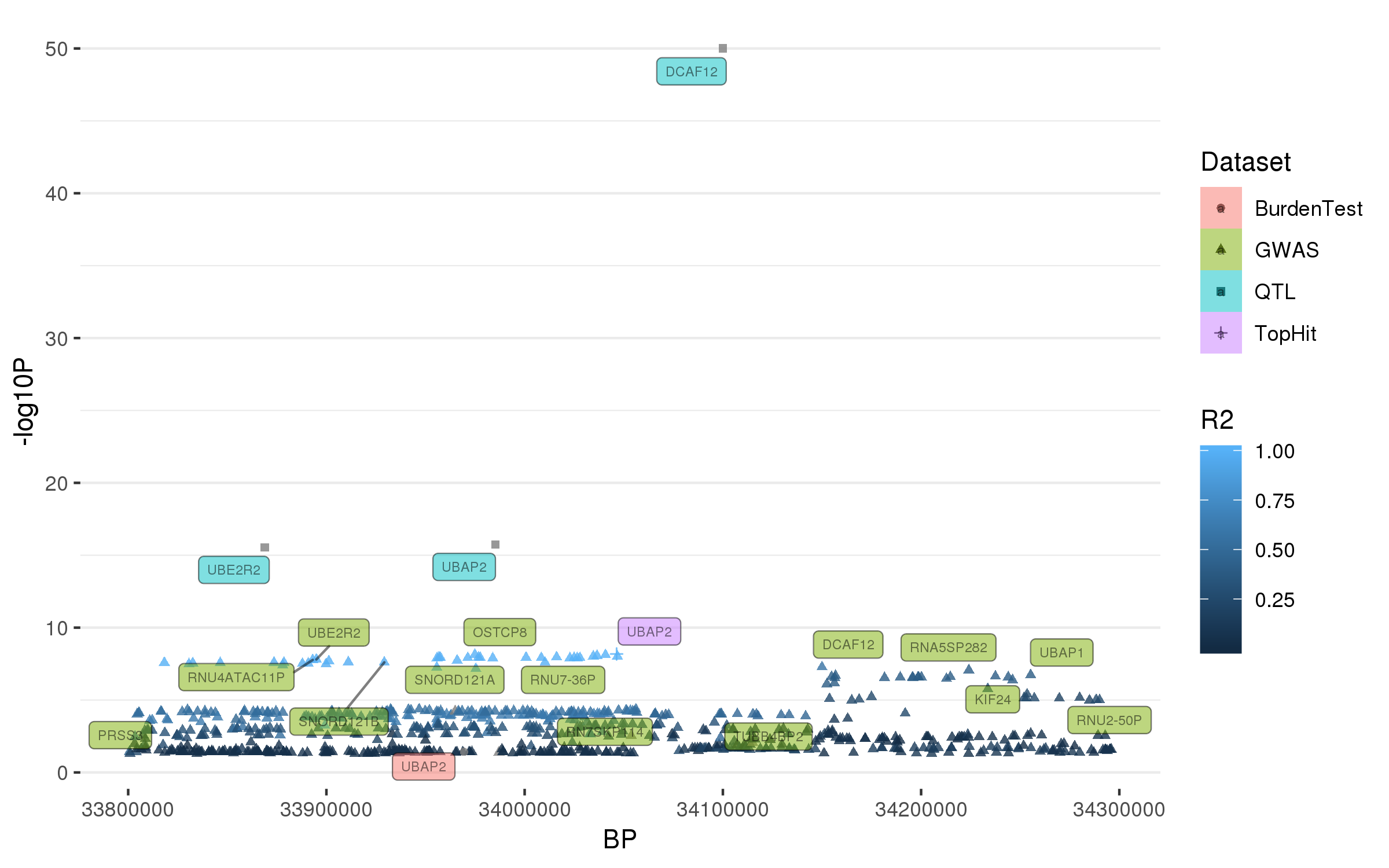

### rs6497339.png

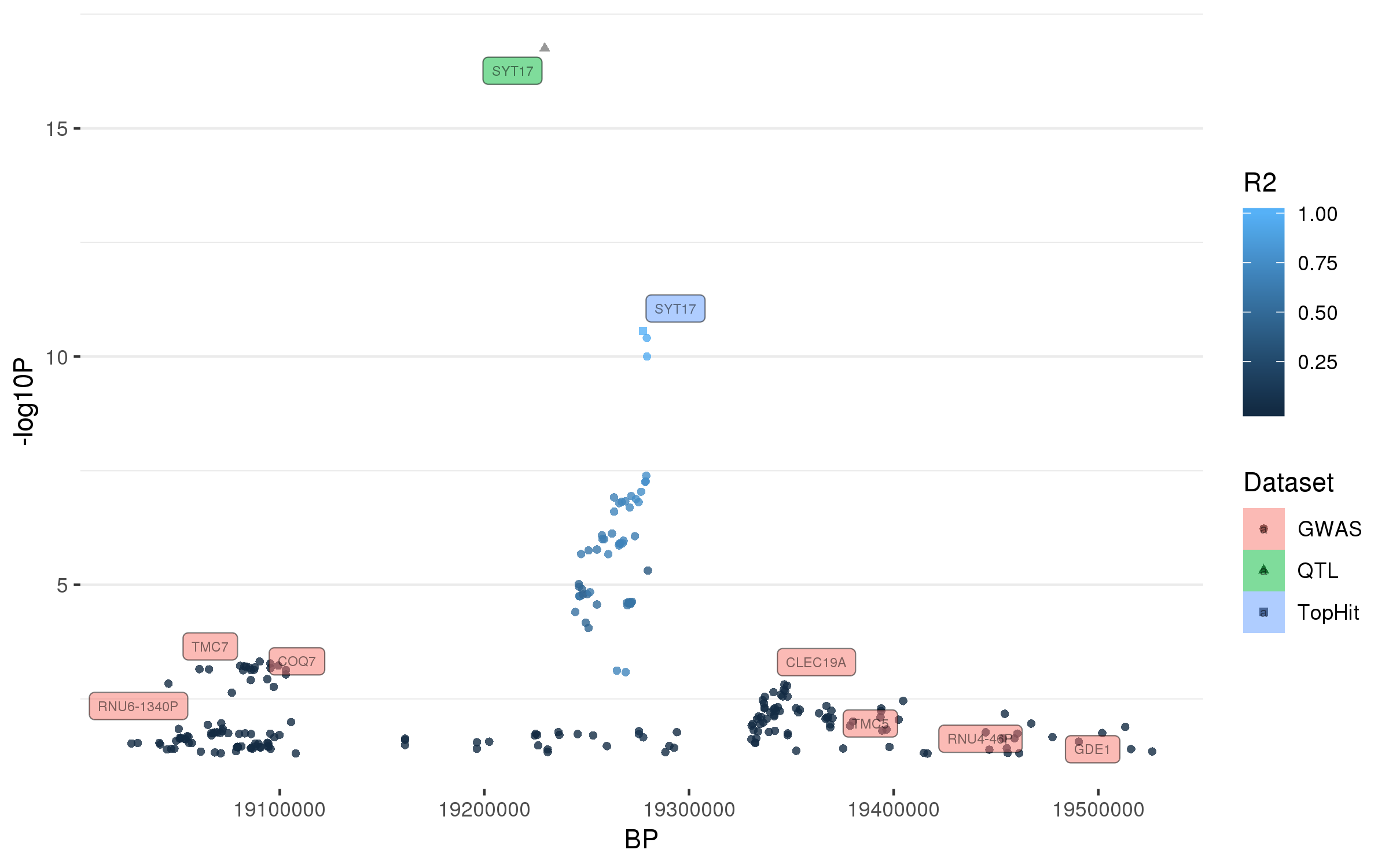

### rs6500328.png

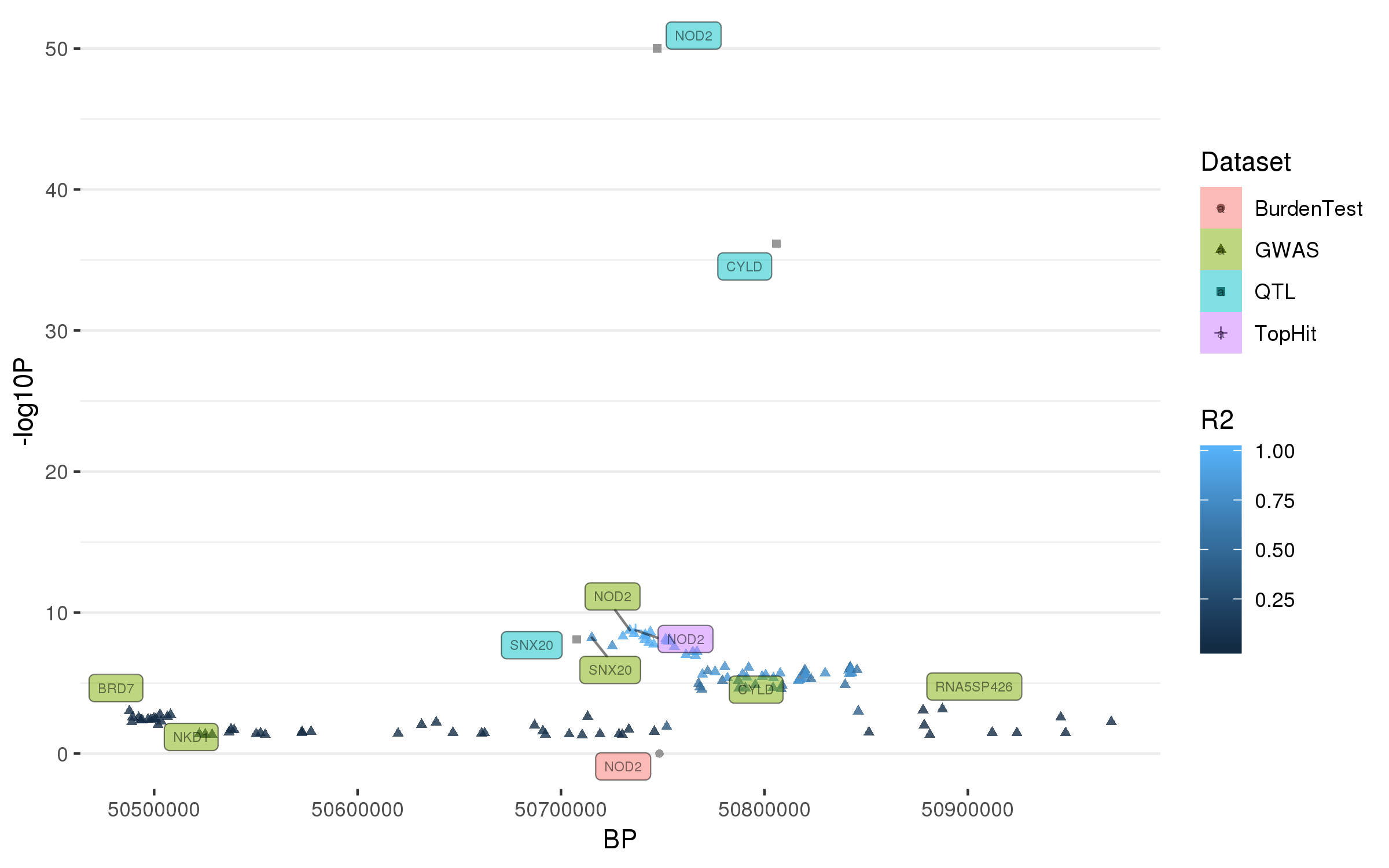

### rs6658353.png

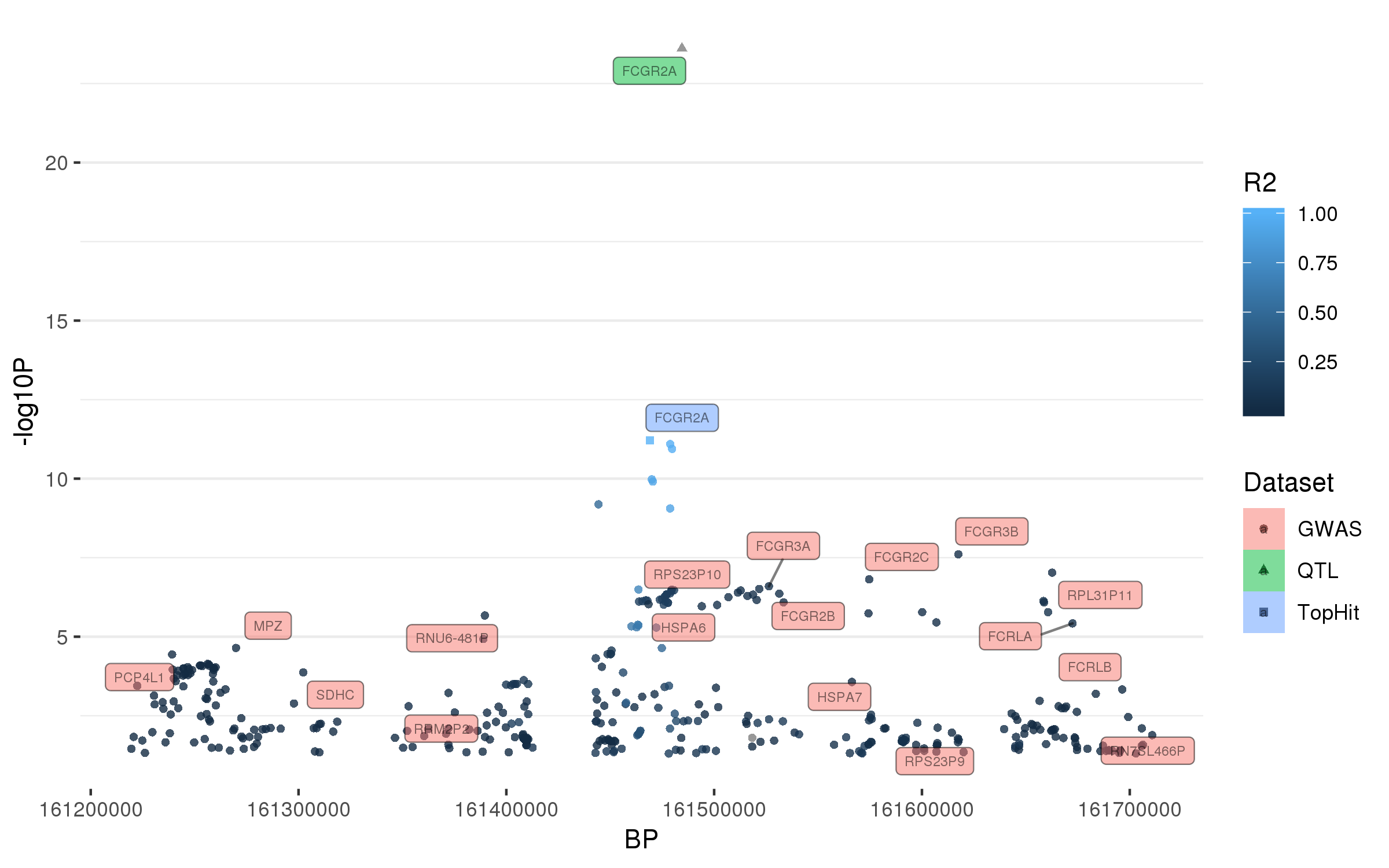

### rs6808178.png

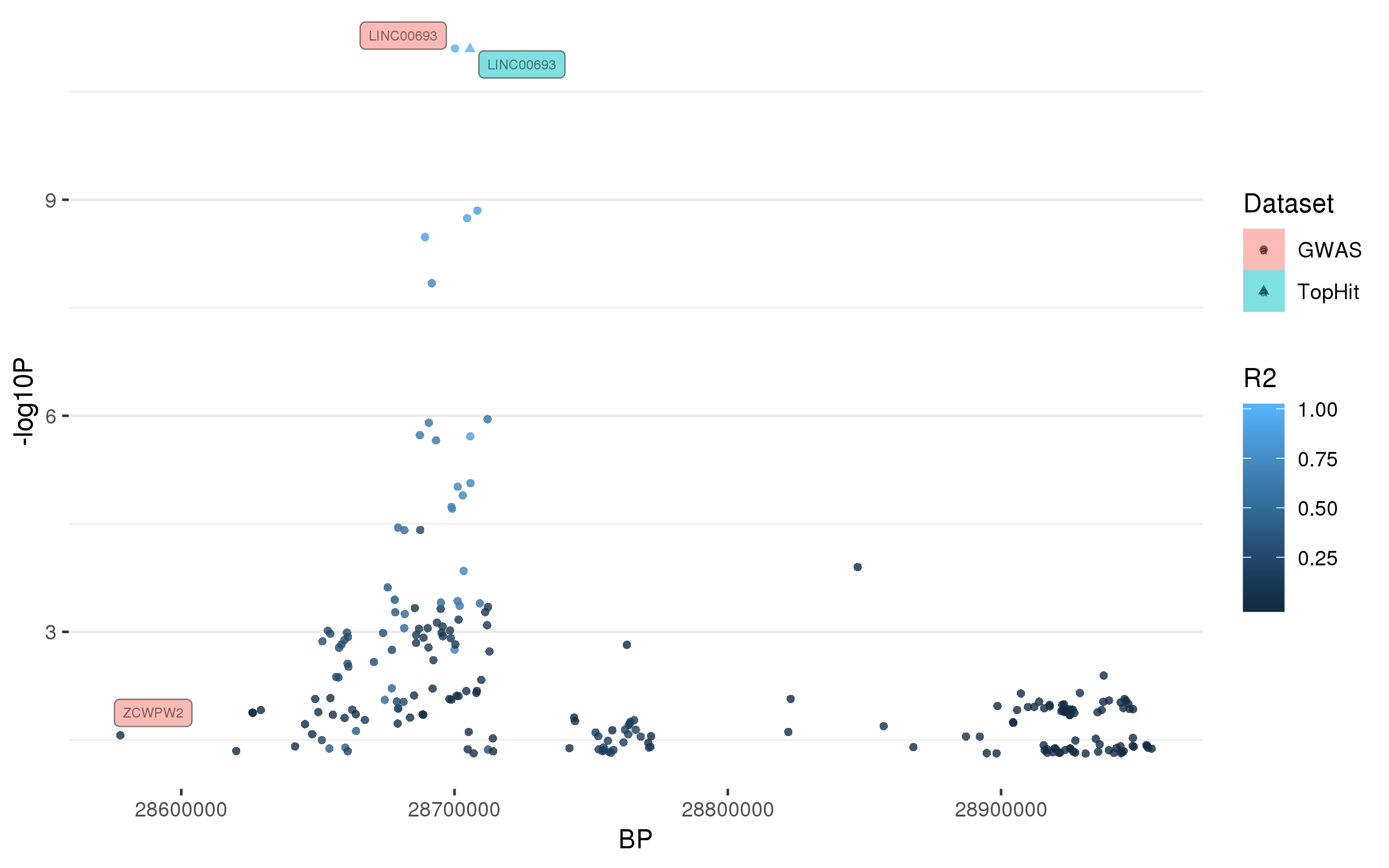

### rs6825004.png

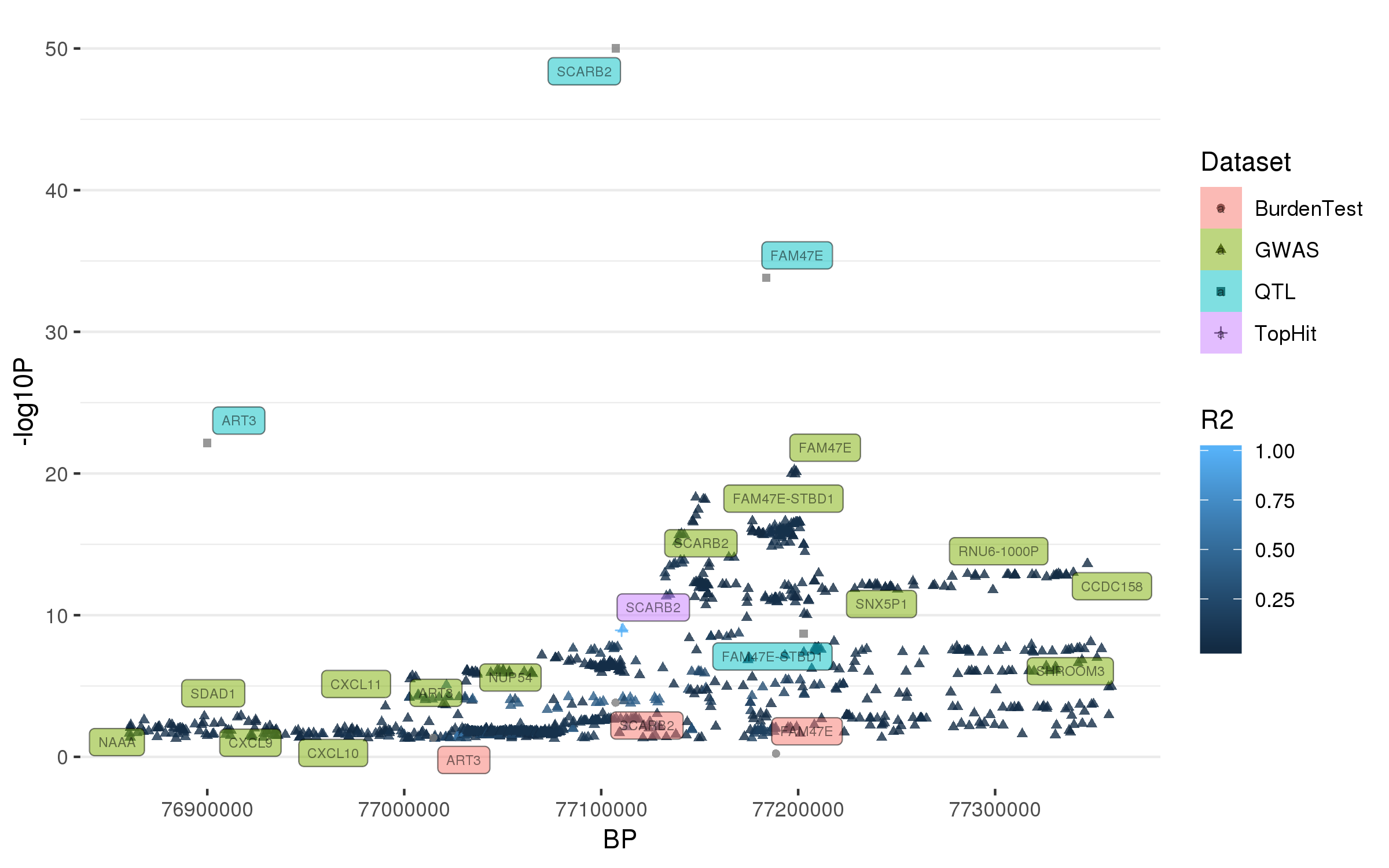

### rs6854006.png

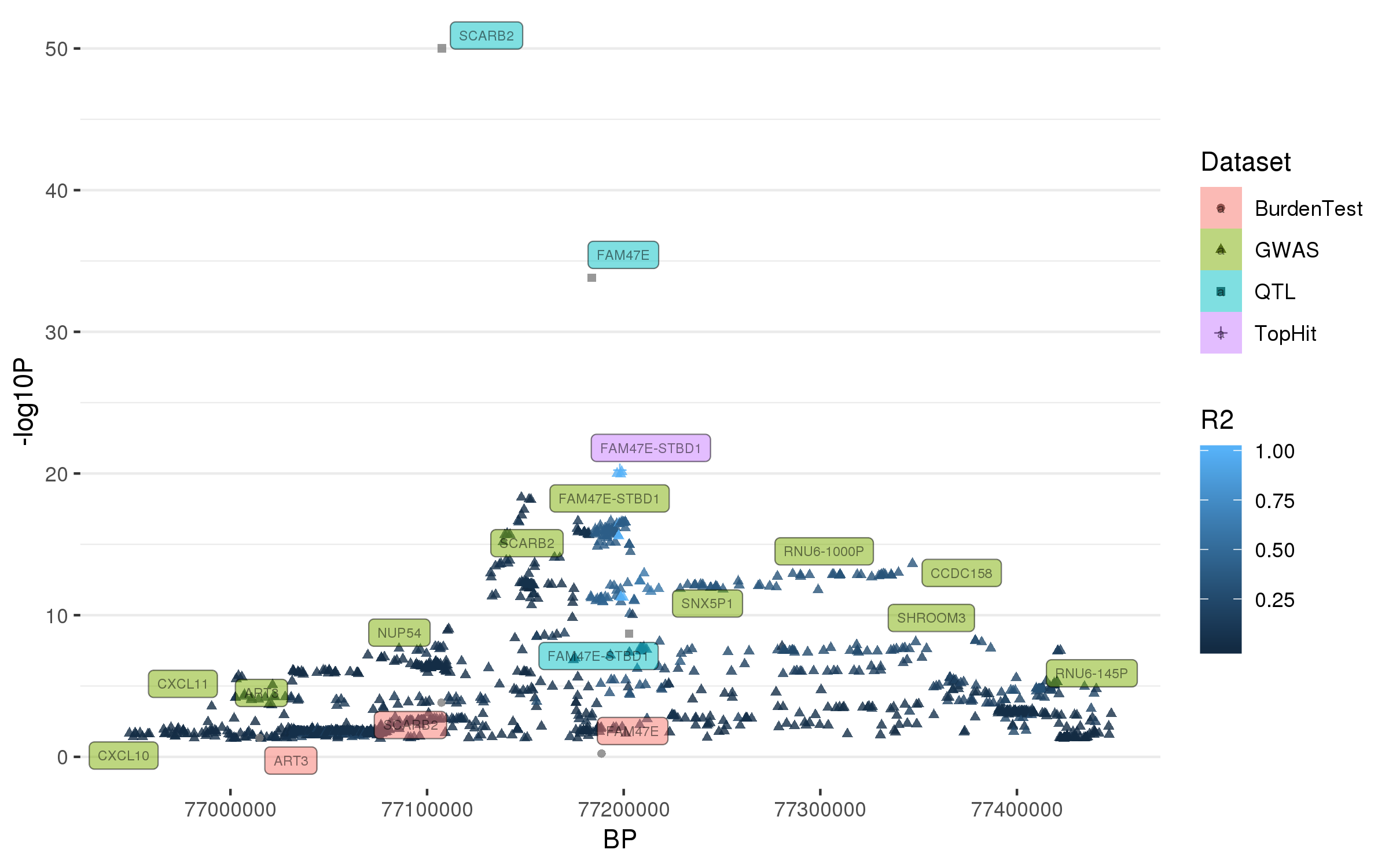

### rs7134559.png

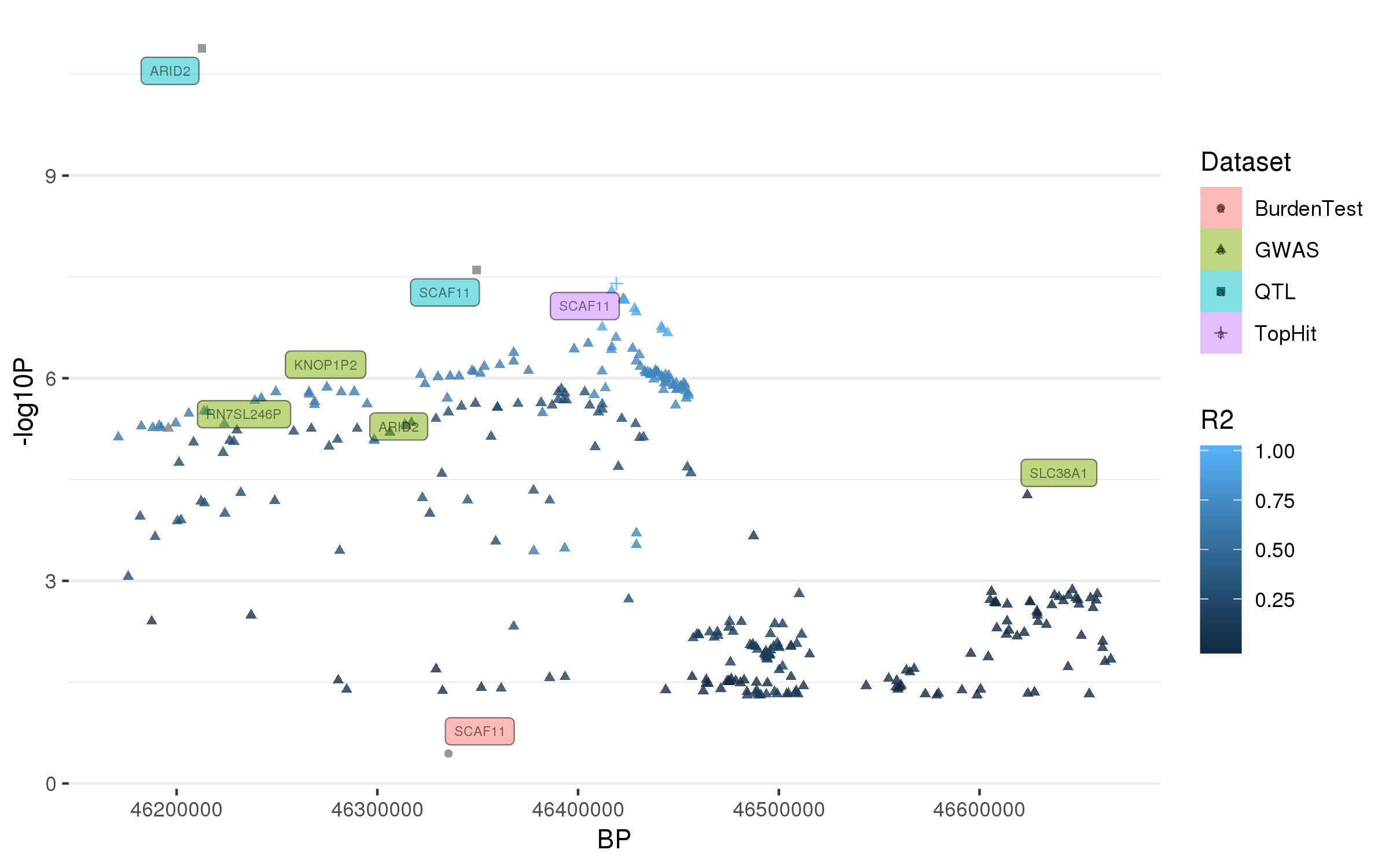

### rs7938782.png

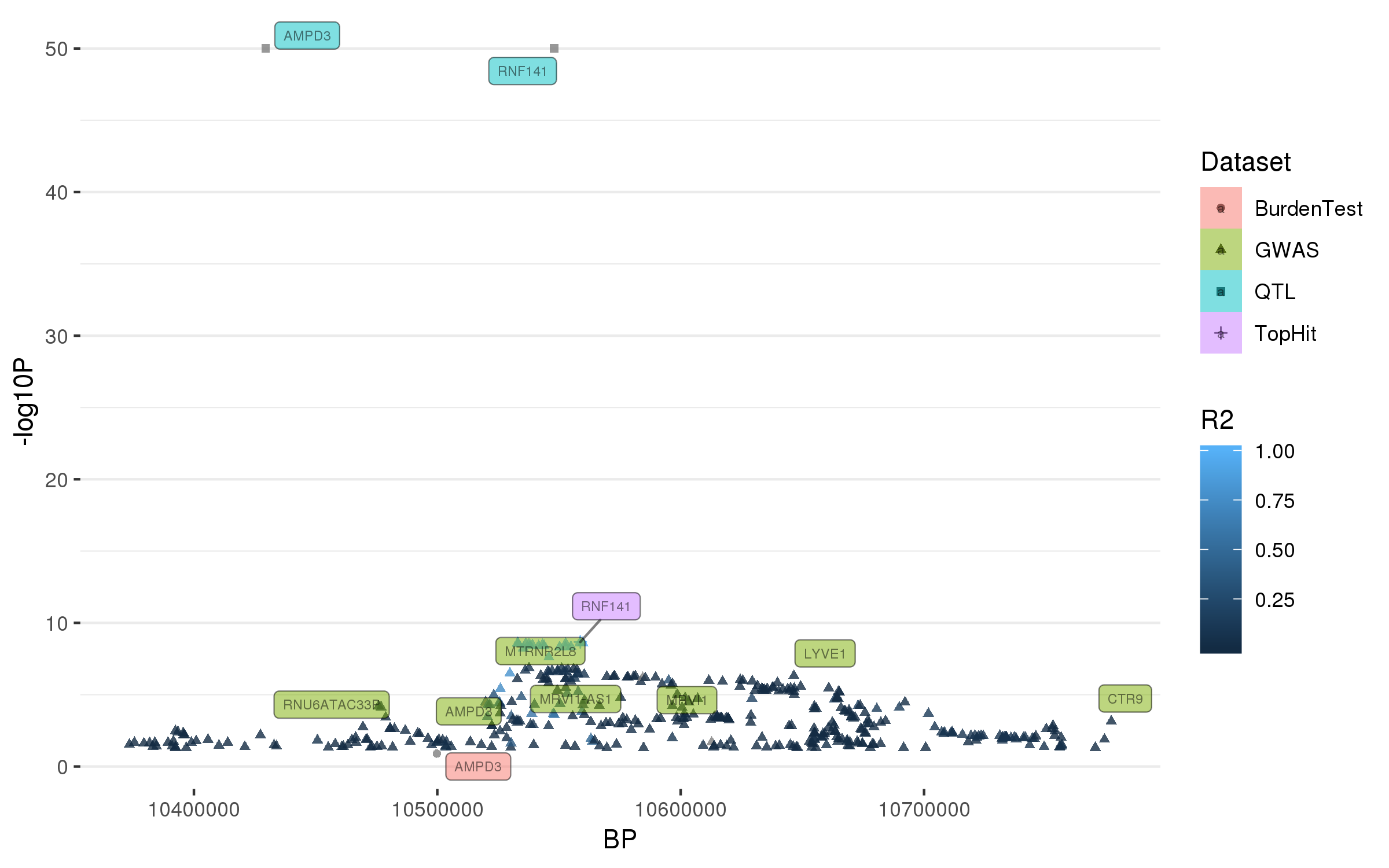

### rs8087969.png

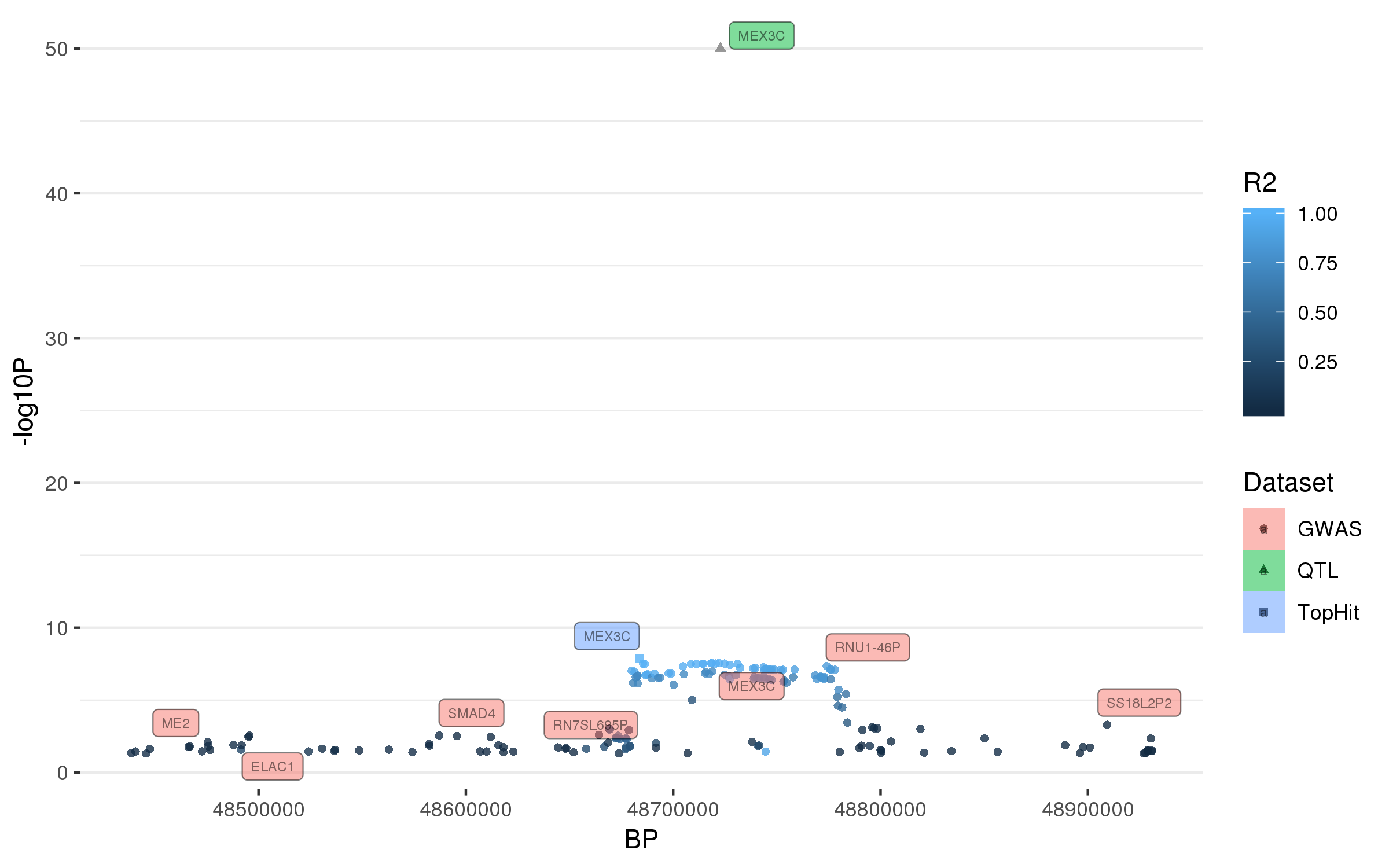

### rs9261484.png

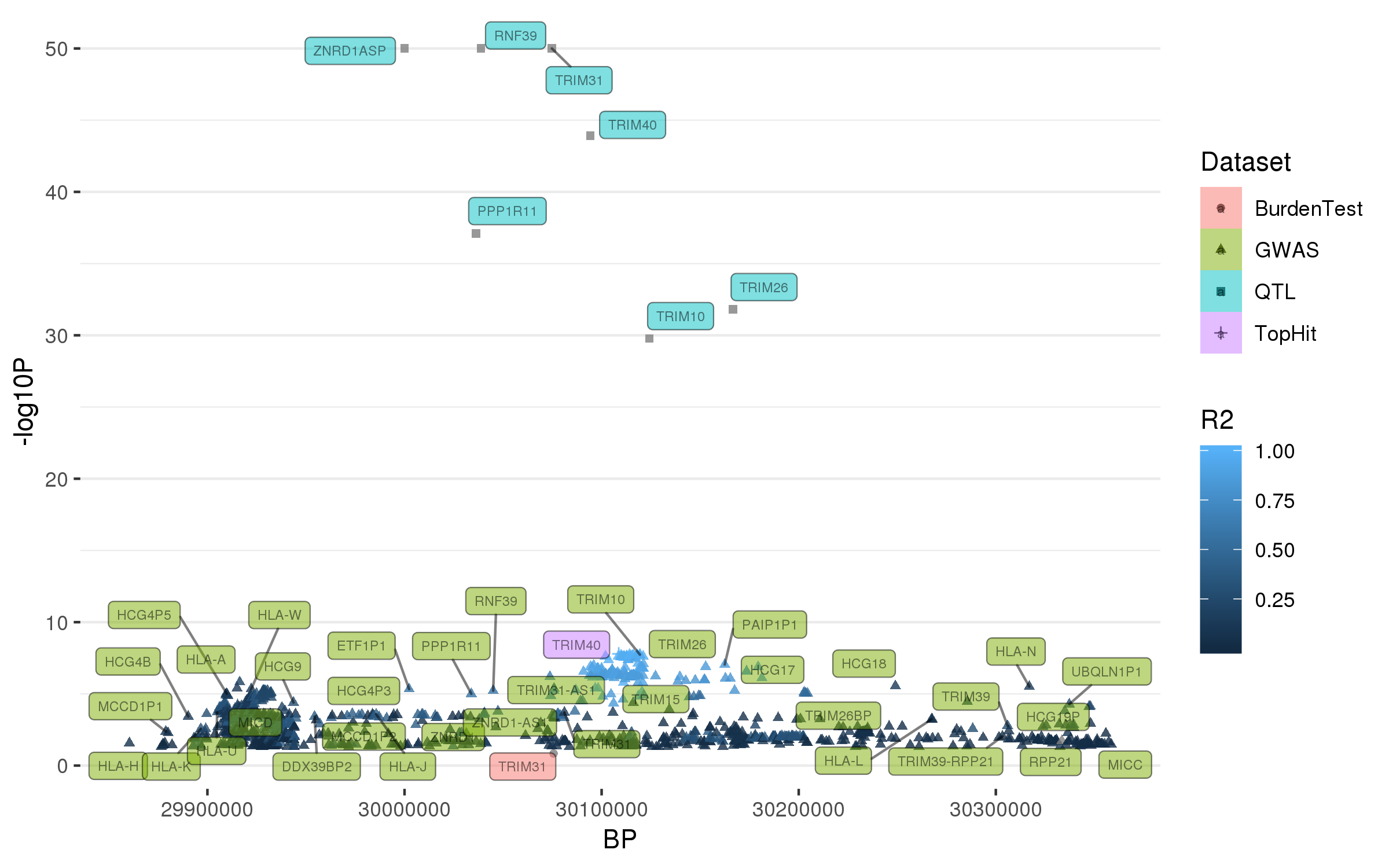

### rs9568188.png

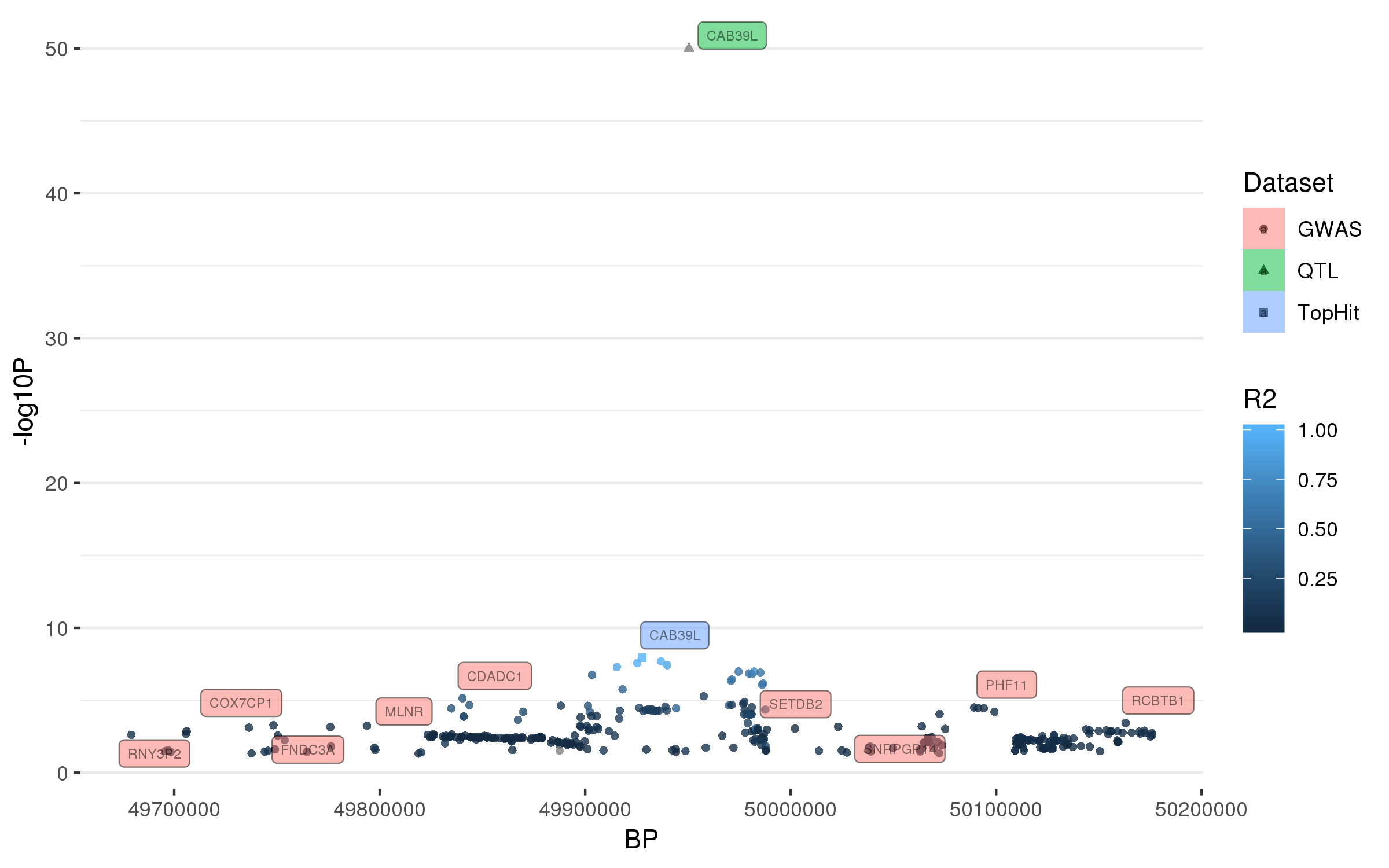

### rs10221156.png

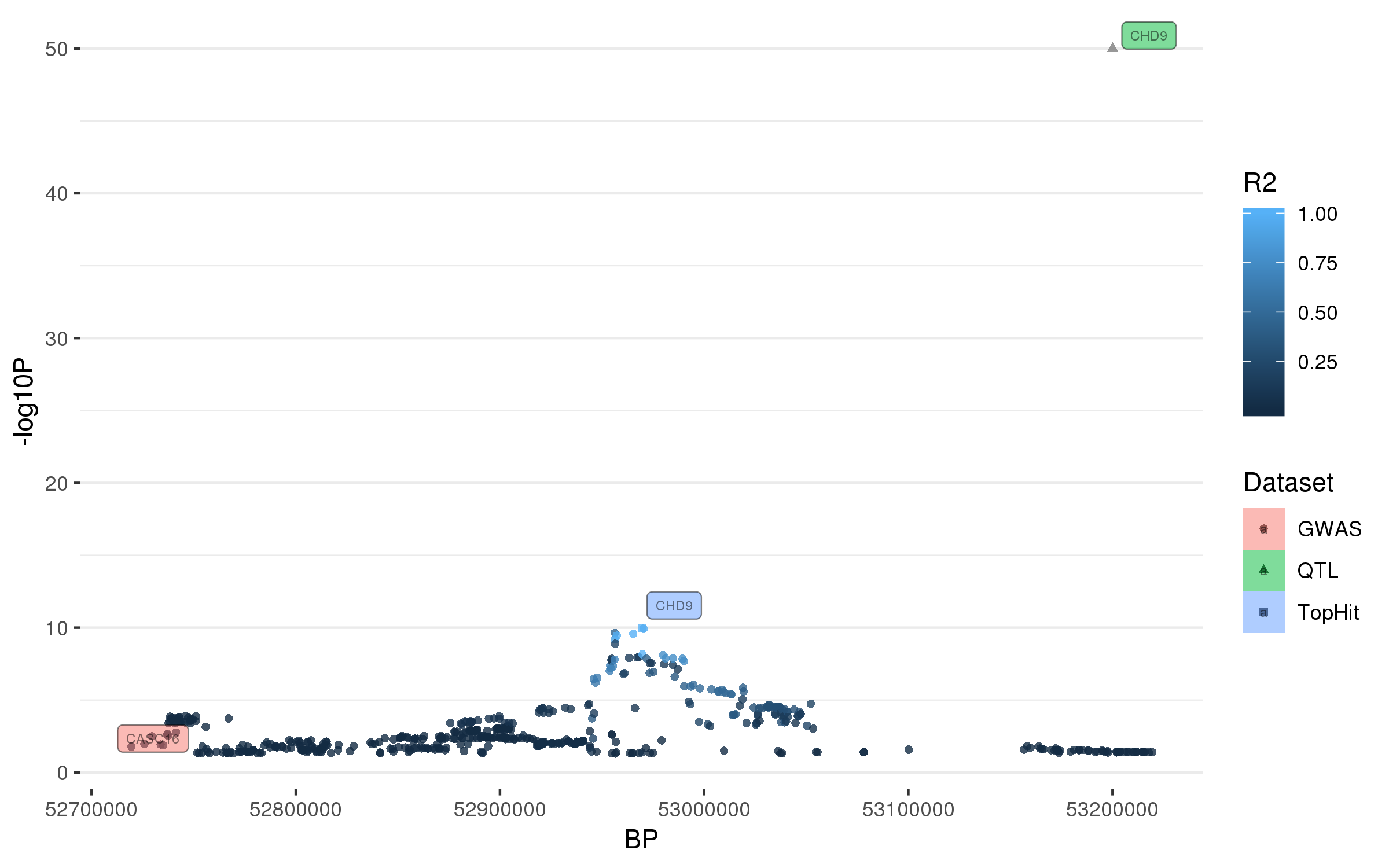

### rs10513789.png

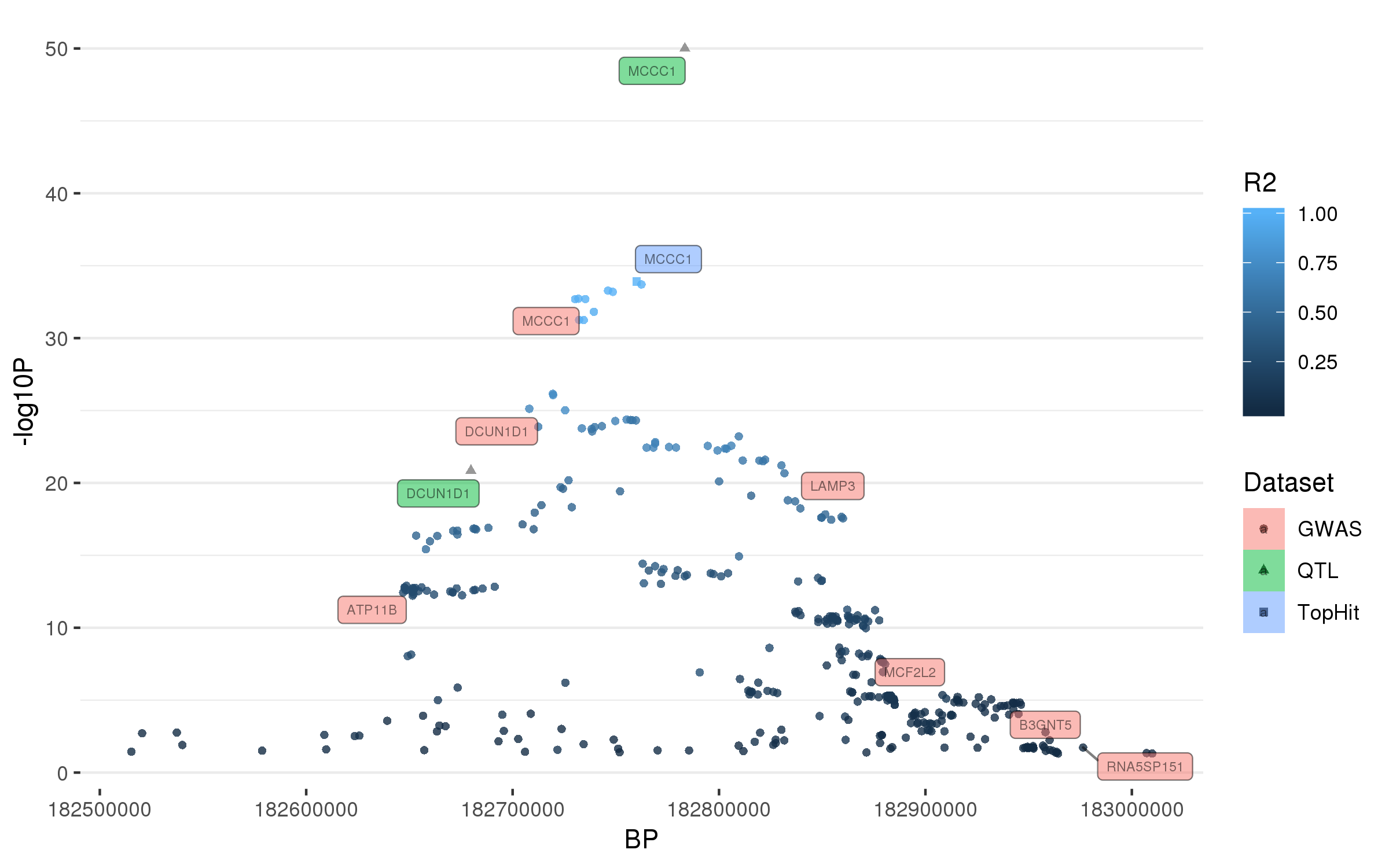

### rs10748818.png

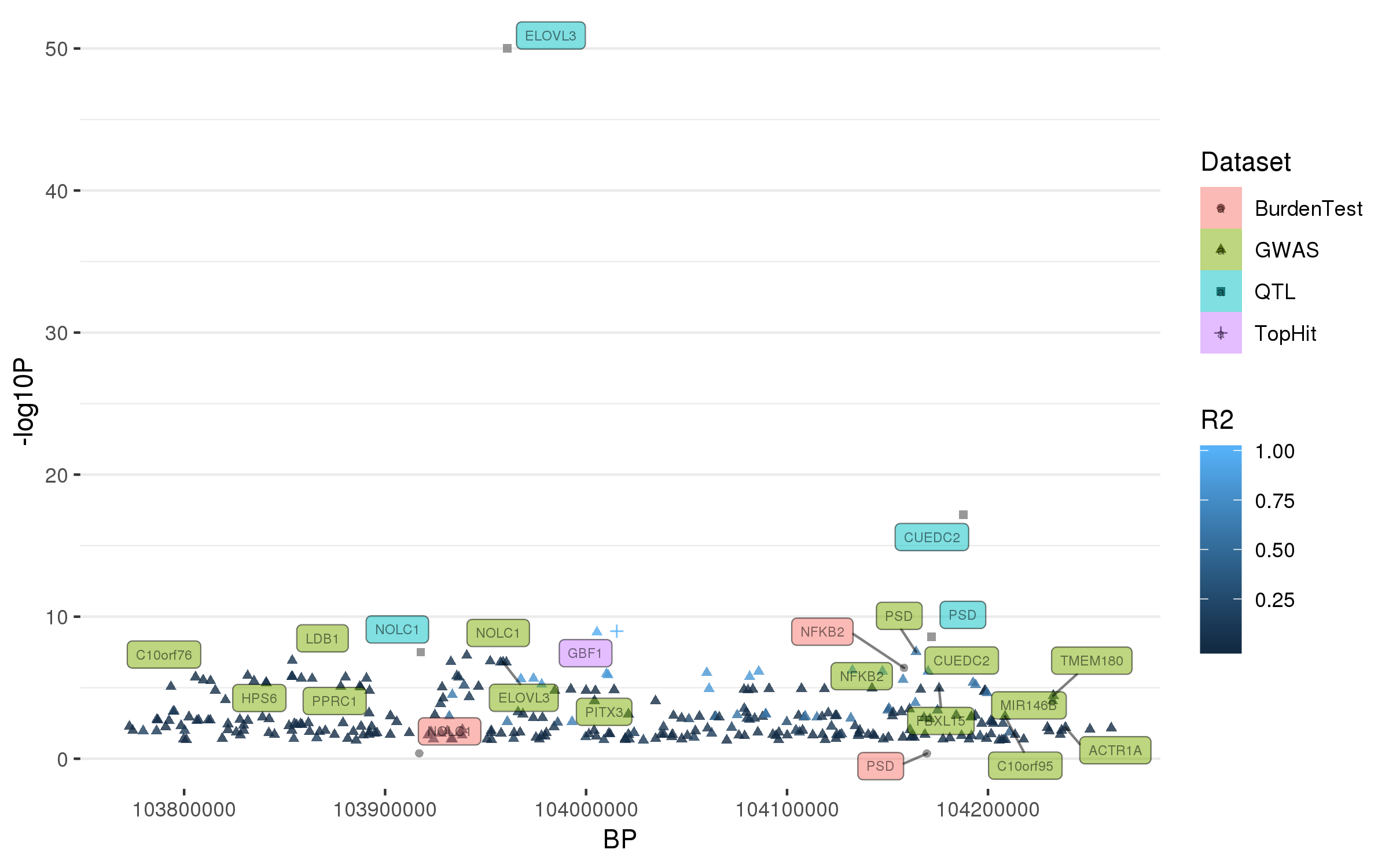

### rs10756907.png

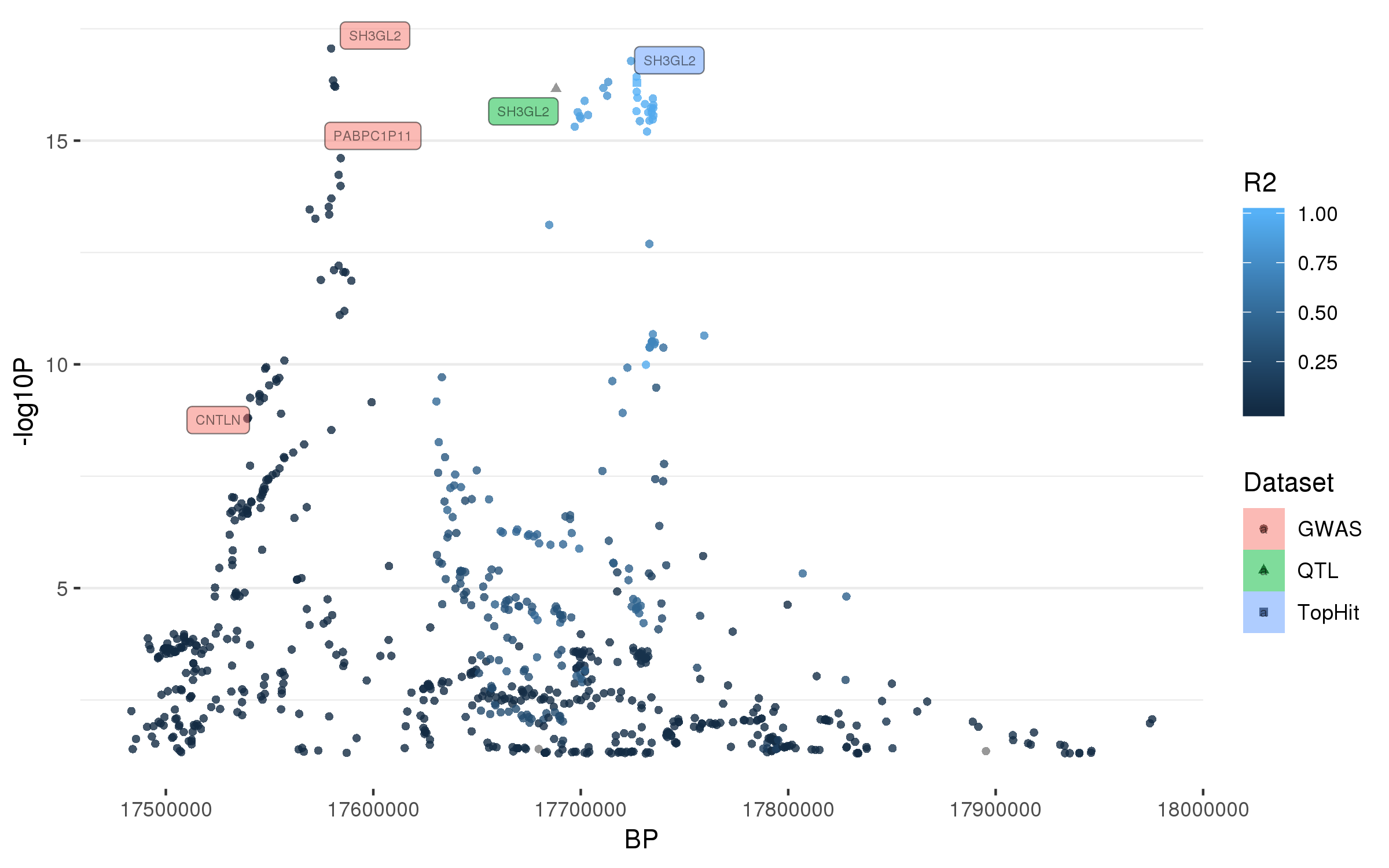

### rs10797576.png

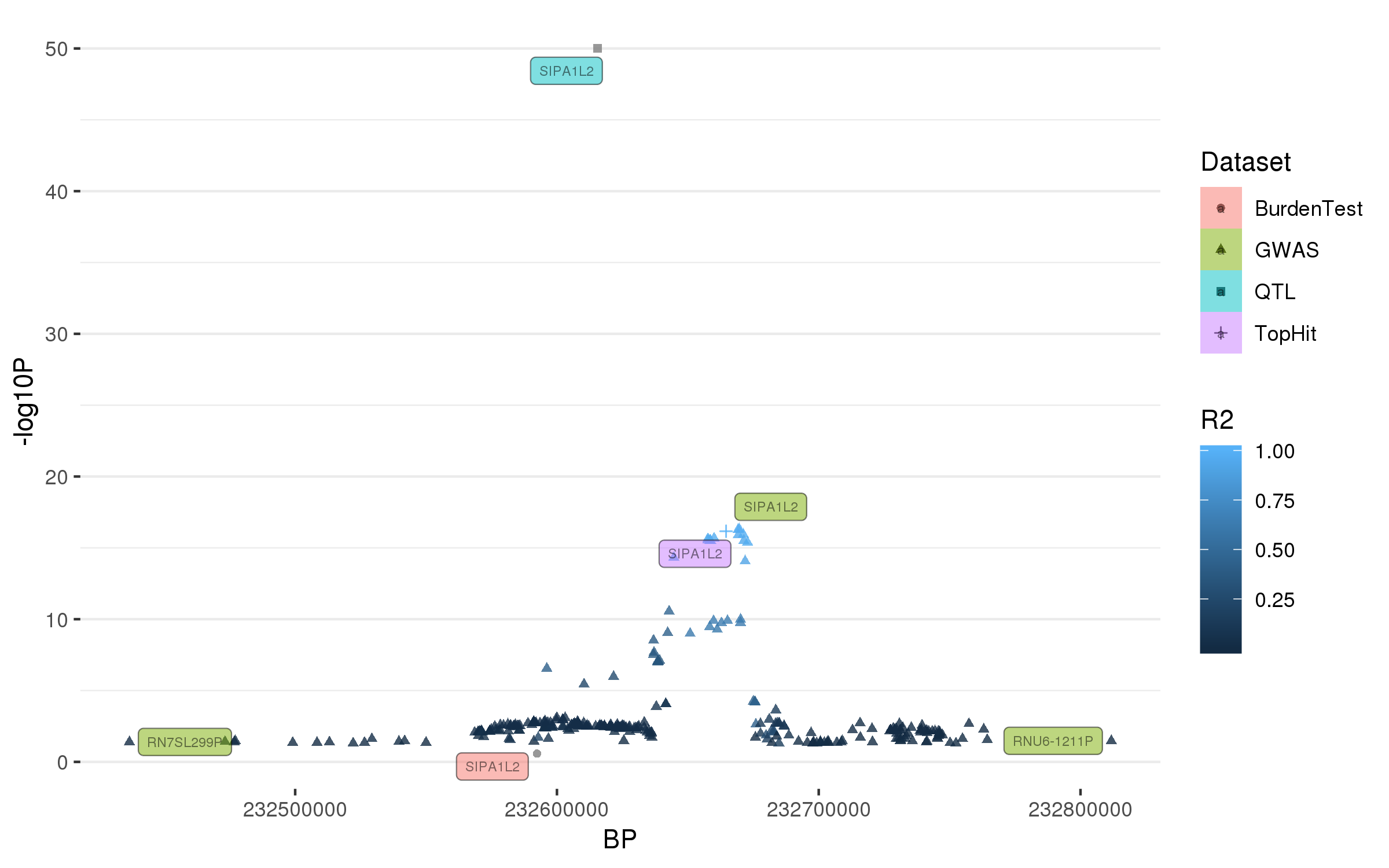

### rs10847864.png

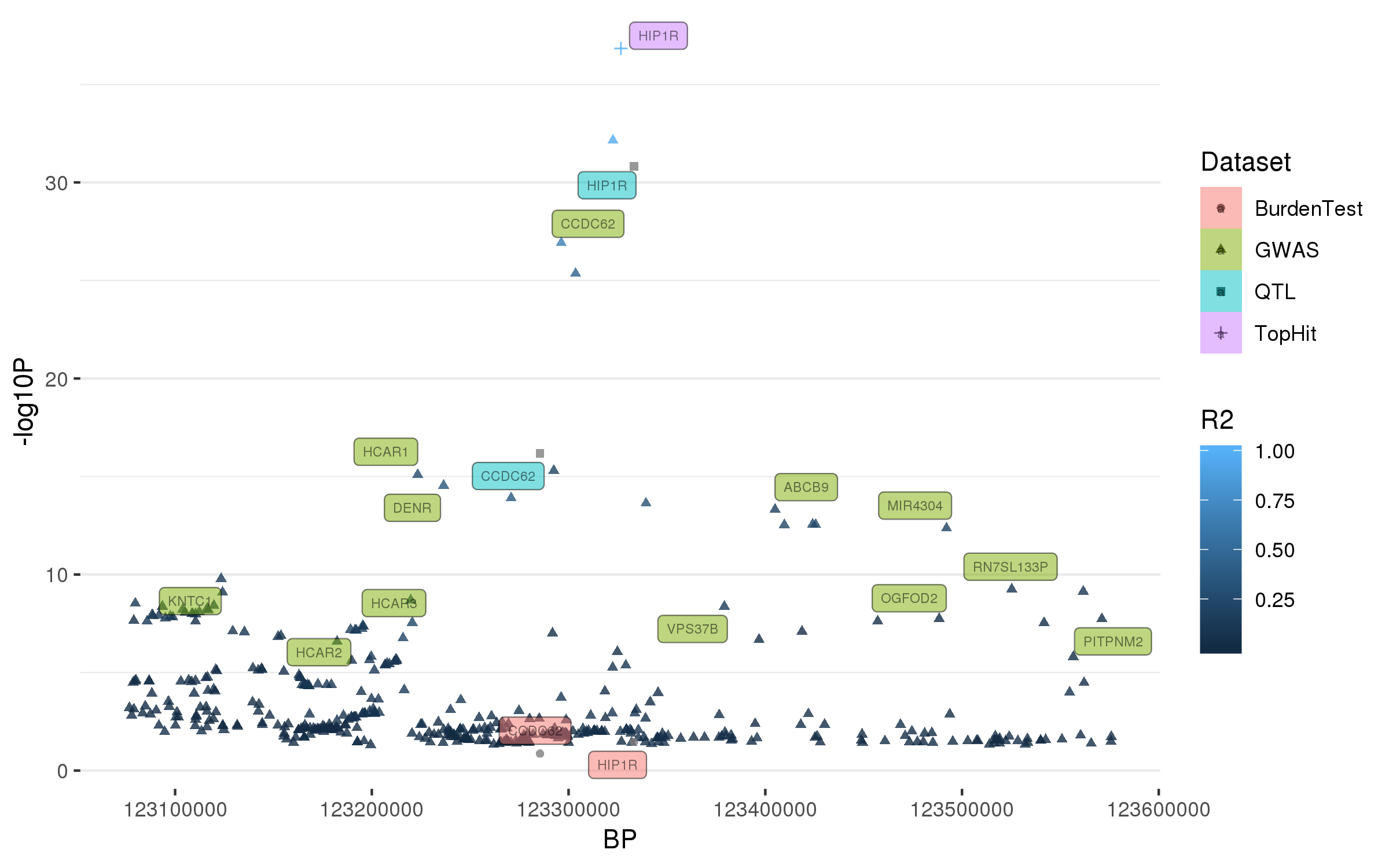

### rs11150601.png

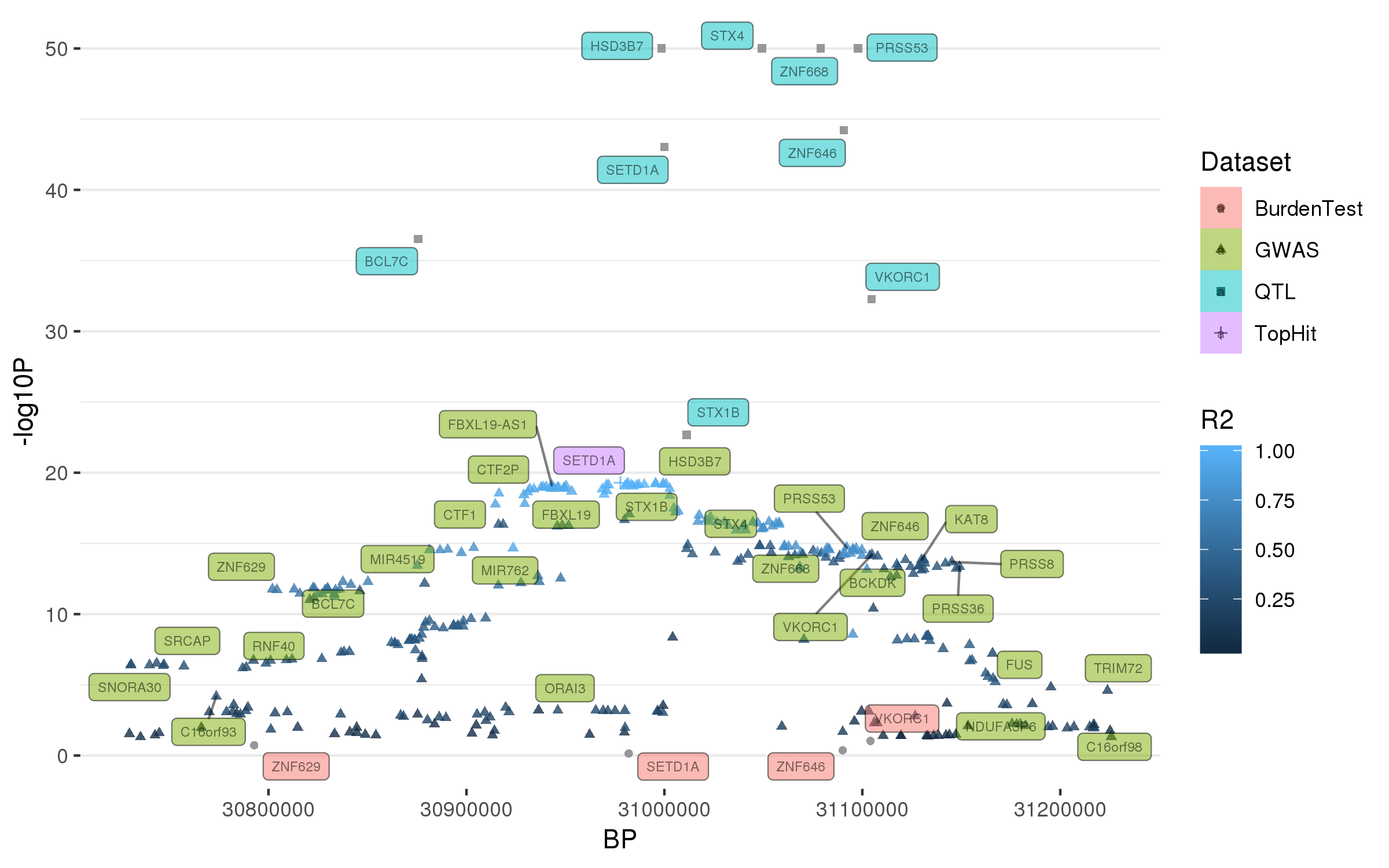

### rs11158026.png

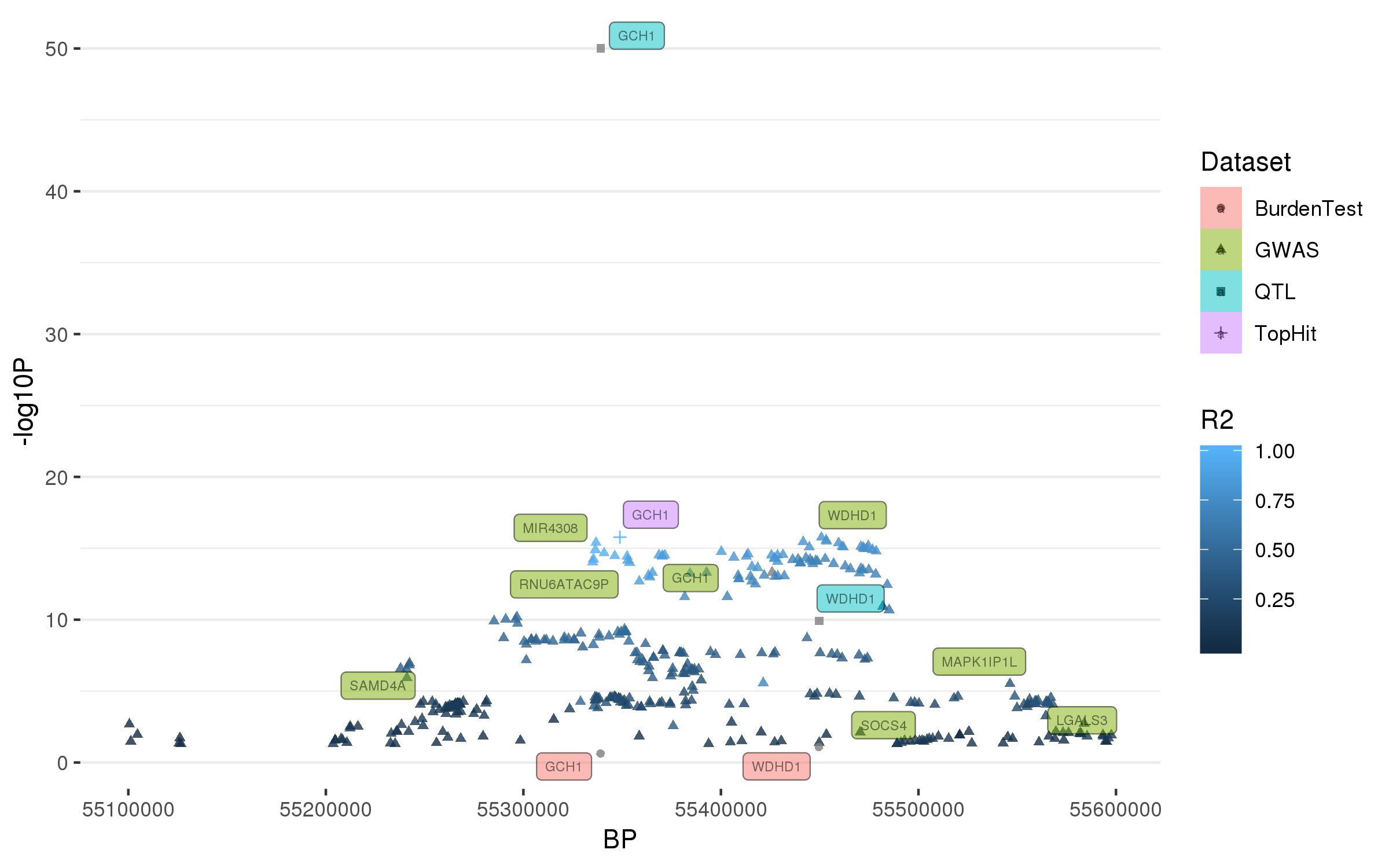

### rs11557080.png

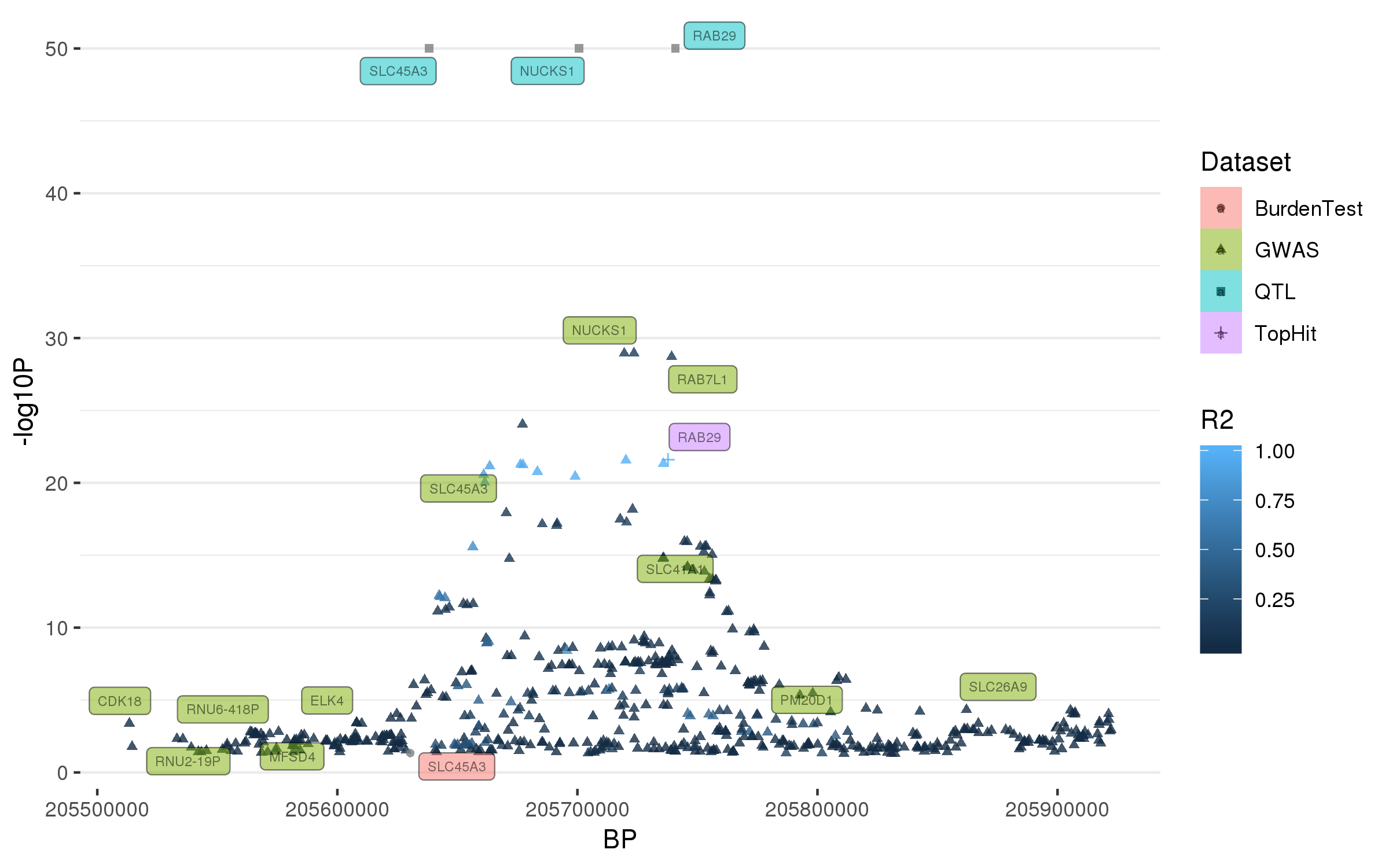

### rs11578699.png

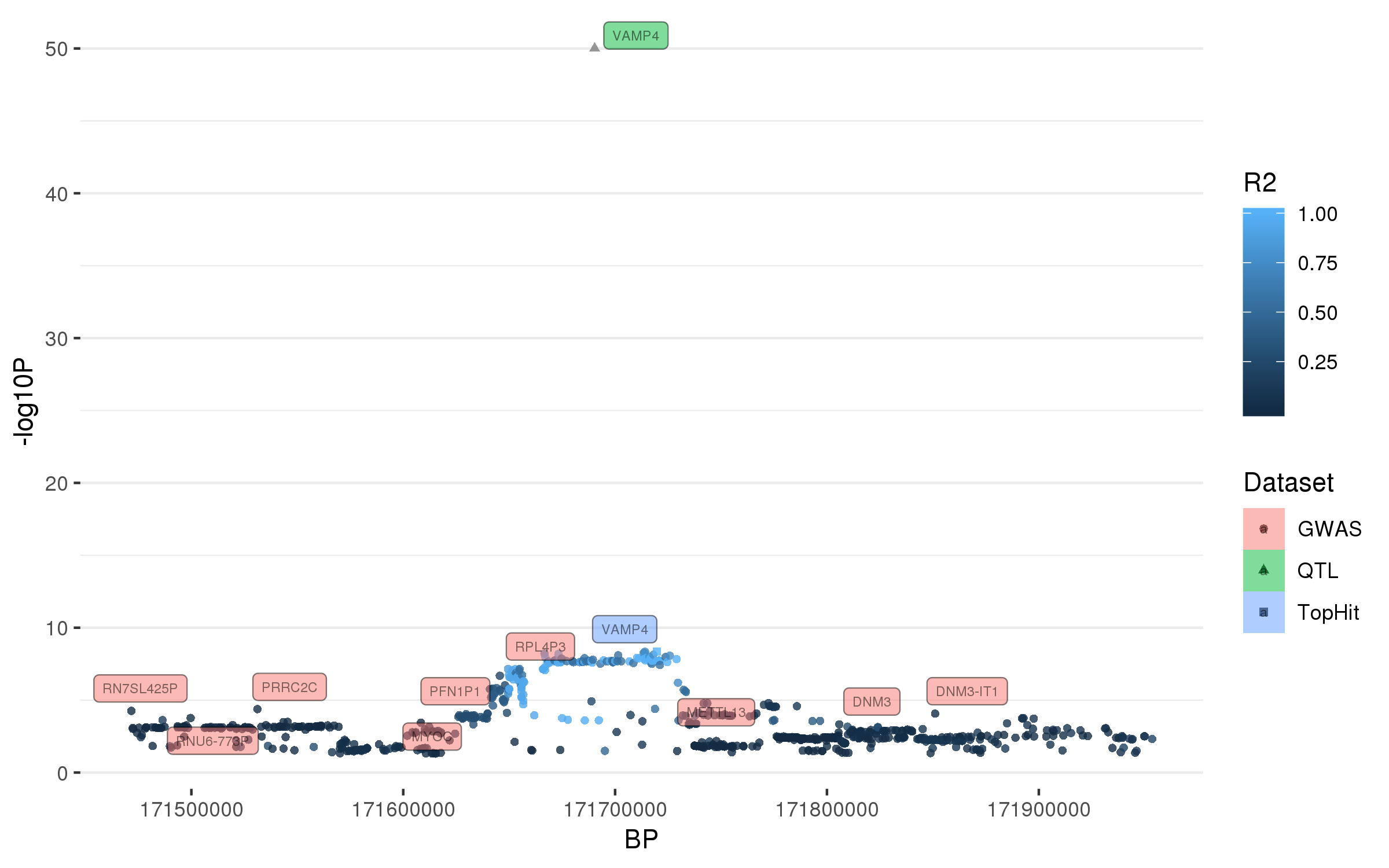

### rs11610045.png

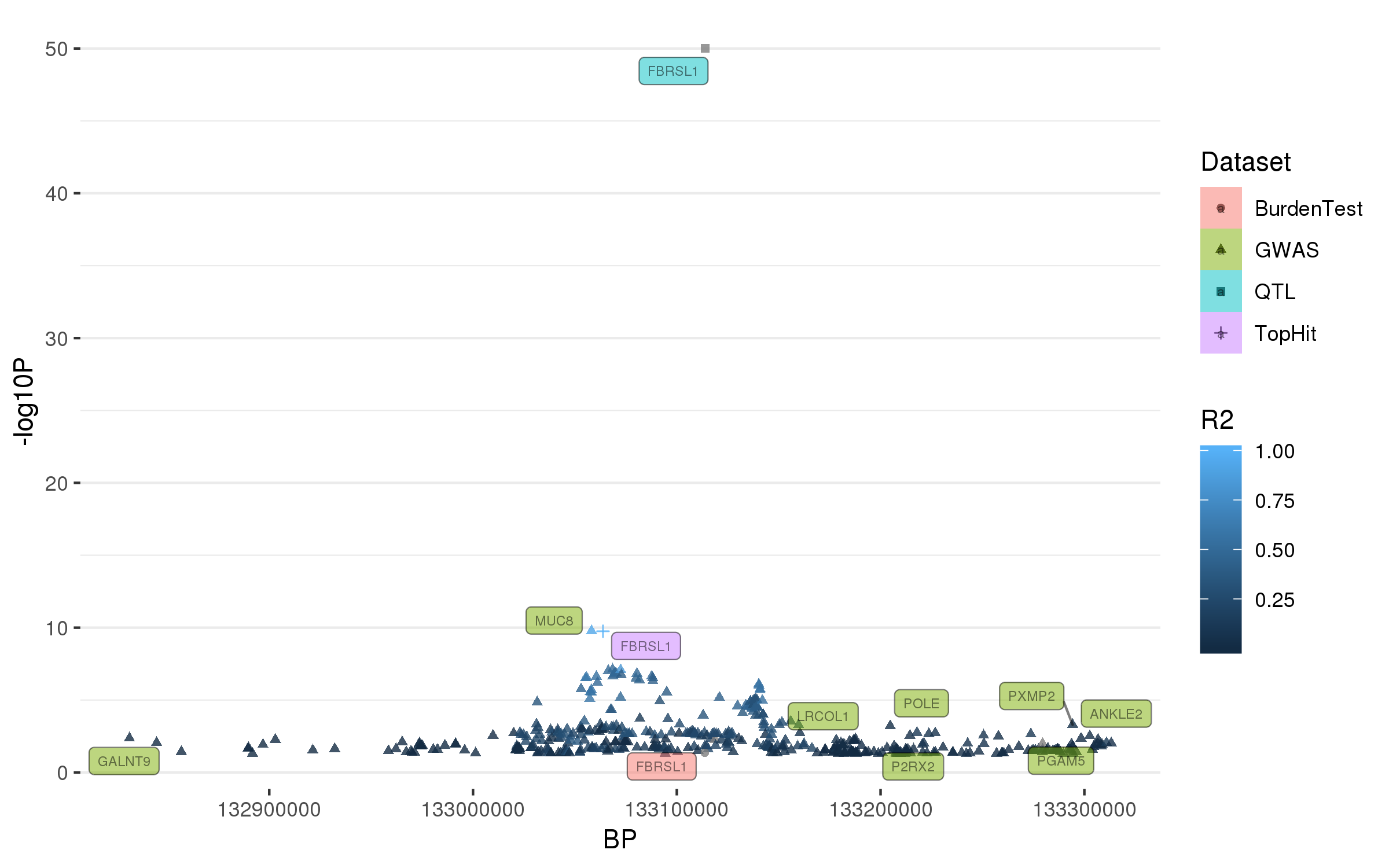

### rs11658976.png

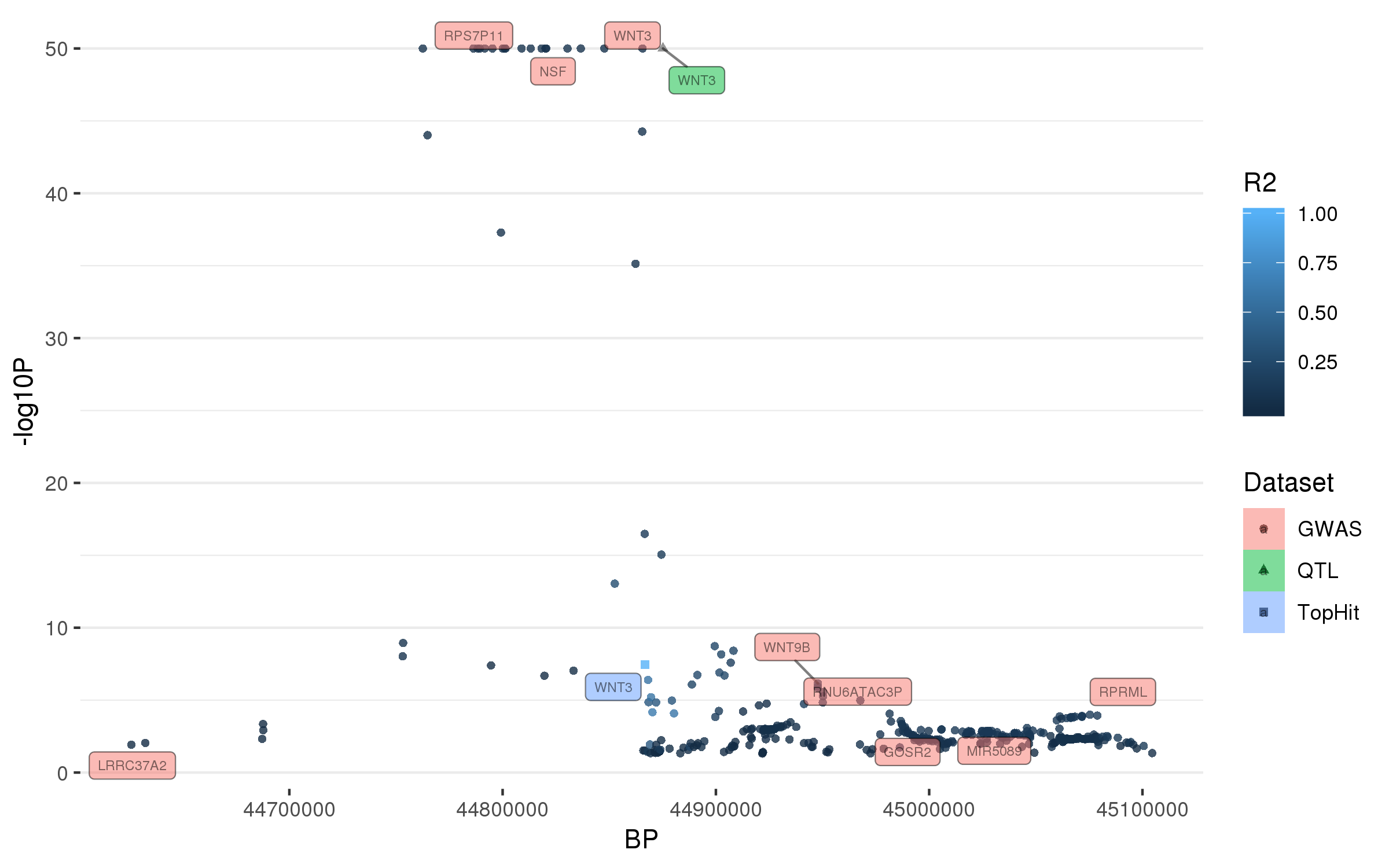

### rs11683001.png

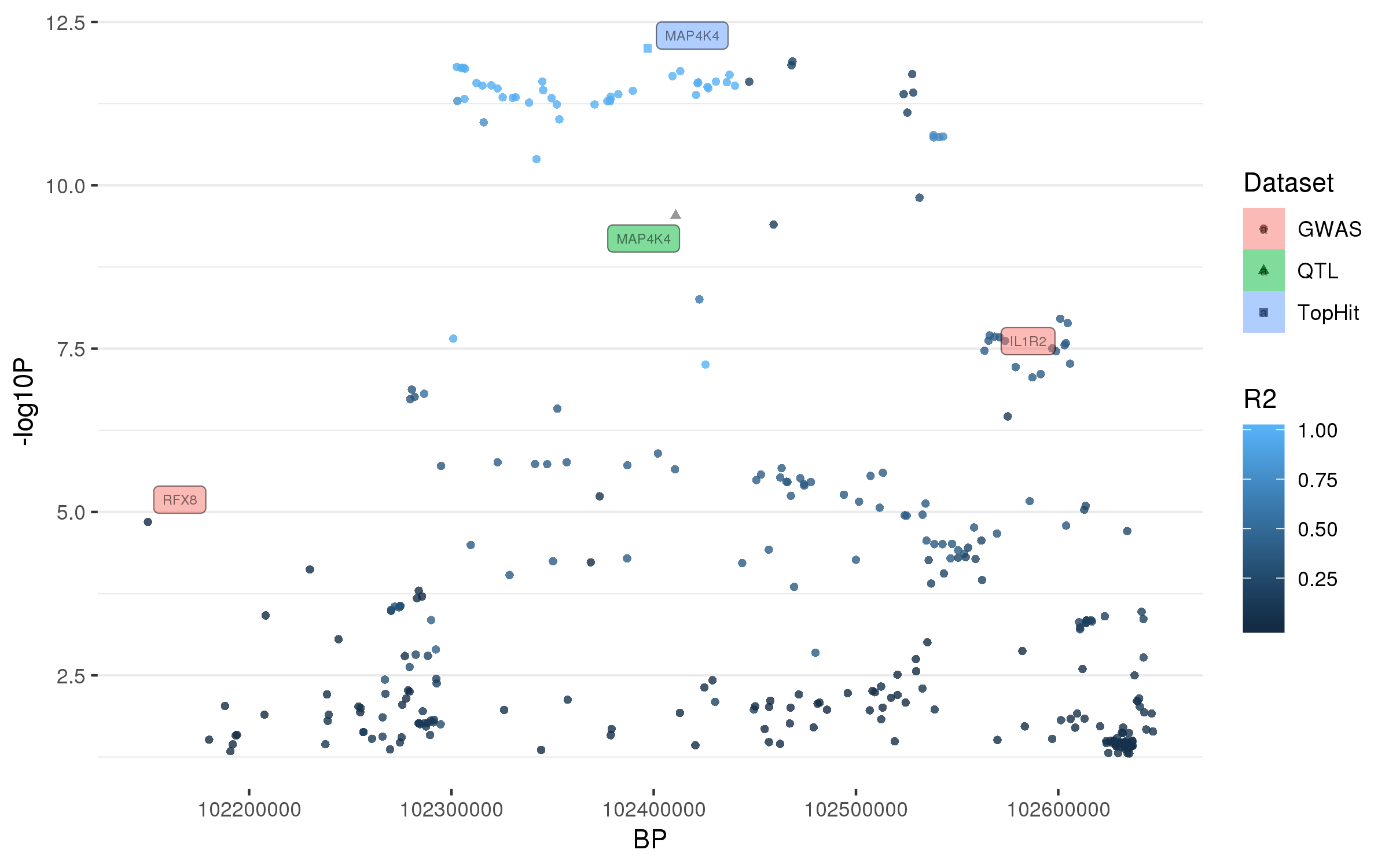

### rs11707416.png

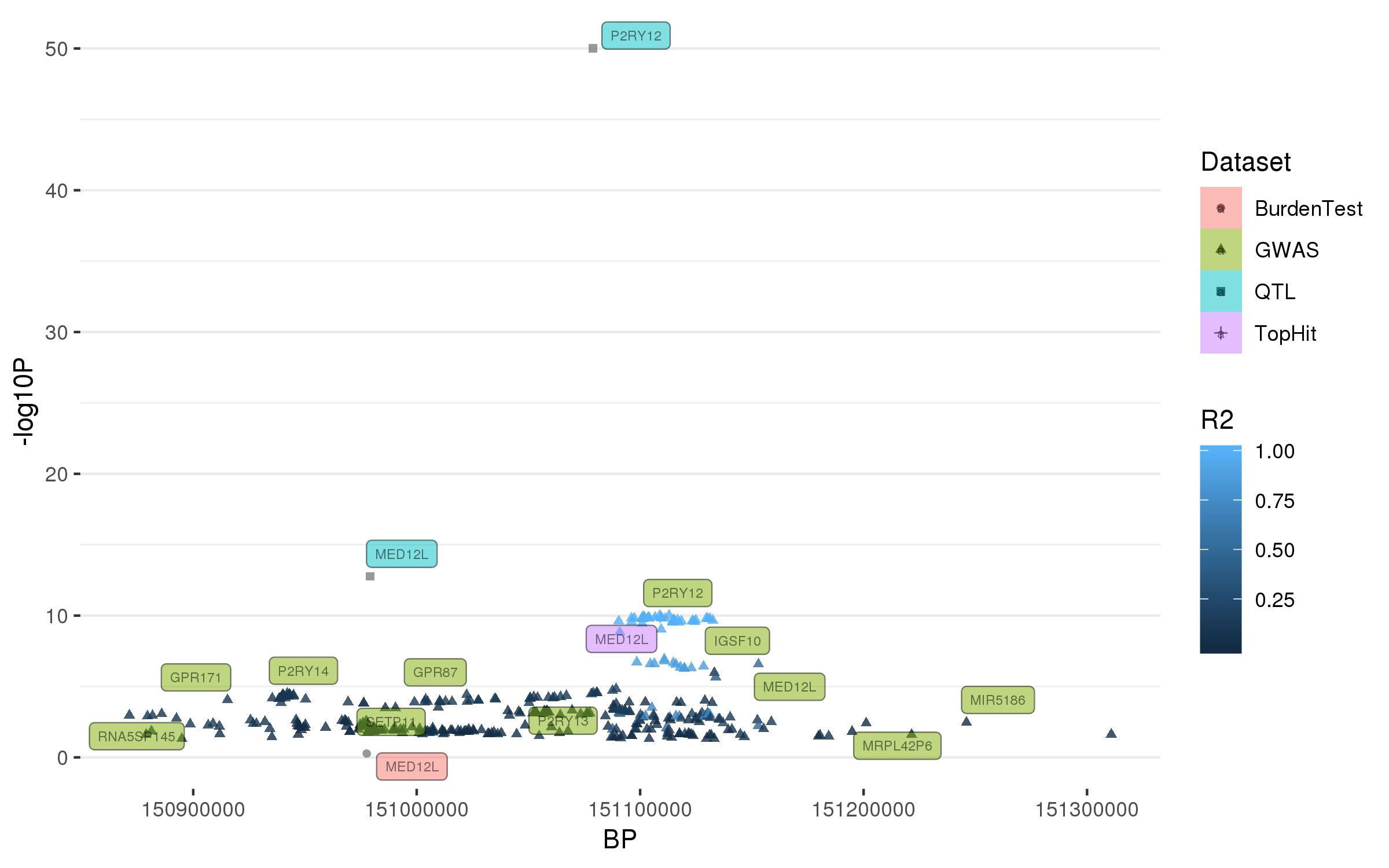

### rs11950533.png

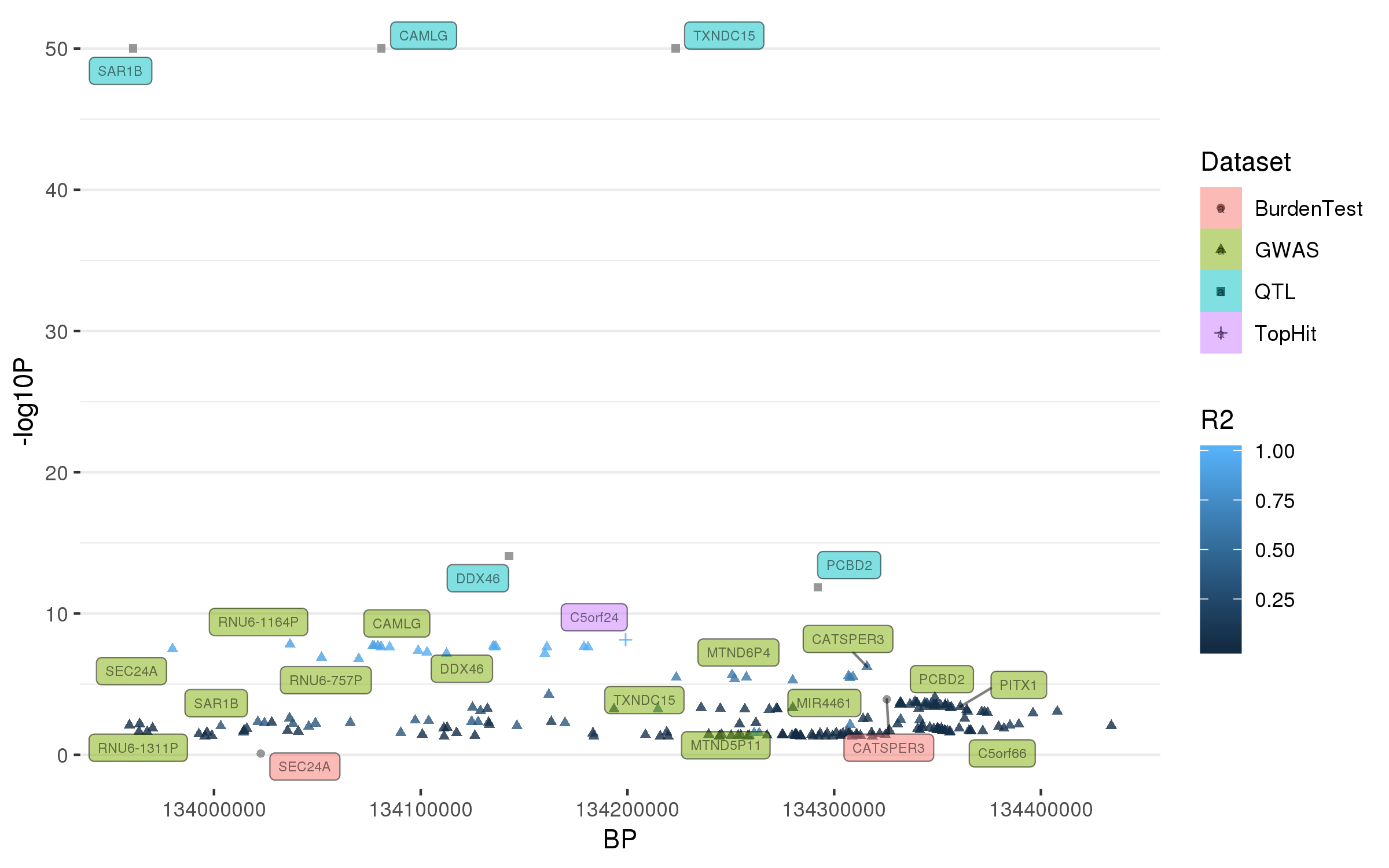

### rs12147950.png

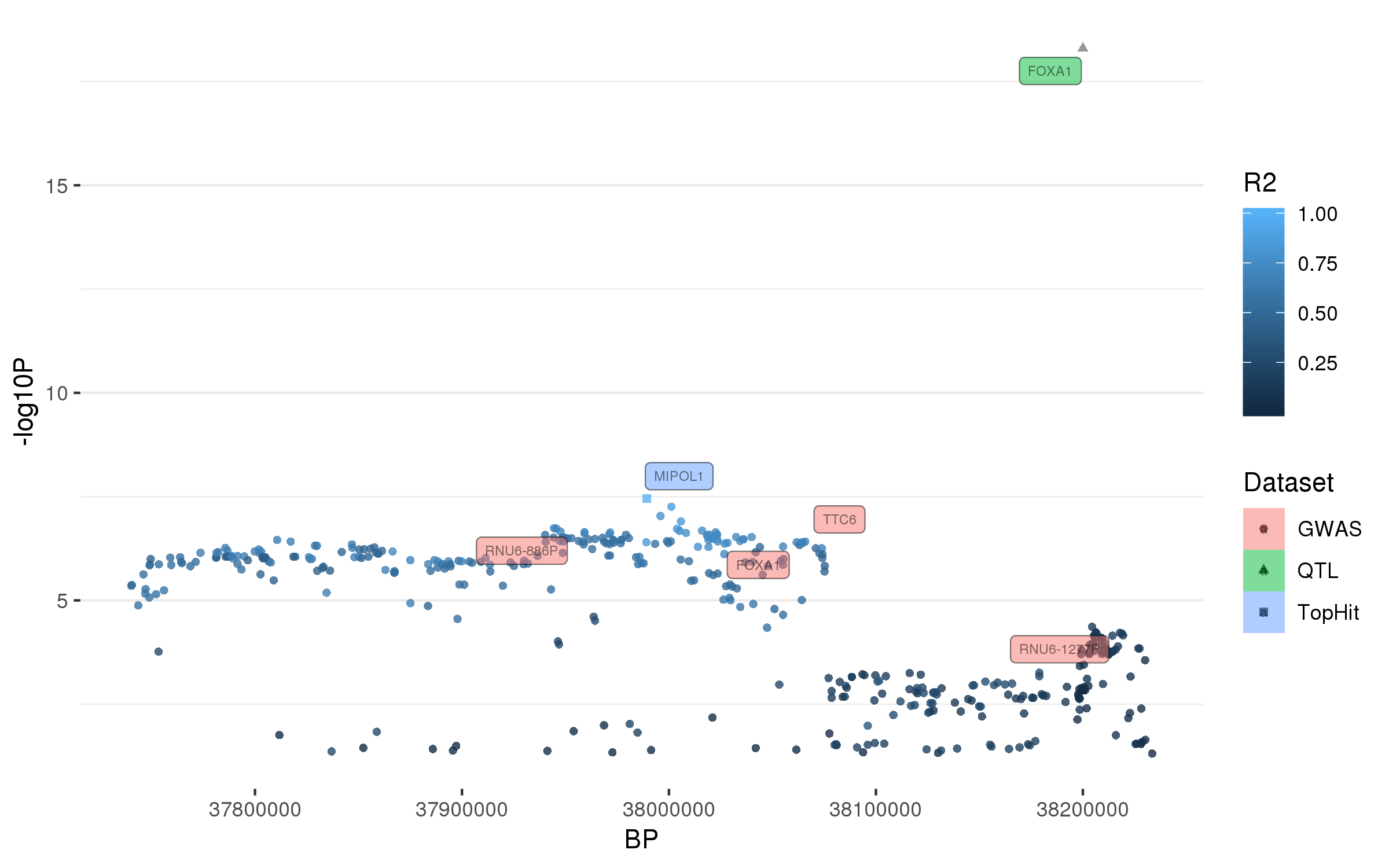

### rs12283611.png

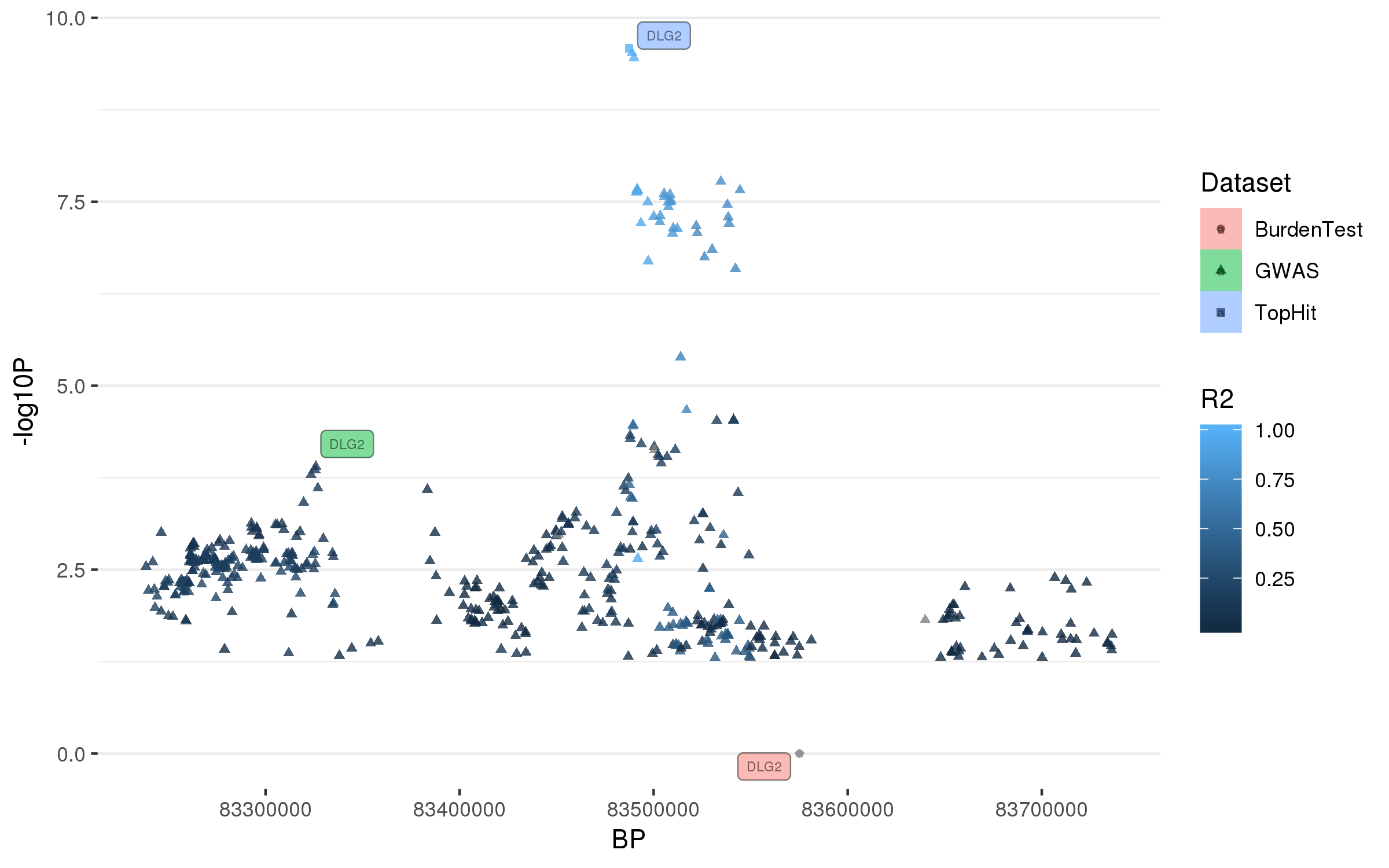

### rs12456492.png

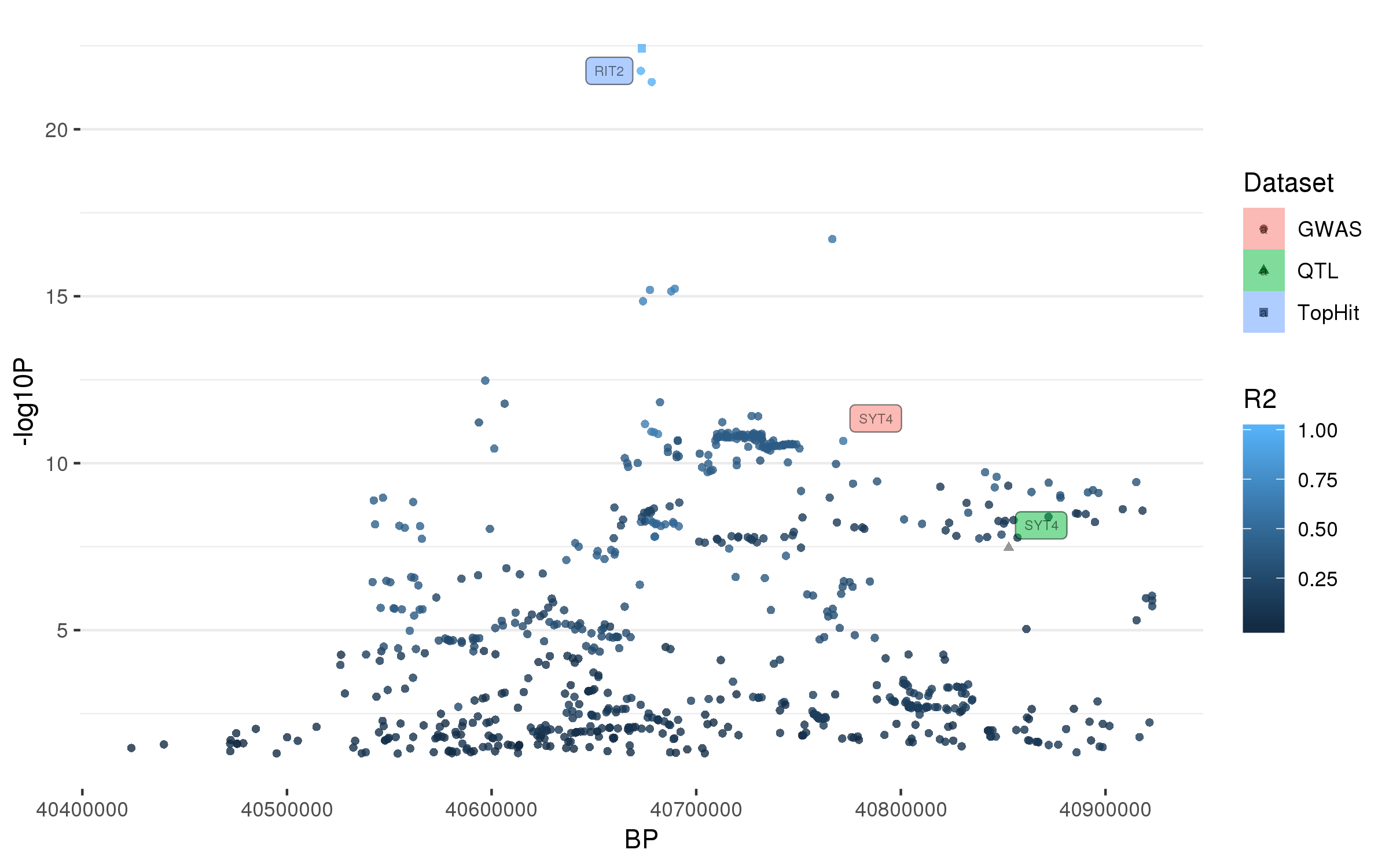

### rs12497850.png

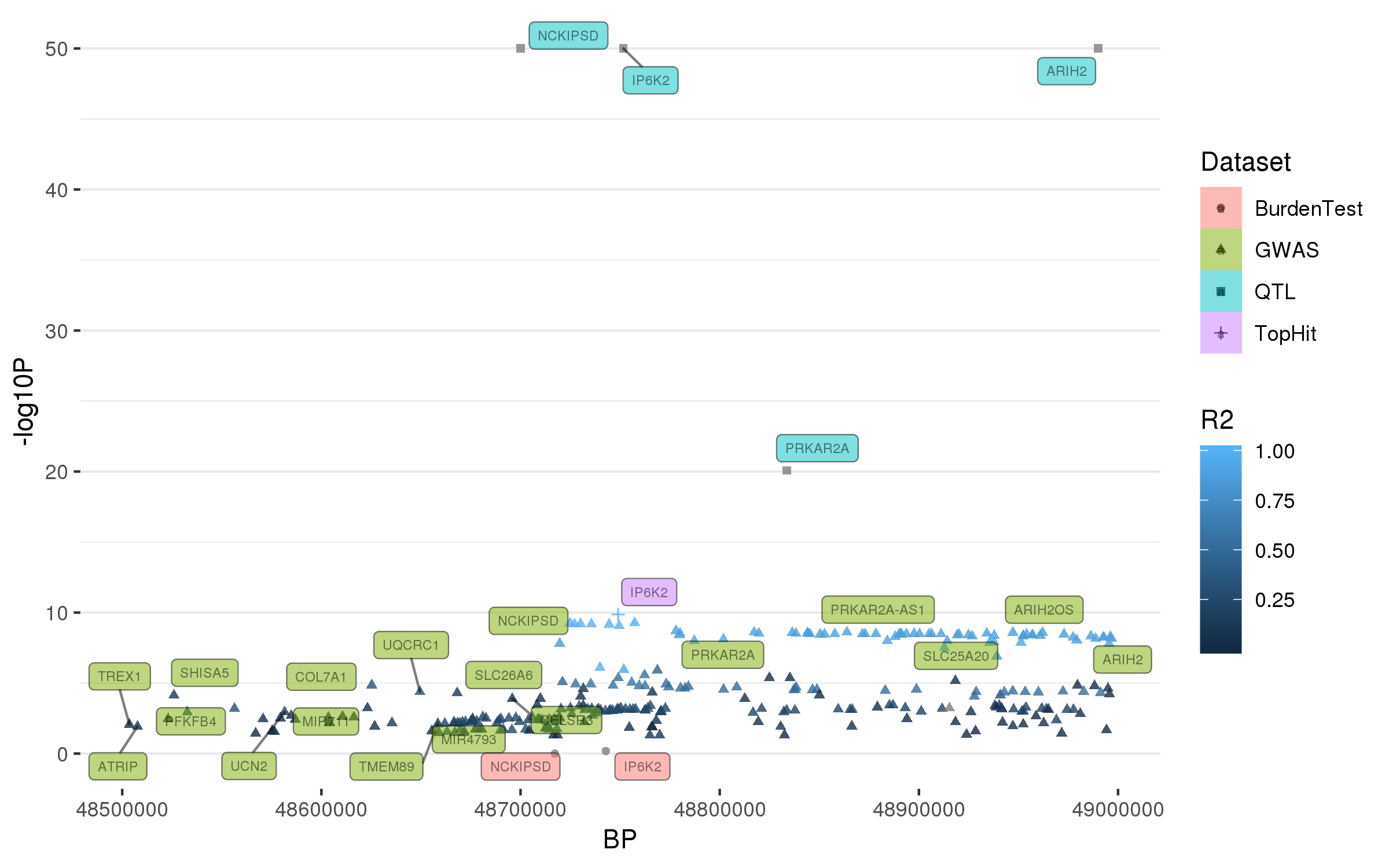

### rs12528068.png

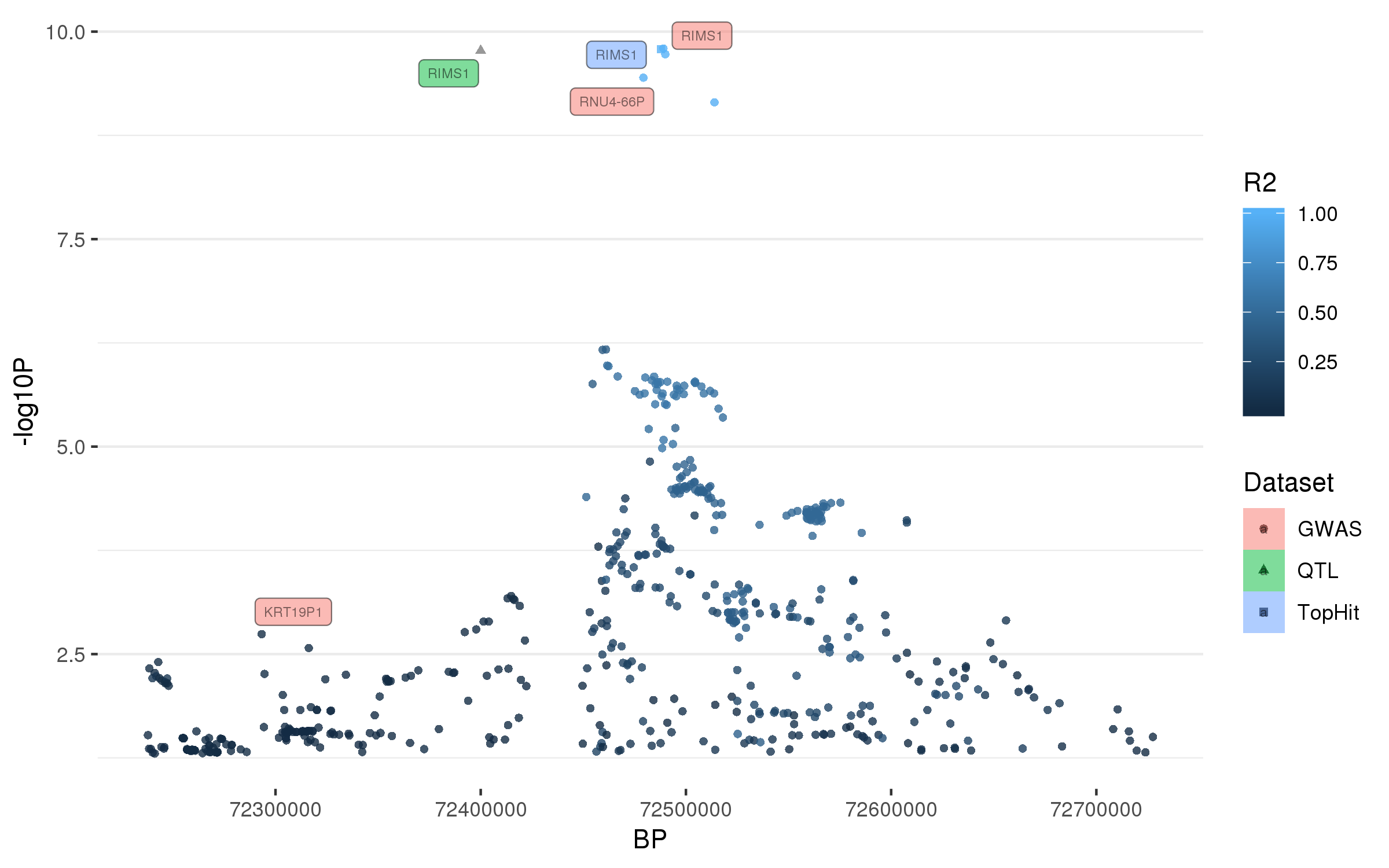

### rs12600861.png

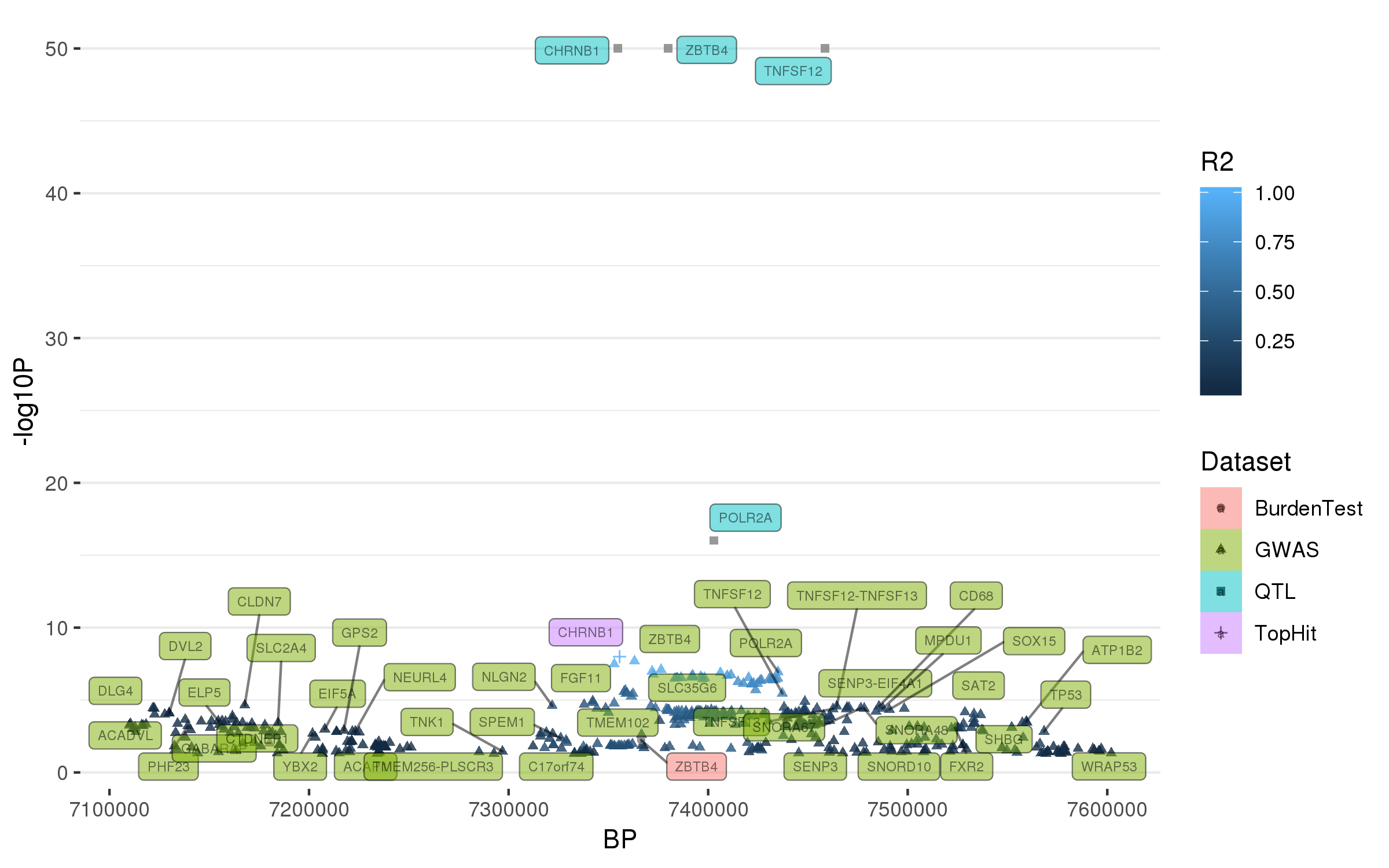

### rs12951632.png

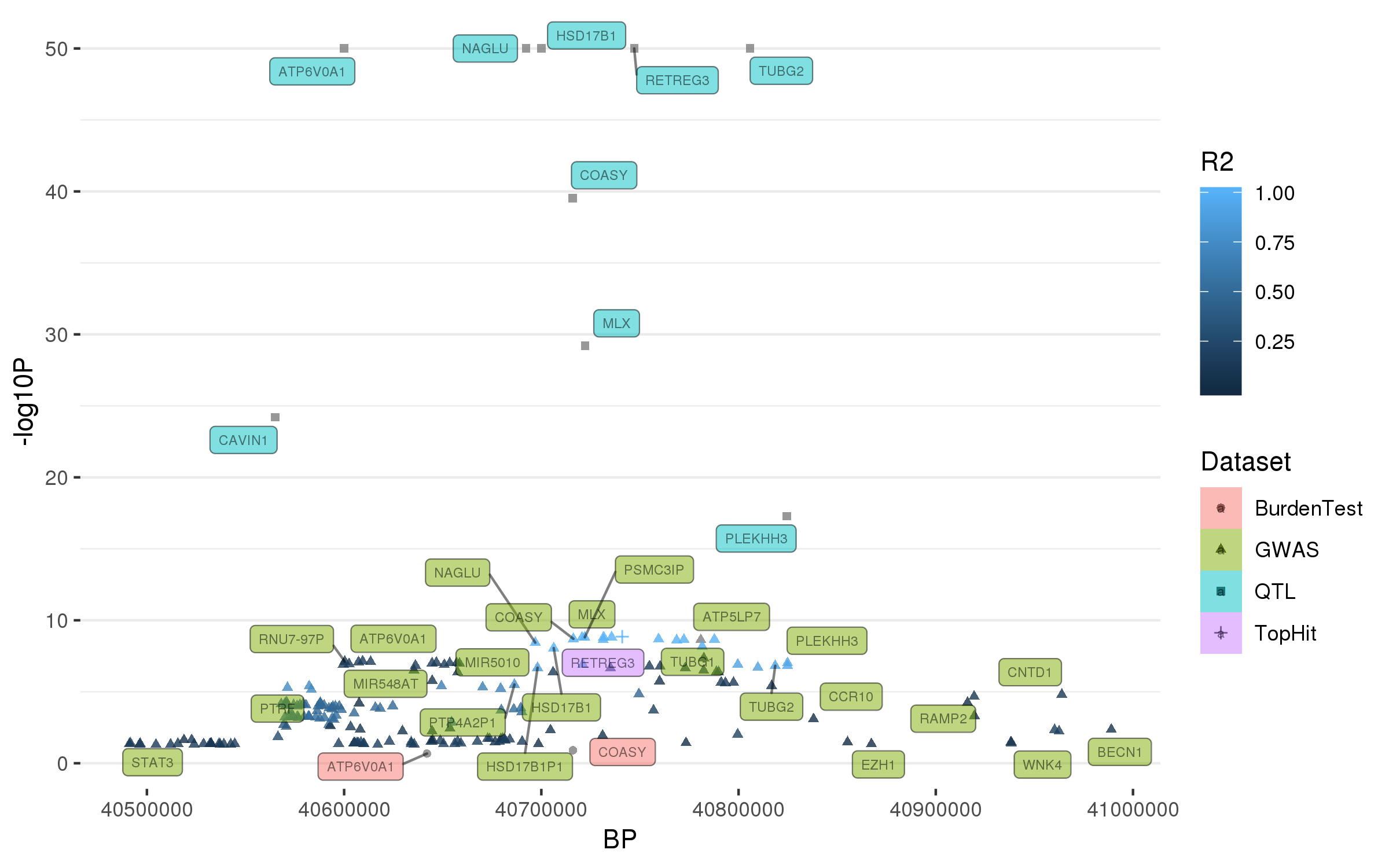

### rs13117519.png

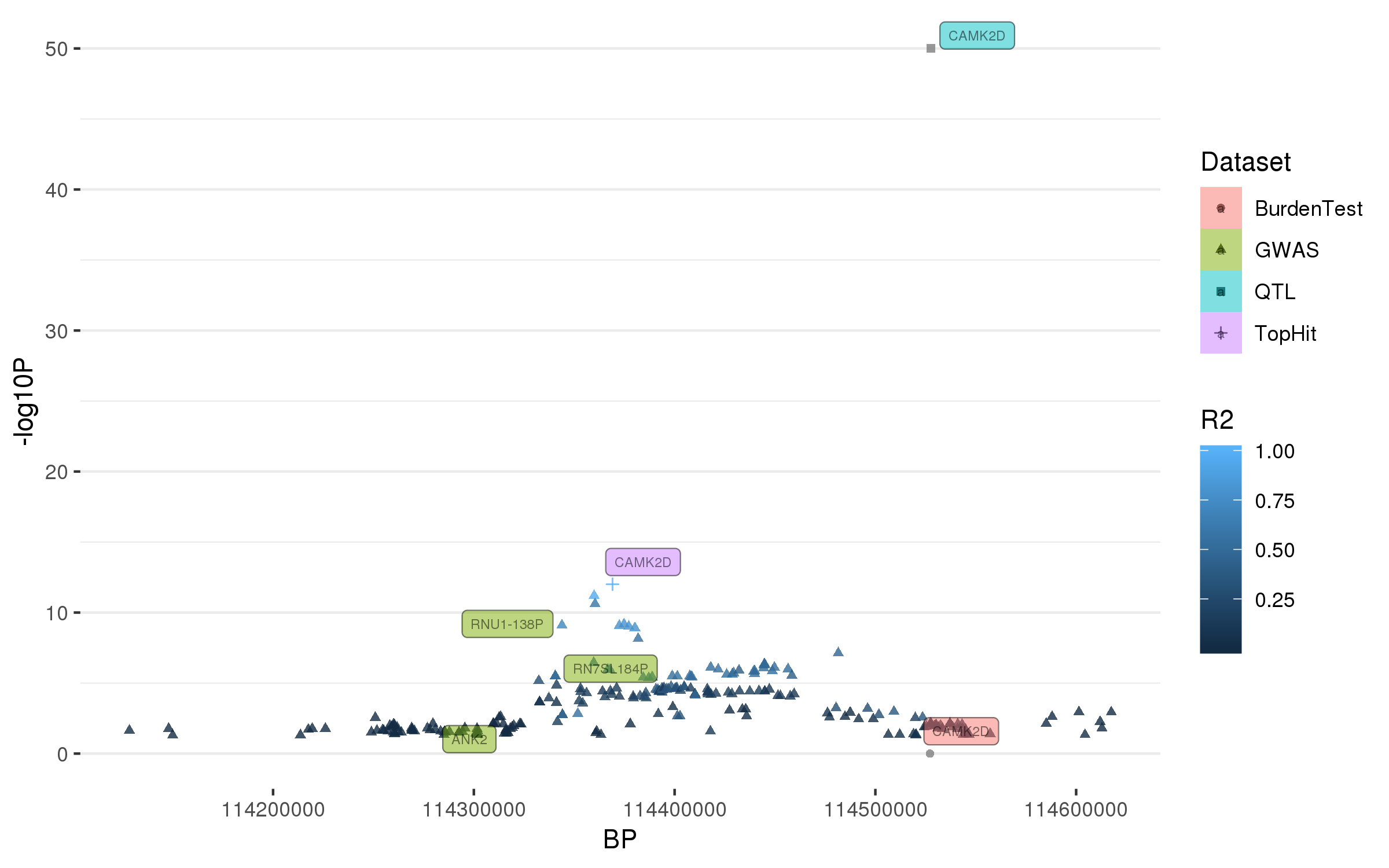

### rs13294100.png

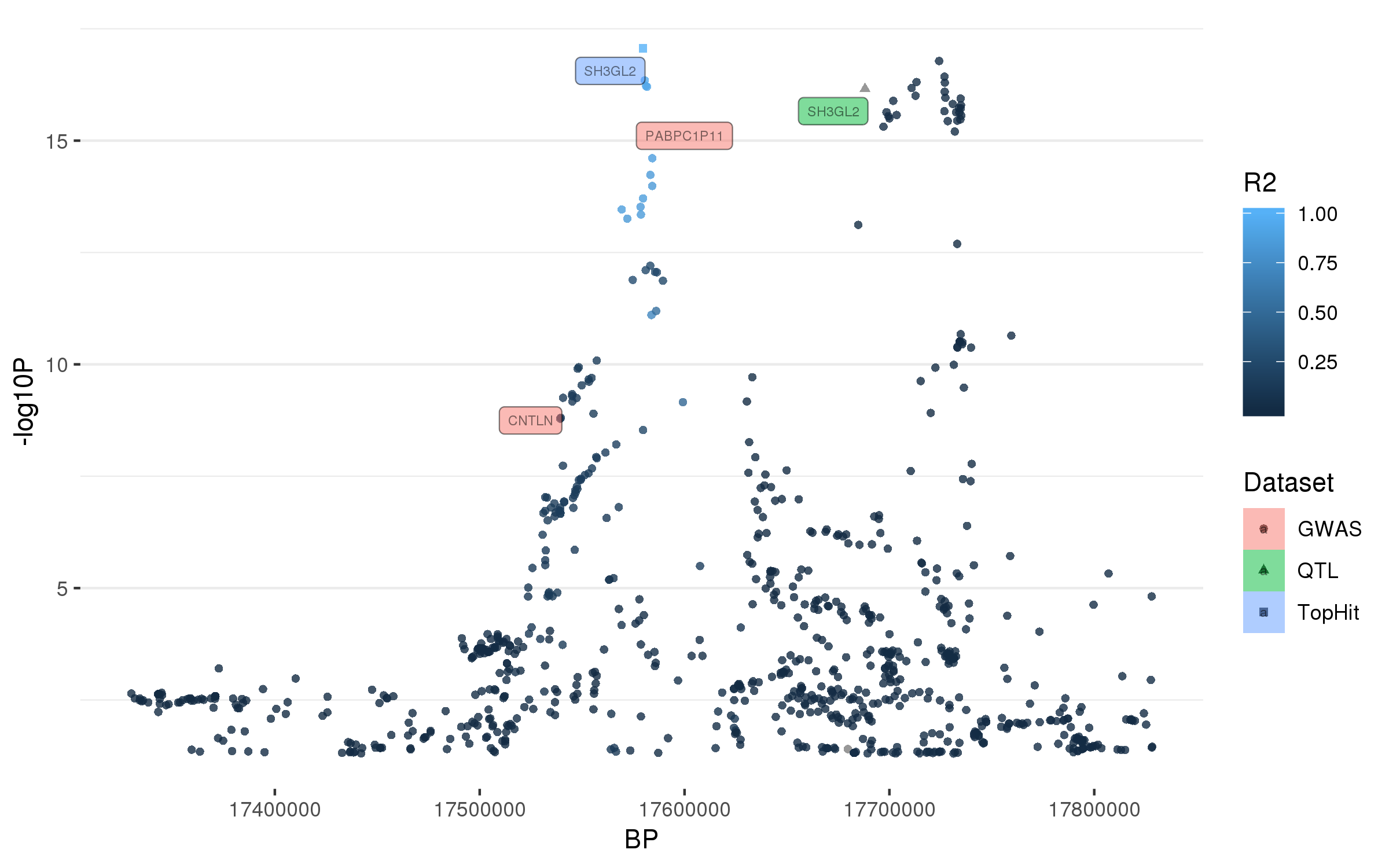

### rs34025766.png

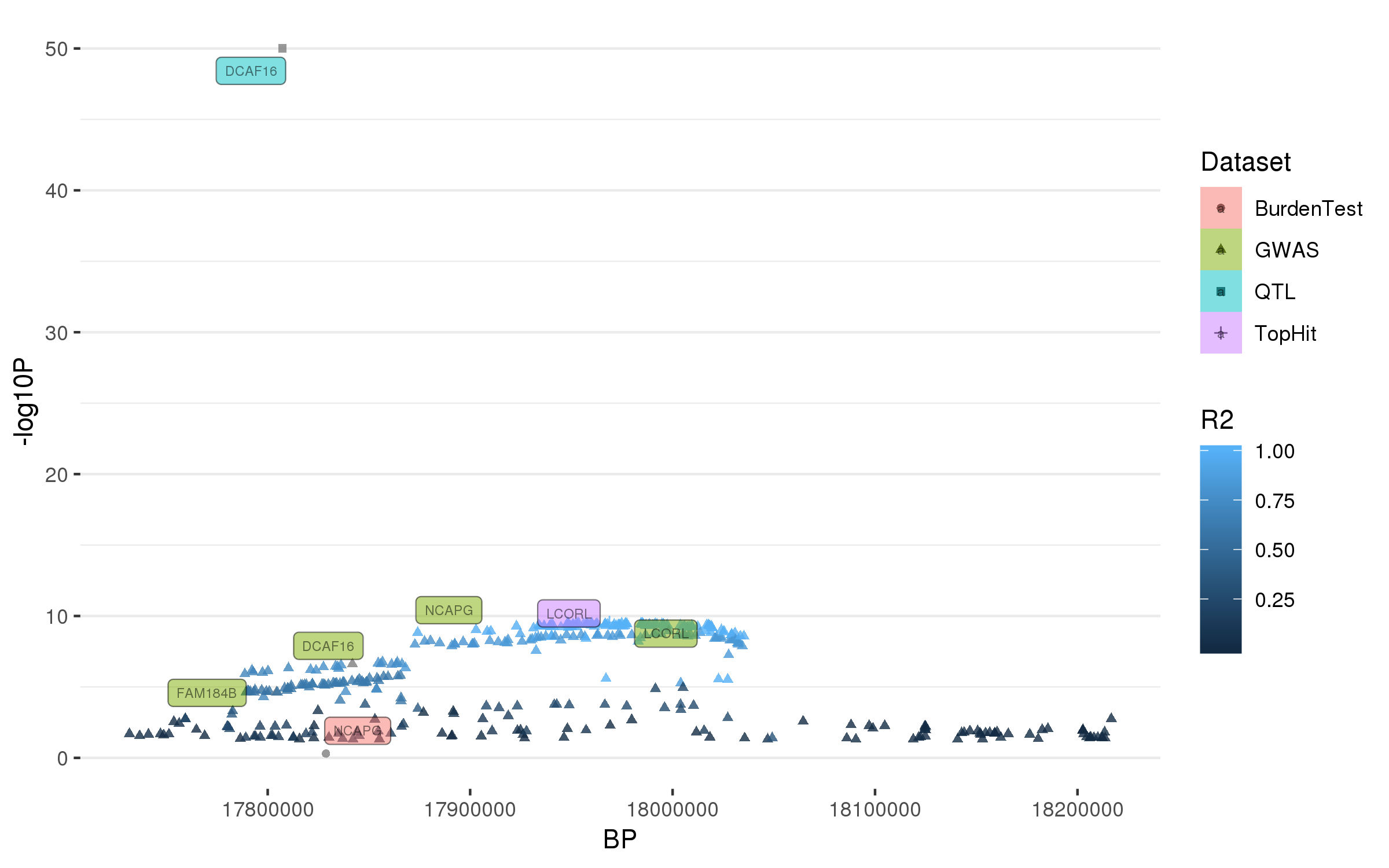

### rs34311866.png

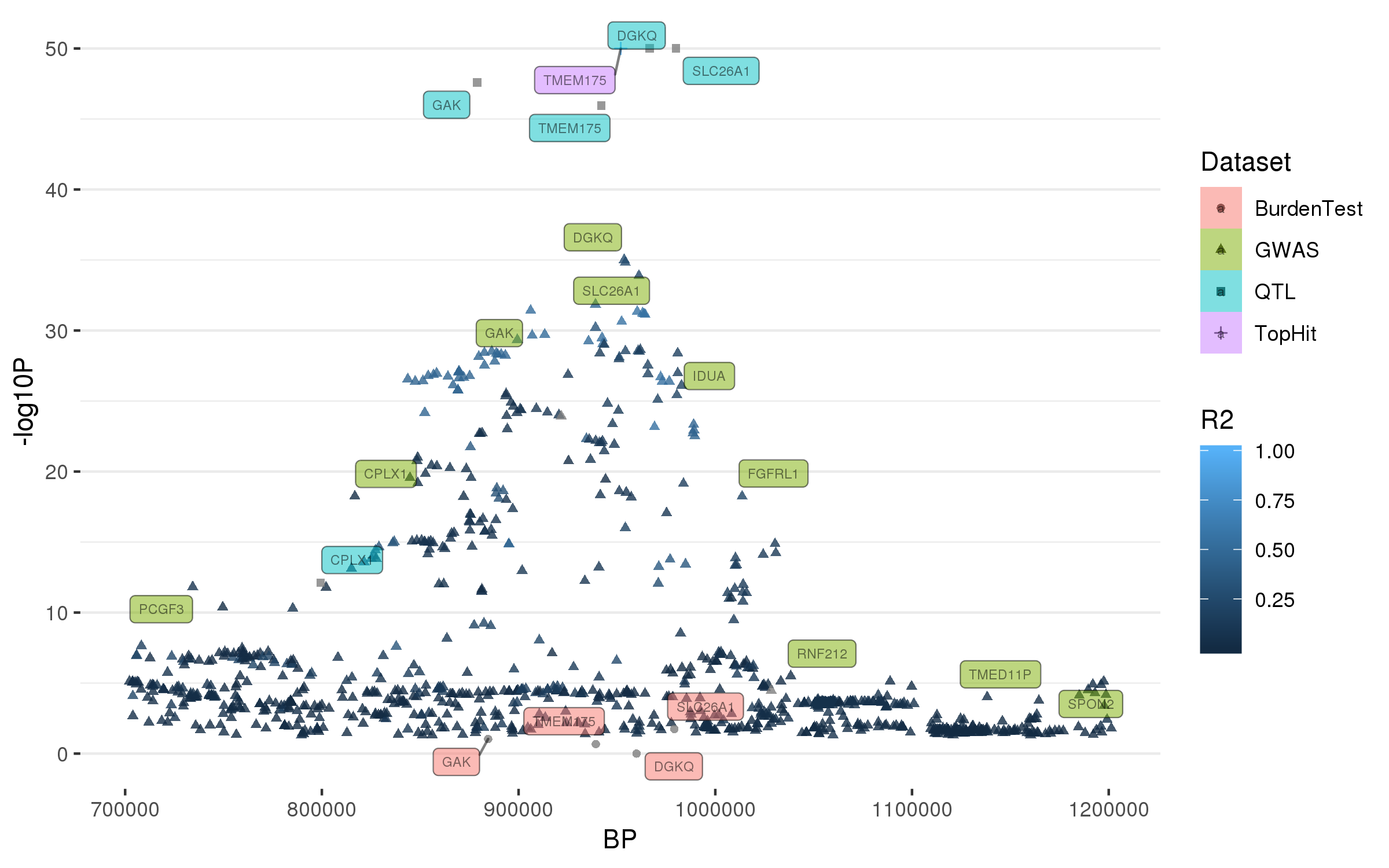

### rs34637584.png

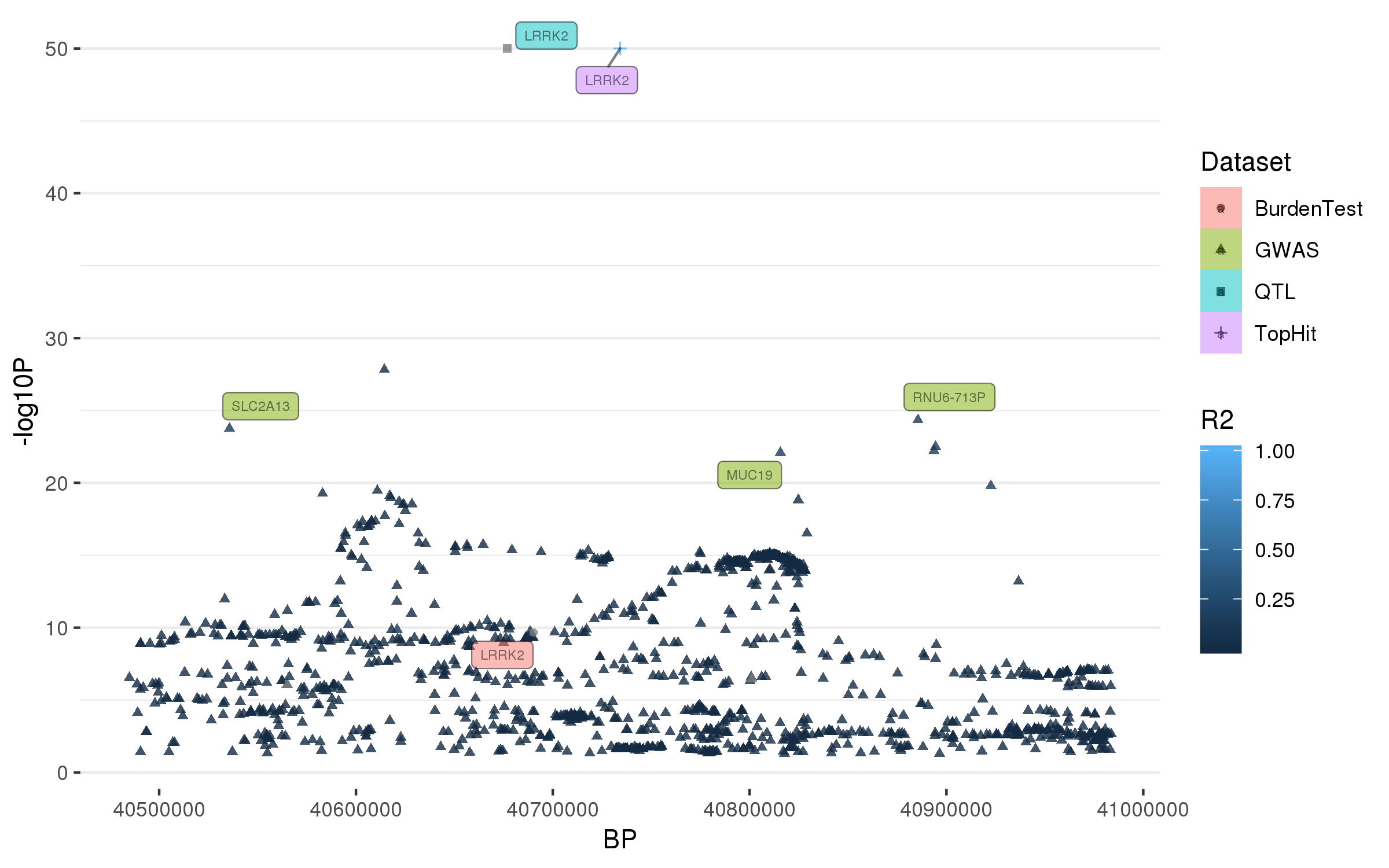

### rs35749011.png

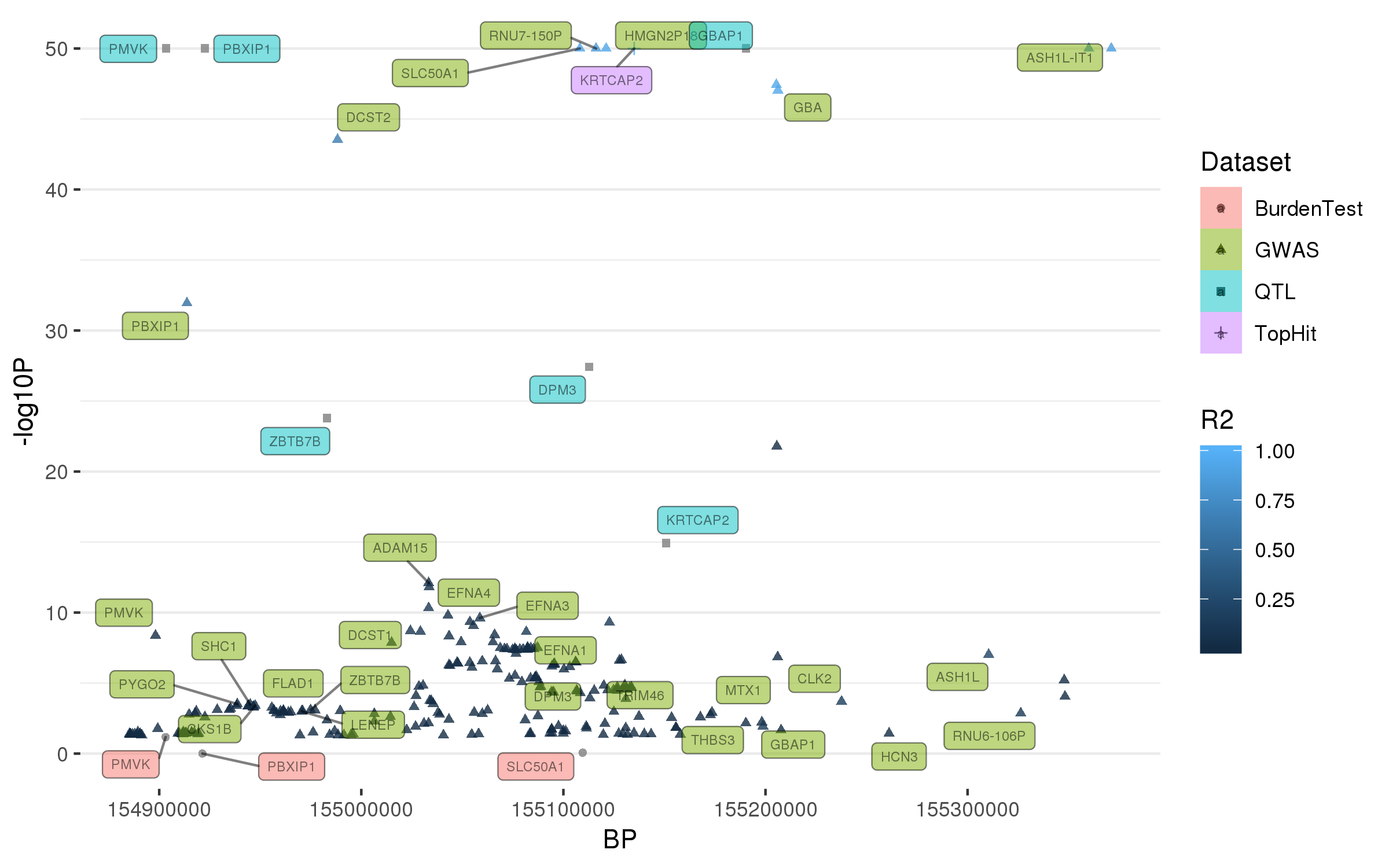

### rs55818311.png

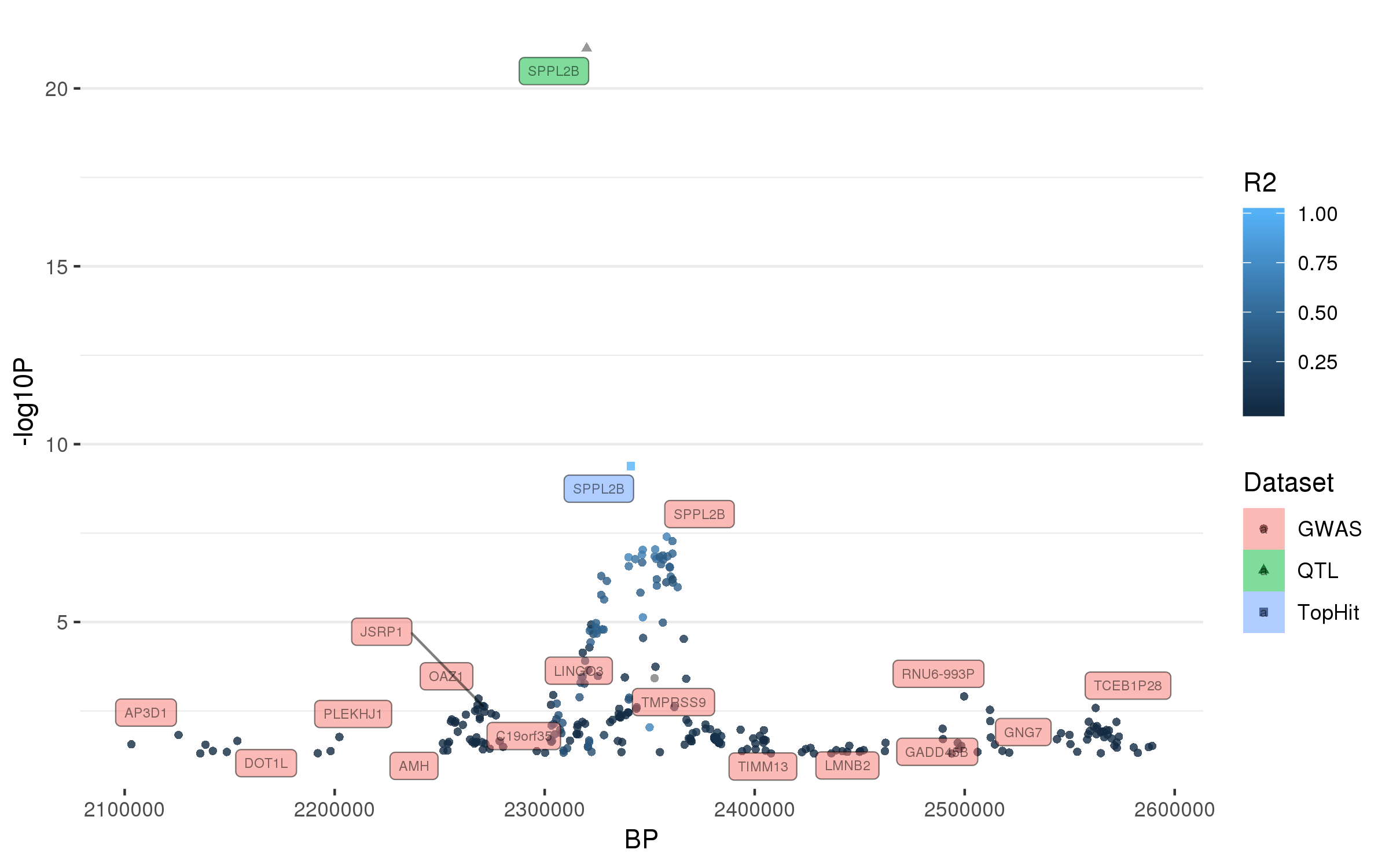

### rs55961674.png

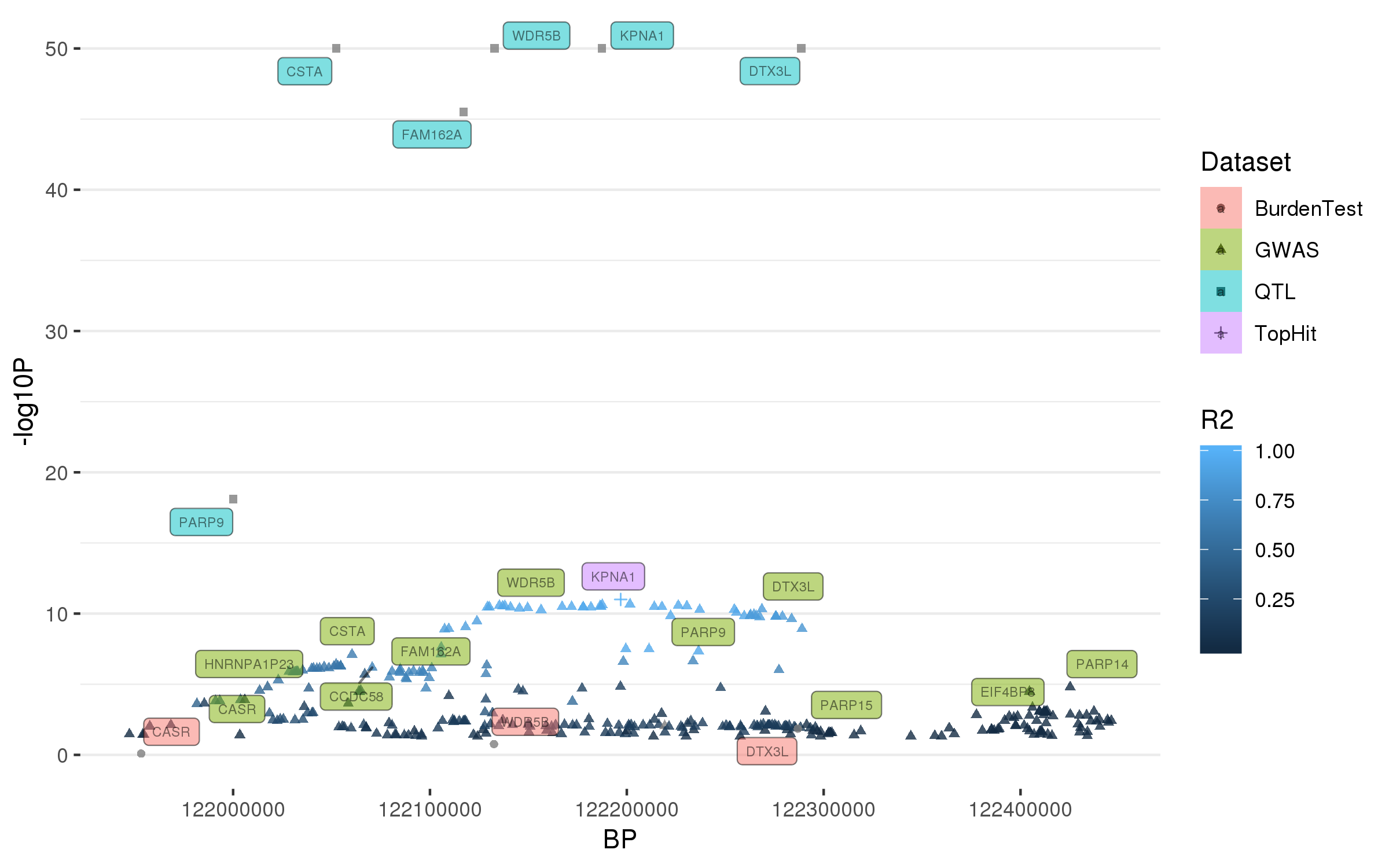

### rs57891859.png

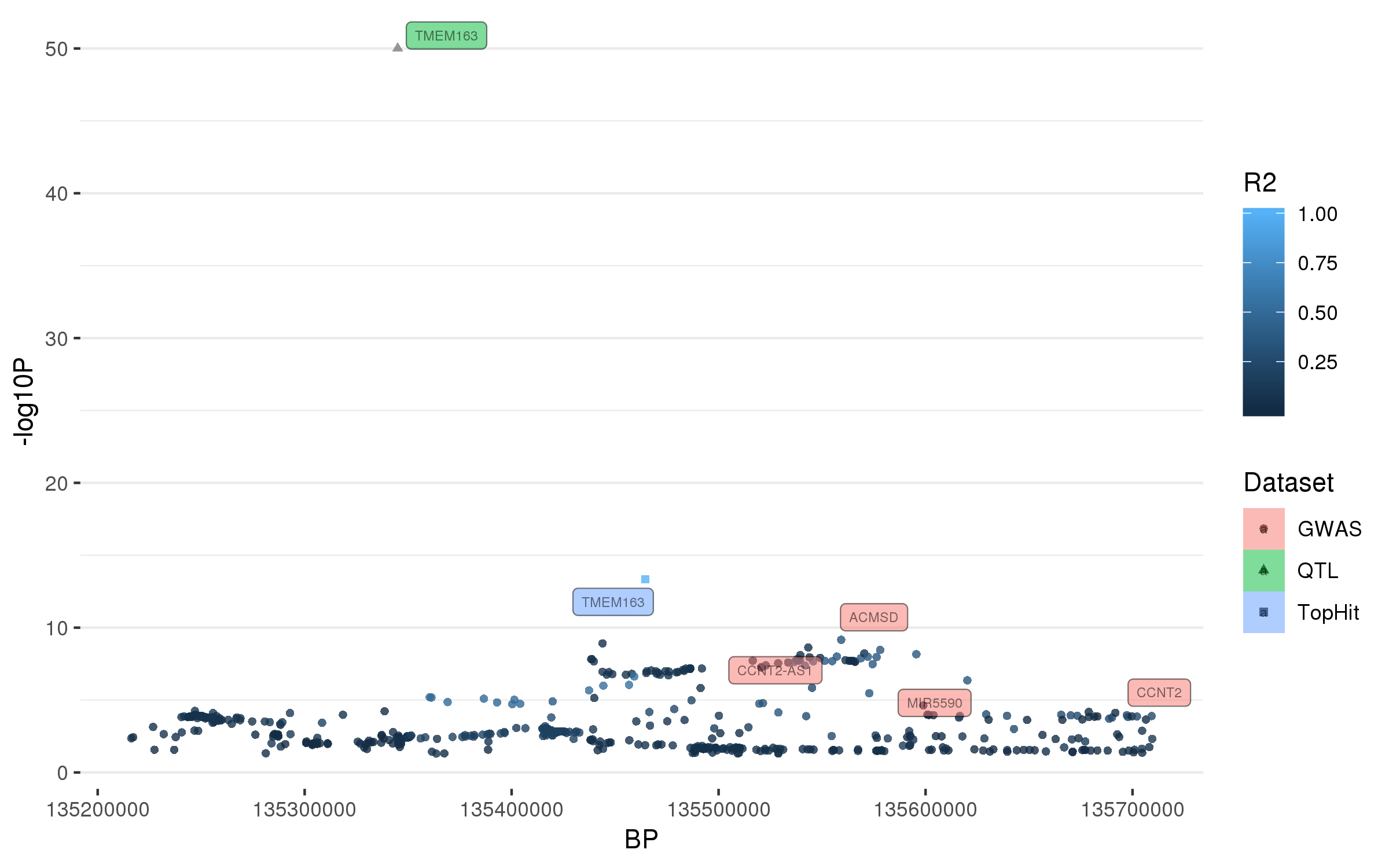

### rs61169879.png

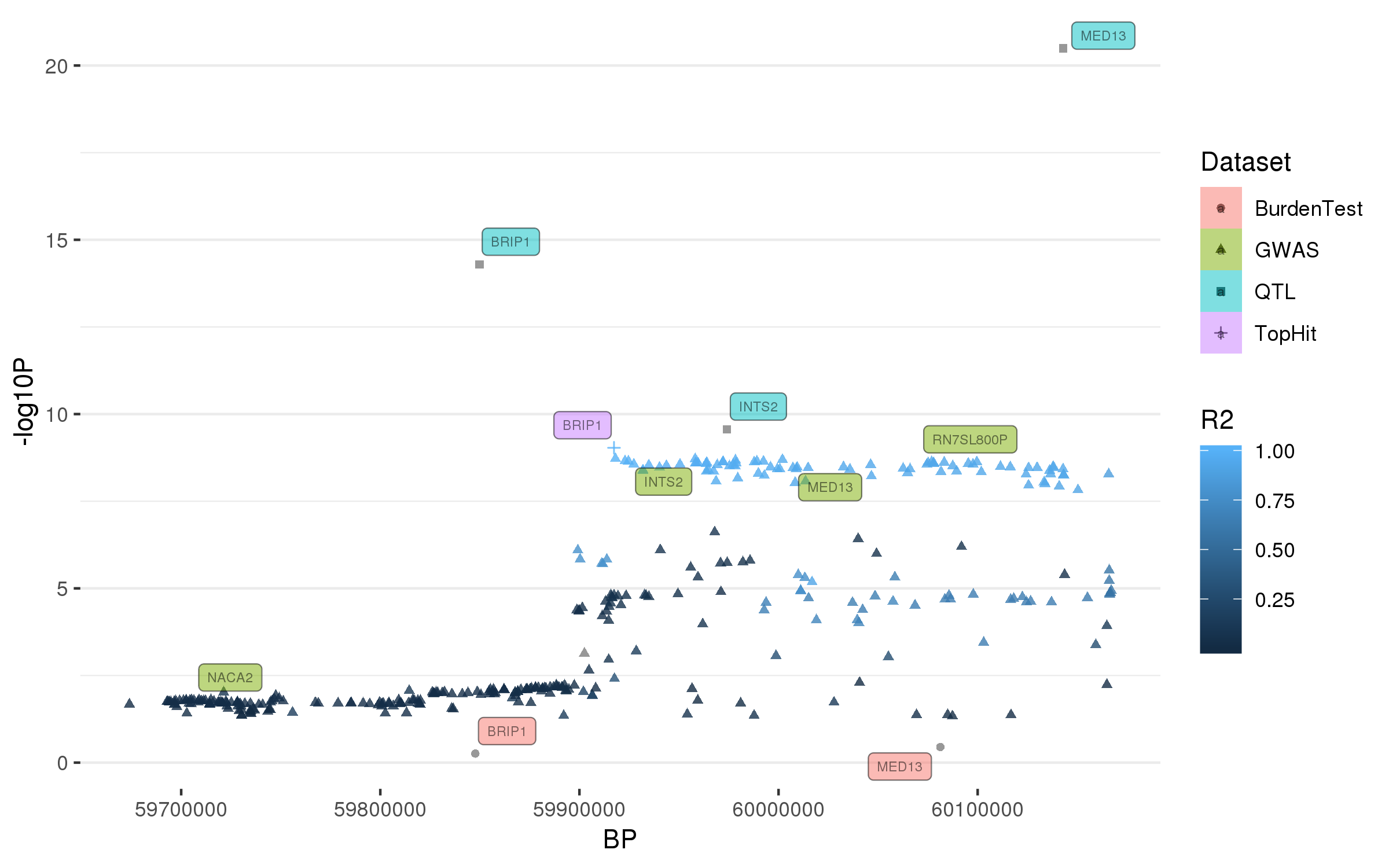

### rs62053943.png

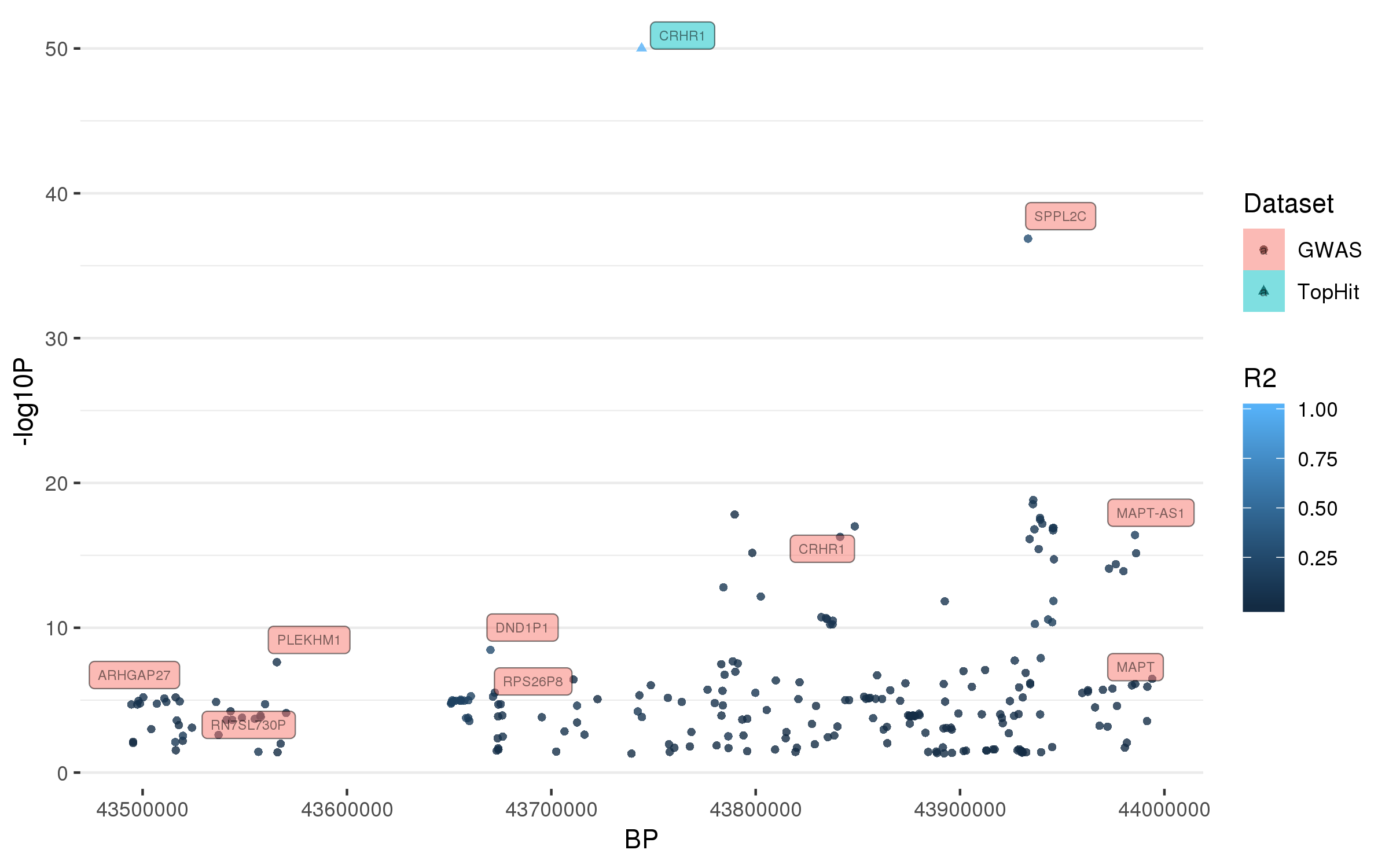

### rs62333164.png

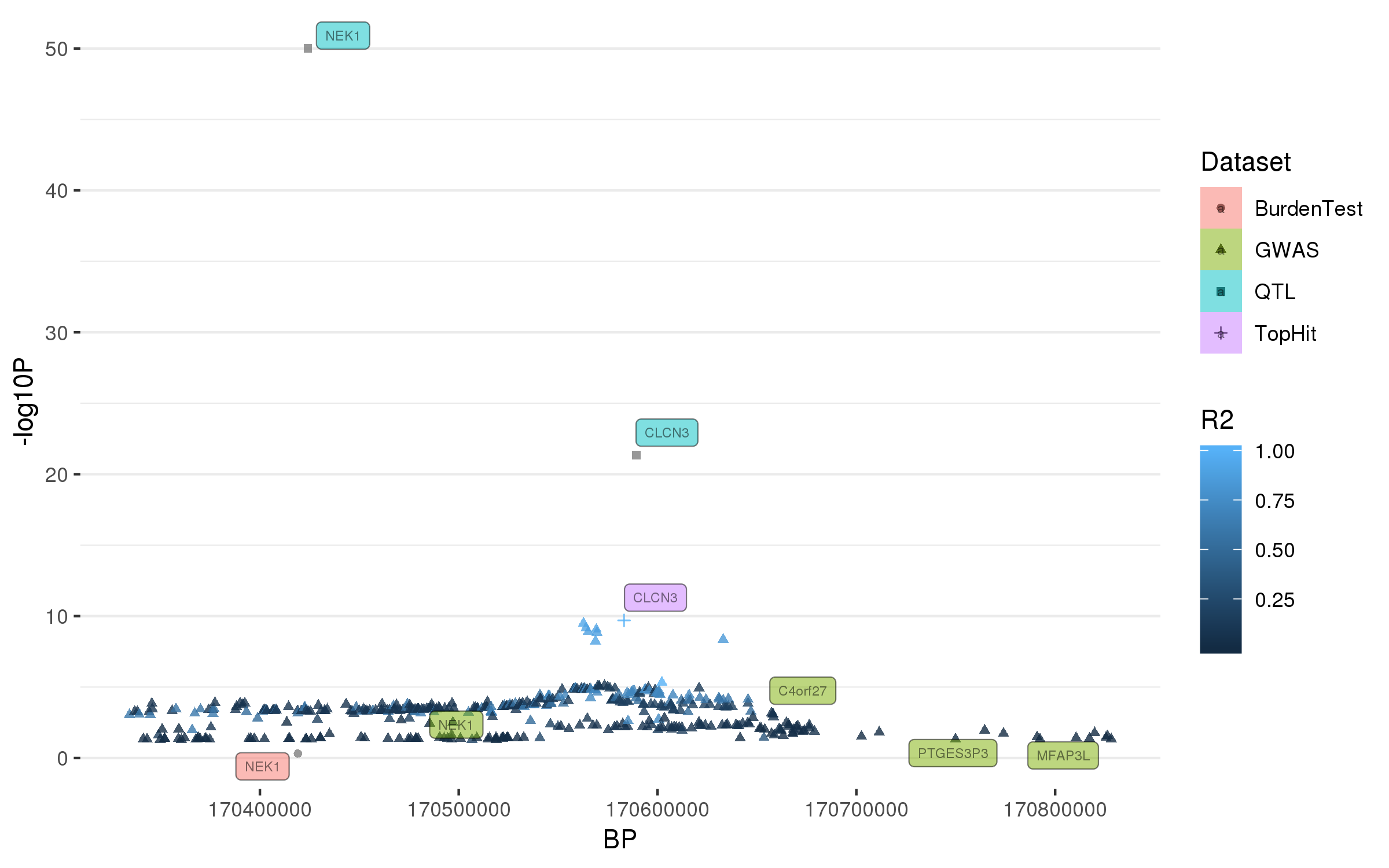

### rs72840788.png

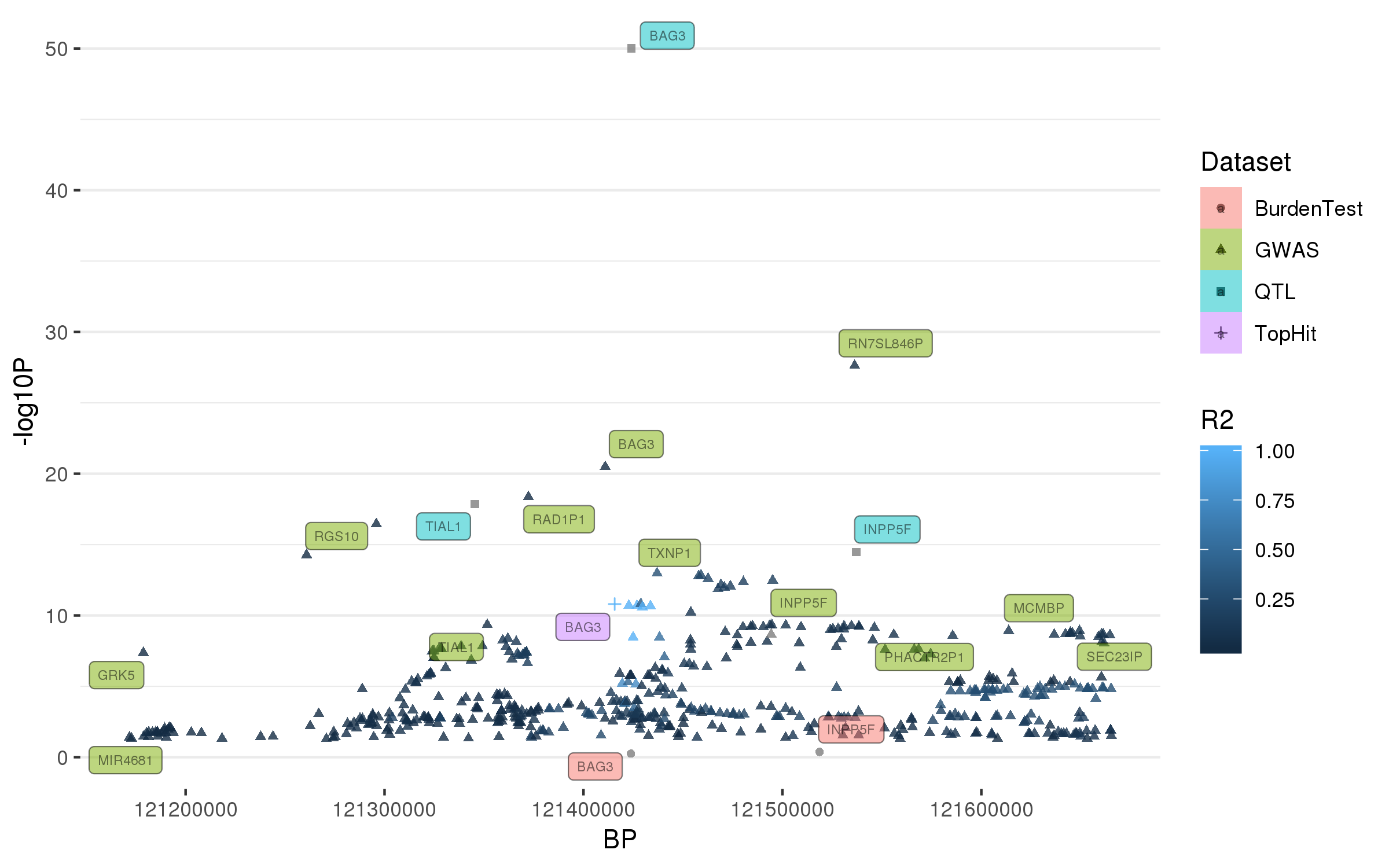

### rs73038319.png

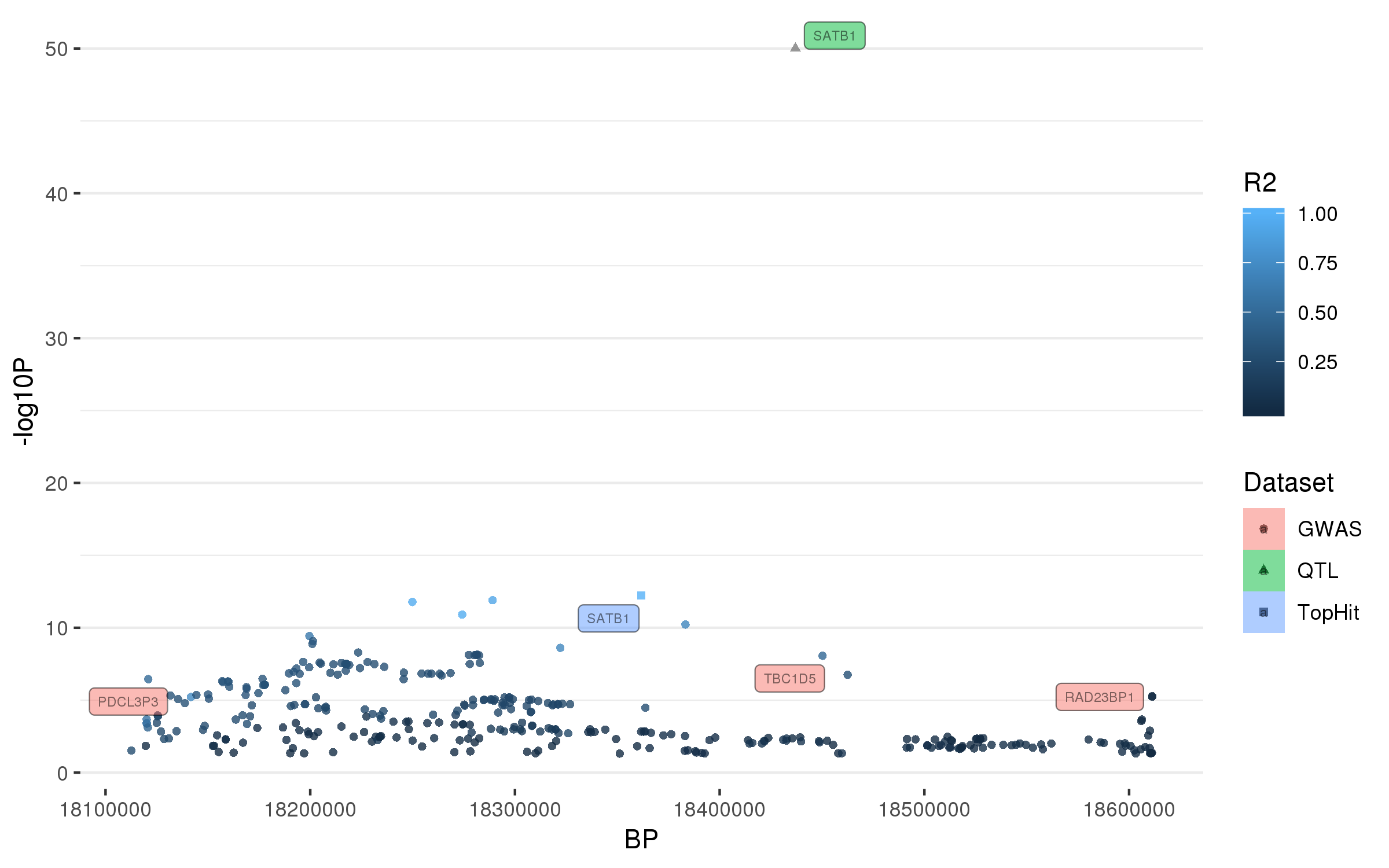

### rs75859381.png

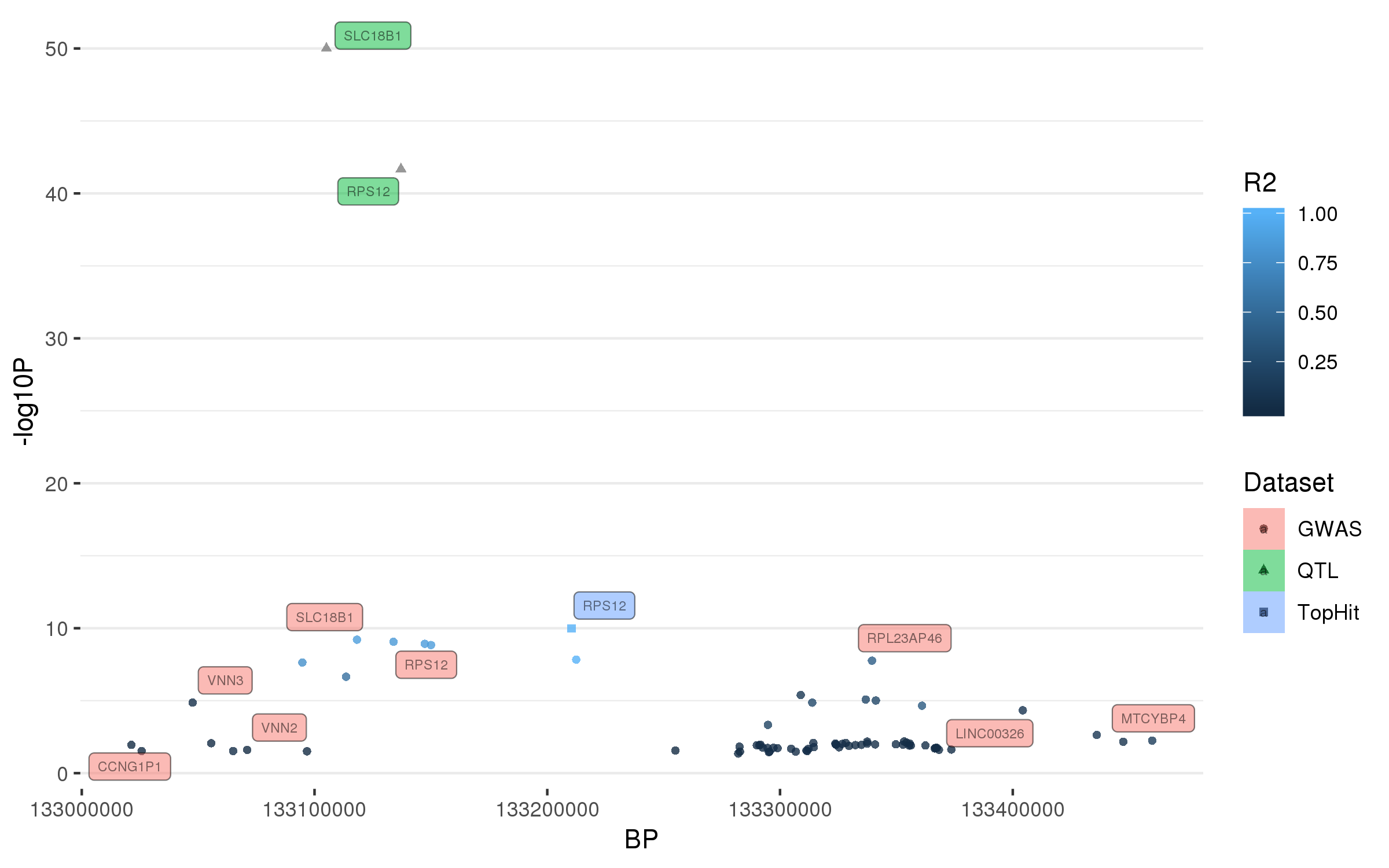

### rs76116224.png

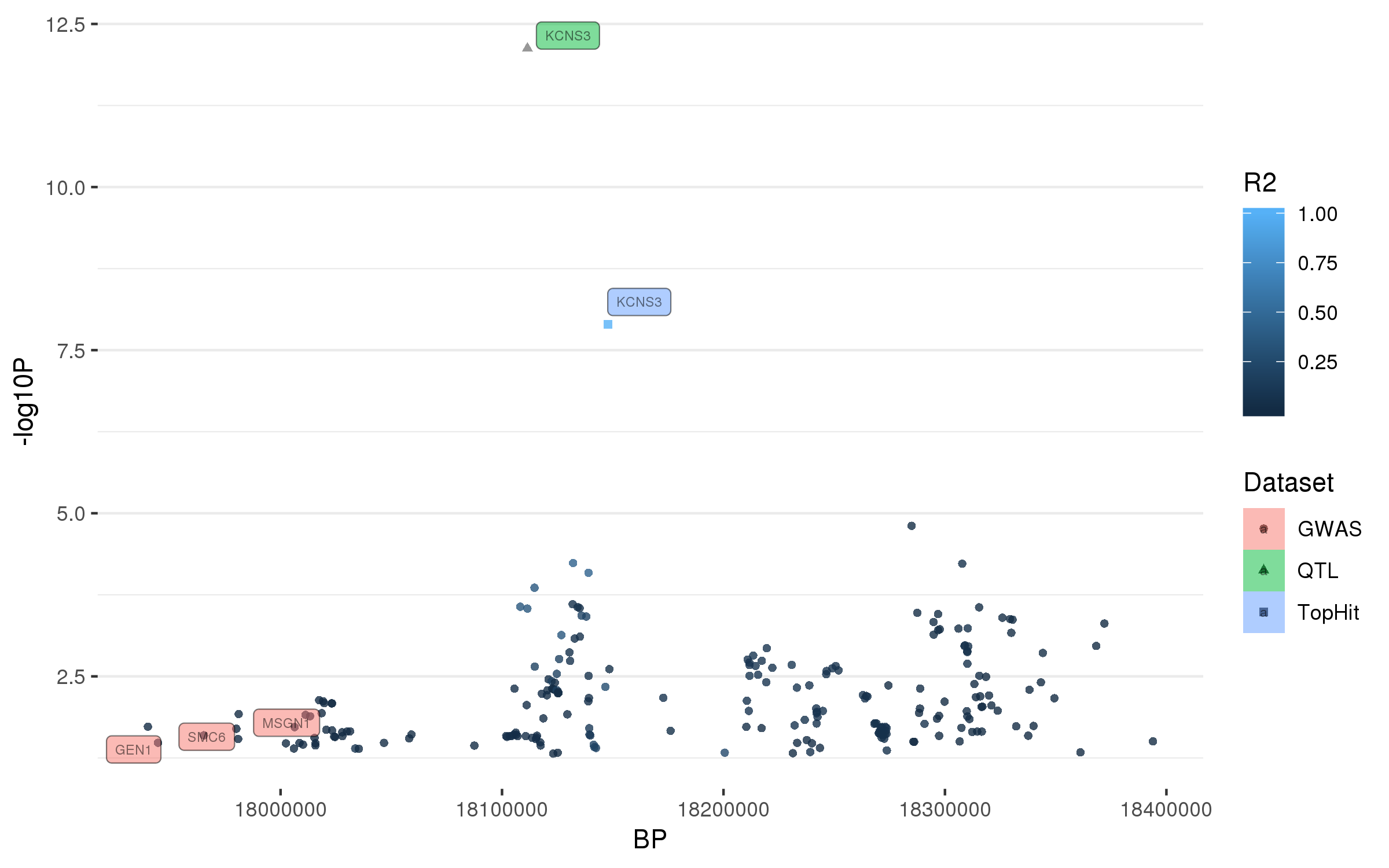

### rs76763715.png

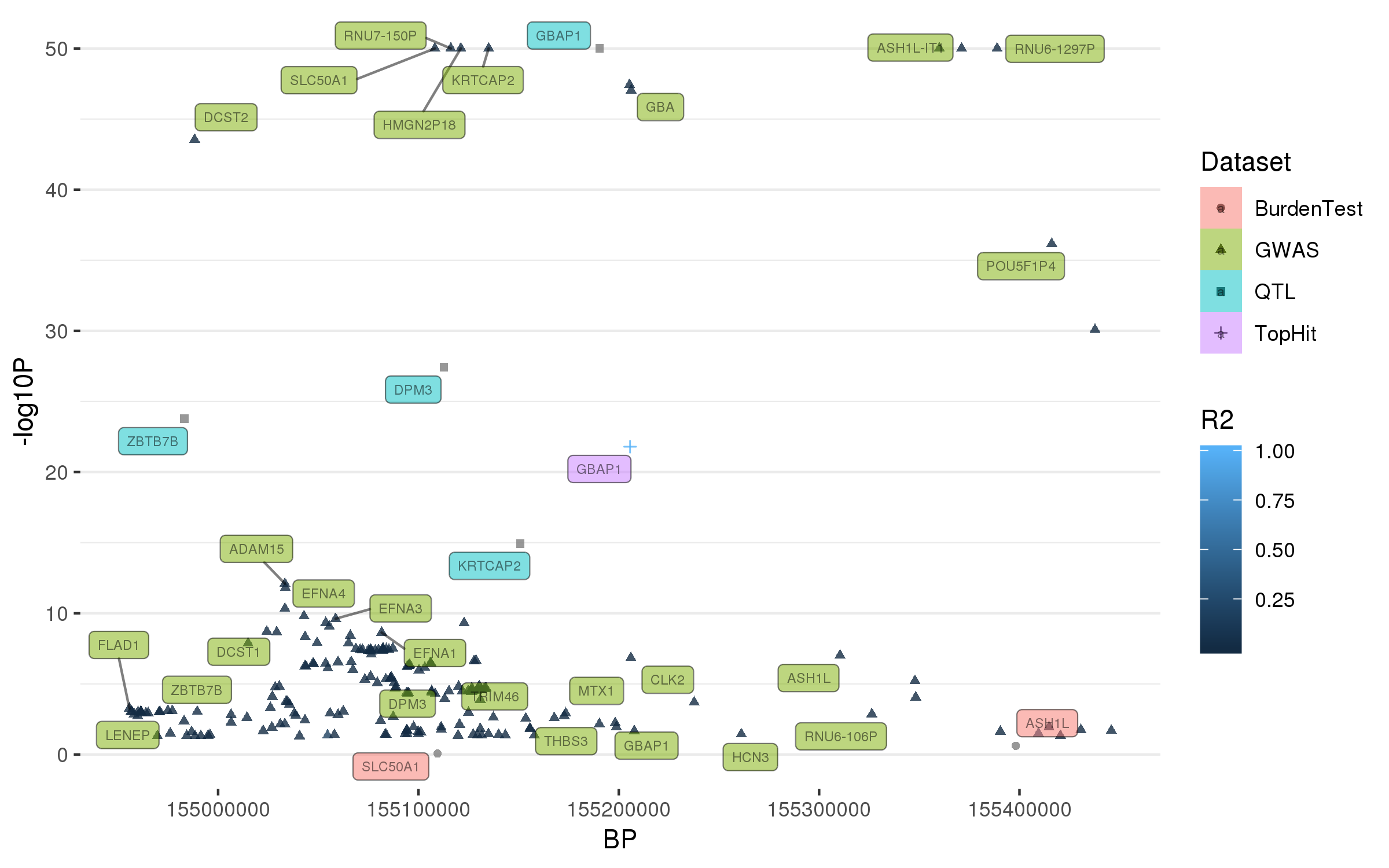

### rs76904798.png

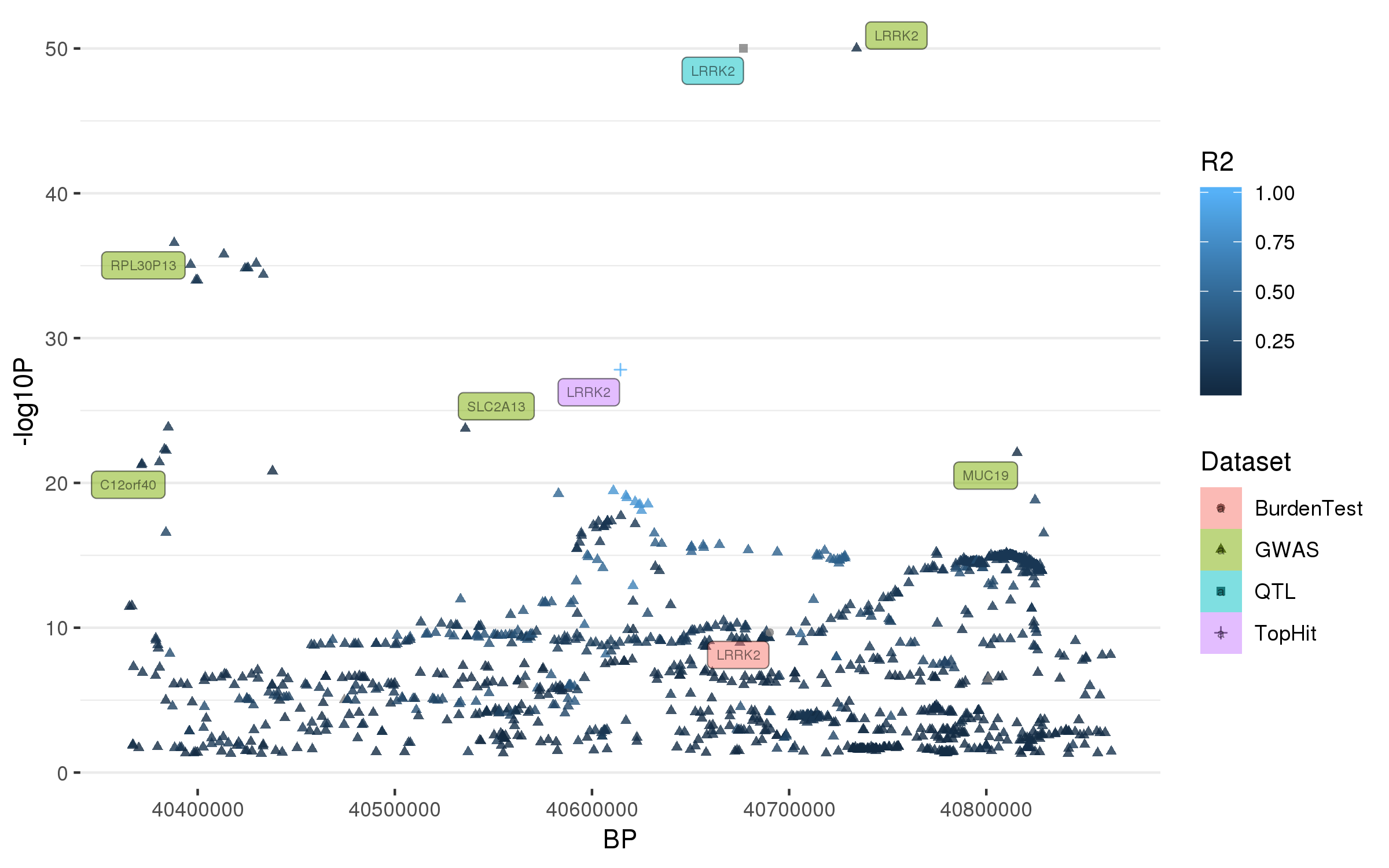

### rs76949143.png

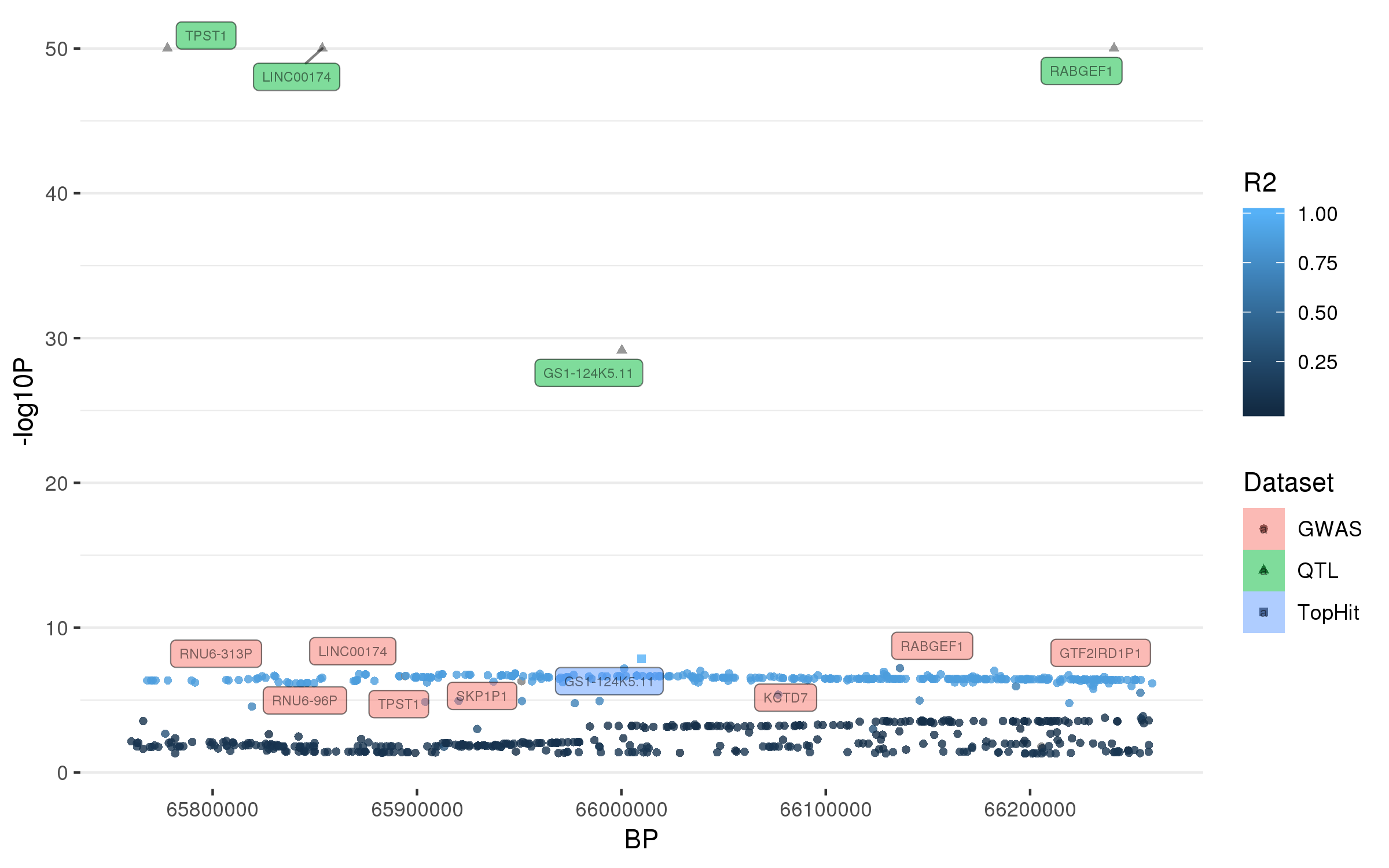

### rs77351827.png

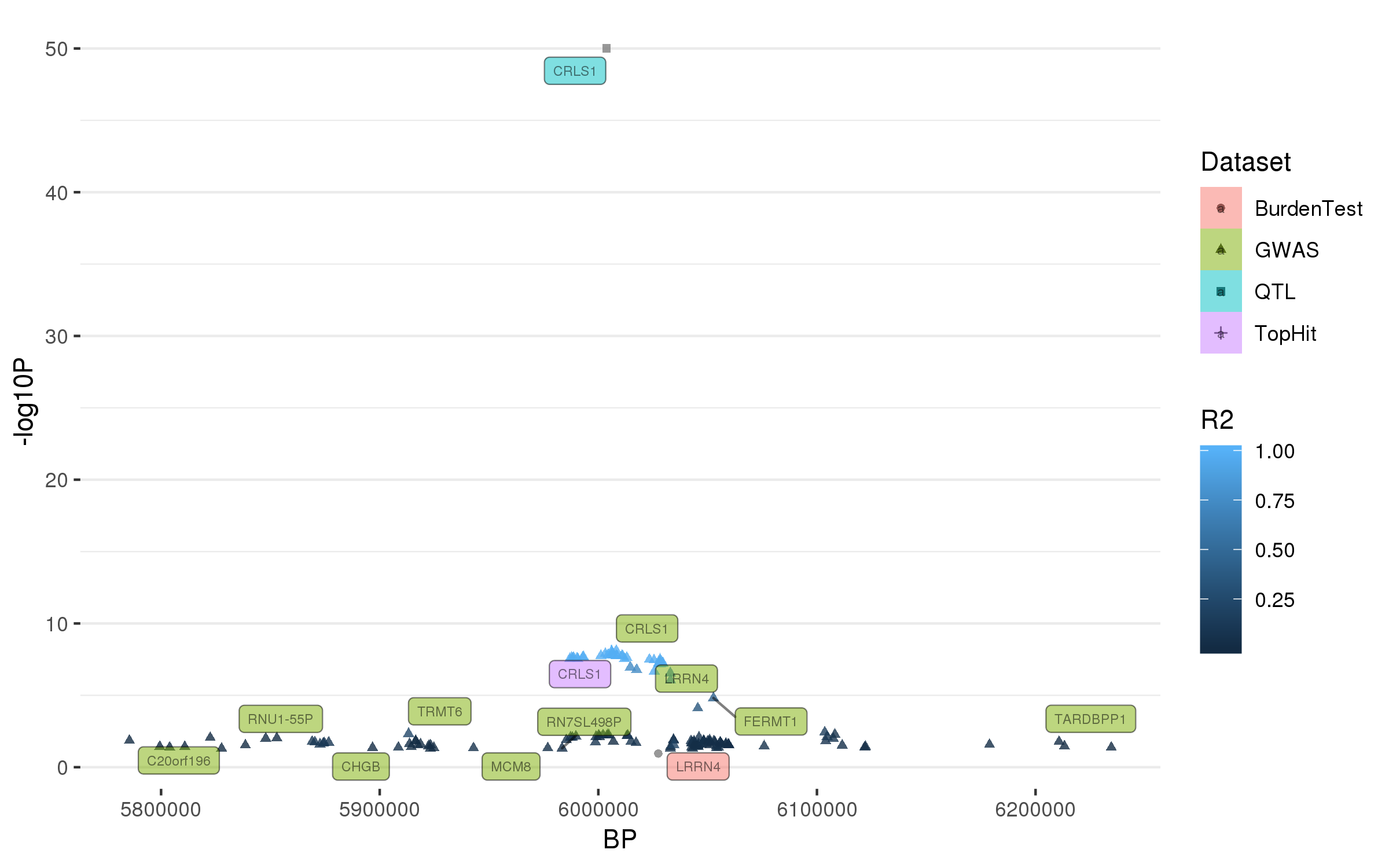

### rs112485576.png

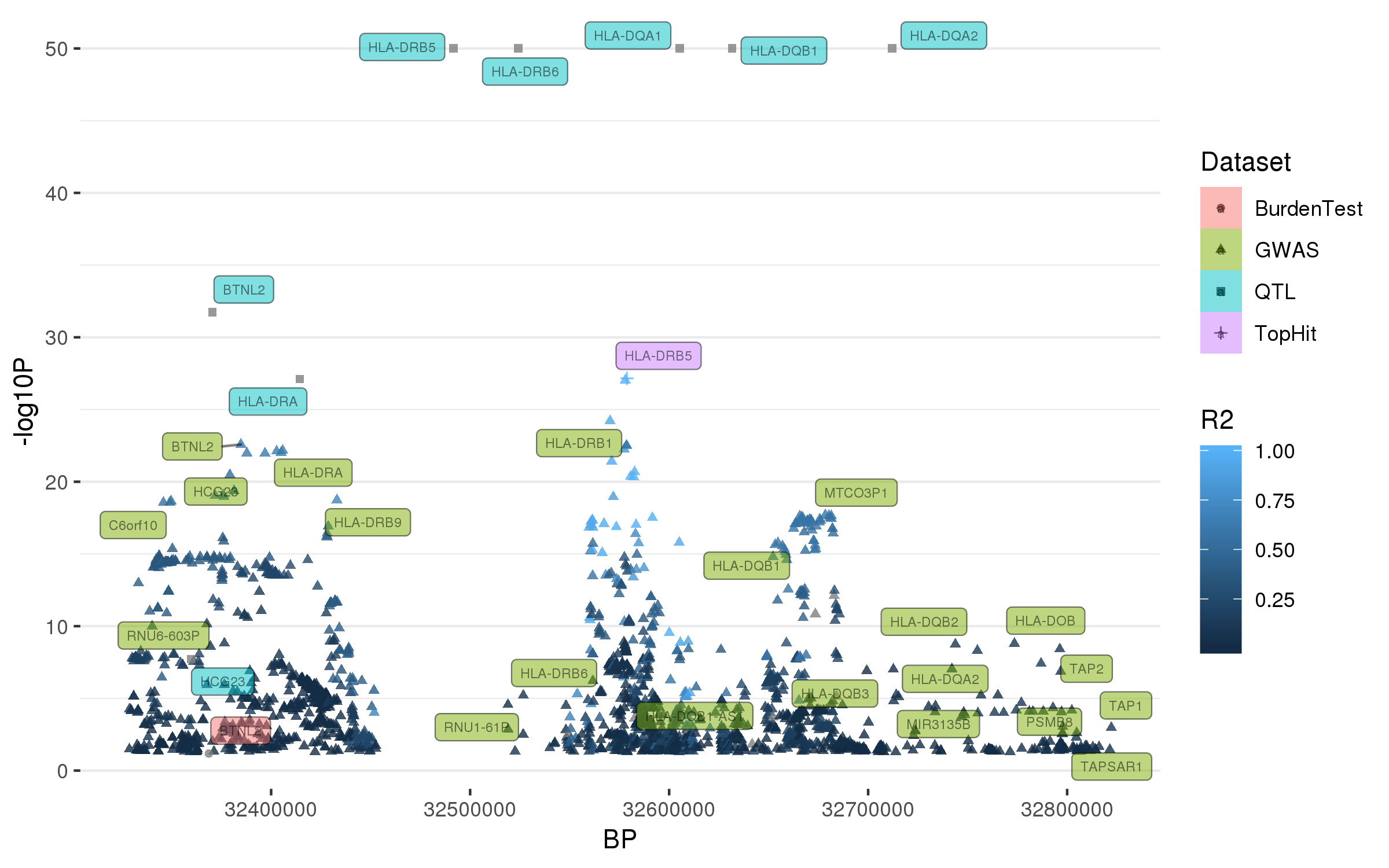

### rs114138760.png

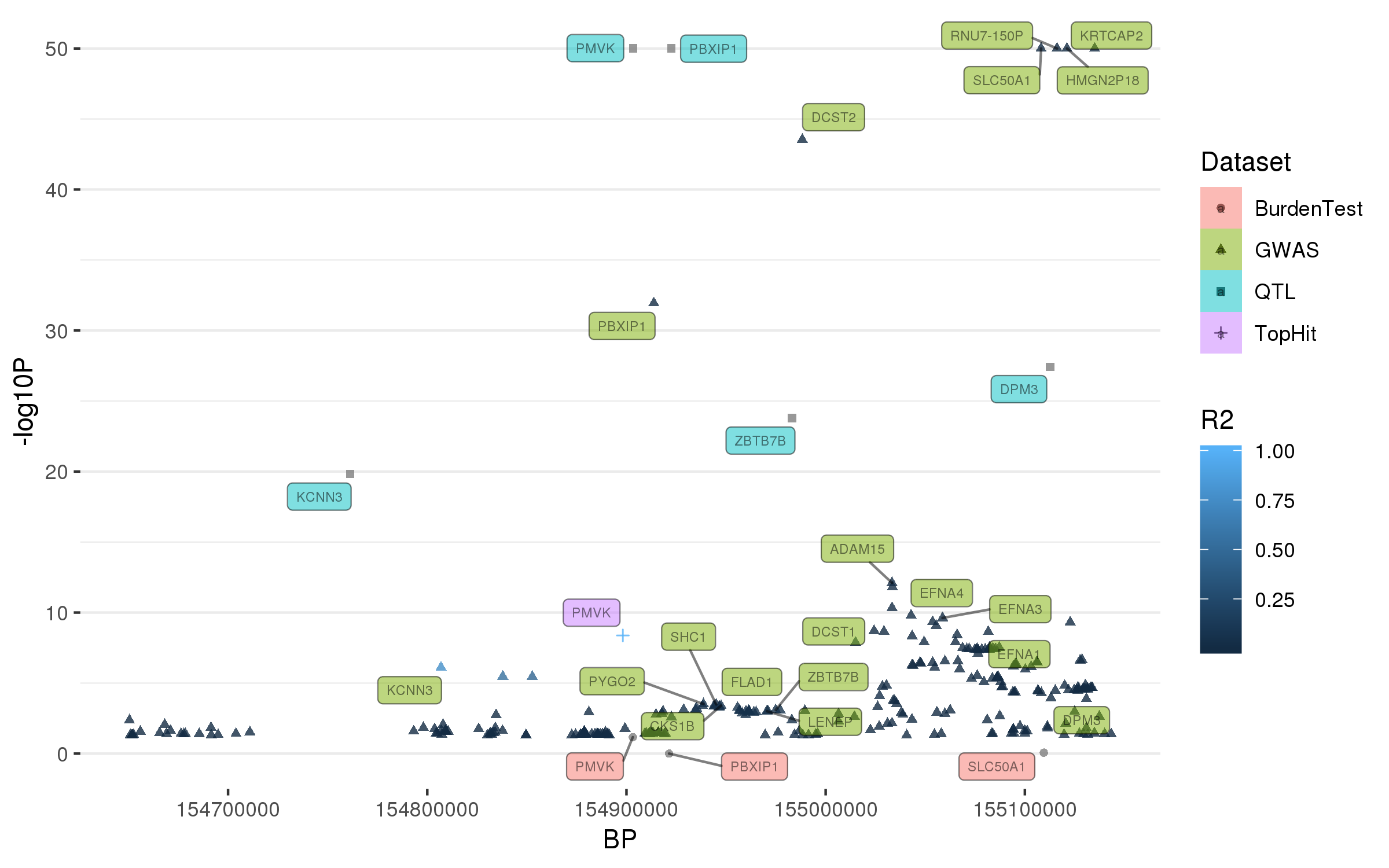

### rs117615688.png

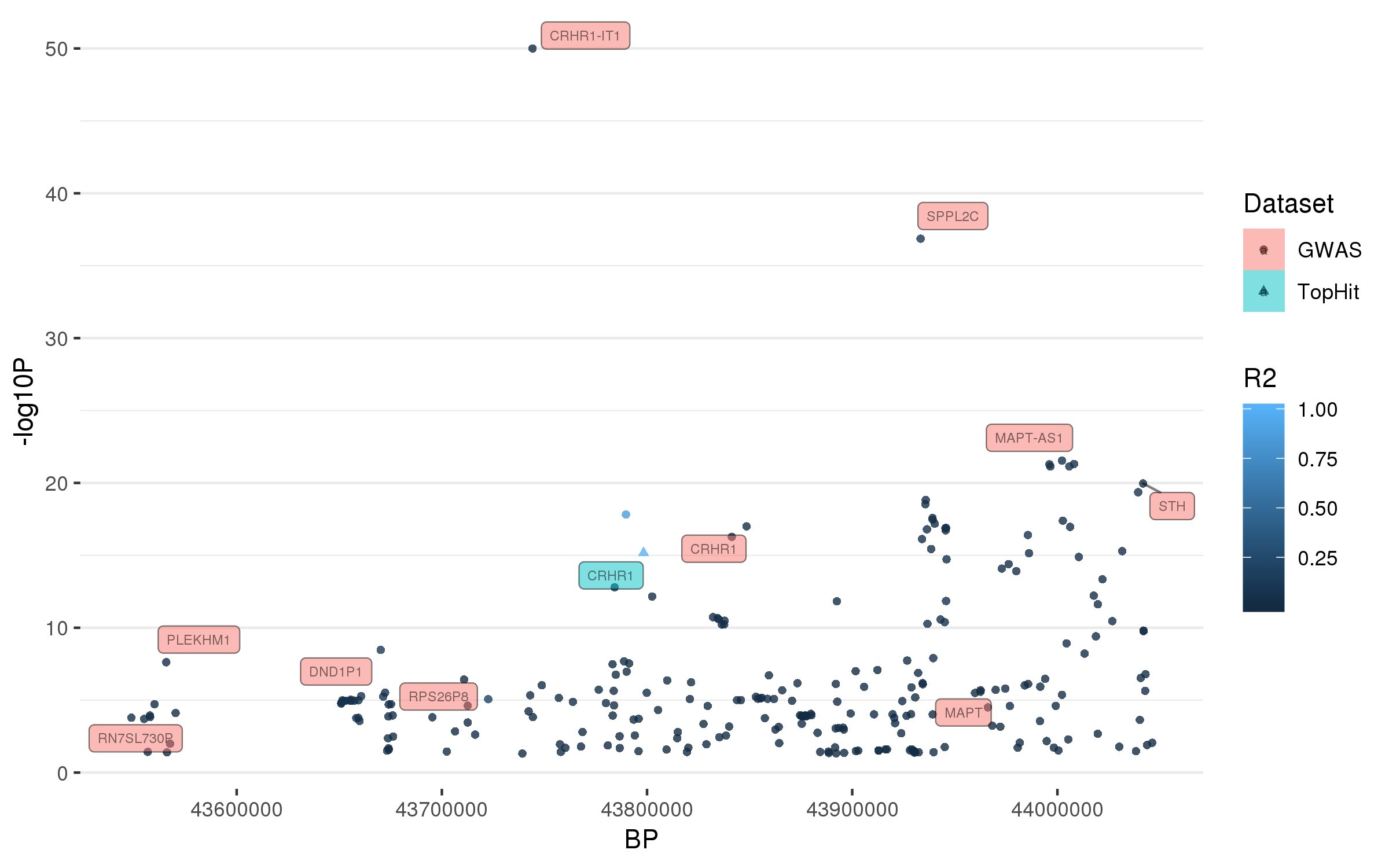

### rs117896735.png

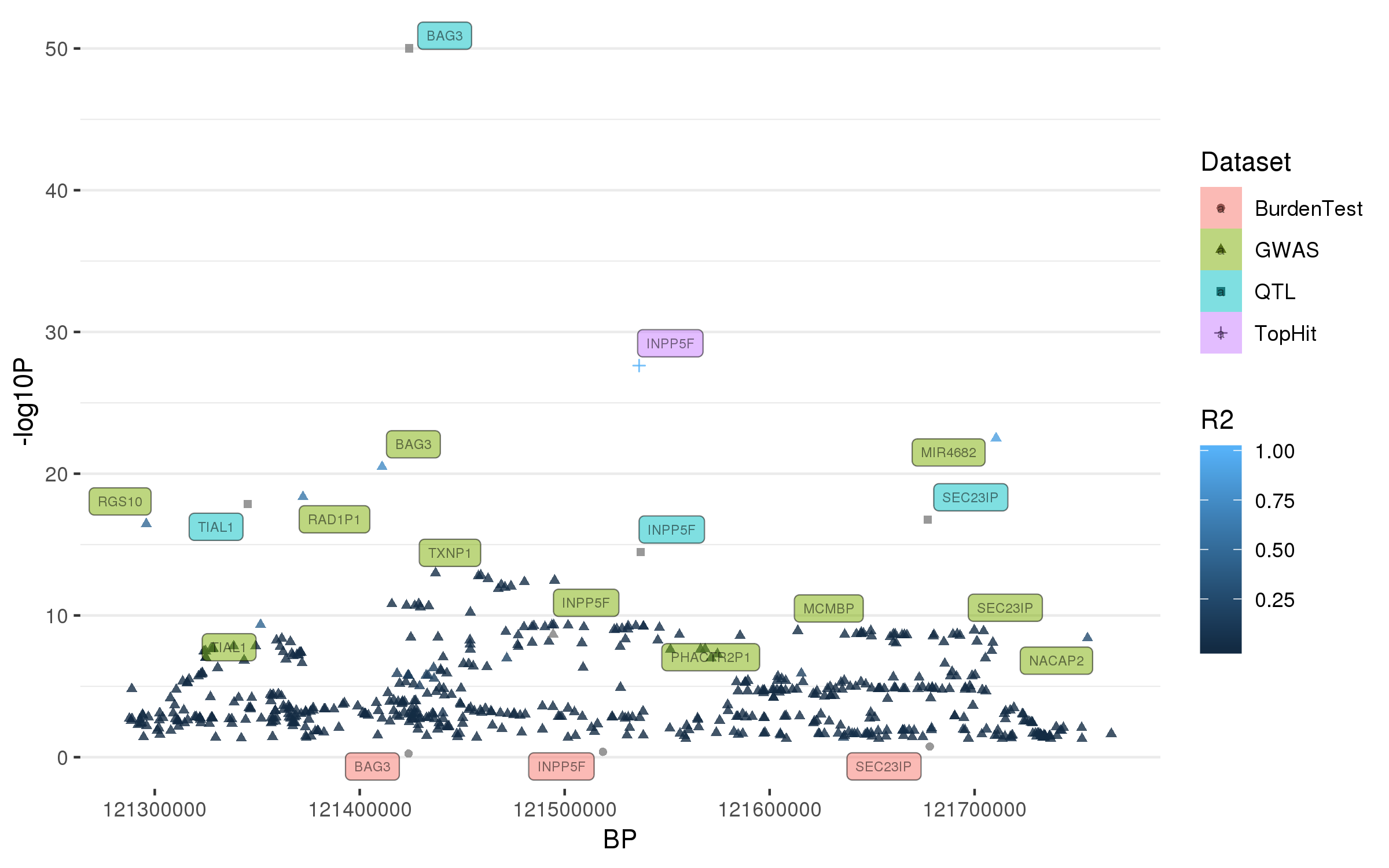
