## Supplementary material for "Expanding Parkinson’s disease genetics: novel risk loci, genomic context, causal insights and heritable risk": Online Supplementary Methods

**Samples and quality control**

Three primary sources of data were used for discovery analyses, these include three previously published studies, 13 new datasets, and proxy-case data from the UK BioBank (UKB). Previous studies include summary statistics from the meta-analyses of GWAS published in Nalls *et al*. 2014, GWAS summary statistics from the 23andMe Web-Based Study of Parkinson's Disease (PDWBS) cohort described in Chang *et al*. 2017, and the publicly available NeuroX dataset from the International Parkinson’s Disease Genomics Consortium (IPDGC) that had been used in the two prior publications as a replication sample. These cohorts have been reported in detail elsewhere ^1,2^. We also included 13 new case-control sample series for meta-analyses through either publicly available data or new collaborations (please see Supplementary Table S1 for details regarding these studies).

All new samples from the 13 new datasets underwent similar standardized quality control for inclusion, mirroring that of the previous studies. Initial sample inclusion criteria include: age at disease onset or last examination at 18 years of age or older, minimum sample call rate of >95%, majority European ancestry confirmed through principal-components, no genetically ascertained relation to other samples in the meta-analysis (proportional sharing at a maximum of 12.5%) at the cousin level or closer (except in the case of the System Genomics of Parkinson's Disease (SGPD) which utilized mixed modeling to account for related samples) and no heterozygosity outliers past +/- 15% (quantified by F estimates in PLINK)^3^ . Studies from IPDGC collaborators (including previously published series) and those in publicly available databases were checked for relatedness and duplicates using identity-by-descent filtering to remove related samples both within and across datasets contributing to this effort (also using PLINK to generate PI_HAT estimates of relatedness and excluding samples with > 12.5% PI_HAT). All publicly available datasets and newly genotyped IPDGC datasets were clustered to identify cryptically related samples between each other as well as the UKB using similar methods in PLINK. These studies screened for relatedness at the National Institute on Aging site include: Baylor College of Medicine / University of Maryland, Finnish Parkinson's, Harvard Biomarker Study (HBS), McGill Parkinson's, Oslo Parkinson's Disease Study, Parkinson's Disease Biomarker's Program (PDBP), Parkinson's Progression Markers Initiative (PPMI), Spanish Parkinson's (from IPDGC), Tubingen Parkinson's Disease cohort (CouragePD), Vance (dbGap phs000394), UK PDMED (CouragePD), UK BioBank (UKB), IPDGC (Nalls et al. 2014 discovery phase) and NeuroX - dbGaP (phs000918.v1.p1). If a sample overlapped in two studies, it was removed from the larger study. Samples derived from 23andMe were checked internally for relatedness both among themselves and publicly available PD datasets using similar methods as previously described in earlier publications before being incorporated into this analysis (studies included in Nalls et al 2014, as well as the NeuroX-dbGaP, HBS, PPMI, PDBP, UKB) ^1,2^. There is a low likelihood for overlap between the SGPD and IPDGC samples due to geographic differences in sample collection (Australia versus Europe and North America). This is summarized in Supplementary Table S1.

For case diagnosis, all of the new cases, except for the post-Chang *et al.* dataset from 23andMe, conformed to the generally used criteria of a clinic visit and standard UK Brain Bank criteria with a modification to allow the inclusion of cases that had a family history of PD. In the most recent dataset from 23andMe (post-Chang *et al*.) cases were ascertained via the criteria of self-report of diagnosis, similar to previous publications ^1,2,4^. Status as “post-Chang” is denoted by enrollment as a case at a time after the analysis for the PDWBS closed. Post-Chang 23andMe samples were imputed using a combination of Finch for phasing (an in-house developed fork of Beagle) and miniMac2 for imputation with all-ethnicity samples from the September 2013 release of 1000 Genomes Phase1 as reference haplotypes^5–7^. Both the Finnish Parkinson's study and post-Chang sample series used population controls to some degree, with inclusion criteria of no self-report of neurological disease, memory loss, tremor or family history of PD when available. UKB proxy-cases were defined as the report of a first degree relative with PD and no International Classification of Diseases (ICD-10) or self-report of actual PD. UKB cases with self-reported PD were excluded, this includes 727 cases that self-reported PD without family history to keep the GWAX analysis homogenous and parsimonious. UKB controls were free of first degree family history of PD or PD by self-report or ICD-10 designation. No censoring was made on parent age in the UKB. Additional summary statistics were generated for the UKB censoring on those who answered “Do not know” or “Prefer not to answer” to any of the possible illnesses. This reduced control counts down to 369,711. The reduced UKB results were highly correlated with the larger set included in all analyses (rG = 0.9865, SE = 0.0011), so we opted to utilize the results including more samples. All participants donated DNA samples and provided informed consent for participation in genetics studies.

Prior to imputation, SNPs were filtered using similarly uniform criteria for inclusion, such as: minimum genotype call rate of >95%, a minor allele frequency (MAF) of >0.1%, a Hardy-Weinberg equilibrium P values >1 E-04 in controls (1% MAF and HWE p<1E-06 for SGPD), and non-random missingness by phenotype or haplotype at P values of >1E-04. Palindromic SNPs were also removed. The Nalls *et al*. 2014 and Chang *et al*. 2017 samples were imputed with Minimac2 using 1000 Genomes phase 1 haplotypes ^7^. All additional sample series except for the post-Chang *et al*. 2017 samples from 23andMe were imputed using the Haplotype Reference Consortium (HRC) on the University of Michigan imputation server under default settings with Eagle v2.3 phasing based on reference panel HRC r1.1 2016^8,9^. UKB genotype data for proxy-cases and controls was downloaded in April 2018 as provided by the analysis group at the Wellcome Trust Centre for Human Genetics at the University of Oxford and is fully detailed at [http://biobank.ctsu.ox.ac.uk](http://biobank.ctsu.ox.ac.uk/)^10^.

**Study-level analyses and meta-analyses**

Summary statistics for each study were generated uniformly using additive allele dosages from imputation in a logistic regression framework adjusting for age at onset in cases (age at most recent examination for controls), biological sex and up to the first five principal components from stepwise modeling per cohort. The SGPD study did not use logistic regression, as an inclusion of related samples required mixed modeling. The Finnish Parkinson's study was unable to include age as a covariate due to collinearity issues as the controls were significantly older than the cases. Age at onset or last exam data was not available for the UK PDMED dataset. Age at last exam was used for the Oslo Parkinson's Disease Study dataset.

UKB data was analyzed slightly differently from standard case-control GWAS because phenotypes were available on relatives of the individuals with genotypes. Genome-wide association study by proxy (GWAX) was carried out as per Liu ^11^, adjusting for age, sex, the first ten principal components, genotyping batch and Townsend index^12^. GWAX is a analysis method shown to carry out reliable and generalizable association analyses on biobank / population scale data utilizing at-risk samples instead of true cases. Summary statistics were also filtered at a MAF > 1% for inclusion. Per SNP effect estimates were transformed using the method from Lloyd-Jones et al. assuming the lifetime probability of disease (K) at 0.02 and then converted to standard case-control scale (with the total proxy cases equivalent to ~4.5K cases) ^13^.

Similar to previous publications, study-level summary statistics were filtered for inclusion criteria of reasonable beta estimates imputation quality score > 0.3 and MAF > 1% ^1,2,14^. We retained 2 variants for study below the MAF minumum of 1%, these include known coding risk factors include rs34637584 (LRRK2, p.G2019S) and rs76763715 (GBA, p.N370S). Our GWAS meta-analyses spanned all three data sources (previously published case-control datasets, new case-control datasets and proxy-case data from the UKB) including 37,688 cases, 18,618 proxy-cases and 1,474,097 controls with data for 7,784,415 SNPs that passed inclusion filtering. Inclusion filtering included variants seen in at least five of the 17 sets of summary statistics and differences in the per variant minimum and maximum MAFs across all studies at greater than the 99th percentile (15% frequency differences). Four of the studies involved in our meta-analysis were genotyped using the NeuroX array, which is enriched for PD risk loci (see Supplementary Table S1) ^15^. The motivation for choosing five out of 17 studies was to reduce potential bias due to targeted genotyping in the four NeuroX array studies.

Overall genomic inflation was minimal with a raw lambda estimate of 1.170 and a lambda scaled to 1,000 cases and 1,000 controls at 1.002^15–17^. Well behaved lambda estimates in conjunction with an LD score intercept of 0.991 and a number of well replicated risk loci spanning larger tracts of the genome (i.e. *MAPT* and *HLA/MHC*), led us to not implement genomic control on the meta-analysis level. All quality control metrics on a per study basis including lambda estimates across strata of minor allele frequencies are summarized in Supplementary Table S1.

As previously stated, we did not correct for genomic control. Both the previous datasets and new datasets exhibited LD score regression intercepts close to 1, with 0.988 for the previous datasets and 0.975 for the new datasets, suggesting that our results are unlikely to be due to population stratification ^18^. As detailed in Supplementary Table S1, scaled lambdas were acceptable across the allele frequency spectrum with minor inflations for studies using genotyping arrays with targeted PD-centric content.

We also queried the 17 novel variants of interest from Chang et al., 2017 for their inclusion under GWAS peaks from this report^2^. We identified proxy variants tagging genome-wide significant peaks in this report (Supplementary Tables S2). Pairwise linkage disequilibrium was calculated from the European subset of the 1000 genomes data. Proxies are summarized in Supplementary Table S3. Four loci are directly validated using the same SNPs, 6 are tagged by a strong proxy (r^2^ > 0.8), 4 tagged by a weaker proxy (0.2 < r^2^ < 0.5) and an additional 3 with no genome-wide significant proxy within 500kb observed (rs353116, rs143918452 and rs78738012). One missing variant, rs78738012, is tagged under one of our peaks of interest by a SNP not passing genome-wide significance near the gene *CAMK2D*. The two untagged variants are likely due to expanding our analysis past genotyped SNPs from the NeuroX array and utilizing updated imputation references.

**Conditional-joint analysis to nominate variants of interest**

To nominate variants of interest, we employed a conditional and joint analysis strategy (GCTA-COJO, http://cnsgenomics.com/software/gcta/) as a means to algorithmically identify variants that best account for the heritable variation within and across nearby loci^19^. This is particularly useful in scenarios where only basic summary statistics are available for a majority of samples in a meta-analysis and additional participant level analyses are logistically prohibitive. For this analysis, we used the full meta-analysis summary statistics in conjunction with the largest single site collection of HRC-level imputed PD and control data as a reference for linkage disequilibrium patterns in the conditional-joint workflow (described below).

Using the consensus IPDGC data cleaning and imputation workflow, we assembled a reference set of 17,188 cases and 22,875 controls at variants overlapping with the locus discovery analysis results that passed quality control on average in 74.3% of samples incorporating soft call genotypes at a minimum imputation quality of 0.30. This set includes data from all samples described in Supplementary Table S1, except the 23andMe post-Chang *et al.*, System Genomics of Parkinson's Disease (SGPD), UK PDMED and UKB sample series. This aggregate dataset included previously described samples series such as the Dutch GWAS, German, UK and US IPDGC series plus the NeuroX-dbGaP and Myers-Faroud datasets from dbGaP^15,20,21^. The COJO analysis was run using default analysis parameters including a significance threshold of P < 5E-8 and a window specification of 1 megabase. Additional analyses described below were utilized to further scrutinize putative associated variants and account for possible differential LD signatures in multiple ways, including utilizing the massive single site reference data from 23andMe in further conditional analyses. If a variant nominated during the COJO phase of analysis was greater than 1 megabase from any of the genome-wide significant loci nominated in Chang *et al*. 2017, we considered this to be a novel locus.

**Additional filtering of nominated variants**

We instituted two additional filters after fixed-effects and COJO analyses. These additional filters exclude variants that 1) had a random-effects *P* value across all datasets > 4.67E-04 and 2) a conditional analysis P > 4.67E-04 using participant level 23andMe data. This multiple testing threshold is based on up to 107 nominated variants at this stage of filtering, of which 90 passed these criteria. Random-effects meta-analysis P-values were generated under the residual maximum likelihood method using the R package metafor^22^. Conditional analyses were carried out using 23andMe pooled data analyses including all available 23andMe data (from Nalls et al. 2014, PDWBS and the post-Chang et al. 2017 datasets combined). For the participant level conditional analyses in 23andMe, all nominated variants per chromosome were included in a single logistic regression model with appropriate covariates, then parameter estimates per variant were extracted. Conditional analyses on a per chromosome interval instead of a locus or megabase interval should adjust for possible longer range LD associations. For more information on variant filtering, please see Supplementary Table S2 summarizing all variants nominated. We nominated variants as sharing a single locus if they are within +/- 250kb of each other.

**Refining heritability estimates and determining extant genetic risk**

The primary tool for risk profiling used here was the R package PRSice2^23^. This package carries out polygenic risk score (PRS) profiling in the standard weighted allele dose manner as we have previously described ^24–30^. In addition, PRSice incorporates permutation testing where case and control labels are swapped in the withheld samples to generate an empirical P. This workflow also uses P-value aware LD pruning to facilitate identifying the best P thresholds for variant inclusion into the PRS below what is commonly regarded as genome-wide significance levels. PRS analyses were conducted in sample series with readily accessible participant level IPDGC GWAS data at the NIH site.

A two stage design was also employed, training on the largest single array study (NeuroX-dbGaP) and then tested on the second largest similar study (HBS). These two targeted array studies were chosen for three reasons: precedent in the previous publications where the NeuroX-dbGaP dataset was used in PRS comparisons; direct genotyping of larger effect rare variants in GBA and LRRK2; participant level genotypes for these datasets are publicly available.

To facilitate risk profiling analyses with less bias, we regenerated the meta-analysis summary statistics excluding the NeuroX-dbGaP and HBS datasets. To select SNPs we ran the PRSice workflow, utilizing beta weights from the meta-analyses excluding our studies of interest. LD clumping was carried out under recommended default settings (window size = 250kb, r^2^ > 0.1). Next 10,000 permutations were used to generate empirical P estimates for each GWAS derived P threshold ranging from 5E-08 to 1E-04, by increments of 5E-08, then again with GWAS derived P thresholds from 1E-04 to 0.5 by increments of 1E-04. For each iteration of the permutation tests in the training dataset, Nagelkerke’s pseudo r^2^ estimates between the PRS and PD were estimated, after adjustment for an estimated prevalence of 0.5% and study-specific eigenvectors 1-5, age and sex as covariates; this prevalence was chosen as a conservative estimate based on global estimates of disease ^31^. Then the summary statistics for these SNPs of interest (1805 overlapping with the HBS dataset after QC) were extracted from a the meta-analysis excluding HBS to generate variant weights for the validation phase of analysis. Next the PRS was tested in the HBS dataset. After this, we also reduced the PRS SNPs to just 90 variants reported as independent GWAS risk variants. We then repeated this workflow using the 88 SNPs passing QC in HBS as an additional test.

Areas under the curve and related metrics for predictive models based on the PRS were generated by combining the predicted case status probability estimates across datasets. To calculate heritability in clinically defined PD datasets, we also used LD score regression under default settings, also employing the LD references for European provided with the software^18^. This workflow was also repeated on a per cohort level and is detailed in the Supplementary Appendix.

**Mendelian randomization to infer functional consequences**

We used MR to test whether changes in methylation and RNA expression of genes physically proximal to genome-wide significant PD risk loci were causally related to PD risk. To nominate genes of interest for MR analyses, we took our putative 90 loci of interest in the large LD reference used for the COJO phase of analysis and identified SNPs in LD with our SNPs of interest at an r^2^ > 0.5 within +/- 1MB (Supplementary Table S5). Once these SNPs were identified, nearest genes were queried from the European Bioinformatics Institute (EMBL-EBI, https://www.ebi.ac.uk) and compiled into a list of 305 possible genes linked to PD risk loci.

MR was used to make functional inferences by integrating discovery phase summary statistics with quantitative trait locus (QTL) association summary statistics across well-curated methylation and expression datasets. We utilized the curated versions of Qi et al., 2018 brain methylation and expression summary statistics, as well as a specific focus on GTEx substantia nigra data, we also made use of the blood expression data from Võsa et al. 2018, all available from the website for summary-data-based Mendelian randomization (SMR, <http://cnsgenomics.com/software/smr/#Overview>) or the eQTLgen consortium (<http://www.eqtlgen.org>) ^32–36^. For all QTL analyses, we utilized the multi-SNP SMR method under default analysis settings. In the analyses using SMR, the large LD reference set from our COJO phase of analysis was used. For each of the four QTL datasets, false discovery rate was used to adjust P-values and account for multiple testing within each dataset. All MR effect estimates are reported on the scale of a standard deviation increase in the exposure variable relating to a similar change in PD risk.

**Rare coding variant burden tests**

A uniformly quality controlled and imputed dataset from the IPDGC (described above) was used to carry out burden tests for all rarer coding variants successfully imputed in an average of 85% of the sample series (17,188 cases and 22,875 controls). These analyses include all variants at a hard call threshold of imputation quality > 0.8. After annotation with annovar, we had a total of 37,503 exonic coding variants (nonsynonymous, stop or splicing) at MAF < 5% and a subset of 29,016 at MAF < 1%^37^. We then extracted proximal genes for SNPs tagging any of our 90 loci using same the LD reference as in the COJO analysis (r^2^ > 0.5 within +/- 1MB, see Supplementary Table S5). For inclusion in this phase of analyses, a gene must have contained at least 2 coding variants. After assembling this subset of 113 testable genes, we used the optimized sequence kernel association test to generate summary statistics at maximum MAFs of 1% and 5%^38^. All burden analyses were adjusted for the first 15 principal components based on common unlinked variants, age, sex and study site. Resulting P values were then adjusted via Bonferroni.

**Network analyses**

Two network-based approaches were utilized to assess connectivity across loci. The first focuses on integrating GWAS summary statistics with expression data (Functional Mapping and Annotation of Genome-Wide Association Studies, FUMA), the second focuses on protein interactions (webgestaltR).

Functional mapping and annotation based on publicly available gene expression and ontology resources were made using FUMA version 1.3.1 ^39^. In brief, summary statistics were analyzed using MAGMA gene property tests to compare enrichment of the average gene expression per tissue in GTEx v7 ^40,41^. Bonferroni correction was applied to tissue enrichment analyses. In total 10,651 gene sets (Curated gene sets: 4734, GO terms: 5917) were tested. Curated gene sets were generated from nine data resources including KEGG, Reactome and BioCarta (see MSigDB for details, http://software.broadinstitute.org/gsea/msigdb/collections.jsp)^42^. GO terms comprised of the three standard categories, biological processes (bp), cellular components (cc) and molecular functions (mf). All parameters were set as default for the competitive test. We employed a more conservative version of the FDR correction for multiple testing by applying it to all pathways from various sources at once. Single cell RNA sequencing data from DropViz (<http://dropviz.org>) was also queried for enrichment using FUMA in an identical manner, spanning 88 possible tissue and cell type combinations ^43^.

To investigate protein components related to genetic risk loci, we utilized the R package webgestaltR to build networks via its network topology analysis and random walk algorithm ^44^. The input data for this analysis was all genes found under the association peaks in our GWAS based on the LD structure in reference samples as described in the previous subsection. Ontologies were extracted from the Biogrid Protein-Protein Interaction Networks^45^. False discovery rate adjustment was used to adjust for multiple testing.

**LD score regression**

To investigate shared genetic correlations of PD with multiple traits and diseases, we employed bivariate LD score regression ^18^. These analyses were carried out under the default settings as previously discussed, using data from the 757 GWAS available via LD Hub as well as biomarker GWAS summary statistics from two additional publications of interest focusing on c-reactive protein and cytokine measures; LD Hub was accessed on June 20th, 2018 (version 1.2.0)^46–48^. P-values from the bivariate LD score regressions were adjusted for false discovery rate to account for multiple testing. We acknowledge that for some traits of interest there is a minor overlap with samples derived from the CHARGE studies utilized in a small portion of discovery data in Nalls et al. 2014 which may influence results slightly in downstream MR analyses ^1^. For evaluation of genetic correlations between PD and UKB derived GWAS, we utilized PD summary statistics with the UKB data excluded to reduce bias.

**Causal Inference**

Traits showing significant genetic correlations with PD were analyzed using MR methods. We excluded the UKB data when a nominated trait was from summary statistics derived from the UKB or including the UKB in a meta-analysis.

When complete GWAS summary statistics were available for traits of interest (relating to smoking and education), we utilized the more powerful bi-directional generalized summary-data-based Mendelian Randomization (GSMR). This approach utilized bi-directional GSMR under default settings with the exception of more stringent HEIDI outlier removal options (global HEIDI threshold at 0.05)^49^. For all MR results (both here and below), effect estimates where PD is the outcome are interpreted as the change in the log odds ratio (beta) of PD for a single standard deviation increase in the polygenic risk score for the exposure.

Summary statistics for PD excluded UKB data at this stage. The previously described PD reference datasets for estimating LD was used, excluding samples with missing data for SNPs of interest. We analyzed GWAS summary statistics for smoking initialization (453,693 records from a self-report survey with 208,988 regular smokers and 244,705 never regular smokers) and current smoking within the UKB Current smoking (CS) contrasted 47,419 current smokers versus 244,705 never regular smokers. The same analysis was carried out incorporating recent GWAS data regarding educational attainment (N = 766,345) and cognitive performance (N = 257,828)^50^. These outcomes were analyzed using methods to mirror that of the UKB PD GWAS dataset. Combined left and right putamen volume from a T2 magnetic resonance imaging GWAS available from Oxford Brain Imaging Genetics (BIG) Server (accessed December 28th, 2018) ^51^.

**Additional sensitivity analyses**

Mirroring our previous workflows used in the initial meta-analysis, we conducted 17 “leave-one-out” meta-analyses (LOOMA), excluding one dataset’s summary statistics each time. We also carried out a similar set of meta-analyses separately for previously published data versus the combined set of 14 unpublished case-control and proxy-case datasets.

We employed a number of sensitivity analyses to further investigate our data and results as opposed to using a two-stage study design in order to maximize discovery power. LD score regression was utilized to calculate the genetic correlation between the meta-analysis of previously published datasets and the meta-analysis of the 14 unpublished datasets^18^. Additionally, all datasets described in Supplementary Table S1 were meta-analyzed using fixed-effects, stratified by diagnostic criteria of either self-reported PD (post-Chang and PDWBS) or clinically ascertained PD (all other datasets excluding data from the UKB and Nalls et al. 2014). Summary statistics stratified by diagnosis were then used to compare genetic correlations across clinically defined, self-reported and proxy-case derived datasets using LD score regression.

Due to the relatively small sample sizes of many of the datasets involved in this study, we could not run LD score regression on all combinations of LOOMAs. Instead, we opted to calculate linear regression models comparing the beta coefficients for the left-out dataset with the beta coefficients from a meta-analysis of the remaining (non-left out) datasets, stratified by all, novel and known PD risk loci.

Similar to above, we calculated random-effects meta-analysis P-values using the residual maximum likelihood method in the R package metafor. We also generated forest plots from this analysis that reflect the distributions of effect estimates across studies per SNP, summarized by the index of heterogeneity (I2) statistics in Table 1 and Supplementary Table S2. This statistic estimates possible variance accounted for by study heterogeneity.

Final sensitivity analyses included “leave-one-out” meta-analyses (LOOMA) comparisons of each dataset to a meta-analysis of the remaining datasets. This analysis focused on comparing the log odds ratios (termed beta here) per SNP identified in the GWAS analyses across all cohorts. After adjusting for multiple test correction for 17 tests (P < 0.003 for significance) in regressions of up to 90 betas per iteration, we noted only 5 departures from significant correlations between the withheld and included datasets. These non-significant results included only novel loci in the Baylor / University of Maryland dataset, the Finnish Parkinson’s dataset, the Harvard Biomarker Study (HBS), the Parkinson’s Disease Biomarkers Program (PDBP) and the Parkinson's Progression Markers Initiative (PPMI). For these five studies, correlations were significant in the known and all loci strata of variants. This is likely be related to statistical power for detecting recently identified risk variants in this subset of smaller studies. While there may be some caution in utilizing UKB proxy-cases, our data shows that the UKB data was significantly representative of other datasets, with high r^2^ estimates across novel (r^2^ = 0.714, 38 variants), known (r^2^ = 0.897, 47 variants) and all variants strata (r^2^ = 0.866, 85 variants) in the LOOMAs. We view these LOOMAs as a means of detecting an outlier study and estimating generalizability in the context of the 92 nominated variants. Forest plots included in the Supplemental Appendix compare each study on a per variant basis.
